# Automated Landmarking via Multiple Templates

**DOI:** 10.1101/2022.01.04.474967

**Authors:** Chi Zhang, Arthur Porto, Sara Rolfe, Altan Kocatulum, A. Murat Maga

## Abstract

Geometric morphometrics based on landmark data has been increasingly used in biomedical and biological research for quantifying complex phenotypes. However, manual landmarking can be laborious and subject to intra and interobserver errors. This has motivated the development of automated landmarking methods. We have recently introduced ALPACA (Automated Landmarking through Point Cloud Alignment and Correspondence), a fast method to automatically annotate landmarks via the use of a landmark template as a part of the SlicerMorph toolkit. Yet, using a single template may not consistently perform well for large study samples, especially when the sample consists of specimens with highly variable morphology, as it is common in evolutionary studies. In this study, we introduce a variation on our ALPACA pipeline that supports multiple specimen templates, which we call MALPACA. We show that MALPACA outperforms ALPACA consistently by testing on two different datasets. We also introduce a method of choosing the templates that can be used in conjunction with MALPACA, when no prior information is available. This K-means method uses an approximation of the total morphological variation in the dataset to suggest samples within the population to be used as landmark templates. While we advise investigators to pay careful attention to the template selection process in any of the template-based automated landmarking approaches, our analyses show that the introduced K-means based method of templates selection is better than randomly choosing the templates. In summary, MALPACA can accommodate larger morphological disparity commonly found in evolutionary studies with performance comparable to human observer.

## INTRODUCTION

Geometric morphometrics (GMM), a statistical approach for shape analysis based on landmark data, has been increasingly used in biological and biomedical fields to disentangle complex phenotypes (Adams et al. 2013; Rolfe et al. 2021b; Zelditch et al. 2012). Nevertheless, collecting landmarks manually can be laborious and time consuming (Aneja et al. 2015; Percival et al. 2019; Porto et al. 2021; Pui and Minoi 2019; Young and Maga 2015). Manual landmarking also inevitably lead to intra- and interobserver errors, which can even disrupt detecting biologically meaningful variations (Daboul et al. 2018; Percival et al. 2014; Porto et al. 2021; Robinson and Terhune 2017). Nowadays, combining landmark data collected by different researchers becomes increasingly common when working with big datasets, so proper error control becomes more urgent for ensuring accuracy, consistency, and reproducibility (Daboul et al. 2018; Fruciano 2016; Porto et al. 2021; Robinson and Terhune 2017).

To resolve the limitations in manual landmarking, researchers have developed various automated landmarking techniques based on image registration (Bromiley et al. 2014; Devine et al. 2020; Maga et al. 2017; Percival et al. 2019; Young and Maga 2015). However, these methods may not always be convenient for biologists to use because most of them require high-end hardware and knowledge of image-processing (Bromiley et al. 2014; Young and Maga 2015; Devine et al. 2020). We have recently introduced an automatic landmarking method, ALPACA (Automated Landmarking through Point cloud Alignment and Correspondence), as part of the SlicerMorph morphometrics toolkit (Porto et al. 2021; Rolfe et al. 2021b). For a detailed explanation of ALPACA and its underlying methods, we refer the readers to the cited papers. In summary, ALPACA is fast and lightweight because it uses sparse point clouds extracted from the original 3D surface models (Porto et al. 2021). These point clouds, while retaining sufficient geometric information, greatly reduce the computational burden so that ALPACA can efficiently run on any recent personal computer and does not require access to specialized hardware such as GPUs or high-performance computing cluster. ALPACA is free to download as part of the open-source SlicerMorph extension in the 3D Slicer software (Kikinis et al. 2014; Porto et al. 2021; Rolfe et al. 2021b).

One limitation of ALPACA, and most other registration-based methods, lies in the usage of single template for landmarking the entire sample (Porto et al. 2021). This is because the accuracy of automated landmarking depends on how well the registration algorithm optimizes the cost of global registration with local shape differences, which becomes more difficult as the template and target specimens become more different in form (Young and Maga 2015; Porto et al. 2021). This limitation is particular well-known in neuroimaging where template-based analysis had been the norm for the last two decades (Rohlfing et al. 2005; Iglesias and Sabuncu 2015; Young and Maga 2015; Doshi et al. 2016). And it is particularly difficult in evolutionary focused biological studies, where researchers frequently deal with highly variable samples from different species. Consequently, it can be difficult to use a single specimen to landmark a variable study sample while maintaining accuracy of landmarking.

If one template is insufficient for landmarking a highly variable study sample, a potential solution is to use multiple templates to have a more comprehensive representation of the whole study sample (Antonelli et al. 2019; Iglesias and Sabuncu 2015; Rohlfing et al. 2005; Young and Maga 2015; Schipaanboord et al. 2019). The multi-template approach has already been utilized in automated image segmentation to better capture variability of anatomical structures within a sample so that the segmentation of these structures can be more accurate (Antonelli et al. 2019; Doshi et al. 2016; Iglesias and Sabuncu 2015; Rohlfing et al. 2005; Schipaanboord et al. 2019; Wang and Yushkevich 2013). Young and Maga (2015) also adopted the multi-template method from automated segmentation for their image-based automatic landmarking research and have achieved an improvement in accuracy comparing to the single-template method. The main issue with the multi-template methods is the increased computational demand, as each subject now needs to be registered as many times as there are templates. While this is a concern for computationally expensive methods takings hours, in ALPACA a single specimen can be landmarked in a matter of few minutes, making ALPACA an effective tool to implement multi-template automated landmarking approach(Porto et al., 2021).

Overall, there are two steps to the MALPACA workflow: the first step, which is optional, is to identify the templates to be used to landmark the rest of the samples. The second step is the execution of the multi-template estimation pipeline, which is essentially running ALPACA independently for each unique template. The final output for the target specimen is the median of all corresponding ALPACA estimates.

Identifying the templates to sufficiently capture the variability of the whole sample are critical for any method that relies on them (Antonelli et al. 2019; Gooding 2021; Schipaanboord et al. 2019). This might be particularly difficult if there is no prior information, such as a pilot study with smaller samples sizes, is available to the investigator. If the investigator needs to manually landmark even a portion of the study sample to determine the morphological variability, and then choose the ones to be used as templates, this time investment may overcome the benefit from automated landmarking. If such a pilot dataset is already in existence, we advise investigators to make use of these priors for their template selection. For the cases where no prior information is available, we implemented a K-means based template selection that uses point-clouds of the surface models of the study population to approximate the morphological variability in an unbiased way. A Generalized Procrustes Analysis (GPA) is applied to these point clouds, followed by the PCA decomposition of Procrustes aligned coordinates. Using all PC scores from the GPA we apply K-means clustering to the data to detect the samples that are closest to the centroids of the identified clusters. The investigator can specify how many templates per group (if data consists of multiple groups or species) are going to be extracted. The investigator then landmarks the selected specimens, and inputs them as the templates for the MALPACA pipeline. More details of how to execute the K-means based template selection procedure can be found in the supplemental online material (Supplement_1).

The first goal of this study is to evaluate whether MALPACA outperforms ALPACA in estimating landmark positions with less error when compared to the “gold standard” (GS) manual landmarks. The second goal is, for cases where no prior information about landmark variability is available, to assess whether K-means can be an acceptable alternative for choosing a set of templates for MALPACA. The structure of the study is summarized in Table 1.

**TABLE 1.**
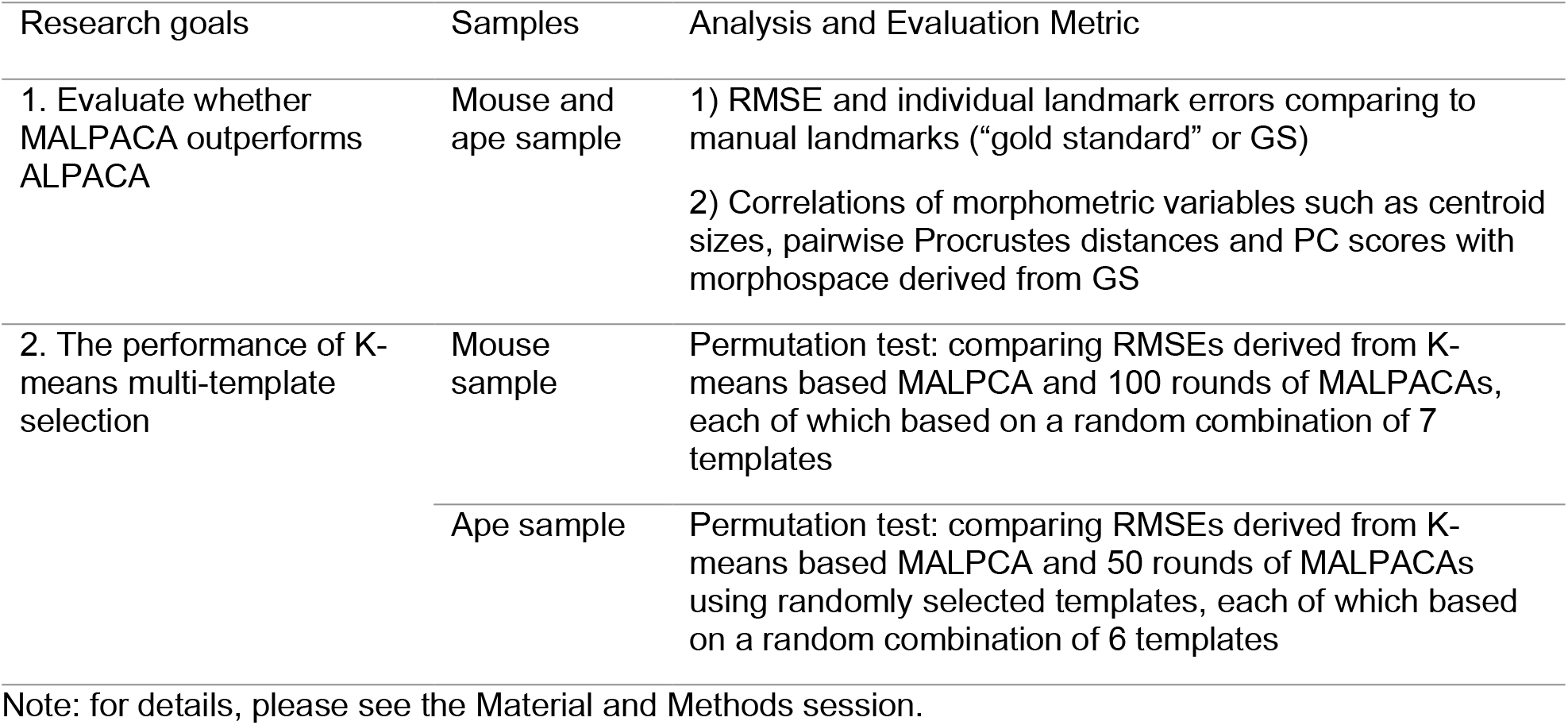
A summary of research goals of this study.

| Research goals | Samples | Analysis and Evaluation Metric |
| --- | --- | --- |
| 1. Evaluate whether MALPACA outperforms ALPACA | Mouse and ape sample | 1) RMSE and individual landmark errors comparing to manual landmarks (“gold standard” or GS)<br>2) Correlations of morphometric variables such as centroid sizes, pairwise Procrustes distances and PC scores with morphospace derived from GS |
| 2. The performance of K-means multi-template selection | Mouse sample | Permutation test: comparing RMSEs derived from K-means based MALPCA and 100 rounds of MALPACAs, each of which based on a random combination of 7 templates |
|  | Ape sample | Permutation test: comparing RMSEs derived from K-means based MALPCA and 50 rounds of MALPACAs using randomly selected templates, each of which based on a random combination of 6 templates |

Note: for details, please see the Material and Methods session.

## MATERIAL AND METHODS

### Samples and templates

Our study uses two samples that represent different types of group structure. The first sample consists of 3D skull models of one representative from 61 inbred laboratory mice from both classical and wild-derived strains (Maga et al. 2017; Porto et al. 2021) (See Tables S1 for list of strains used in the study). This dataset is used to mimic a population level morphometric analysis, where no distinct grouping structure exists (or expected) in the resultant morphospace. All specimens had been manually annotated with 51 landmarks once by single observer. Throughout this study we will refer the manually annotated datasets are as “gold standard” (GS). Seven templates are selected to landmark the remaining 54 specimens by MALPACA using the K-means template selection module. In addition, a synthetic template, an average mouse model, was the template used in the original ALPACA evaluation study (Porto et al., 2021).

The second sample contains 3D skull models of 52 great ape specimens from three species: 11 *Pan troglodytes*, 23 *Gorilla gorilla*, and 18 *Pongo pygmaeus* (For details, see Porto et al. 2021 and Rolfe et al. 2021a as well as Table S2 for list of specimens used in this study). The dataset is courtesy of the 3D Digitization program of the Smithsonian Institution. Each specimen has been annotated twice manually with 41 landmarks by a single observer. The mean of these two manual landmark sets is used as the GS. K-means template selection was used to identify two templates from each species to landmark the remaining 46 specimens (9 *Pan*, 21 *Gorilla*, and 16 *Pongo*).

### Methods

#### K-means-based multi-template selection

This study utilizes K-means clustering as the basis for templates selection (Young and Maga 2015). The raw data for K-means are the full set of principal component (PC) scores derived from the decomposition of GPA aligned coordinates from sparse point clouds. To achieve this, point-to-point correspondences must be established across point clouds so that they can be treated as landmark configurations. This is a three-step process:

1. A reference model is tightly registered to the target(s) using the global and ICP rigid registration steps from ALPACA (Rusinkiewicz and Levoy 2001; Rusu et al. 2009; Zhou et al. 2018; Porto et al. 2021).
2. A sparse point cloud is extracted from the reference. This is done by deleting points in the reference model until the distances between any pairs of points are not smaller than a user-defined threshold (Schroeder et al. 2006). In our sample, this leads reference point clouds with 791 and 674 points for mouse and ape datasets respectively.
3. For each point in the reference point cloud, the point closest to the target model is extracted (Schroeder et al. 2006). This results in a sparse point cloud from the target model that has point-to-point correspondence with the reference point cloud.

These point clouds are then treated as landmark configurations and submitted to a Generalized Procrustes Analysis (GPA). The shape coordinates achieved from the GPA of point clouds are submitted to a PCA. (Rolfe et al., 2021)

The full set of PC scores are input into the K-means algorithm to select a user-defined number (k) of templates. The K-means algorithm, which is implemented using the SciPy Python package, iteratively looks for an optimal way to partition specimens into k clusters so that the mean within-cluster Euclidean distance between specimens and the centroid is minimized (Virtanen et al. 2020). Within each cluster, the specimen closest to its centroid is selected as a template, so that k-templates are selected (Young and Maga 2015). It should be noted that for K-means template selection procedure to work correctly, 3D models of all specimens should be complete and segmented to have the same anatomical content. Because the workflow uses the entire model to generate point clouds, inclusion of incomplete specimens or extraneous anatomical elements will create spurious outcome. Step-by-step instruction for executing K-means template selection using the mouse data and landmarks are available as SOM (Supplement_2).

#### MALPACA

To initiate MALPACA user simply needs to input the location of 3D models and corresponding landmark set of specimens designated as templates. For each template, ALPACA is run independently on the target specimens to landmark them. (for details of ALPACA, see Porto et al. 2021). After a target specimen is landmarked by each template, its final landmark coordinate is estimated using the median of individual estimates of from templates. MALPACA module returns both the final median landmark estimate and estimates from individual templates as files on disk. While we chose median to estimate final landmark location due to its robustness to outliers, providing all the output gives the user flexibility to implement their own estimation method. Step-by-step instructions on how to execute the multi-template landmarking can be found in the SOM2 section 2.

### Evaluating MALPACA performance

#### Root Mean Square Error (RMSE)

To assess whether MALPACA outperforms ALPACA, we quantified how closely these two estimates approximate the GS landmark location. This is measured as the square root of the mean sum of squared errors (RMSEs) between GS and estimated landmark position Thus, RMSE serves as a single value to summarize the overall deviations between all estimated and GS landmarks for a specimen. While the RMSEs are calculated in the scale of the data (millimeters in this case), where appropriate we also report them as percentage of the centroid size of the specimen so that they can be understood in context of the body size. Because ALPACA returns estimated landmarks in target specimens coordinate system, error assessment does not require any further alignment or superimposition. Statistical significance of whether MALPACA-derived RMSEs are smaller than ALPACA-derived ones are assessed via one-sided Welch t-test.

#### Size and shape variables

To further evaluate the performance of MALPACA-estimated landmarks in morphometric analysis, we perform separate GPAs for MALPACA estimates and GS landmarks. We then calculate correlations between estimated and GS landmarks in pairwise Procrustes distances, principal components (PCs), and centroid sizes are also assessed. Procrustes distances quantify overall shape differences. PCs are components ranked by proportions of the total variance they explain. Centroid sizes are common size measures. Morphometric analyses commonly use these variables to represent overall shape and size variations (Zelditch et al. 2012). Joint GPA between MALPACA and GS landmarks are also carried out to visualize differences between estimated mean shapes.

#### Assessing the magnitude of digitation errors in MALPACA

As indicated above, both intra and interobserver errors are common in manual landmarking. Thus, looking at the RMSE of MALPACA errors is not sufficient, as these RMSE need to be evaluated in context of manual digitization errors. Our mouse sample was only landmarked a single time, as such there was no possibility of calculating observer errors. Instead, we relied on intraobserver errors computed by Percival et al.’s (2019) as the reference for manual errors as thirty-four of the landmarks in their study are also replicated in our analysis. As for the ape sample, each specimen was landmarked twice by the same observer. The RMSEs between these two manual landmark sets are used as an estimate of intraobserver errors. It should be noted that intraobserver errors tend to be much less than interobserver errors associated with manual landmarking (Percival et al., 2014).

#### Robustness of K-means based template selection

This study uses permutation analysis to evaluate whether K-means is an acceptable method to select templates when no prior information is available. Each permutation is based on using a random choice of specimens as templates to run a MALPACA (100 permutations for the mouse sample, and 50 permutations for the ape sample). We then used the RMSE from our K-means based MALPACA method and compared them to all permutated RMSEs using randomly selected templates (5,400 RMSEs for the mouse sample, and 2,300 RMSEs for the ape sample).

The evaluation of MALPACA performance is carried out in R (R Core Team 2021). RMSEs of the data are calculated using the SlicerMorphR package (https://github.com/SlicerMorph/SlicerMorphR). Generalized Procrustes Analysis (GPA) is performed using the geomorph R package for calculating correlations between estimated and manually placed landmarks in centroid sizes, pairwise Procrustes distances, and principal component scores (Adams et al. 2021).

## RESULTS

All MALPACA and ALPACA analyses were run on the same 64-bit Windows 10 desktop with an Intel Core i5 3.10 GHz CPU, and 16 GB RAM. For mouse dataset execution time was 69s for a single sample or 7.2h for all study which included the template selection.

### MALPACA performance over ALPACA

The analysis of the mouse sample shows that MALPACA outperforms each ALPACA run by estimating landmarks significantly closer to the GS (Fig. 3). The mouse MALPACA yields significantly smaller RMSEs than those generated by any ALPACA based on a K-means selected template or the synthetic template (all p-values from one-sided Welch t-tests < 8.52 × 10^−7^) (Fig. 3; Fig. S2;

**FIGURE 1.**
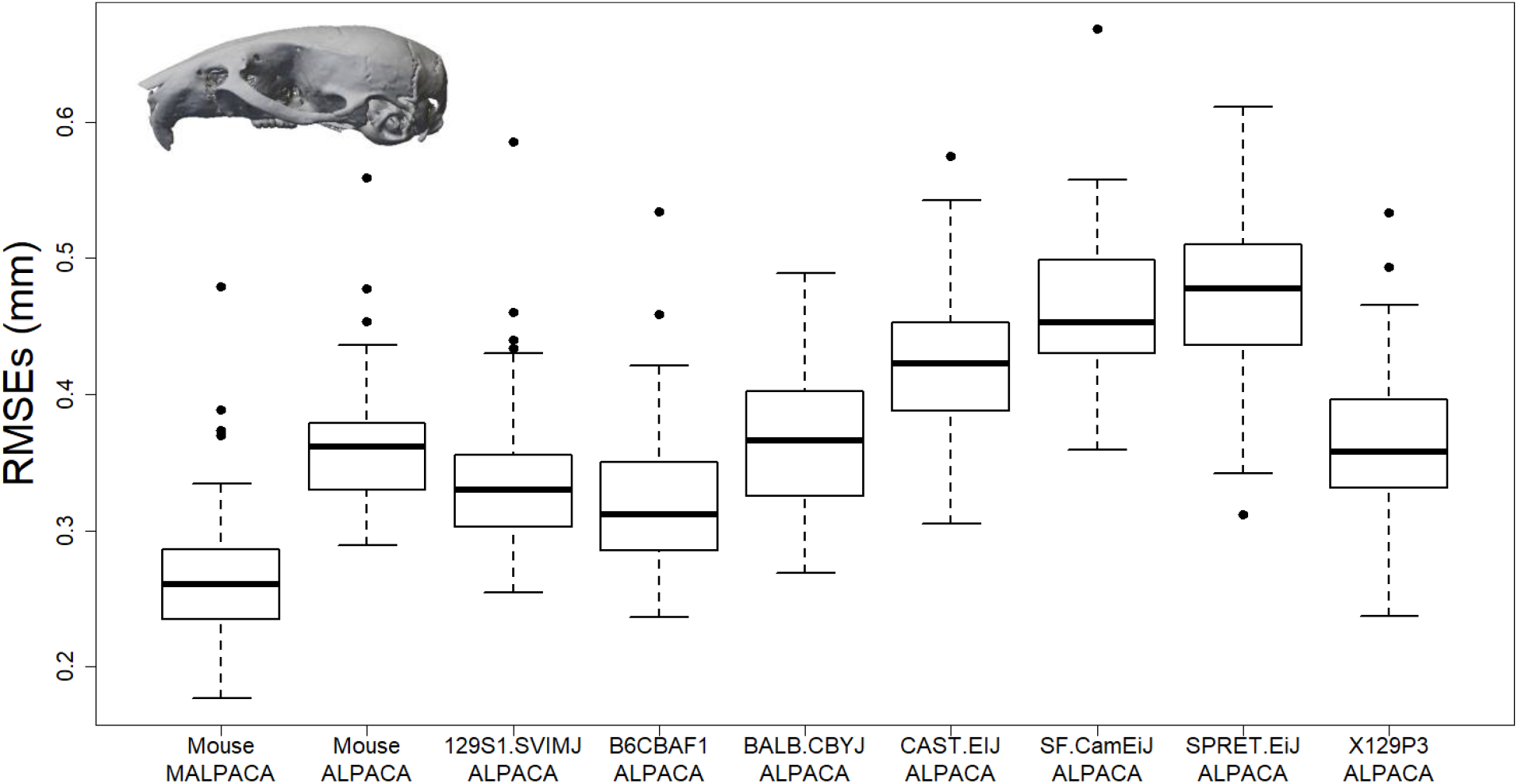
Comparison of RMSEs between estimated landmarks and the “Gold Standard” for **the mouse sample**. “ALPACA”: Mouse ALPACA is estimated using the synthetic mouse template used in the original ALPACA paper. Other boxes are ALPACA estimates using specified template. To see RMSEs expressed as percentage of specimen centroid sizes, please see Fig. S2.

**FIGURE 2.**
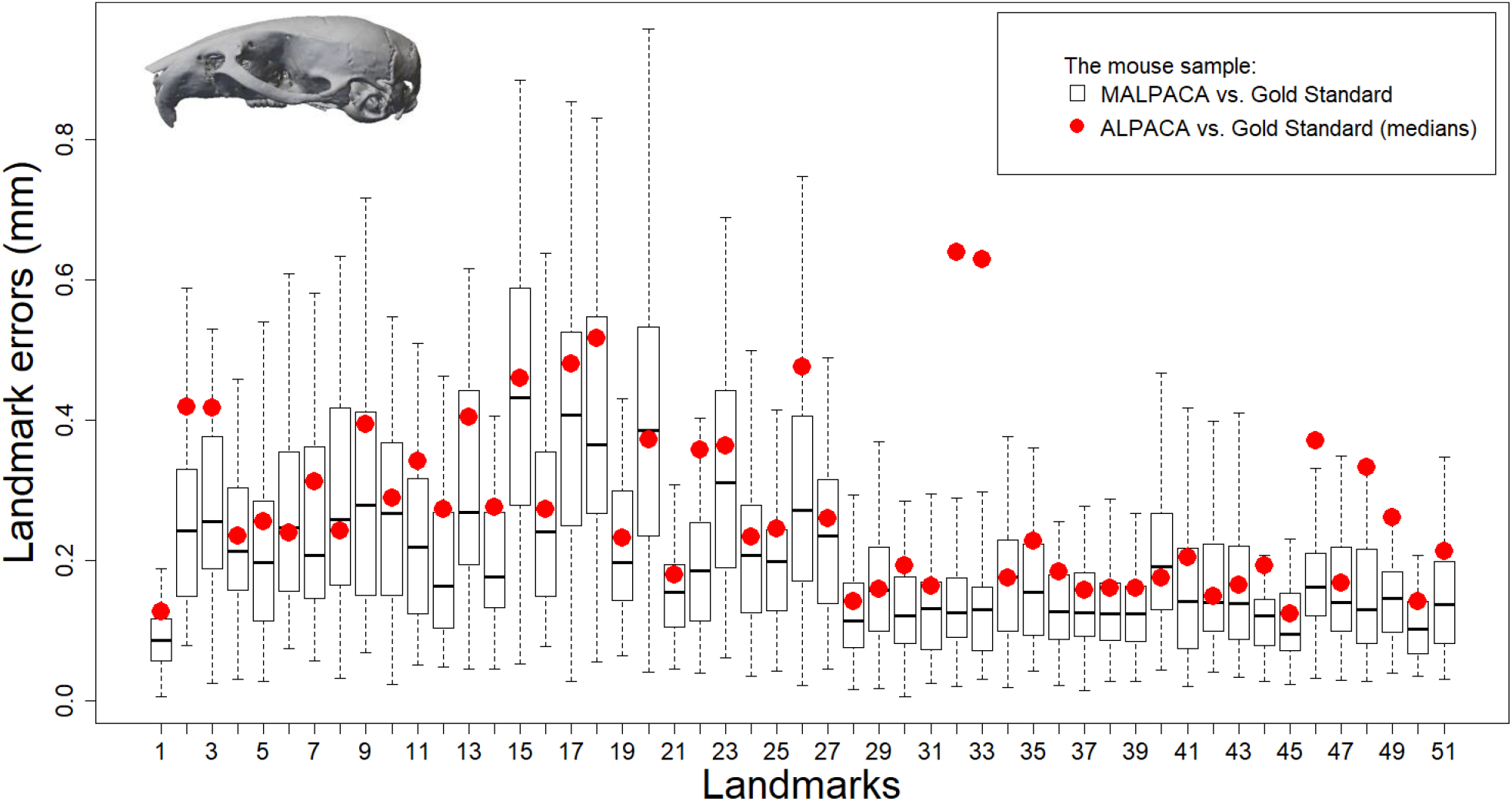
Difference of estimated landmarks from GS for **the mouse sample**. Errors represent by Euclidean distance (for errors expressed in percentage of centroid sizes, see Fig. S3). Boxes represent errors between MALPACA estimates and the Gold Standard (GS) landmarks. Red dots: median errors between the estimates of the synthetic template ALPACA and the GS.

**FIGURE 3.**
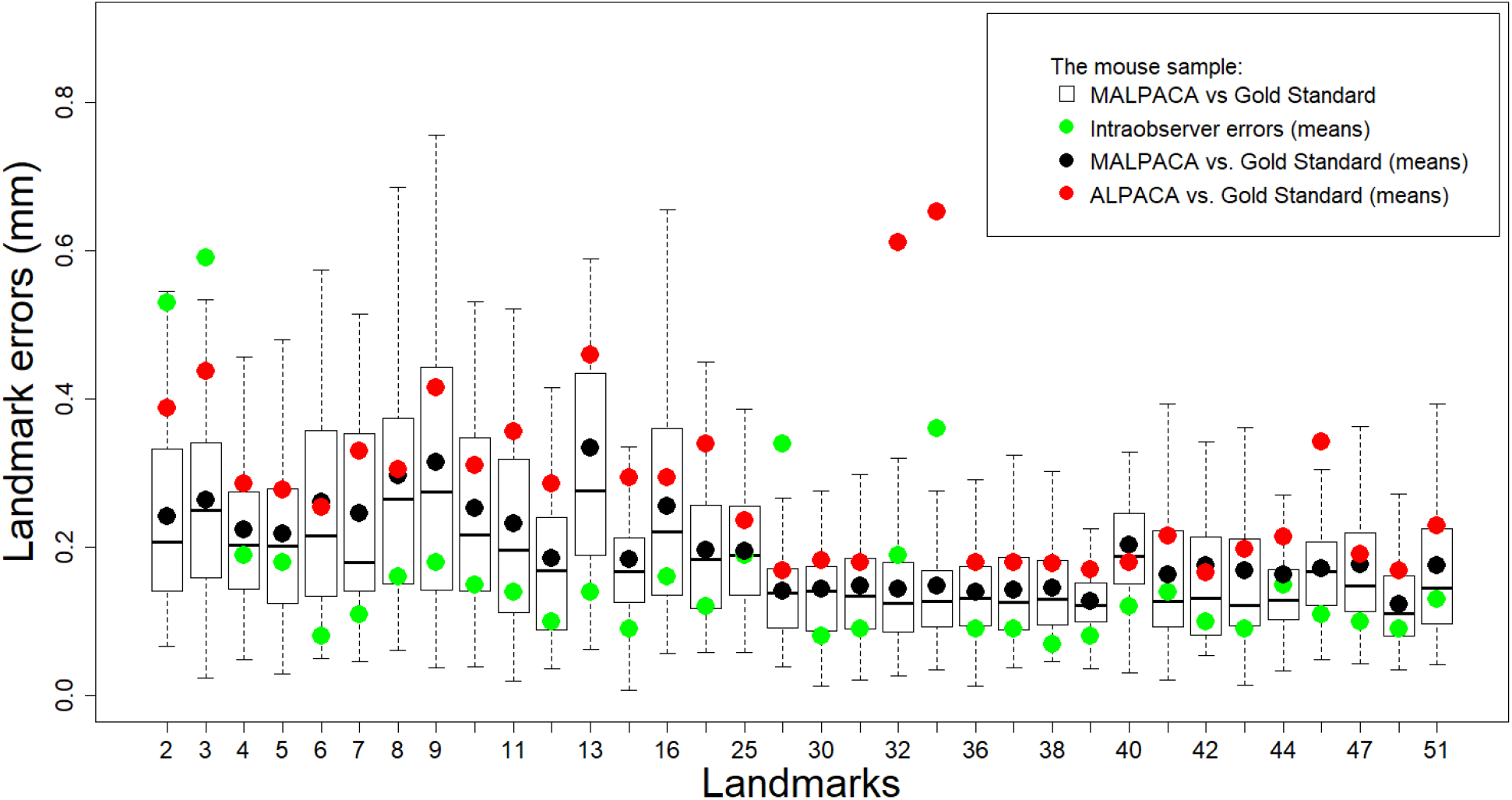
Individual landmark errors of **the mouse sample** comparing to intraobserver errors. Similar to Fig. 3, boxes represent errors between MALPACA and the GS for each landmark. Black dots: mean errors of mouse MALPACA. Green dots: mean intraobserver errors computed by Percival et al. (2019). The labels on the x-axis are landmarks in this study that overlap with those from Percival et al. (2019). Note that in b), to be in line to Percival et al.’s (2019) intraobserver errors, we performed two joint GPAs: one for MALPACA estimates and manual landmarks, and the other for ALPACA estimates and manual landmarks. The MALPACA-manual and ALPACA-manual errors are derived from these joint GPAs and are converted to distances measured in millimeter by scaling the coordinates with centroid sizes.

TABLE 2). For 37 out of 51 landmarks, the MALPACA-based errors are significantly smaller than the ALPACA using the synthetic template (p-values < 0.05 for one-sided t-tests for assessing if MALPACA-errors are smaller) (Fig. 3; Table S4). For the remaining 14 landmarks, the MALPACA-based errors are not significantly larger than the ALPACA ones (p-values > 0.37 for one-sided Welch t-tests assessing if ALPACA-based errors are smaller) (Fig. 3; Table S5).

**TABLE 2.** P-values from one sided Welch t-tests that examines whether ape MALPACA RMSEs are significantly smaller than RMSEs derived from ALPACA using individual mouse template.

| Mouse MALPACA one-sided Welch t-test for RMSEs | p-value |
| --- | --- |
| Vs. ALPACA (synthetic template) | $1.283 \times 10^{-16}$ |
| Vs. 129S1.SVIMJ ALPACA | $8.493 \times 10^{-10}$ |
| Vs. B6CBAF1 ALPACA | $8.522 \times 10^{-7}$ |
| Vs. BALB.CBYJ ALPACA | $2.539 \times 10^{-16}$ |
| Vs. CAST.EIJ ALPACA | $3.847 \times 10^{-28}$ |
| Vs. SF.CAMEIJ ALPACA | $3.496 \times 10^{-36}$ |
| Vs. SPRET.EIJ ALPACA | $5.313 \times 10^{-35}$ |
| Vs. X129P3.J ALPACA | $2.211 \times 10^{-16}$ |

To test for differences in the common morphometric variables (Procrustes distances, centroid sizes, PC ordination scores), we have investigated correlations of these variables from the separate generalized Procrustes alignments of MALPACA estimates and GS landmarks. For the mouse data, our results show that the Procrustes distances derived from MALPACA estimates and GS landmarks are highly correlated in pairwise distances with a 0.8 correlation coefficient (denoted as r) (Fig. 5a). Centroid sizes calculated from MALPACA estimates and GS landmarks are almost identical with the correlation coefficients exceed 0.99 (Table S6). For all these shape and size variables, the correlations of MALPACA derived estimates with GS exceed that of ALPACA and GS (Fig. 5; Table S6).

**FIGURE 4.**
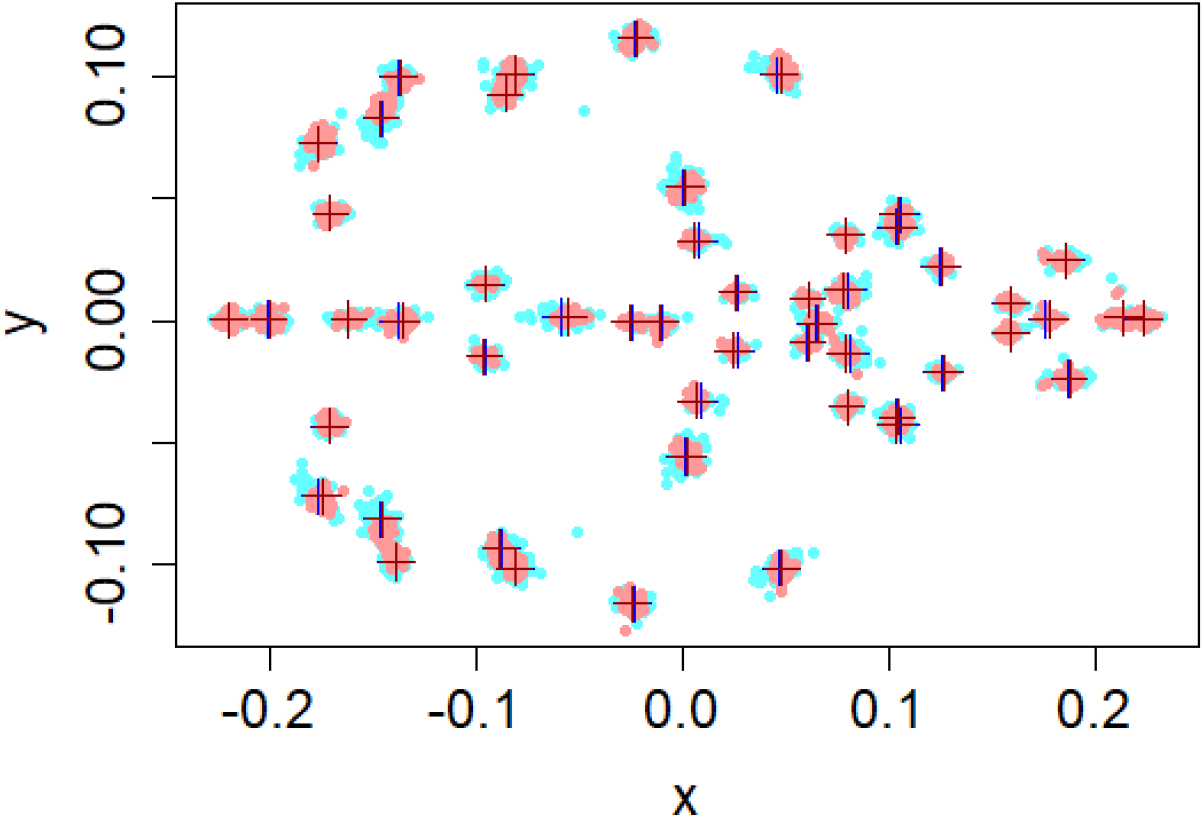
Superimposition of joint GPA for mouse MALPACA and manual landmarks (XY dimensions) based on **the mouse sample**. Light blue dots: all manual landmarks. Dark blue cross: mean manual landmarks. Light red dots: all MALPACA estimated landmarks. Deep red cross: mean MALPACA estimated landmarks.

**FIGURE 5.**
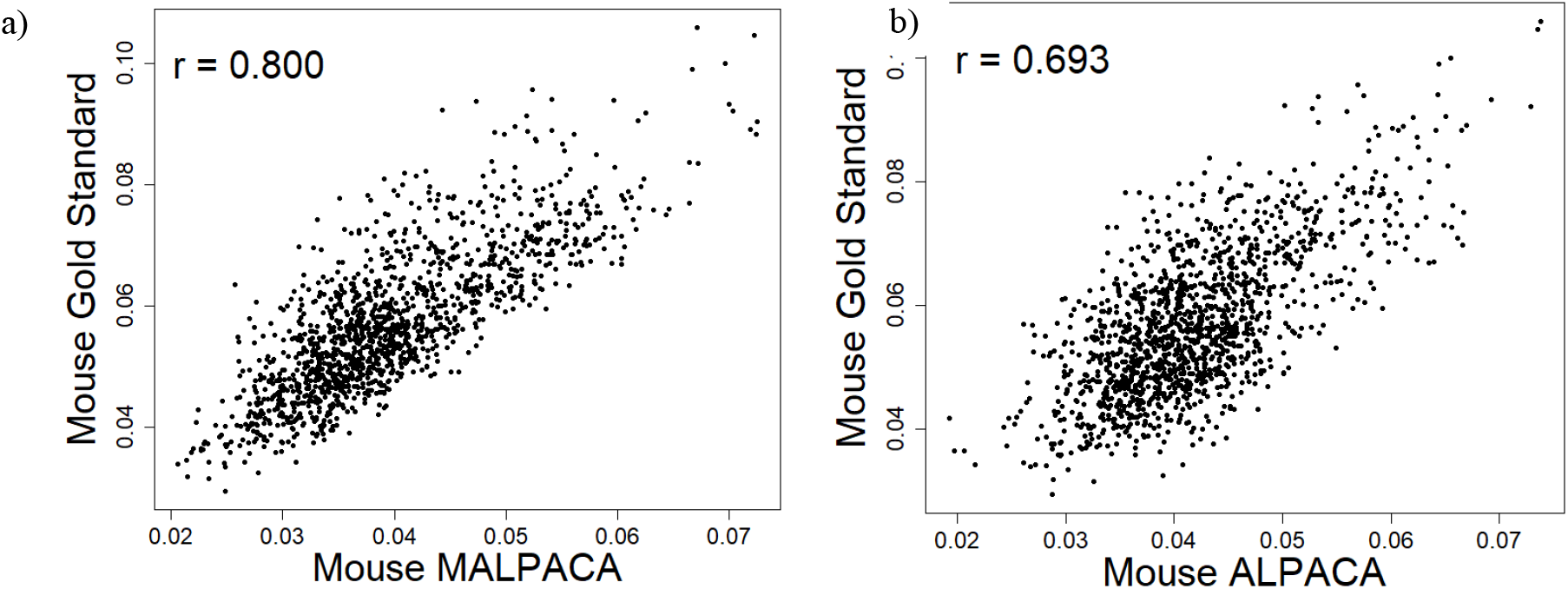
Plots of estimated against Gold Standard pairwise Procrustes distances based on **the mouse sample**: a) MALPACA vs. Gold Standard; b) ALPACA vs. Gold Standard based on the synthetic template. “r” = correlation coefficient. The plot is based on distances between all pairs of specimens within the mouse sample derived from separate General Procrustes Analyses of estimated and Gold Standard landmarks.

To compare the similarity of morphospaces derived from MALPACA and GS landmarks, we relied on the strength of the correlations of corresponding PC scores from different datasets. The MALPACA-GS correlation is especially strong in PC 1 scores as the correlation coefficient in PC 1 scores reaches 0.947 (Fig. 6a). MALPACA and GS landmarks also yield high correlation in PC 2 scores (r = 0.83), with a gradual decline of correlation coefficients with the other PC scores. The ALPACA-GS correlations in scores of PC 1 and PC 2 are still strong but relatively weaker than the MALPACA-GS correlations (Fig. 6b).

**FIGURE 6.**
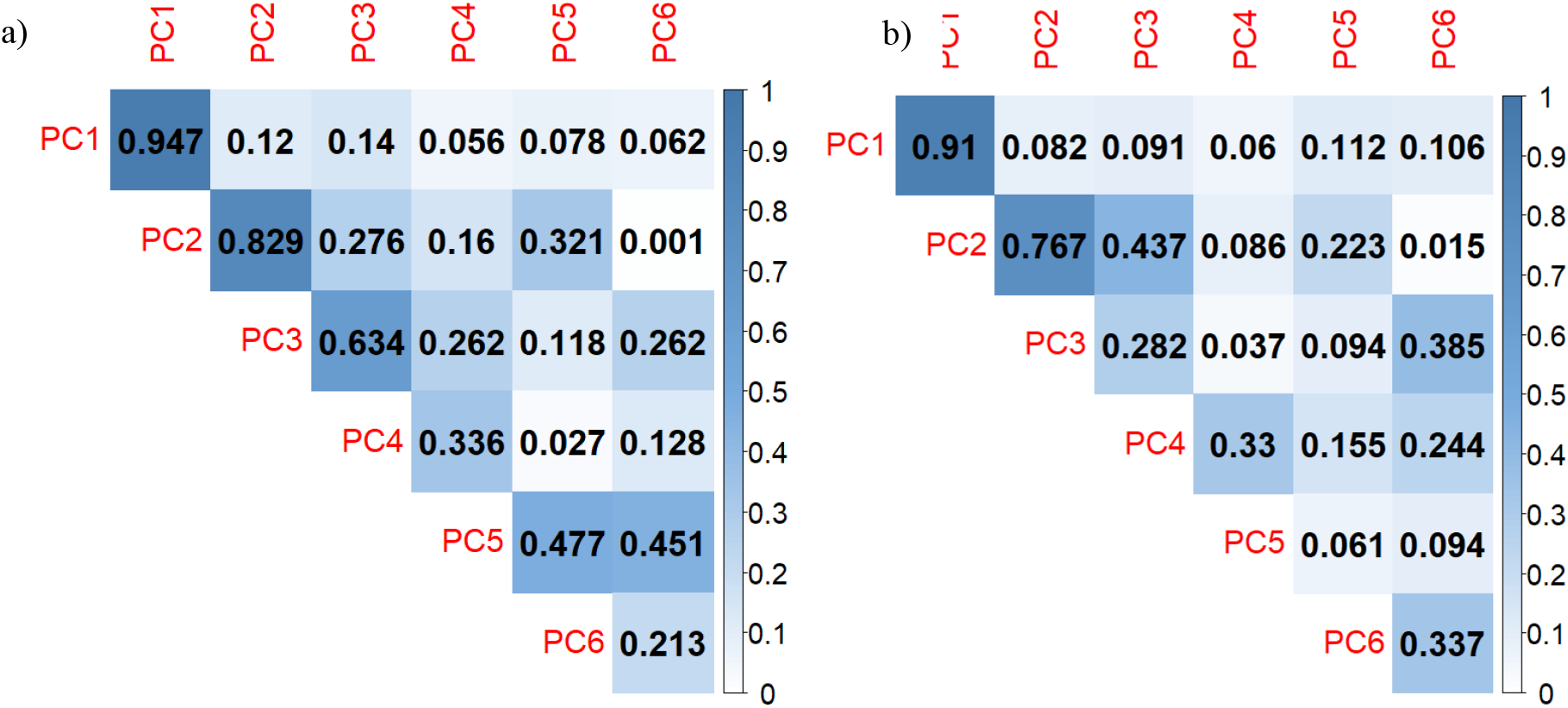
Correlations in the scores of the first six PCs (principal components) between estimated and Gold Standard landmarks based on **the mouse sample**: a) MALPCA vs. Gold Standard; b) ALPACA (the synthetic template) vs. Gold Standard. PC scores are derived from separate GPAs of estimated and manual landmarks.

Overall, the ape MALPACA-ALPACA performance evaluations are in line with the results from the mouse dataset. MALPACA yields significantly smaller RMSEs than does any ALPACA based on a K-means selected template (all p-values < 0.02) (Fig. 7; Fig. S4; TABLE 3). The mean shapes of MALPACA-based and GS landmarks derived from their joint GPA are highly consistent (Fig. 9). All centroid sizes from MALPACA and ALPACA show nearly perfect correlations with the GS as the correlation coefficients exceed 0.99, while the MALPACA-GS correlation is the highest (Table S7). Both pairwise Procrustes distances and PC 1 scores derived from MALPACA estimated landmarks very strongly correlates with those derived from GS (r = 0.912 and 0.985 respectively) (Fig. 10). The MALPACA estimates and GS also yield strong correlations in PC 2, PC 3 and PC 4 scores as the correlation coefficients all exceed 0.8 (Fig. 10b).

**TABLE 3.** P-values from one sided Welch t-tests that examines whether ape MALPACA RMSEs are significantly smaller than RMSEs derived from ALPACA using individual ape template.

| Ape 6-template MALPACA RMSEs | p-value |
| --- | --- |
| Vs. Pan 1 template USNM084655 ALPACA | $3.152 \times 10^{-7}$ |
| Vs. Pan 2 template USNM176236 ALPACA | $1.306 \times 10^{-7}$ |
| Vs. Gorilla 1 template USNM590953 ALPACA | 0.0179 |
| Vs. Gorilla2 template USNM599167 ALPACA | $4.329 \times 10^{-5}$ |
| Vs. Pongo 1 template USNM142185 ALPACA | $1.817 \times 10^{-13}$ |
| Vs. Pongo 2 template USNM153830 ALPACA | $2.738 \times 10^{-6}$ |

**FIGURE 7.**
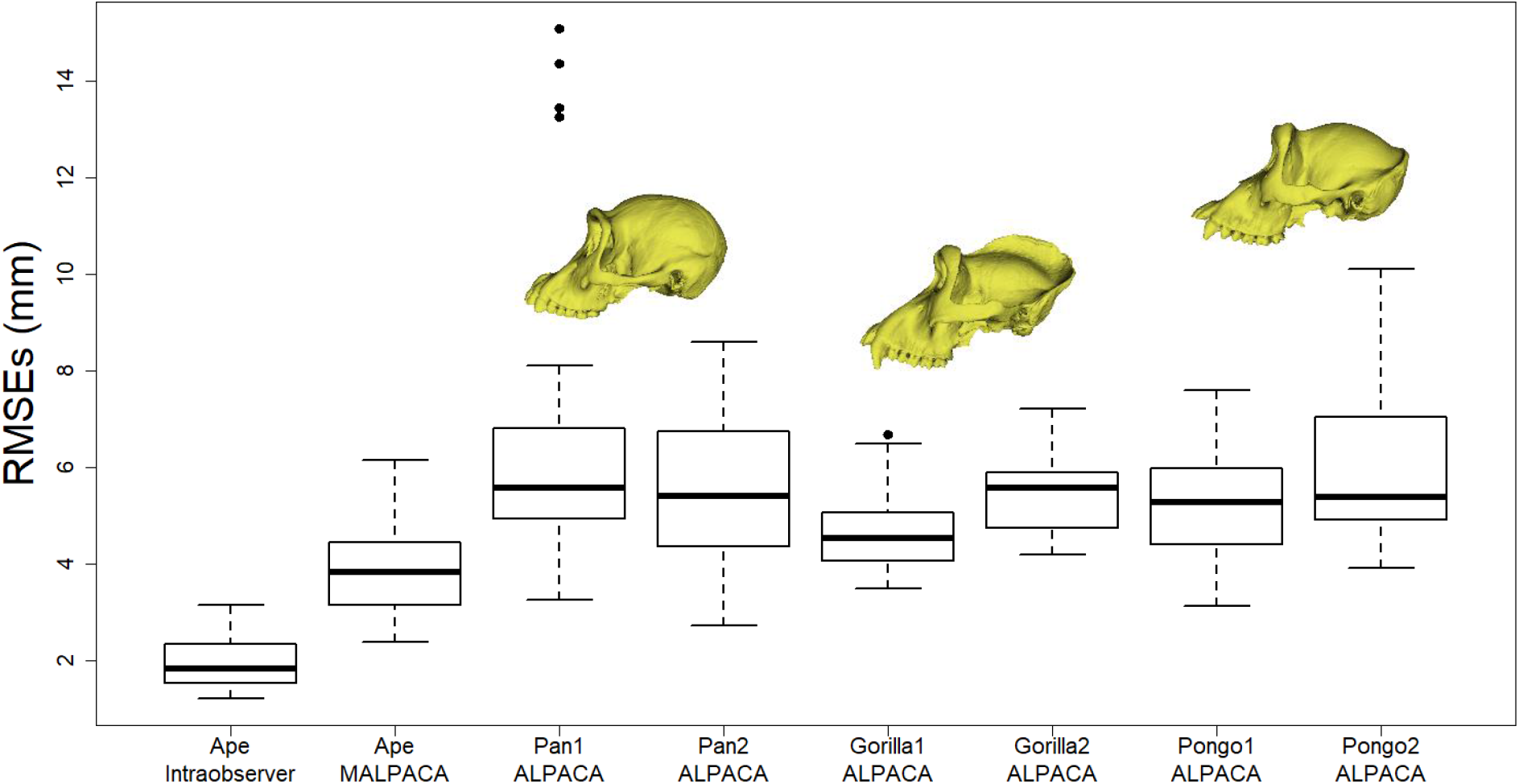
Ape MALPACA and ALPACA performance measured by RMSEs between estimated and manual landmarks. “Ape Intraobserver” refers to the RMSEs between two manual landmark datasets of the ape sample. See Fig S4 for RMSEs as percentage of centroid sizes. See Table 4 for the template used for each ALPACA based on a K-means selected template.

**FIGURE 8.**
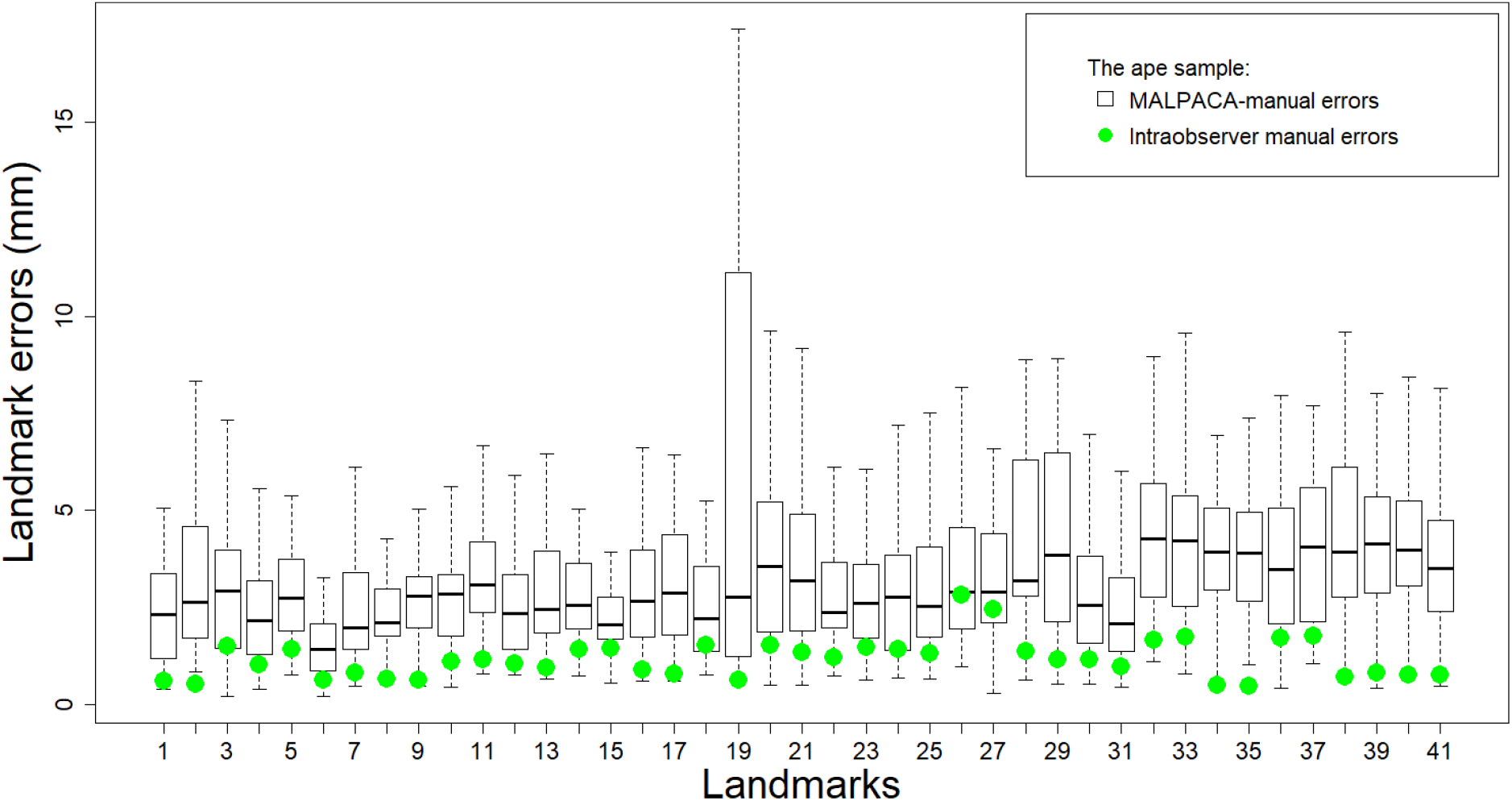
Individual landmark errors of **the mouse sample** (for errors in percentage of centroid sizes, see Fig. S3). Boxes represent errors between MALPACA estimates and the Gold Standard (GS) landmarks. Green dots represent median intraobserver manual landmark errors between two manual landmark sets.

**FIGURE 9.**
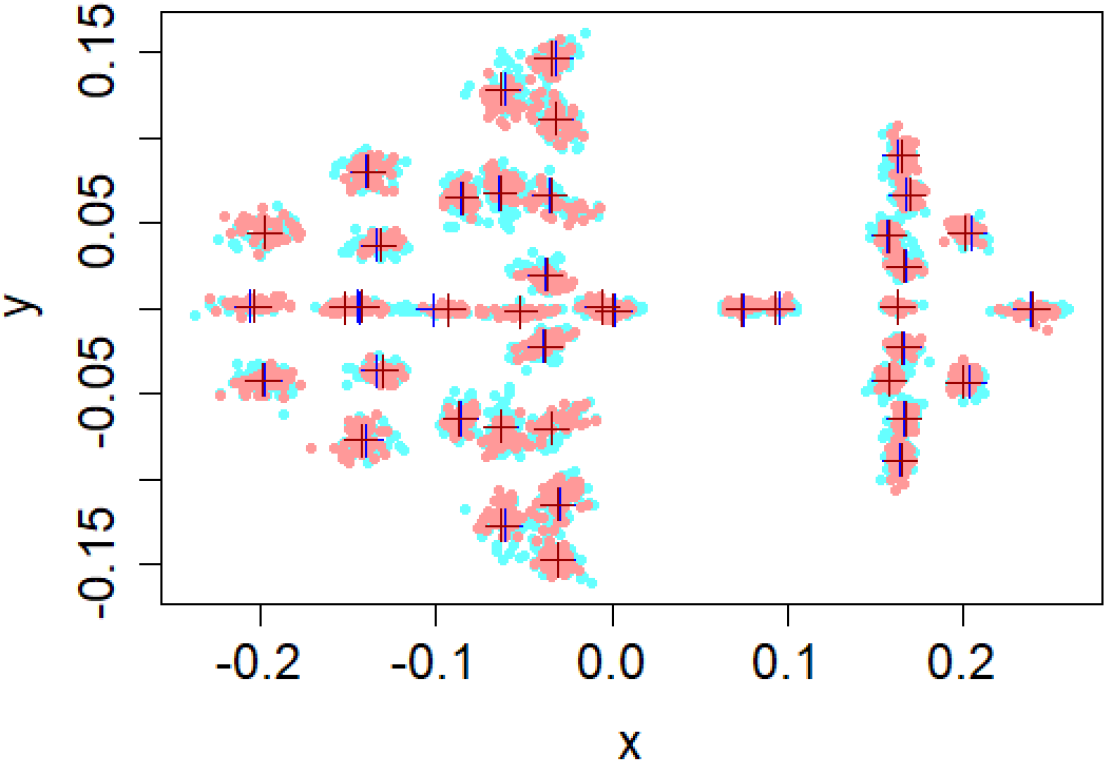
Superimposition of joint GPA for ape MALPACA and manual landmarks (XY dimensions) based on **the ape sample**. Light blue dots: all manual landmarks. Dark blue cross: mean manual landmarks. Light red dots: all MALPACA estimated landmarks. Deep red cross: mean MALPACA estimated landmarks.

**FIGURE 10.**
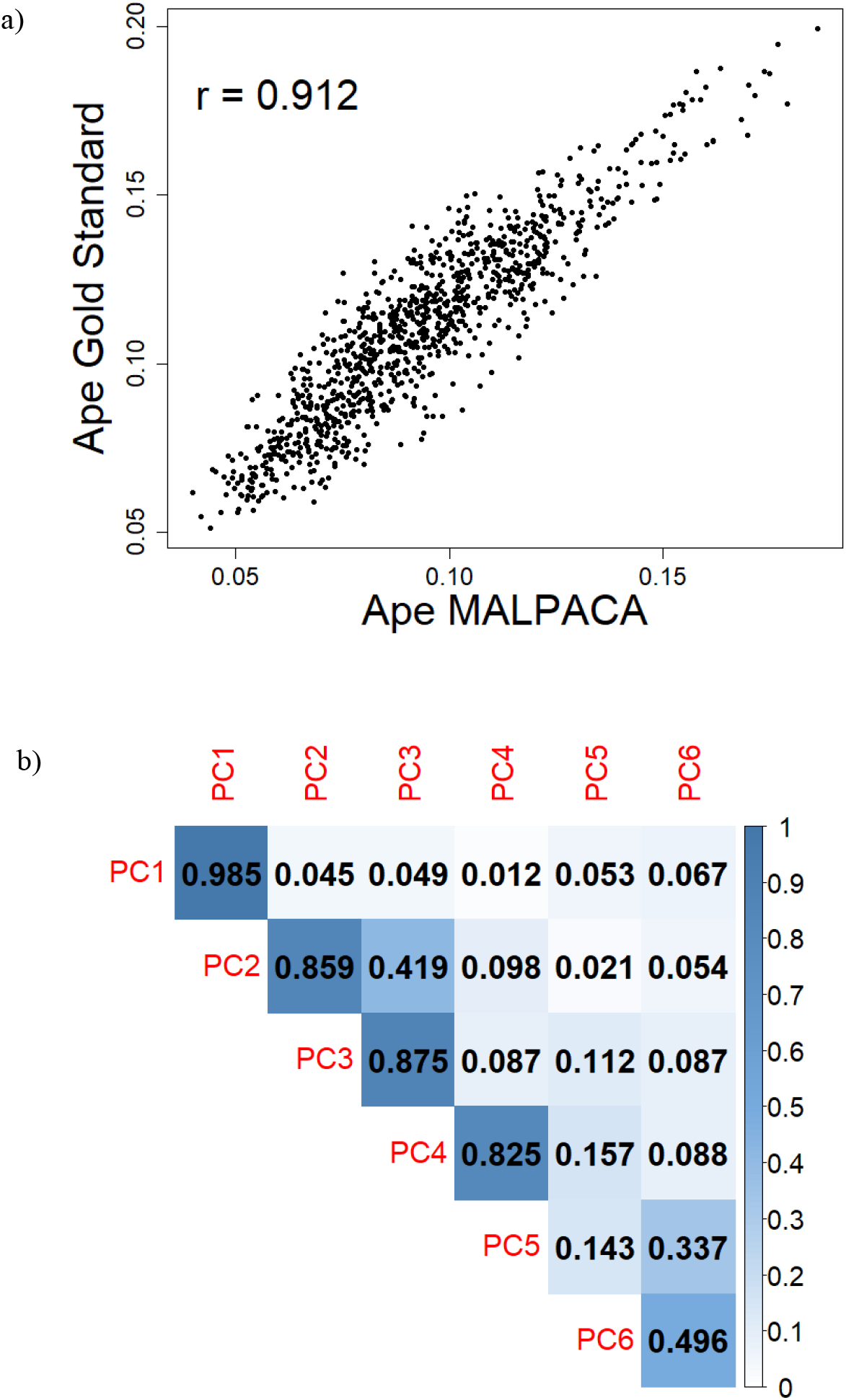
MALPACA-Gold Standard correlations based on **the ape sample**. a) Estimated against Gold Standard pairwise Procrustes distances; b) Correlations in the scores of the first six PCs (principal components) between estimated and Gold Standard landmarks

### Manual landmark errors

Fig. 3 compares the mouse manual landmarking errors (intraobserver errors) calculated by Percival et al (2019) to the errors between estimated and GS landmarks in our study for the 34 landmarks that are shared between these two studies. In general, intraobserver errors are the smallest with a mean error 0.163. The ALPACA-GS errors are much larger as the mean error reaches 0.285. The MALPACA-GS method outperforms ALPACA-GS with a mean error 0.197 and is more comparable to the manual landmarking error.

For the ape sample, manual intraobserver errors are measured by RMSEs, between two landmark sets. Again, the manual errors are significantly smaller than the errors between any set of estimates and GS landmarks (Fig. 8; Fig. S5). The one-sided Welch t-test for assessing whether the manual errors are smaller than the MALPACA-manual errors yield a p-value of 2.381 × 10^−19^. Comparing individual landmark errors show that for 38 of 41 landmarks, the intraobserver manual errors are significantly smaller than MALPACA-manual errors (p-values < 0.05 based on one-sided t-test that assesses if individual intraobserver landmark errors are smaller) (Fig. 8; Table S8).

### Permutation tests and the performance of K-means based templates

Permutation tests are carried out to assess whether K-means based templates outperforms randomly selected templates as a way to determine the efficacy of K-means multi-template selection (Table 4). For the mouse sample, 100 permutations are carried out to generate 100 different random combinations of templates. Consequently, MALPACAs is run 100 times on the set of 54 specimens, generating 5,400 RMSE scores. For the ape sample, 50 permutations are run for the set of 46 specimens, generating 2,300 RMSE scores. For the mouse dataset, 25 out of 54 specimens (46.2%) had a K-means based MALPACA estimate with an RMSE smaller than the 50th percentile of the 5,400 RMSEs from the permutation analysis (Fig. S6). The comparison of K-means versus random template selection for the ape dataset is consistent with the analysis of the mouse sample. Overall, 20 out of 46 specimens (43.5%) had a MALPACA estimate with RMSE error score below the 50th percentile of the 4,600 RMSEs from the permutation analysis (Fig. S7). In addition, the ranges of RMSEs derived from the ape and mouse K-means based MALPACA both overlap with the RMSEs derived from the permutation analysis (Table 4). These results show that K-means based selection of templates for MALPACA has a comparable level of performance to random selection of templates.

**TABLE 4.**
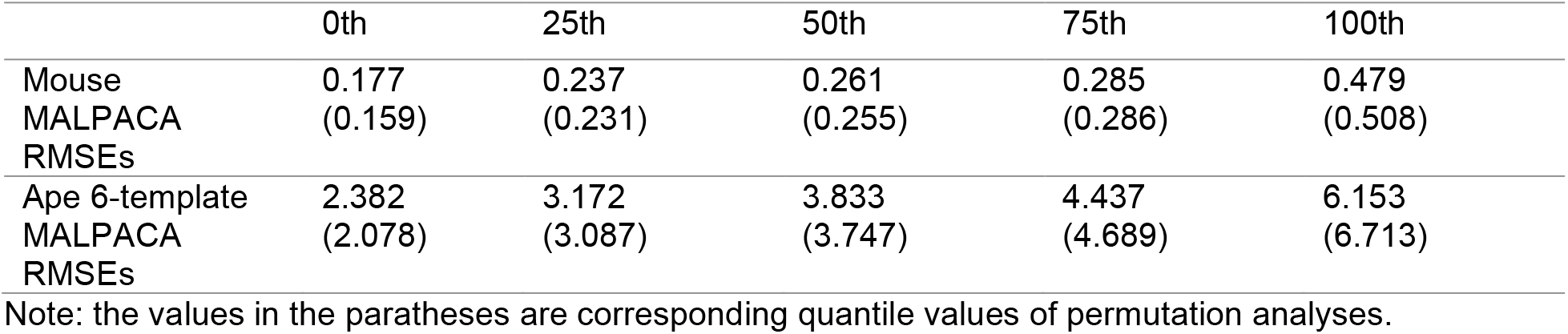
RMSEs of K-means-based MALPACAs comparing to the permutated RMSEs.

## DISCUSSION

### The Efficacy of K-means-based MALPACA

Template selection for properly representing the whole sample is a critical factor in determining the performance of any template based method to avoid potential biases in the outcome especially when the sample is highly variable (Antonelli et al. 2019; Gooding 2021; Schipaanboord et al. 2019). For example, in the field of automatic segmentation based on neuroimage registrations, various multi-atlas selection methods have been designed with differential accuracy for delineating organs (Antonelli et al. 2019; Schipaanboord et al. 2019). In this study, we proposed a convenient method to identify multiple samples that can be used as templates for the study population by applying K-means clustering to the PCA scores derived from the downsampled point cloud data of their 3D models. It is expected that this approach will be able to capture gross patterns of overall shape variations when there is no prior information available to the investigator to guide the template selection.

Our study confirms the expectation that MALPACA using K-means selected templates in general outperforms ALPACA because these templates can capture gross variations within a sample. In both the analyses of the mouse and ape samples, the MALPACA-derived landmark estimates are closer to the GS (manual) landmarks than any ALPACA. In most cases, MALPACA also produces pairwise Procrustes distances and centroid sizes closer to those of GS compared to ALPACA. Moreover, morphospaces derived from MALPACA-based PCs are more similar to GS-based PCs than those produced by ALPACA.

However, there are certain assumptions behind this approach that are important to consider. First, input models should contain the same corresponding structures across samples. For example, if 3D models of skull variably contain pieces of vertebrate, parts of mandibular joint, other non-cranial elements across samples, they will influence the analysis, because ultimately the analysis is dependent on the extracted point cloud. If a sample is partial or broken (e.g., missing a large section alveolar row on maxilla), it should be left out of the template selection analysis, because the correspondence of the point cloud generated from this sample is unlikely to match to that of others. It is important to note that actual ALPACA/MALPACA pipeline is robust with respect to small missing regions in specimens, and such specimens can still be used in the automated landmarking pipeline, provided that the landmark set does not contain any landmark that falls into the missing section. But they should be avoided during template selection procedure.

It should also be noted that templates selected by K-means may not be among the most optimal template sets for MALPACA. We carry out permutation analysis to perform a series of MALPACAs based on randomly selecting specimens as templates for both the mouse and ape sample. The K-means based MALPACA’s performance is intermediate compared to MALPACA based on randomly selected templates. Nevertheless, K-means can effectively avoid selecting poorly performed sets of templates. Furthermore, as shown in this study, MALPACA using K-means selected templates consistently generates more accurate landmarks than using a single template. Overall, Investigators can entirely skip the template selection procedure, if they have other means to determine what templates to use (e.g., prior data, similar genetic background etc.). However, if uncertainty around what templates should be used, the K-means multi-template selection method present in this study provides a reasonable solution for choosing specimens as templates for automated landmarking.

### Landmarking Errors and Consistency

In both the mouse and ape samples, the deviations between MALPACA and GS are obviously much larger than the errors between two manual landmarking trials. This is as expected, as intraobserver errors created by the same expert are usually very small. Still, the MALPACA-GS errors in landmark positions are more similar to the intraobserver errors than ALPACA-GS errors.

On the other hand, inter-observer errors are usually larger than intraobservers errors. These errors can be as significant as some biologically meaningful variations, such as intraspecific variations and sexual dimorphism (Robinson and Terhune 2017; Daboul et al. 2018; Percival et al. 2019). Furthermore, interobserver errors can be unpredictable as they have a variety of causes, including subjective understanding of anatomical variations, vague landmark protocols, and different landmark annotation tools (Robinson and Terhune 2017). In recent years, combining data collected by multiple researchers has become increasingly common. Consequently, while research collaboration and data sharing are greatly increasing time efficiency for data collection and sample size for higher statistical power, controlling interobserver errors is becoming more difficult and complicated. This can create issues in consistency and reproducibility. MALPACA, by contrasts, produces highly consistent landmarks when the same templates are used. Researchers can focus on carefully landmarking a short list of templates that can be shared with others. In this way, MALPACA can greatly facilitate data sharing while also ensuring consistency and reproducibility.

### *Multi-template estimates enable* post hoc *analyses of results*

One advantage of our multi-template pipeline is its potential for designing *post-hoc* analyses to assess landmarking quality. When there is only one estimate for a target, it is not easy to evaluate how well the automated landmarking performed without visualizing the estimate on the target model, or alternatively, manually landmarking the target specimen and calculating the errors. Both approaches are too tedious or downright unfeasible for a large study.

Here we provide an example of how individual estimates can be conveniently used for *post-hoc* analyses using simple heuristics in lieu of having gold standards. One of the *Pan* templates (USNM176236) chosen by K-mean template selection is a juvenile and yielded outlier estimates for four adult *Gorilla* specimens (Fig.7). This is because differences between this juvenile *Pan* template and the adult *Gorilla* specimens are large hence the global registration in ALPACA poorly aligned their point clouds.

In order to test whether these outliers may negatively impact the final output, we performed a *post hoc* test to determine and remove outlier estimates given by individual templates. Procedures described here are available as R functions incorporated to SlicerMorphR package.

To determine whether an estimated landmark for a specimen is an outlier, we first calculate the Euclidean distance between this landmark and its corresponding MALPACA estimate. We then defined a heuristic threshold as two standard deviations above the mean of the pooled distances between each individual estimate and its corresponding final output for one specimen. If the distance for an estimated landmark exceeds the threshold, this landmark is considered as an outlier. This function successfully pinpoints the outlier estimates for the four *Gorilla* specimens derived from the juvenile *Pan* template, along with other outliers that exceed the threshold (Fig. S8). It should be noted that the new estimates after removing all the outliers is not significantly better than the original result (p-value = 0.5751; see Fig. S8). This is because the MALPACA landmark estimates are based on medians, which are more robust to outliers than means.

Still, the issue derived from using a juvenile *Pan* template to landmark adult *Gorilla* specimens is indicative of situation when morphologically well-differentiated species exist within a study sample and including results from the templates of other species may have the potential to hamper the accuracy of the final estimates for specimens from each species. As a result, when templates from all distinct species are used for landmarking the whole sample (the “default” approach of MALPACA), the final output may be sub-optimal. Thus, for the three-species ape sample, we designed a “species-specific” approach that performs three separate MALPACA runs, each of which landmarks one species only using the two templates of that species (“species-specific MALPACA” in the following text). With the output estimates given by each individual template in hand, this was easily done by calculating median values between the two landmark sets of each ape species derived from its two templates.

We then compared the performance of the species-specific MALPACA to the “default” MALPACA using the ape sample by calculating RMSEs between landmarks generated by each approach and the GS. We also assessed correlations in centroid sizes, pairwise Procrustes distances, and PC scores. Detailed results are present in Fig. S10. In general, the results of the species-specific MALPACA are highly consistent with the “default” MALPACA. On the other hand, the species-specific MALPACA estimates did yield scores of PC 2 and PC 3 more correlated to those yielded by manual landmarks as the correlation coefficients both exceed 0.9. Overall, we encourage users to try different approaches based on estimates of individual template based on the structures of their samples. Since the MALPACA algorithm essentially runs multiple instances of ALPACA, it shares all the advantages of ALPACA in being lightweight and user friendly (Porto et al. 2021). Graphic user-interfaces for MALPACA and K-means multi-template selection is included in the existing open-access SlicerMorph module to facilitating the usage and exploration of these methods.

### Future Directions

K-means multi-template selection is determined by the sparse point clouds with point-to-point correspondence, which ultimately depends on the registration between each specimen and the reference. For convenience, this study selects the first one in the specimen list as the reference. If a different specimen is selected as the reference, the registration will be slightly different as will the point clouds. This may lead to a slightly different set of templates. Thus, it is important to assess how different choices of reference in point cloud generation may influence K-means multi-template selection, hence the final results of MALPACA. Furthermore, in this study, the efficacy of MALPACA is based on evaluating RMSE in landmark placement and a few size and morphometric variables. Because MALPACA is particularly suitable for studying highly variable and multi-species samples compared to single-template methods and is fast and lightweight, it would be interesting to use MALPACA in address questions in evolutionary and systematic biology with large sample size, such as phylogenetic and ontogenetic analysis, and then compared to the usage of manual landmarks.

## CONCLUSIONS

In this study, we confirm that MALPACA outperforms single-template ALPACA in generating landmarks closer to manually placed ones for both the single-population mouse and multi-species ape samples. The K-means based multi-template selection method proposed in this study can also generate a template set with good performance when researchers have no prior knowledge for optimal template selection.

MALPACA inherits the advantages of ALPACA in being light weight and easy to use but is more accurate and suitable for landmarking morphologically variable samples, such as the multi-species samples commonly encountered in evolutionary and systematic studies. Overall, MALPACA offers the potential for large-scale collaboration and data sharing for morphometric analysis while ensuring accuracy, consistency and reproducibility, thus contributing to make morphometrics fully embrace the “era of big data”.

## Supporting information

Supplementary Online Material Section 1

Supplementary Online Material Section 2

## ACKNOWLEDGEMENTS

This project was partly supported by grants from National Science Foundation (DBI/1759883, OAC/1939505), and National Institute for Dental and Craniofacial Research (DE027110) to AMM. We thank the Smithsonian’s Division of Mammals and Human Origins Program as well as Dr Matt Tocheri and Dr Kristofer Helgen for the scans of USNM specimens used in this research (http://humanorigins.si.edu/evidence/3d-collection/primate). These scans were acquired through the generous support of the Smithsonian 2.0 Fund and the Smithsonian’s Collections Care and Preservation Fund.

