## Supplementary Online Material Section 1 for "Automated Landmarking via Multiple Templates"

SUPPLEMENTARY MATERIALS: SECTION 1

TABLE S1. Specimen ID for the mouse sample (sample size = 61).

| Mouse ID | Mouse ID | Mouse ID |
|---|---|---|
| **129S1.SVIMJ (Template)** | C57BL.10J | NOD.SHILTJ |
| 129X1.SVJ | C57BL6.J | NOR.LTJ |
| A.J. | C57BLKS.J | NU.J |
| Ark.J | C57L.J | NZB.BINJ |
| B6AF1.J | CAF1.J | NZBWF1.J |
| B6C3F1.J | **CAST.EIJ (Template)** | NZO.HiLtJ |
| **B6CBAF1.J (Template)** | CB6F1.J | NZW.LACJ |
| B6D2F1.J | CBA.CAJ | PERC.EiJ |
| B6FVBF1.J | CBA.J | PL.J |
| B6SJLF1.J | CZECHII.EIJ | PWD.PhJ |
| B6129PF1.J | DBA.1J | PWK.PhJ |
| B6129SF1.J | DBA.2J | **SF.CamEiJ (Template)** |
| **BALB.CBYJ (Template)** | FVB.NJ | SJL.J |
| BALB.CH | I.LNJ | SKIVE.EiJ |
| BTBR.T.Itpr3tf | KK.HIJ | SM.J |
| BUB.BNJ | LEWES.J | **SPRET.EiJ (Template)** |
| C3D2F1.J | LG.J | SWR.J |
| C3H.HEJ | LP.J | TALLYHO.JNGJ |
| C3H.HEOUJ | MOLF.EiJ | **X129P3.J (Template)** |
| C3HEB.FEJ | MOLG.DnJ | |
| C57BL.6NJ | MRL.MPJ | |

Note: The seven specimens selected by K-means as templates are bold and marked by “template” in the parenthesis. The remaining 54 specimens were used for MALPACA pipeline. All mouse models and associated manual LM sets used in this study can be found at https://github.com/SlicerMorph/Mouse_Models

TABLE S2. Specimen IDs and species designation for the ape sample (sample size = 52).

| *Pan troglodytes* | *Gorilla gorilla* | *Pongo pygmaeus* |
|---|---|---|
| **USNM084655 (Template)** | USNM174715 | **USNM142185 (Templates)** |
| USNM174701 | USNM174722 | USNM142188 |
| USNM174703 | USNM176209 | USNM142189 |
| USNM174704 | USNM176211 | USNM142194 |
| USNM174707 | USNM176216 | USNM145300 |
| USNM174710 | USNM176217 | USNM145302 |
| USNM176228 | USNM176219 | USNM145303 |
| **USNM176236 (Template)** | USNM220060 | USNM145307 |
| USNM220062 | USNM220324 | USNM145308 |
| USNM220063 | USNM252575 | USNM145309 |
| USNM220065 | USNM252577 | USNM153805 |
| | USNM252578 | USNM153806 |
| | USNM252580 | USNM153822 |
| | USNM297857 | USNM153824 |
| | USNM582726 | **USNM153830 (Template)** |
| | USNM590942 | USNM197664 |
| | USNM590947 | USNM399047 |
| | USNM590951 | USNM588109 |
| | **USNM590953 (Template)** | |
| | USNM590954 | |
| | USNM599165 | |
| | USNM599166 | |
| | **USNM599167 (Template)** | |

Note: the two templates selected by K-means per species (six templates total) are bold and marked by by "template" in the parenthesis. The remaining 46 specimens are being landmarked.

TABLE S3. Settings for K-means multi-template selection.

| MALPACA | Spacing Factor (K-means) | Reference | Iterations | Set seed? |
|---|---|---|---|---|
| Mouse sample | 0.4 | 129S1.SVIMJ | 10,000 | Yes |
| Ape sample | 0.3 | USNM084655 | 10,000 | Yes |

Note: MALPACA uses the default settings listed in the ALPACA module of the SlicerMorph extension. See Porto et al. (2021) for detail. Spacing factor determines the point cloud density for K-means. See section 2 of the supplementary material for details.

a)

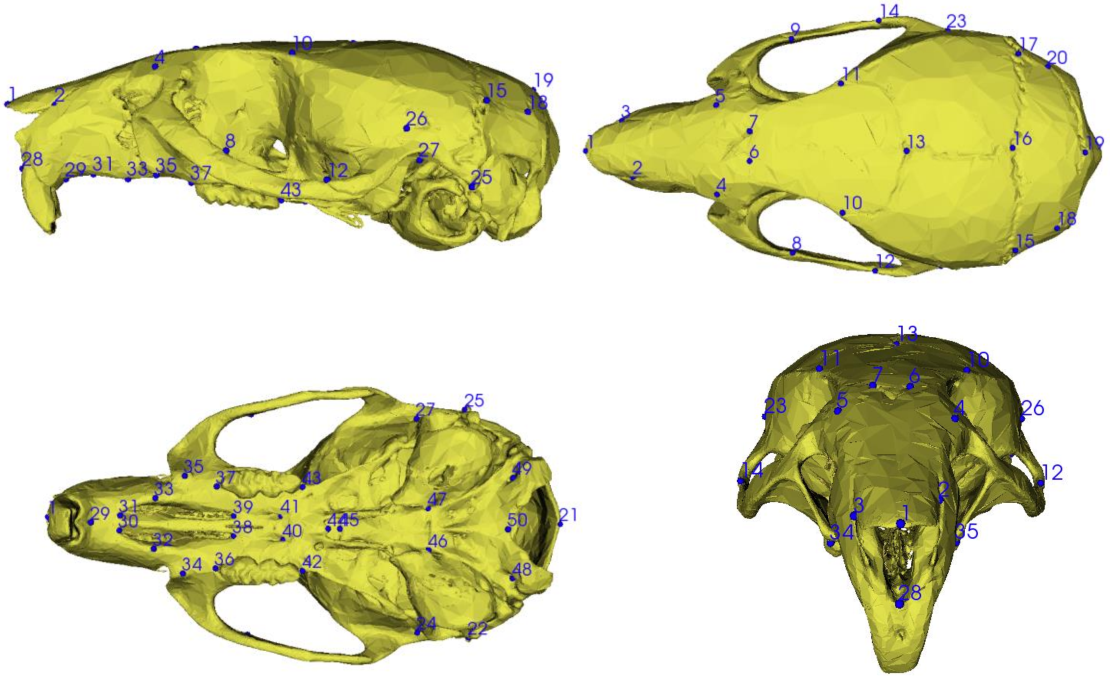

1 2 4 10 15 19 18 26 8 27 28 29 31 33 35 37 12 25 43
9 14 23 17 20 11 3 5 7 1 6 13 16 19 2 4 10 8 12 15 18
27 25 35 37 43 33 49 1 29 31 39 41 47 21 30 38 40 44 45 50 32 34 36 42 46 48 24 22
13 11 7 6 10 23 5 4 26 14 12 3 2 1 34 35 28


b)

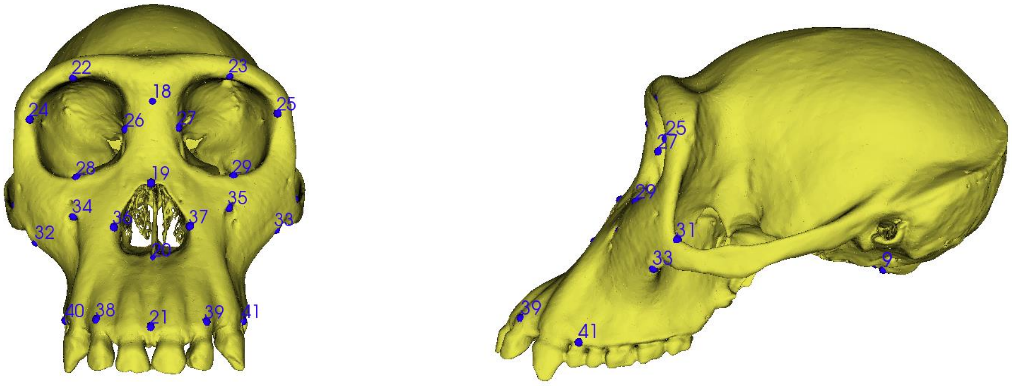

22 23 18 24 26 27 25 28 19 29 34 35 32 36 37 33 20 40 38 21 39 41
25 27 29 31 33 9 39 41

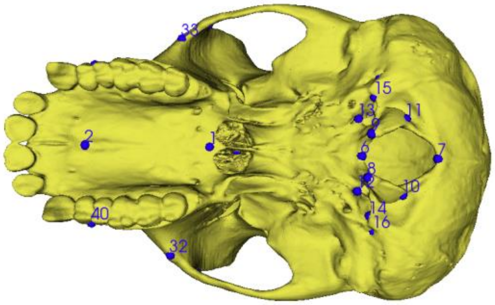


FIGURE S1. Anatomical landmarks used in this study for a) the mouse sample and b) the ape sample.

TABLE S4. P-values of one-sided Welch t-tests that compare if MALPACA individual landmark errors are smaller than ALPACA errors

| Landmark (Mouse) | p-value | Landmark (Mouse) | p-value |
|---|---|---|---|
| 1 | $7.262 \times 10^{-5}$ | 28 | 0.007361715 |
| 2 | $1.738 \times 10^{-6}$ | 30 | 0.00246 |
| 3 | $3.674 \times 10^{-6}$ | 31 | 0.00107 |
| 4 | 0.0175 | 32 | $1.305 \times 10^{-36}$ |
| 5 | 0.0256 | 33 | $7.705 \times 10^{-40}$ |
| 7 | 0.00738 | 35 | $2.802 \times 10^{-5}$ |
| 9 | 0.0315 | 36 | 0.000229 |
| 10 | 0.0324 | 37 | 0.0410 |
| 11 | 0.000713 | 38 | 0.00968 |
| 12 | 0.00369 | 39 | 0.000966 |
| 13 | 0.00373 | 41 | 0.00153 |
| 14 | $5.728 \times 10^{-5}$ | 44 | 0.00636 |
| 18 | 0.00198 | 45 | 0.0190 |
| 19 | 0.0256 | 46 | $2.342 \times 10^{-15}$ |
| 21 | 0.0419 | 48 | $1.554 \times 10^{-14}$ |
| 22 | $3.912 \times 10^{-9}$ | 49 | $1.377 \times 10^{-9}$ |
| 25 | 0.00139 | 50 | 0.0005536854 |
| 26 | 0.000737 | 51 | 0.001063129 |
| 27 | 0.0305 | | |

Note: MALPACA errors versus manual landmarks significantly smaller than ALPACA errors in the 37 landmarks listed.

TABLE S5. P-values of one-sided Welch t-tests that compare if ALPACA individual landmark errors are smaller than MALPACA errors for the mouse sample

| Landmark (Mouse) | p-value | Landmark (Mouse) | p-value |
|---|---|---|---|
| 6 | 0.379 | 24 | 0.886 |
| 8 | 0.505 | 29 | 0.774 |
| 15 | 0.359 | 34 | 0.927 |
| 16 | 0.934 | 40 | 0.332 |
| 17 | 0.765 | 42 | 0.365 |
| 20 | 0.396 | 43 | 0.819 |
| 23 | 0.716 | 47 | 0.806 |

Note: Only listing landmarks not listed in Table S4, i.e., MALPACA estimates do not show significantly smaller errors than ALPACA ones in these 14 landmarks according to the one-sided t-test in Table S4.

TABLE S6. Correlations between estimated and manual landmarks in centroid sizes for the mouse sample

| | Correlation coefficients |
|---|---|
| MALPACA | 0.997 |
| ALPACA (Synthetic template) | 0.993 |
| 129S1.SVIMJ ALPACA | 0.996 |
| B6CBAF1 ALPACA | 0.994 |
| BALB.CBYJ ALPACA | 0.996 |
| CAST.EIJ ALPACA | 0.995 |
| SF.CAMEIJ ALPACA | 0.995 |
| SPRET.EIJ ALPACA | 0.994 |
| X129P3.J ALPACA | 0.995 |

TABLE S7. Correlations between estimated and manual landmarks in centroid sizes for the ape sample

| | Correlations with manually placed landmarks in centroid sizes |
|---|---|
| MALPACA | 0.999 |
| Pan 1 ALPACA (USNM084655) | 0.997 |
| Pan 2 ALPACA (USNM176236) | 0.998 |
| Gorilla 1 ALPACA (USNM590953) | 0.998 |
| Gorilla 2 ALPACA (USNM599167) | 0.997 |
| Pongo 1 ALPACA (USNM142185) | 0.997 |
| Pongo 2 ALPACA(USNM153830) | 0.997 |
| Species-specific MALPACA | 0.994 |

TABLE S8. P-values of one-sided Welch t-tests that compare if manual landmark errors are smaller than MALPACA-manual errors

| Landmark (ape) | P-values | Landmark (ape) | P-values |
|---|---|---|---|
| 1 | $3.254 \times 10^{-8}$ | 22 | $8.522 \times 10^{-8}$ |
| 2 | $2.0607 \times 10^{-11}$ | 23 | $8.463 \times 10^{-7}$ |
| 3 | 0.000681 | 24 | $1.405 \times 10^{-6}$ |
| 4 | 0.0491 | 25 | $3.940 \times 10^{-6}$ |
| 5 | $1.952 \times 10^{-5}$ | 26 | 0.4099142 |
| 6 | $2.225 \times 10^{-6}$ | 27 | 0.2822061 |
| 7 | $3.960 \times 10^{-7}$ | 28 | $4.081 \times 10^{-5}$ |
| 8 | $4.316 \times 10^{-13}$ | 29 | $4.869 \times 10^{-6}$ |
| 9 | $1.015 \times 10^{-15}$ | 30 | $7.860 \times 10^{-8}$ |
| 10 | $1.205 \times 10^{-7}$ | 31 | $4.909 \times 10^{-6}$ |
| 11 | $7.325 \times 10^{-10}$ | 32 | $7.872 \times 10^{-11}$ |
| 12 | 0.00415 | 33 | $3.388 \times 10^{-8}$ |
| 13 | $3.233 \times 10^{-5}$ | 34 | $3.688 \times 10^{-17}$ |
| 14 | $1.093 \times 10^{-5}$ | 35 | $4.092 \times 10^{-18}$ |
| 15 | 0.00520 | 36 | $1.908 \times 10^{-5}$ |
| 16 | $4.318 \times 10^{-10}$ | 37 | $3.464 \times 10^{-6}$ |
| 17 | $7.573 \times 10^{-11}$ | 38 | $9.577 \times 10^{-13}$ |
| 18 | 0.227 | 39 | $1.183 \times 10^{-13}$ |
| 19 | $1.562 \times 10^{-8}$ | 40 | $1.218 \times 10^{-14}$ |
| 20 | 0.000237 | 41 | $3.310 \times 10^{-13}$ |
| 21 | $5.922 \times 10^{-5}$ | | |

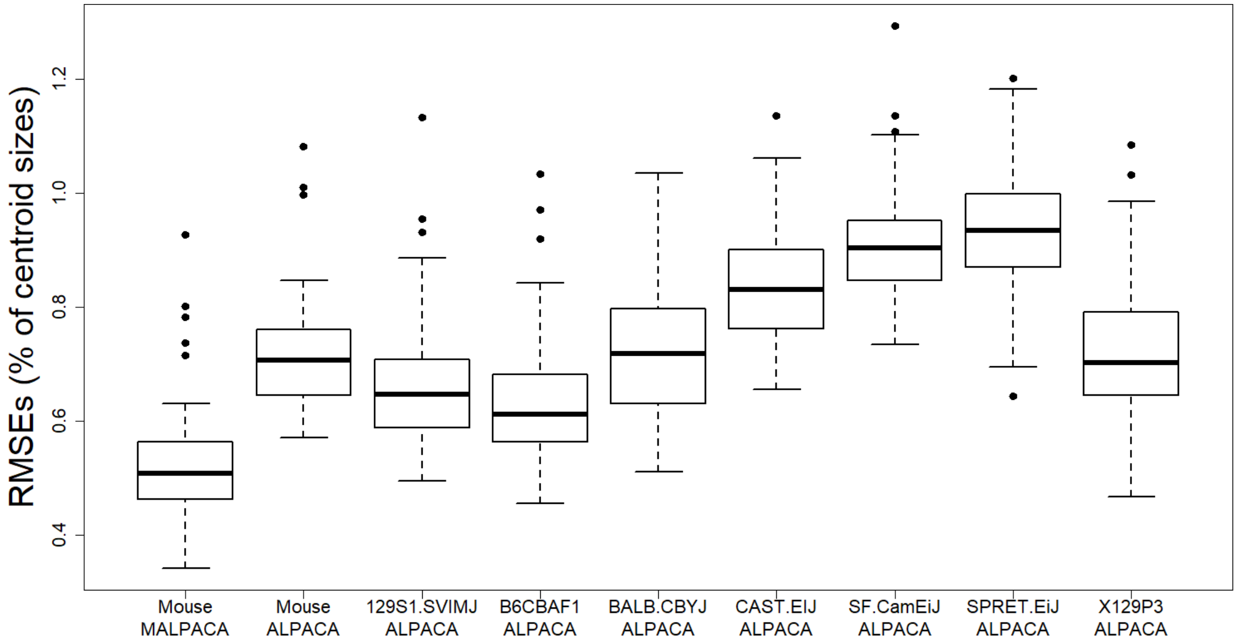


FIGURE S2. Comparison of RMSEs as percentage of centroid sizes between estimated landmarks and the “Gold Standard” for the mouse sample. Centroid size of each specimen is calculated using the Gold Standard landmark set. **Mouse ALPACA**: ALPACA based on synthetic mouse template used in Porto et al., 2021. Other boxes are ALPACA estimates using specified template.

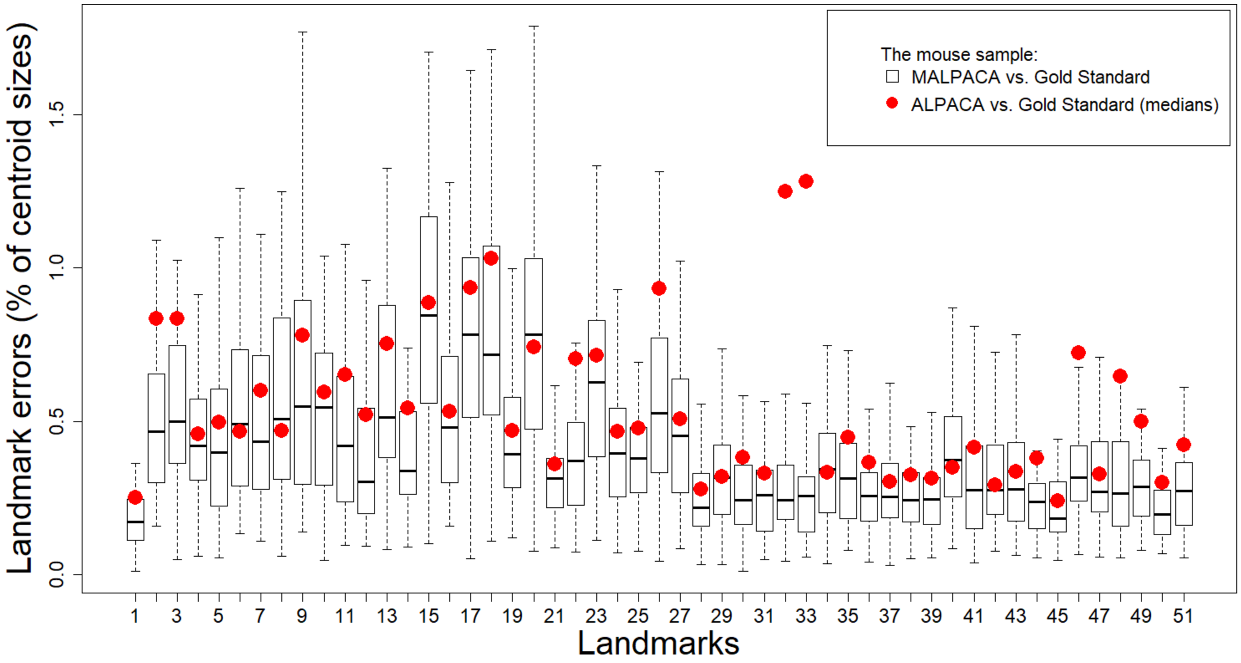


FIGURE S3. Individual landmark errors of the mouse sample as percentage in centroid sizes. Boxes represent errors between MALPACA estimates and the Gold Standard (GS) landmarks. Red dots: median errors between the estimates of the synthetic template ALPACA and the GS. Centroid sizes are calculated from the Gold Standard landmark set.

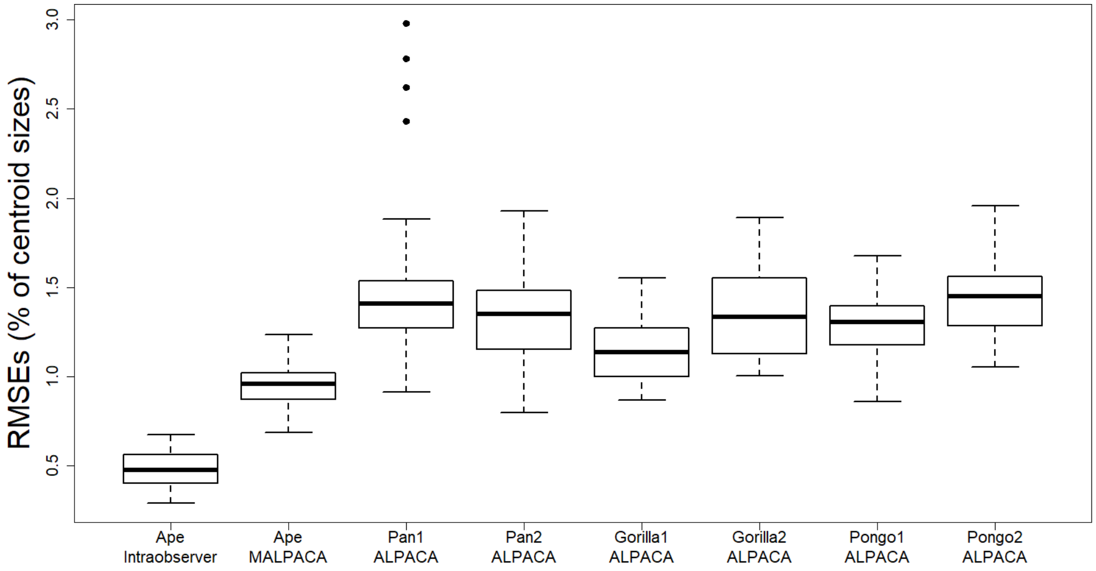


FIGURE S4. Ape MALPACA and ALPACA performance measured by RMSEs (percentage of centroid sizes) between estimated and manual landmarks. “Manual” refers to the RMSEs between two manual landmark datasets of the ape sample. See Table 4 for the template used for each ALPACA based on a K-means selected template set. Centroid sizes are calculated from the Gold Standard landmark set.

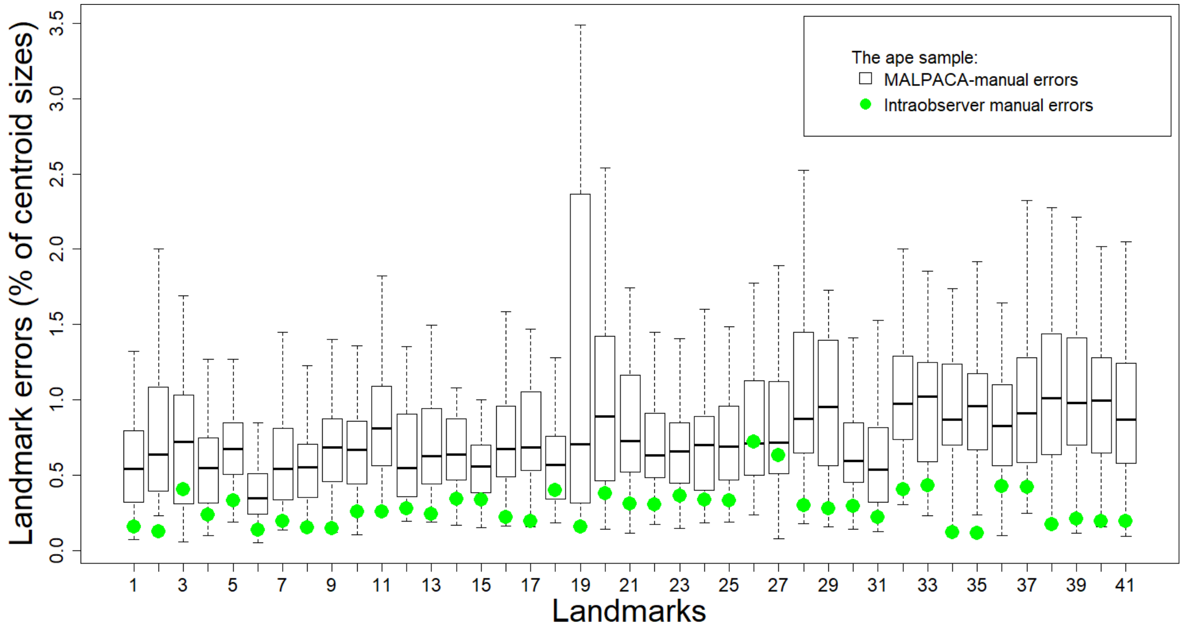


FIGURE S5. Individual landmark errors of the mouse sample as in percentage of centroid sizes. Boxes represent errors between MALPACA estimates and the Gold Standard (GS) landmarks. Green dots represent median intraobserver manual landmark errors between two manual landmark sets. Centroid sizes are calculated from the Gold Standard landmark set.

FIGURE S6 (See FigS6.pdf). Boxplots for the mouse permutation test. The red vertical line in each graph represents the RMSE between each specimen's landmark set generated by the K-means based MALPACA and the Gold Standard. The boxplot represents the pooled 5,400 RMSEs between estimates generated by randomly selected seven templates in each round of the permutation test and the Gold Standard.

FIGURE S7 (See FigS7.pdf). Boxplots for the ape permutation test. The red vertical line in each graph represents the RMSE between each specimen's landmark set generated by the K-means based MALPACA and the Gold Standard. The boxplot represents the pooled 2,300 RMSEs between estimates generated by randomly selected six templates (two templates per ape species) in each round of the permutation test and the Gold Standard.

FIGURE S8 (See FigS8.pdf). Boxplots for the ape individual estimates and the threshold for choosing outliers. Each graph shows the boxplot of distances between individual estimates of one ape specimen to the corresponding MALPACA final output. The red horizontal line represents the threshold, which is two standard deviations above the mean of the distances for that specimen.

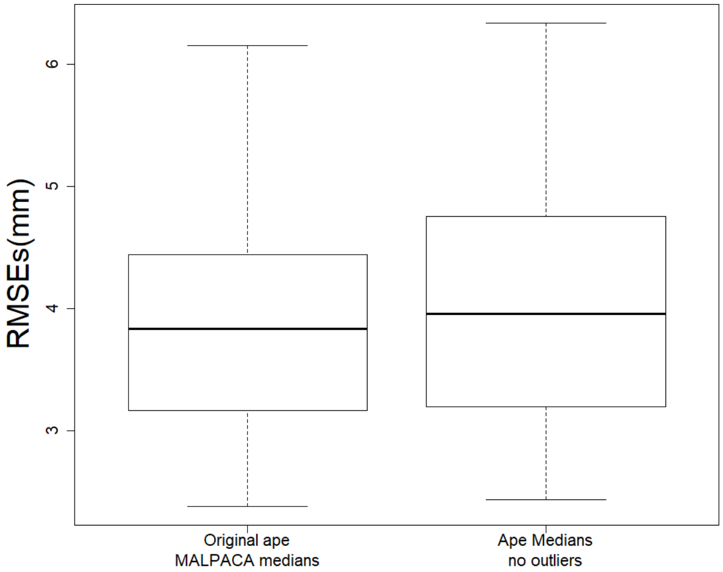


FIGURE S9. The boxplot for comparing the performance the ape MALPACA and the new median estimates achieved after removing outlier estimates demonstrated in Fig. S8. Comparison is based on calculating RMSEs between estimates and the Gold Standard. Two-sided Welch t-test shows that the RMSEs yielded from these two analyses are not significantly different with a p-value 0.5751.

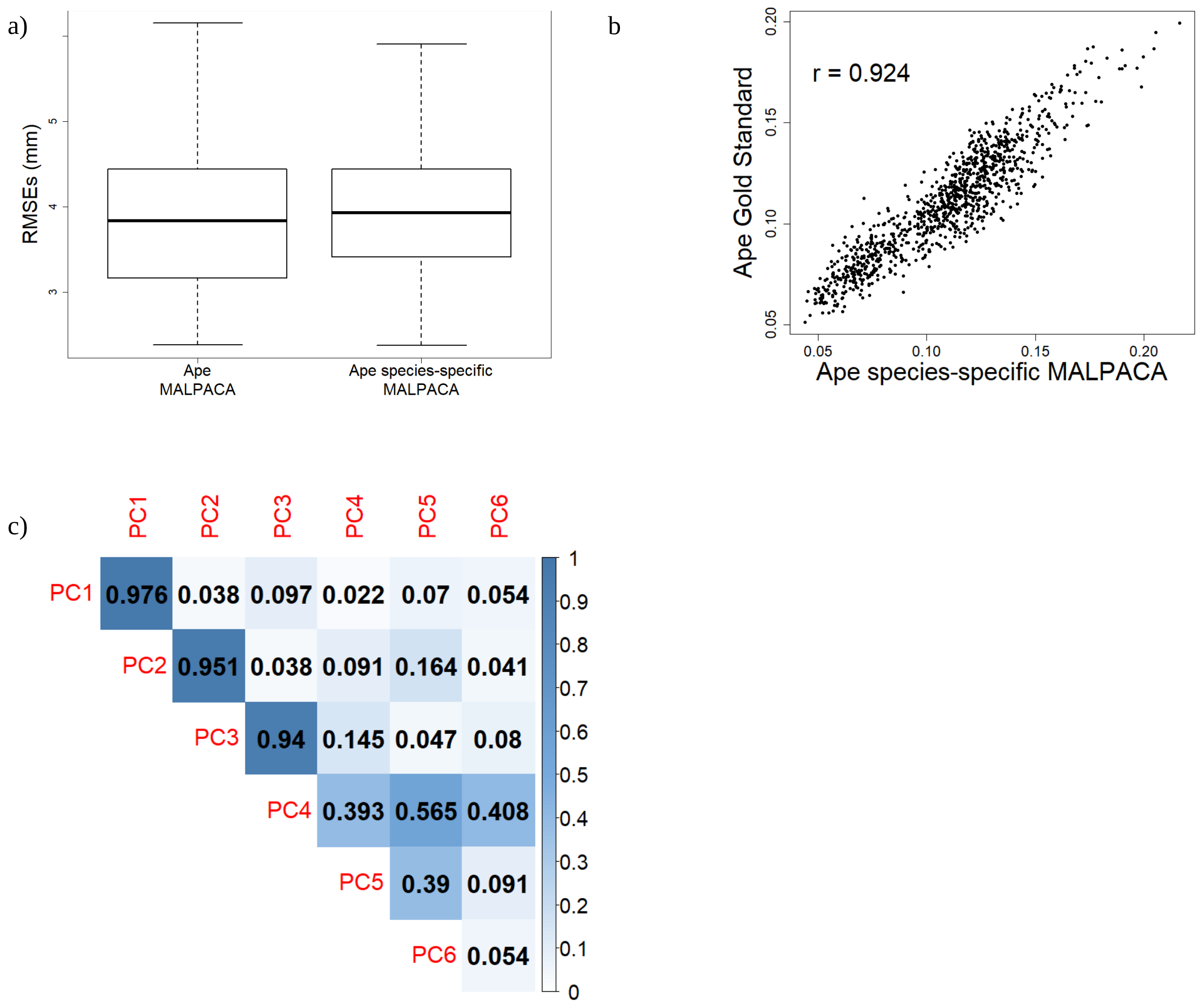


FIGURE S10. Performance of ape species-specific MALPACA comparing to manual landmarks. Species-specific MALPACA refers to performing a MALPACA for one species only using templates of that species. a) MALPACA- and species-specific MALPACA-manual RMSEs. b) Correlation in the pairwise Procrustes distances derived from species-specific MALPACA and manual landmarks. c) Correlations in PC scores derived from species-specific MALPACA and manual landmarks.
