## Supplementary Online Material Section 2 for "Automated Landmarking via Multiple Templates"

### SUPPLEMENT 2: STEP-BY-STEP INSTRUCTIONS FOR RUNNING MALPACA

**To the editor and reviewers:**

Installation and setup instruction in this SOM is temporary, as the current code being reviewed is not incorporated to the main branch of SlicerMorph. When the review of the paper is finalized, we will push these changes to the main SlicerMorph repository and make it publicly available, at which point these instructions will be superseded by the official SlicerMorph installation and update documentation in the repository https://github.com/SlicerMorph/SlicerMorph#installation). Users following these official methods will have all the necessary packages.

We have provided two archives of SlicerMorph with the MALPACA and necessary python libraries bundled for MacOS and Windows. Linux version of MALPACA is forthcoming.

- To obtain SlicerMorph with MALPACA on Windows 10, please use this link.
- To obtain SlicerMorph with MALPACA on MacOS, please use this link.

Extract the contents of the downloaded archive to a convenient location and click the **Slicer** executable at the top level of the archive to initiate the 3D Slicer/SlicerMorph session. Introductions to the 3D Slicer UI and how to navigate between modules can be found at https://slicer.readthedocs.io/en/latest/user_guide/user_interface.html.

**Sample Data to run MALPACA:** Entire set of mouse skull models and associated manual LMs used in the study can be found at https://github.com/SlicerMorph/mouse_models. Please clone or download this repository as the remaining steps will use them as sample data. As noted in the data availability statement, Smithsonian Institution does not allow us to re-distribute the 3D models we have extracted from the CT scans of their ape collection. Please contact Smithsonian Institution Digitization Program Office (https://dpo.si.edu/) to obtain access to the ape skull scans.

Please follow the rest of the document to:

1. Run the K-means multi-template selection procedure. This step can be skipped, if the user already knows what samples are going to be used as templates.
2. Use the ALPACA in batch mode using the multi-template method.
3. Use the SlicerMorphR R package to do quality control on the MALPACA estimate. We provide an example script to detect, remove outlier and redo the estimates. This step is optional, and users can implement their own quality control pipelines using the example provided.

#### I. K-means multi-template selection (Optional Step)

1. Switch to the "Templates Selection" tab (yellow). In the "Templates Selection Setup" collapsible section, specify entries to enable buttons in the following sections:

- Models Directory (red): specify the folder that contains the original 3D models in ". ply" format. If you have downloaded the mouse models repository from github, this would be the contents of the 'models' folder.
- Point clouds output directory (green): specify the folder for storing downsampled point clouds for subsequent K-means multi-templates selection.
- Templates output directory (blue): specify the folder for storing the selected templates

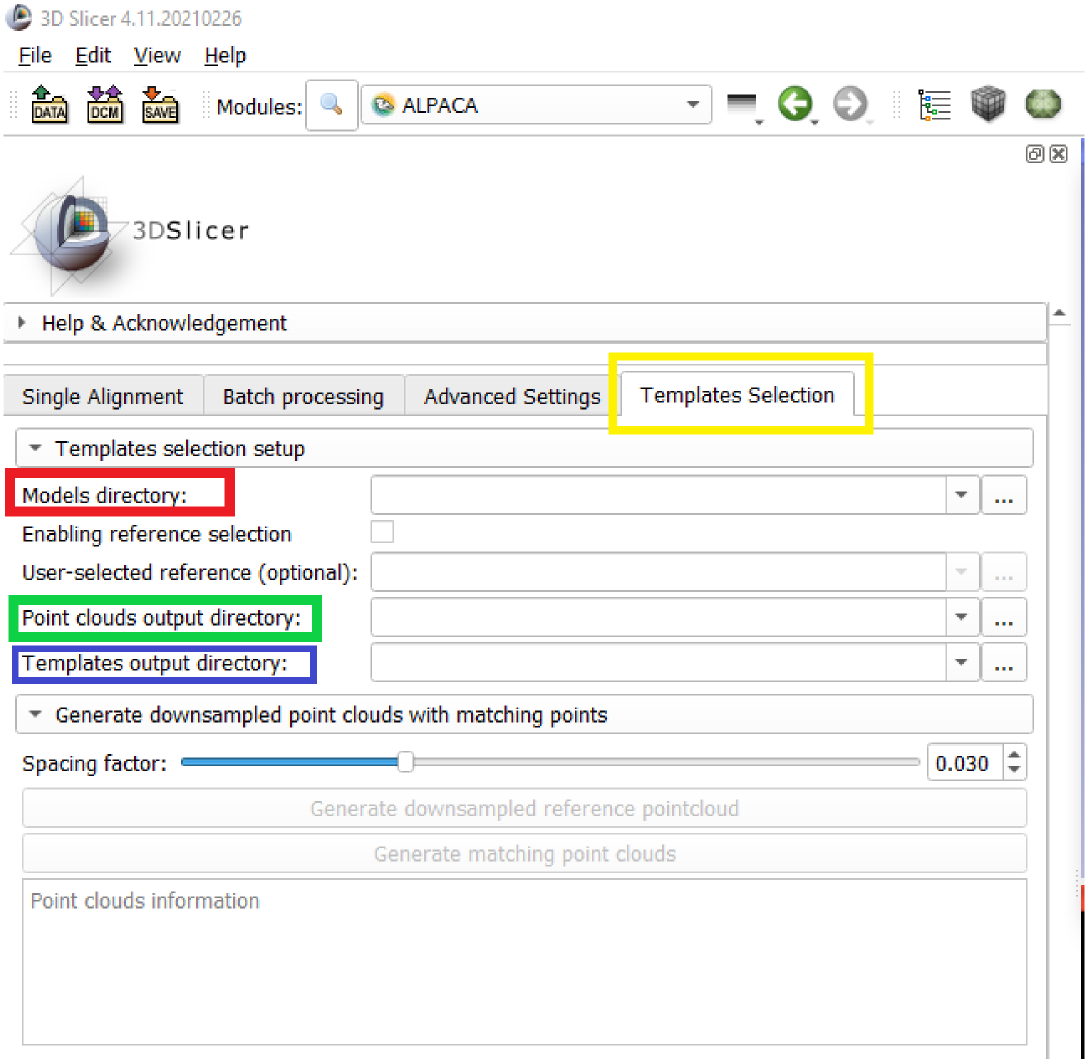

- (**Optional**) Checking the “Enabling reference selection” checkbox (red) will enable “User-selected reference” entry. This allows users to select a specimen in the “Models Directory” (red arrow) as the reference for registering specimens as well as generating downsampled point clouds with point-to-point correspondence.
    - If this checkbox is unchecked or no specific reference is selected, the first specimen in the “Models Directory” folder will serve as the reference by default.

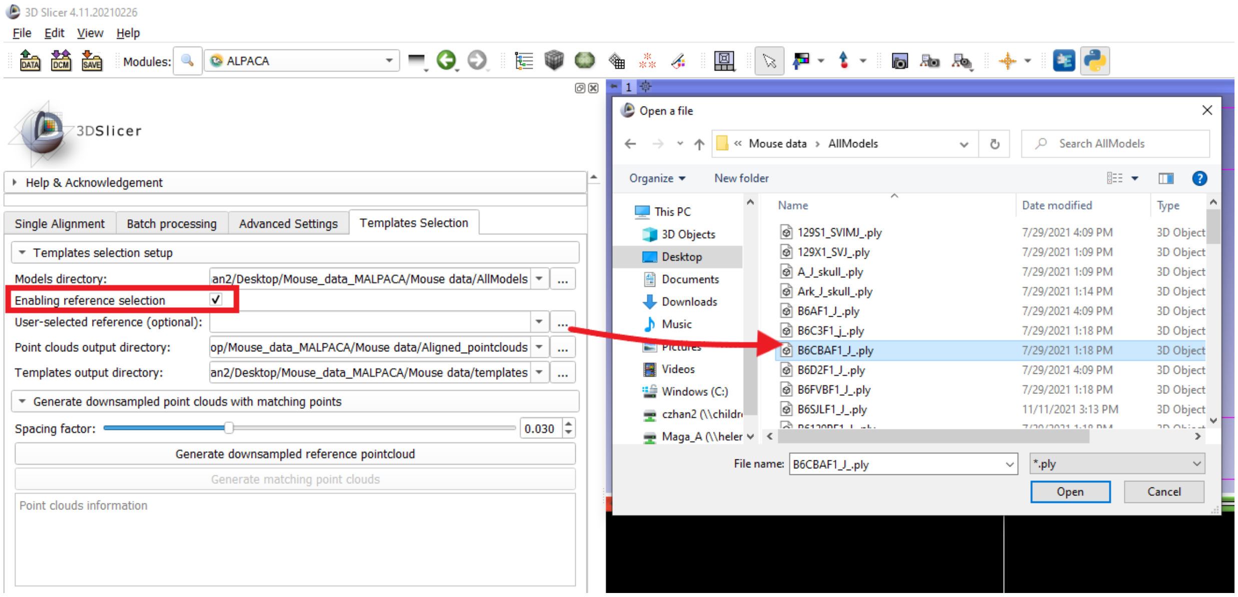


2. Generating downsampled point clouds with point-to-point correspondence.

- Specify a “Spacing factor” value using the slider (red) or simply input a value in the box at the right side. This value determines how sparse the downsampled reference point cloud will be. The larger the spacing factor value, the sparser the point cloud will be. For new samples, this requires some experimentation on user’s end (See explanation in next steps). For mouse sample spacing of 0.03 is suggested.

- Click the “Generate downsampled reference pointcloud” button (blue). After the downsampled reference point cloud is generated, the box below (green) will display information about the name of the reference model and the number of points in the reference point cloud. The “Generate matching point clouds” button will also be enabled.

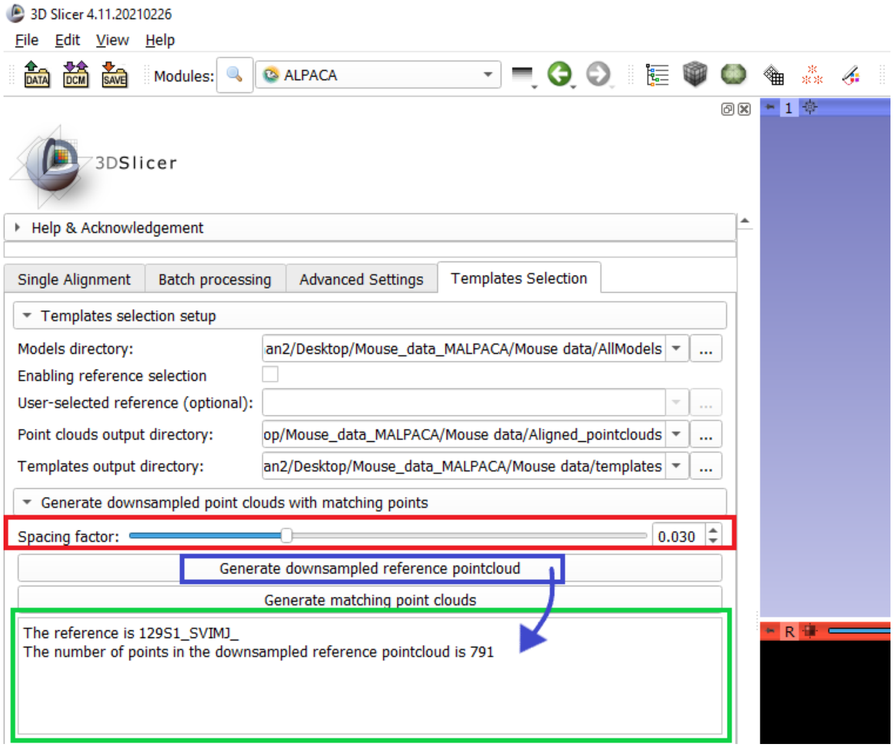


- Click the "generate matching point clouds" button ("red"). This will extract point clouds from each target specimen listed in the "Models directory" based on the reference. These point clouds with point-to-point correspondence will be stored in "fcsv" format as landmark files in the folder specified in 2) of Step 1.
  - Note that occasionally a point in the target may be repeatedly fetched if it is equally close to multiple points in the reference point cloud. Consequently, the number of unique points in a target will be smaller than that of the reference.
  - Since the number of points in a point cloud is usually large, having a few redundant points are unlikely to significantly impact results of morphometric analysis. Thus, the error threshold that measures the number of redundant points in a target point cloud is set up as 1%. For example, if the reference has 800 points and the number of unique points in the target is 793, the result will be suggested as acceptable (the error rate does not exceed the 1% threshold).
  - The box below the "Generate matching point clouds" button displays information about the number of unique points in generated point clouds, including specimens with number of unique points smaller than that of the reference and whether their error rates exceed the threshold.
  - Users can opt to ignore the warning when some point clouds have error rates exceeding 1%. If not, they can try increasing the spacing factor to achieve sparser point clouds or check possible deficiencies in the original models.

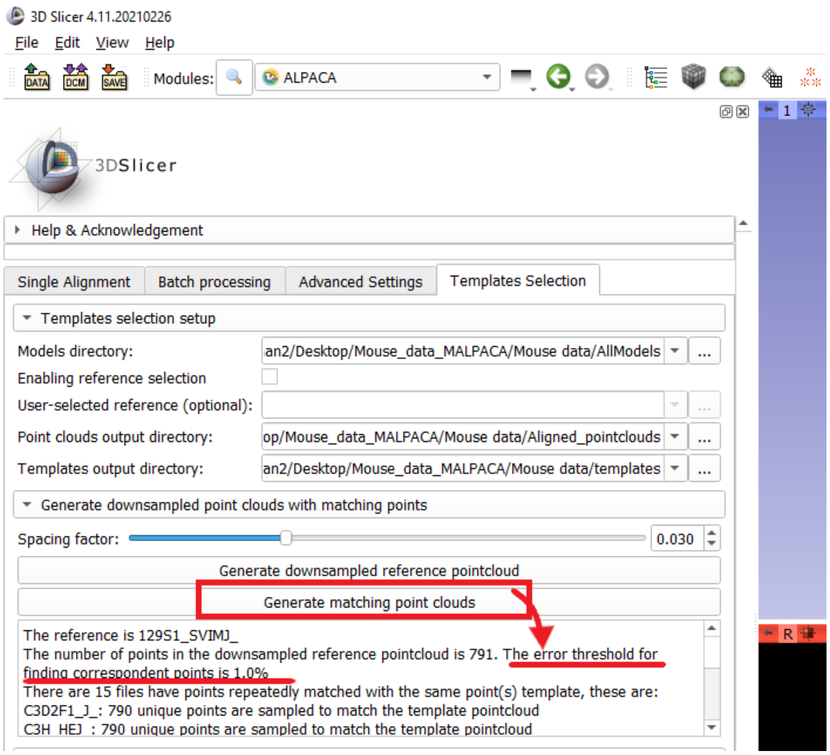

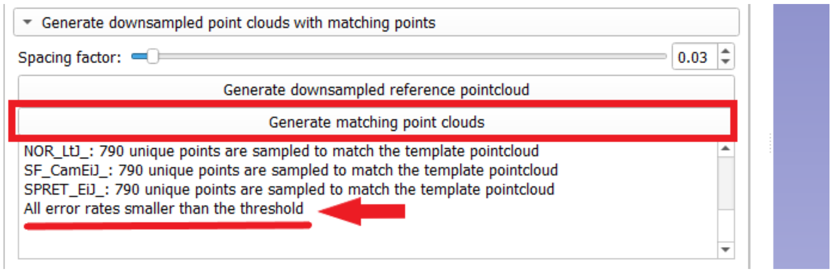


3. Specify whether there are multiple groups within the sample in the "Multi-template selection" section. If users have the fcsv files of the point cloud data in hand, they can specify the directory in the "Point clouds output directory" in Step 1 without re-generating these point clouds.

- Check "One group for the whole sample" (red arrow) if no pre-known groups exist within the sample or users do not want to divide the sample into groups. In this case, the user-input "Number of templates per group" (blue) is the number of templates for the whole sample.

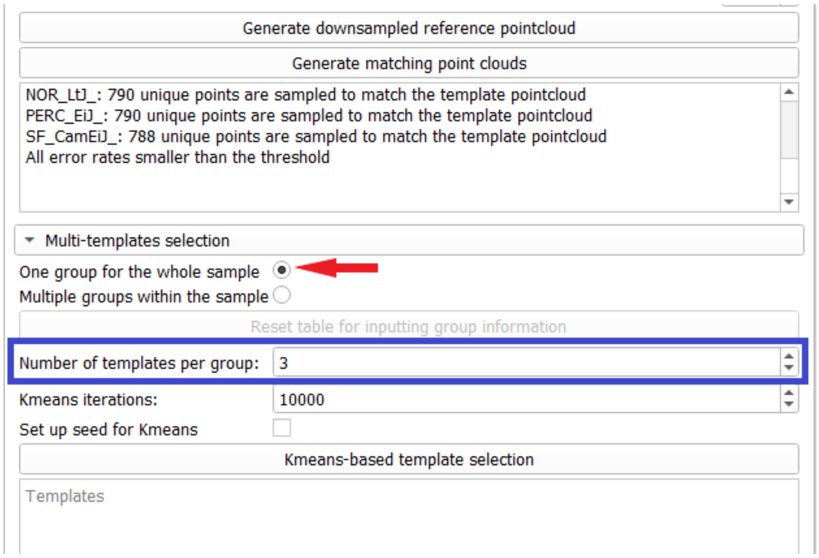


- Check the “Multiple groups within sample” (green arrow) if for dividing a sample into multiple groups based on prior knowledge. By checking this option:
  - A table for manually entering group labels for each specimen will be generated (right). A K-means algorithm will be executed per group to select a user-defined number of templates specified in “Number of Templates per group” (blue).
  - The “Reset table for inputting group information” button (yellow) will be enabled. Click it to reset the group information table.

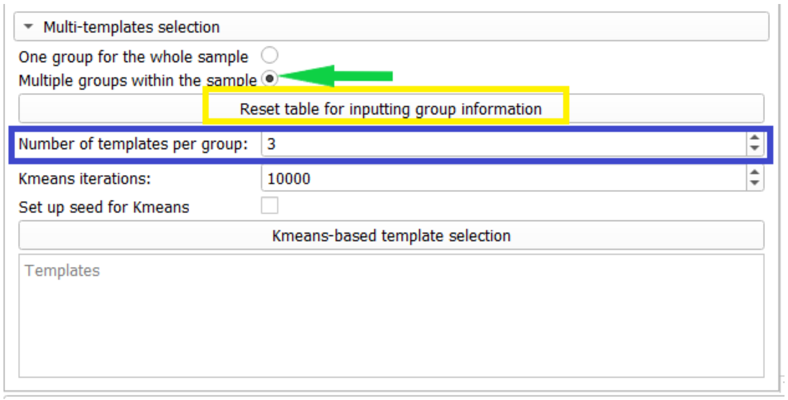


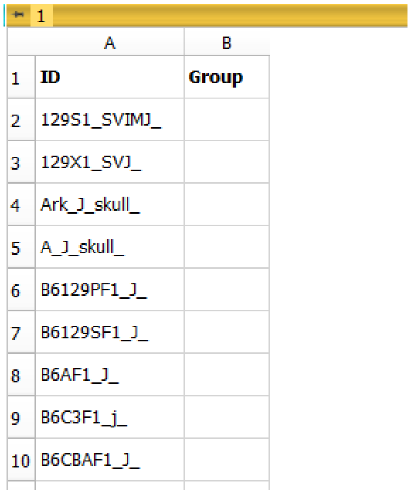

| | A | B |
|---|---|---|
| 1 | ID | Group |
| 2 | 129S1_SVIMJ_ | |
| 3 | 129X1_SVJ_ | |
| 4 | Ark_J_skull_ | |
| 5 | A_J_skull_ | |
| 6 | B6129PF1_J_ | |
| 7 | B6129SF1_J_ | |
| 8 | B6AF1_J_ | |
| 9 | B6C3F1_j_ | |
| 10 | B6CBAF1_J_ | |

4. Set up number of iterations (red) and a seed (optional) (yellow) for K-means algorithm

- The default number of K-means iterations is 10,000. Iteration number can be increased for consistent results of K-means.
- For ensuring reproducibility, users can set up a seed for K-means by checking the “Set up seed for K-means” box (blue). However, this may also influence other random-based functions in 3D Slicers. Therefore, for running other 3D Slicer modules, it is recommended to open a new Slicer session.

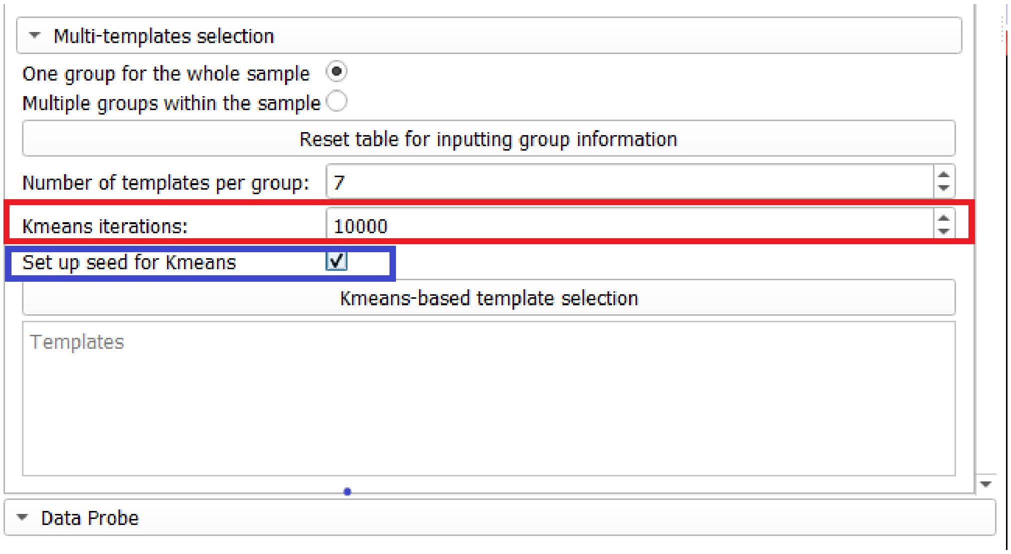


5, Click “Kmeans-based template selection” button (red) will execute K-means algorithm for multi-template selection. The models selected as templates will be saved in the folder specified in “Templates output directory” in Step 1. The display box (yellow) below shows specimens selected as the templates.

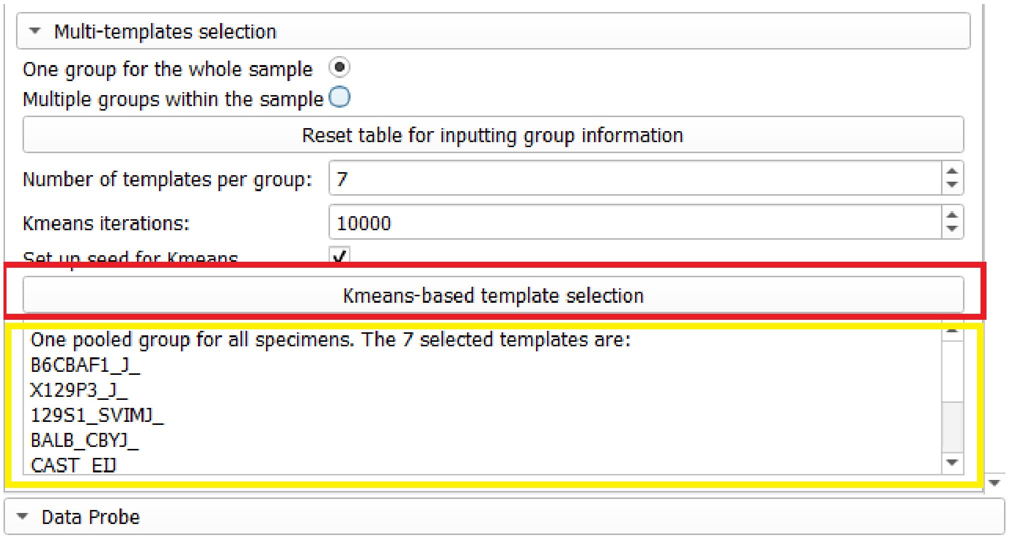


- If “One group for the whole sample” is checked in step 4, a plot of PC1 and PC 2 based on a Generalized Procrustes Analysis of the point cloud data will be displayed. Specimens are black squares. Templates are light color, diamond-shaped points that overlay the squares.

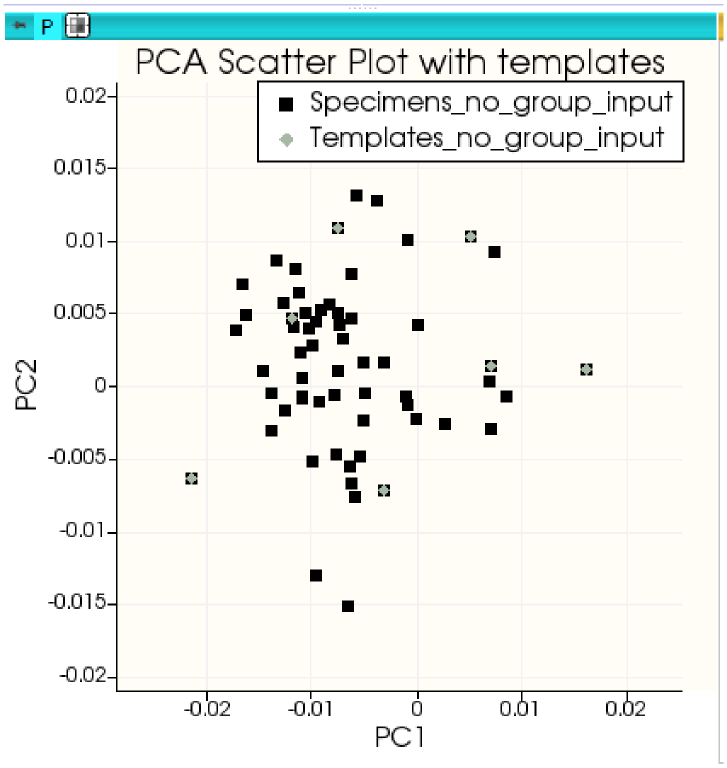


- If "Multiple groups within the sample" is checked (green arrow) and group identity for each specimen is entered, the display box (yellow) will show selected templates for each entered group. The PC plot will also display specimens in each user-defined group in a unique color.

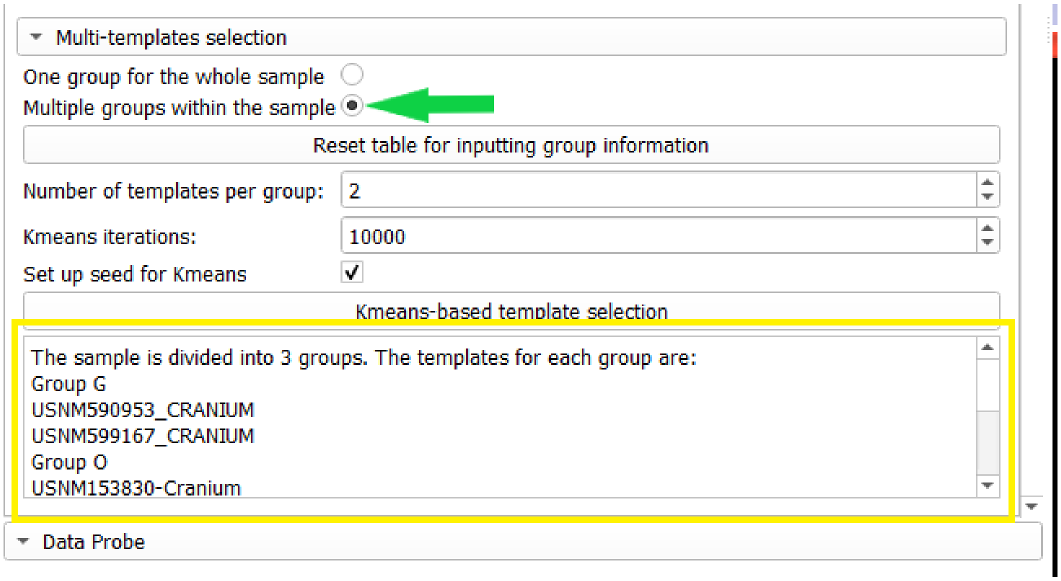

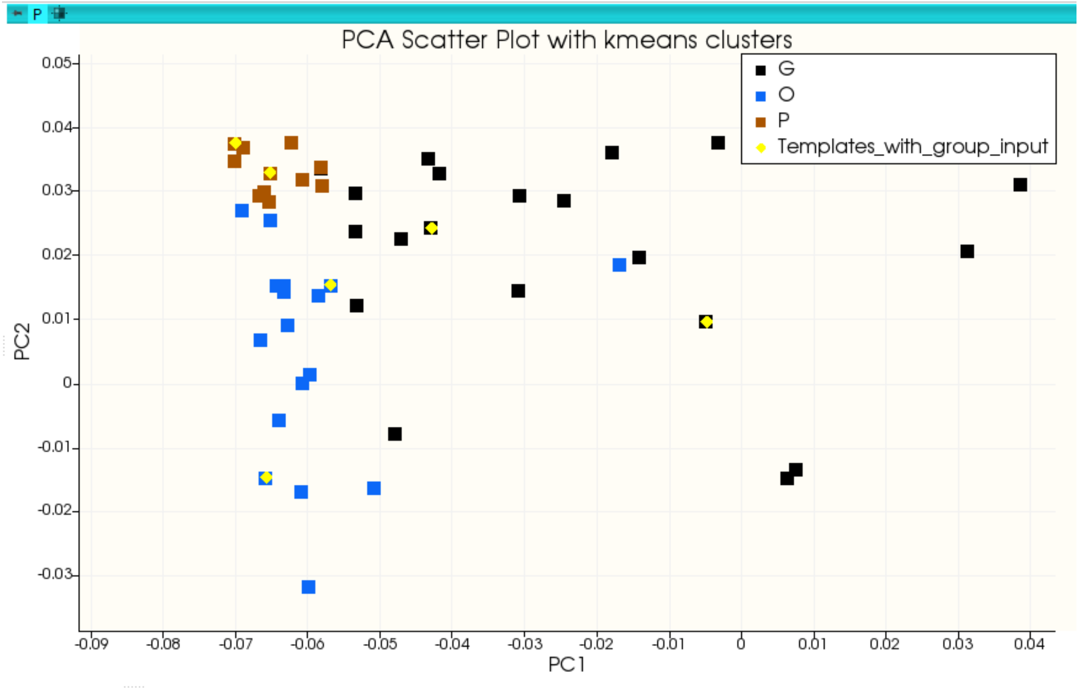
PCA Scatter Plot with kmeans clusters
G
O
P
Templates_with_group_input
PC1
PC2
0.05
0.04
0.03
0.02
0.01
0
-0.01
-0.02
-0.03
-0.09
-0.08
-0.07
-0.06
-0.05
-0.04
-0.03
-0.02
-0.01
0
0.01
0.02
0.03
0.04

#### II. Executing ALPACA in Multi-Template mode (MALPACA)

6. Switch to the ALPCA module and choose the Batch Processing tab (red). In the "Method" entry (dark blue), open the drop-down menu to select **"Multi-Template (MALPACA)"** option from the dropdown menu.

7. Select required input and output directories.

- In the "Source mesh(es)" entry (yellow), select the folder that contains .ply files of the templates.
- In the "Source landmarks" entry (green), select the folder that contains fcsv landmark files for the templates. The names of the template model and landmark files must be identical.
- In the "Target mesh directory" entry (dark grey), select the folder that contains .ply files of the target meshes. These are the specimens to be landmarked by MALPCA.
- In the "Target output landmark directory" (light blue), select the folder for MALPACA output.

8. Click the "Run auto-landmarking" button (red arrow) to execute MALPACA. Slicer may appear to be in the "no response" condition. This is because MALPACA is running.

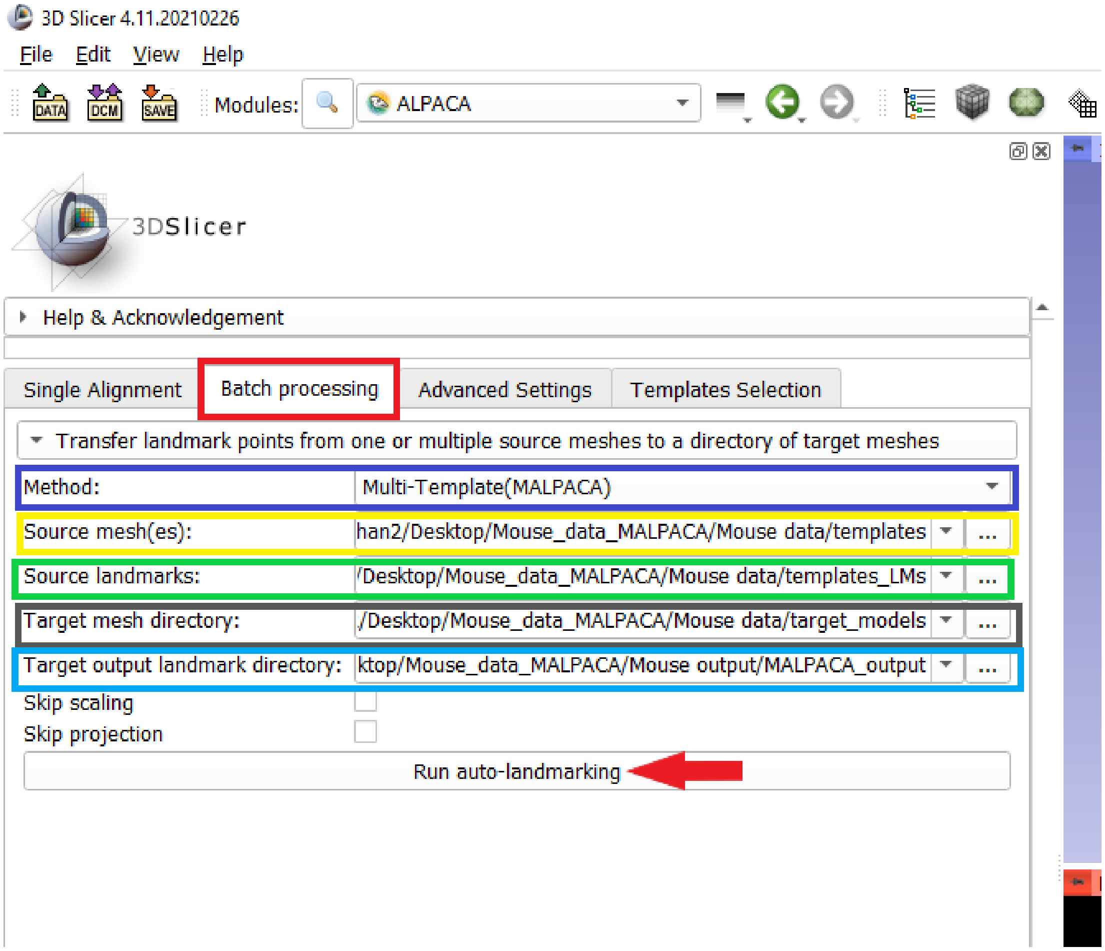

9. Open the target output landmark directory specified in Step 7 that stores the MALPACA output.

- The "advancedparameters.txt" file stores the MALPACA settings.

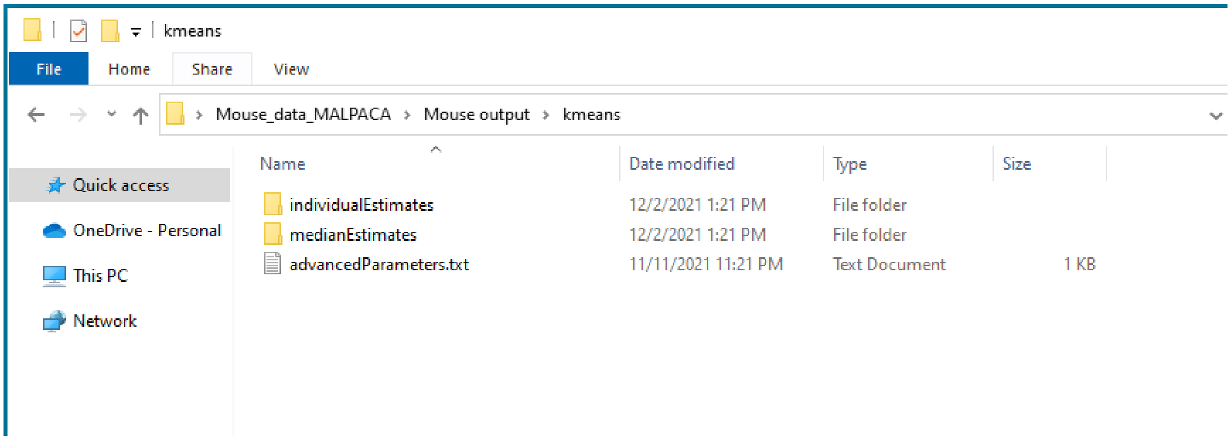


- The "individual estimates" folder contains landmarks estimated by each individual template stored in the fcsv format. For each file name, the postfix is the template that generate this estimated landmark file. For example, "129X1_SVJ_B6CBAF1" suggests that the estimated landmarks of the specimen 129X1_SVJ is derived from using the template B6CBAF1.

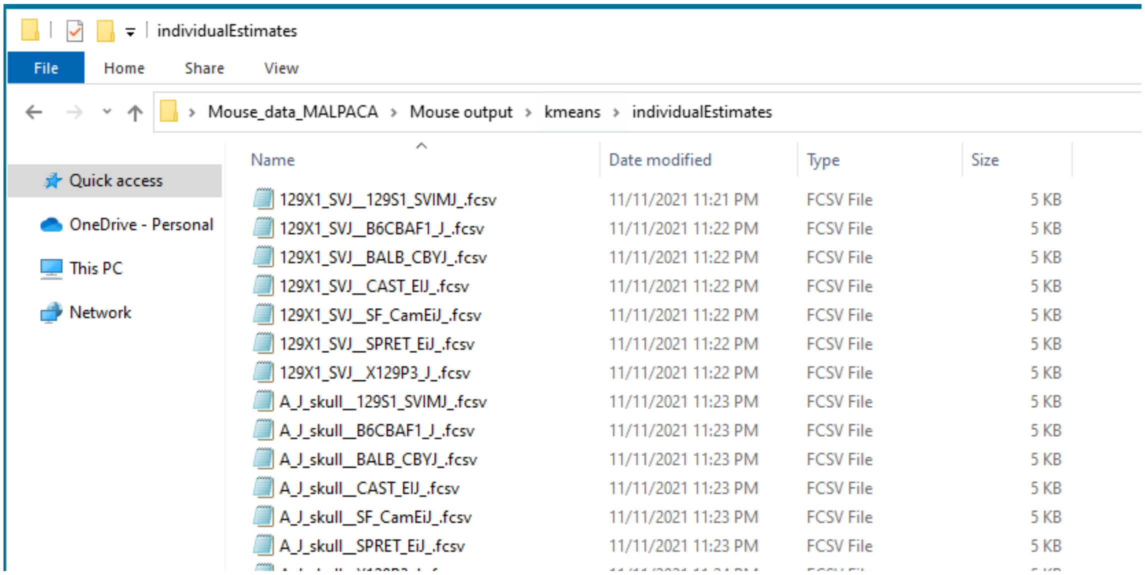


- The "medianEstimates" folder contains the final output of MALPACA storing in fcsv format. Each landmark file name has a suffix "_median", suggesting it is the median of the estimates derived from using all templates.

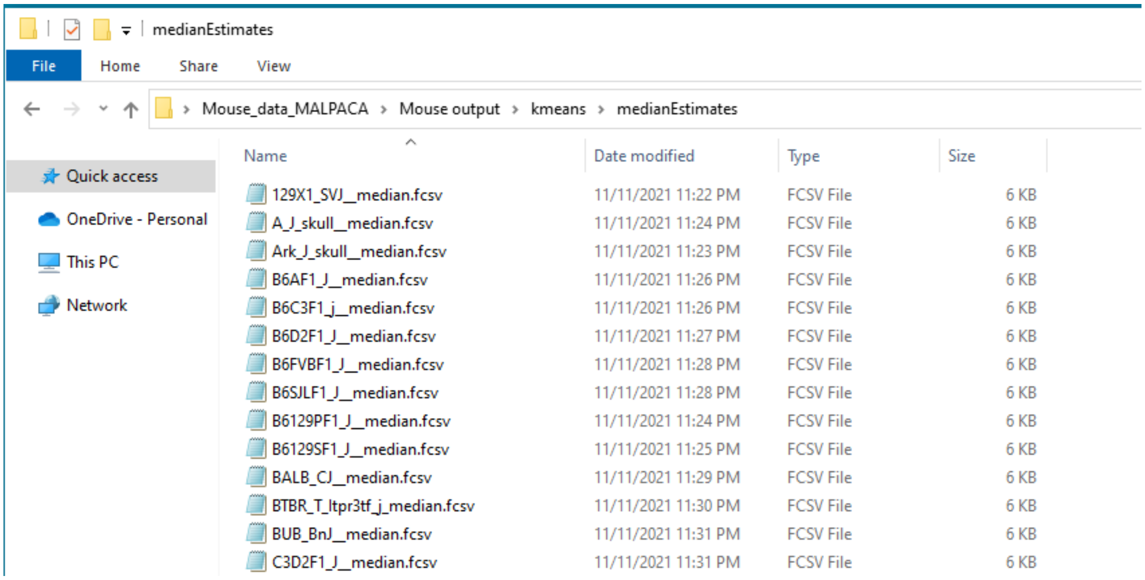
medianEstimates
File
Home
Share
View
Mouse_data_MALPACA
Mouse output
kmeans
medianEstimates
Quick access
OneDrive - Personal
This PC
Network
Name
Date modified
Type
Size
129X1_SVJ__median.fcsv 11/11/2021 11:22 PM FCSV File 6 KB
A_J_skull__median.fcsv 11/11/2021 11:24 PM FCSV File 6 KB
Ark_J_skull__median.fcsv 11/11/2021 11:23 PM FCSV File 6 KB
B6AF1_J__median.fcsv 11/11/2021 11:26 PM FCSV File 6 KB
B6C3F1_j__median.fcsv 11/11/2021 11:26 PM FCSV File 6 KB
B6D2F1_J__median.fcsv 11/11/2021 11:27 PM FCSV File 6 KB
B6FVBF1_J__median.fcsv 11/11/2021 11:28 PM FCSV File 6 KB
B6SJLF1_J__median.fcsv 11/11/2021 11:28 PM FCSV File 6 KB
B6129PF1_J__median.fcsv 11/11/2021 11:24 PM FCSV File 6 KB
B6129SF1_J__median.fcsv 11/11/2021 11:25 PM FCSV File 6 KB
BALB_CJ__median.fcsv 11/11/2021 11:29 PM FCSV File 6 KB
BTBR_T_Itpr3tf_j_median.fcsv 11/11/2021 11:30 PM FCSV File 6 KB
BUB_BnJ__median.fcsv 11/11/2021 11:31 PM FCSV File 6 KB
C3D2F1_J__median.fcsv 11/11/2021 11:31 PM FCSV File 6 KB

#### III. Landmarking Quality Control Pipeline (OPTIONAL)

One of the benefits of multi-template pipeline is the ability to assess confidence in the estimates. Here we demonstrate a pipeline based on a simple heuristic (mean RMSE + 2.SD RMSE) to flag individual estimates as potential outliers. It should be noted that because MALPACA is using median to obtain final LM positions, it is robust to an occasional outlier (i.e., removal of the outlier and re-estimate of the position should have no appreciable effect on final LM estimate). However, in cases where an ill-chosen template produces consistently poor results, diagnostic plots like the one shown below would be helpful to identify the problem (abundance of LMs above threshold for that template) and re-run the MALPACA pipeline by replacing the template by another sample, or simply re-estimating without the outlier.

10. Open R or RStudio console, install SlicerMorphR package using the code devtools::install_github('chz31/SlicerMorphR'), then run 'library(SlicerMorphR)' to load the package.

11. To try the functions for extracting outliers, users can load the ape MALPACA data from Github, either by clone the repository or using the following code to download the zip file and unzip it:

> save.dir='C:/tmp'  #enter your directory to download the MALPACA data sample
> setwd(save.dir)
> zip_url <- "https://github.com/SlicerMorph/Mouse_Models/archive/refs/heads/main.zip"
> download.file(url = zip_url, destfile = "Mouse_Models.zip")
> unzip(zipfile = "Mouse_Models.zip")

The following steps use the sample ape data loaded in Step 3. Users can switch to their own data.

12. Set up MALPACA output path and the directory that stores the template landmark or model files (specified in step 7 if using your own data):

> MAL_outDir_git<- paste(save.dir, 'Mouse_Models-main/ape_malpaca', sep="/")  #Set up MALPACA output path
> templates_path_git<- paste(save.dir, 'Mouse_Models-main/ape_malpaca/templates_LMs', sep='/')  #Set up

13. Read all median estimates and estimates derived from individual templates.

> LMs <- read.malpaca.estimates(MALPACA_outputDir = MAL_outDir_git, templates_Dir = templates_path_git)

> allEst <- LMs$allEstimates #Store all individual estimates in a 4D array [i, j, k, n]. i = number of templates; j = dimension; k = number of templates; n = sample size. For example:

```
> allEst[1:2, , 1, 3] #The LM1 and LM2 estimated by template 1 for target specimen 3
          x       y       z
1 -15.37080 428.172 159.411
2  -1.88649 365.697 180.744
```

> MAL_medians <- LMs$MALPACA_medians #Store all median estimates in a 3D array [i, j, n]. i = number of templates; j = dimension; n = sample size. For example:

```
> MAL_medians[1:2, , 3] #The median estimates for LM 1 and LM3 of the target specimen 3
          x        y        z
1 -7.218665 417.9610 158.4400
2 -1.850612 366.3141 180.6046
```

14. Use the 'extract.outliers()' function in SlicerMorphR to find and remove outliers from estimates derived from individual templates. This function calculates the distance between each landmark estimated by an individual template to its corresponding MALPACA median estimate. If the distance exceeds the threshold, then this estimate is marked as an outlier. By default, the threshold is 2 × (standard deviation) + mean of the pooled distances for each specimen. 'NA' is then assigned to its coordinates in the 4D array that stores all estimates of individual templates. For the parameters:

- 'allEstimates' is the 4D array that stores all estimates of individual templates, generated by 'read.malpca.estimates()'
- 'MALPACA_medians' is the 3D array that stores MALPACA median estimates generated by 'read.malpaca.estimates()'.
- 'outputPath' specifies the output directory for storing output the pdf files that contains the boxplots of distances between each individual estimate to its corresponding MALPACA median estimate.
- 'ZScore' specify the threshold for determining outliers. The default value is 2. This means that the threshold is (2  (standard deviation) + mean) of pooled distances between all pairs of individual estimates and the corresponding MALPACA median estimates.
- For the ape sample data, run:
  > 'outliers <- extract.outliers(allEstimates = allEst, MALPACA_medians = MAL_medians, outputPath = outPath)'
- Results using the ape sample data are shown below. The boxplot shows the distances between every individual landmark estimate and its corresponding MALPACA median output for one ape specimen. Each column of dots represents the distances for the estimates of the six ape templates. The red horizontal line represents the threshold. Points above the threshold are outlier estimates. For the pdf file in the 'outputPath', each page represents a boxplot for one target specimen. The legends are at the first page.

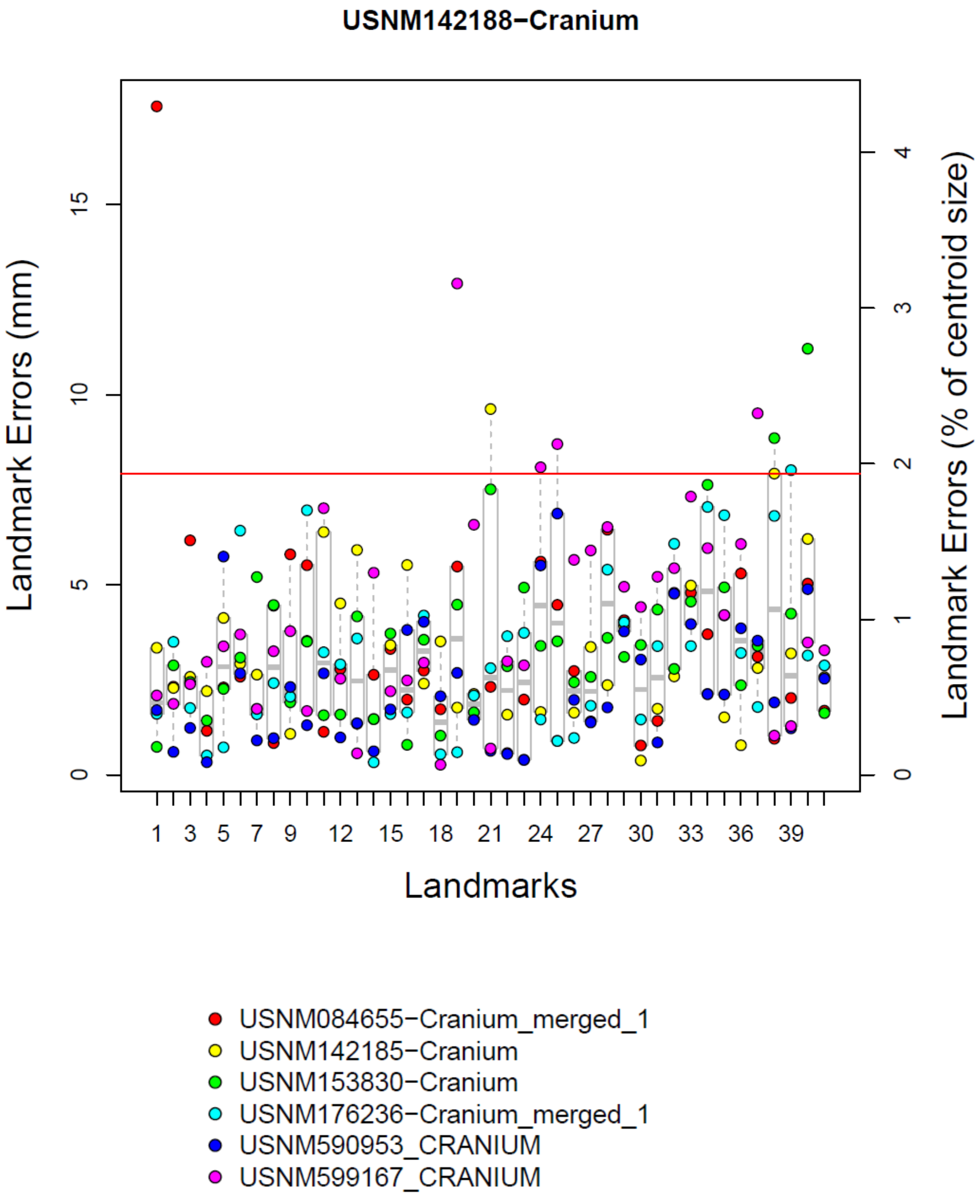

USNM142188−Cranium
Landmark Errors (mm)
Landmark Errors (% of centroid size)
Landmarks
0
5
10
15
1
2
3
4
1
3
5
7
9
12
15
18
21
24
27
30
33
36
39
USNM084655−Cranium_merged_1
USNM142185−Cranium
USNM153830−Cranium
USNM176236−Cranium_merged_1
USNM590953_CRANIUM
USNM599167_CRANIUM

```
> outliers$estimates_no_out[18:23, , 2, 7]
           x        y        z
18   3.123262 441.5676 226.7014
19   1.254022 417.1126 206.4035
20  -2.445040 385.6000 193.9190
21         NA       NA       NA
22 -17.194445 457.1618 238.1673
23  28.504499 455.5570 234.8030
> #Show LM18 to LM23 estimated by template 2 for the 7th target specimen.
> #Note the coordinates of LM21 are marked as NA because it is an outlier
```

```
> outliers$outlier_info[[7]]
$`USNM084655-Cranium_merged_1`
NULL

$`USNM142185-Cranium`
[1] 21 39

$`USNM153830-Cranium`
[1] 21

$`USNM176236-Cranium_merged_1`
NULL

$USNM590953_CRANIUM
[1] 19 36 38

$USNM599167_CRANIUM
[1] 19 25 36 37

>
> #Print out the outlier LMs generated by each template for the 7th target specimen
> #"NULL" means a template generates no outlier estimate
```

15. Use the "get.medians()" function from SlicerMorphR to generate new median estimates for each specimen, storing in a 3D array [i, j, n]. i = number of templates; j = dimension; n = sample size.

```
> medians <- get.medians(AllLMs = outliers$estimates_no_out, outlier.NA = TRUE)
> #AllLMs = input 4D array that stores estimates of individual templates
> #outlier.NA by default is False. Setting up 'True' specifies there is NA in the input 4D array.
```
