## Supplemental Figure 6 for "Automated Landmarking via Multiple Templates"

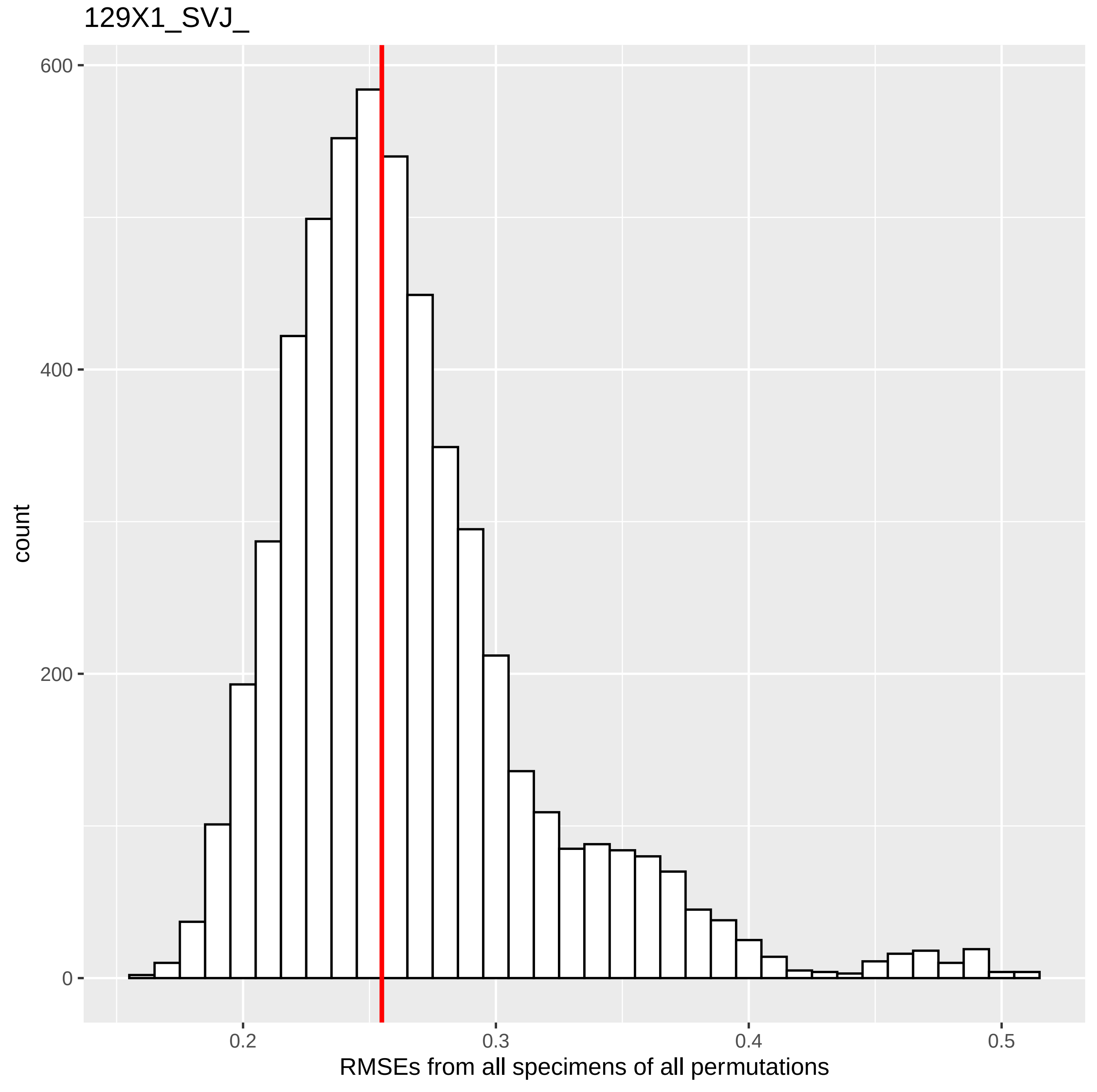
129X1_SVJ_
600
400
200
0
count
0.2
0.3
0.4
0.5
RMSEs from all specimens of all permutations

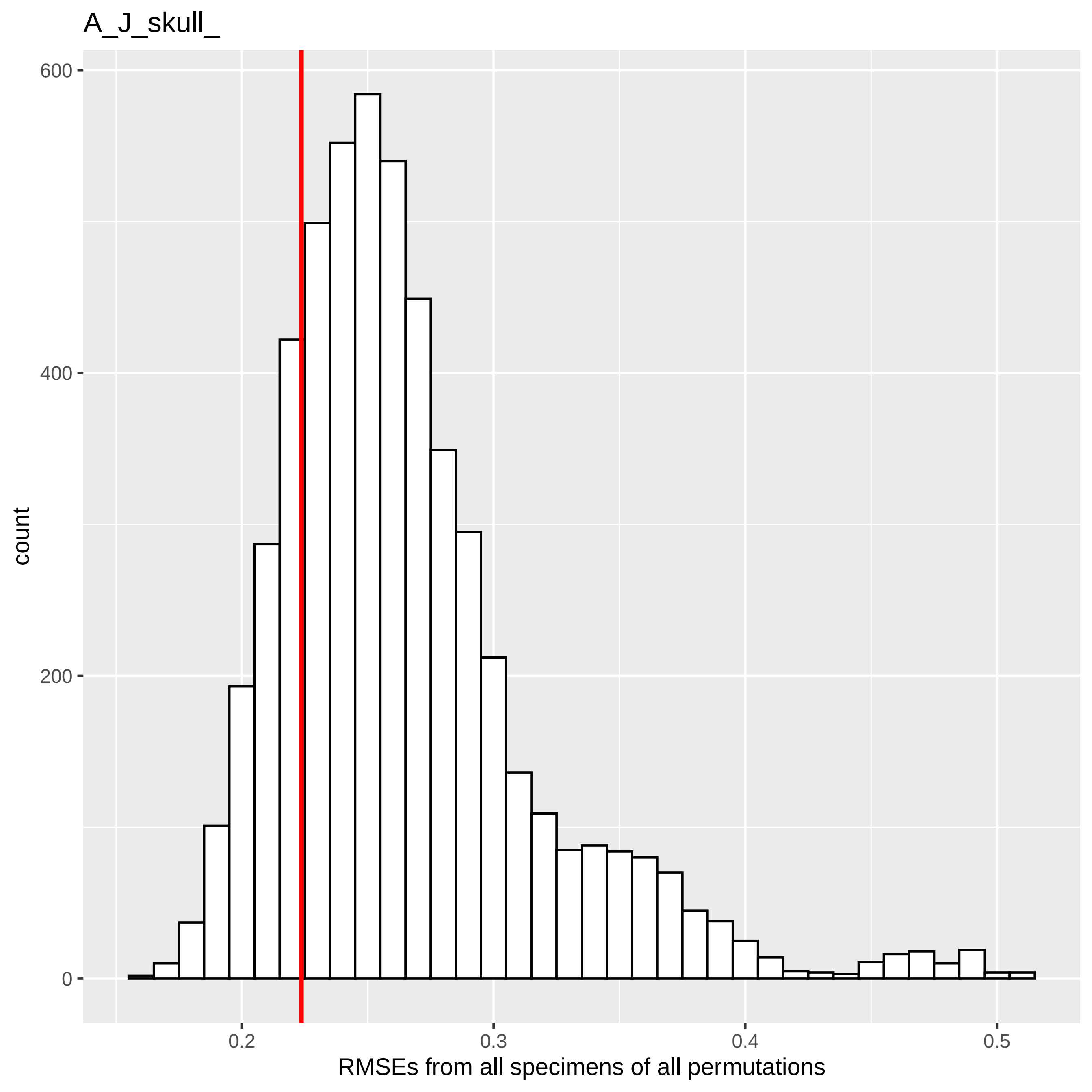
A_J_skull_
count
600
400
200
0
0.2
0.3
0.4
0.5
RMSEs from all specimens of all permutations

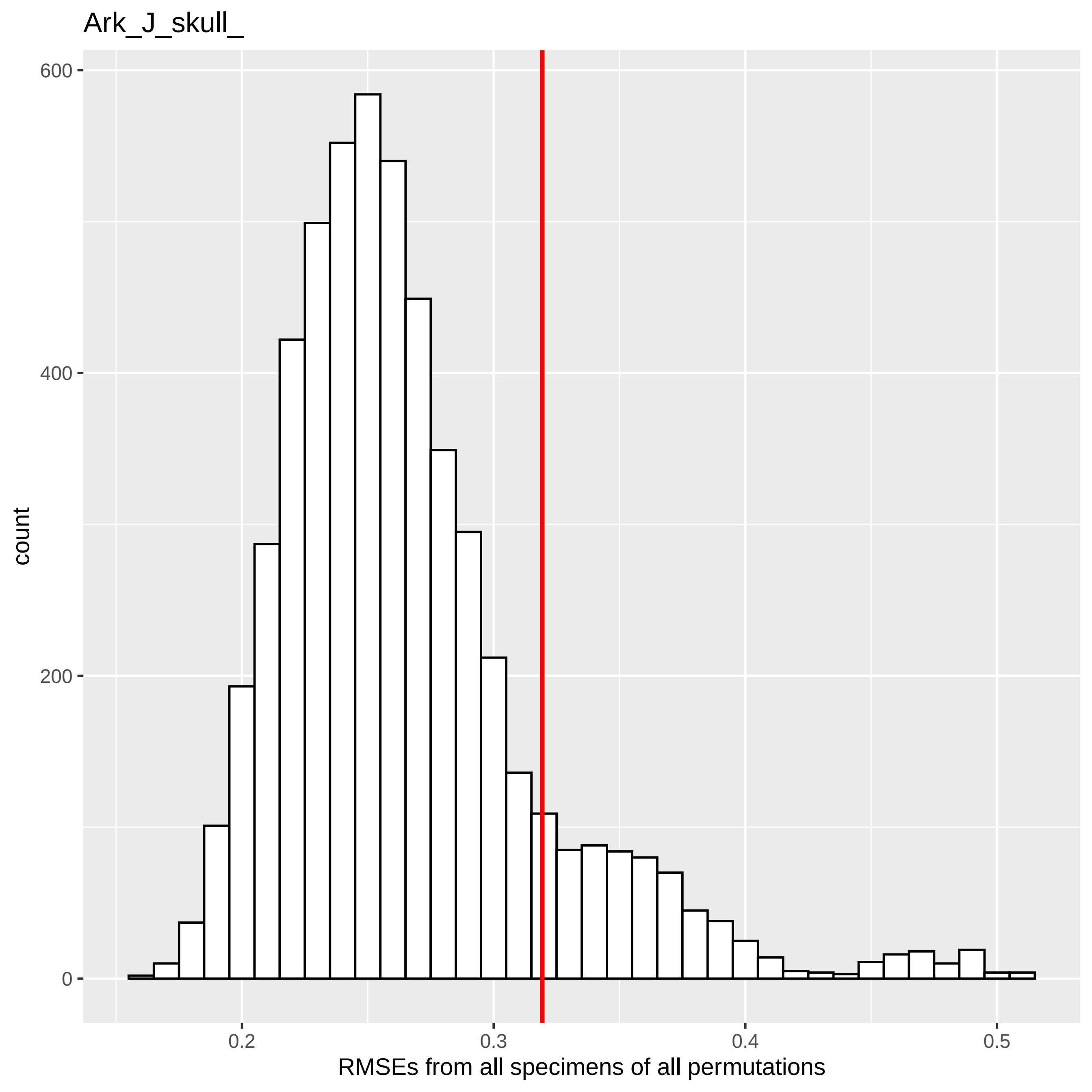

Ark_J_skull_
600
400
200
0
count
0.2
0.3
0.4
0.5
RMSEs from all specimens of all permutations

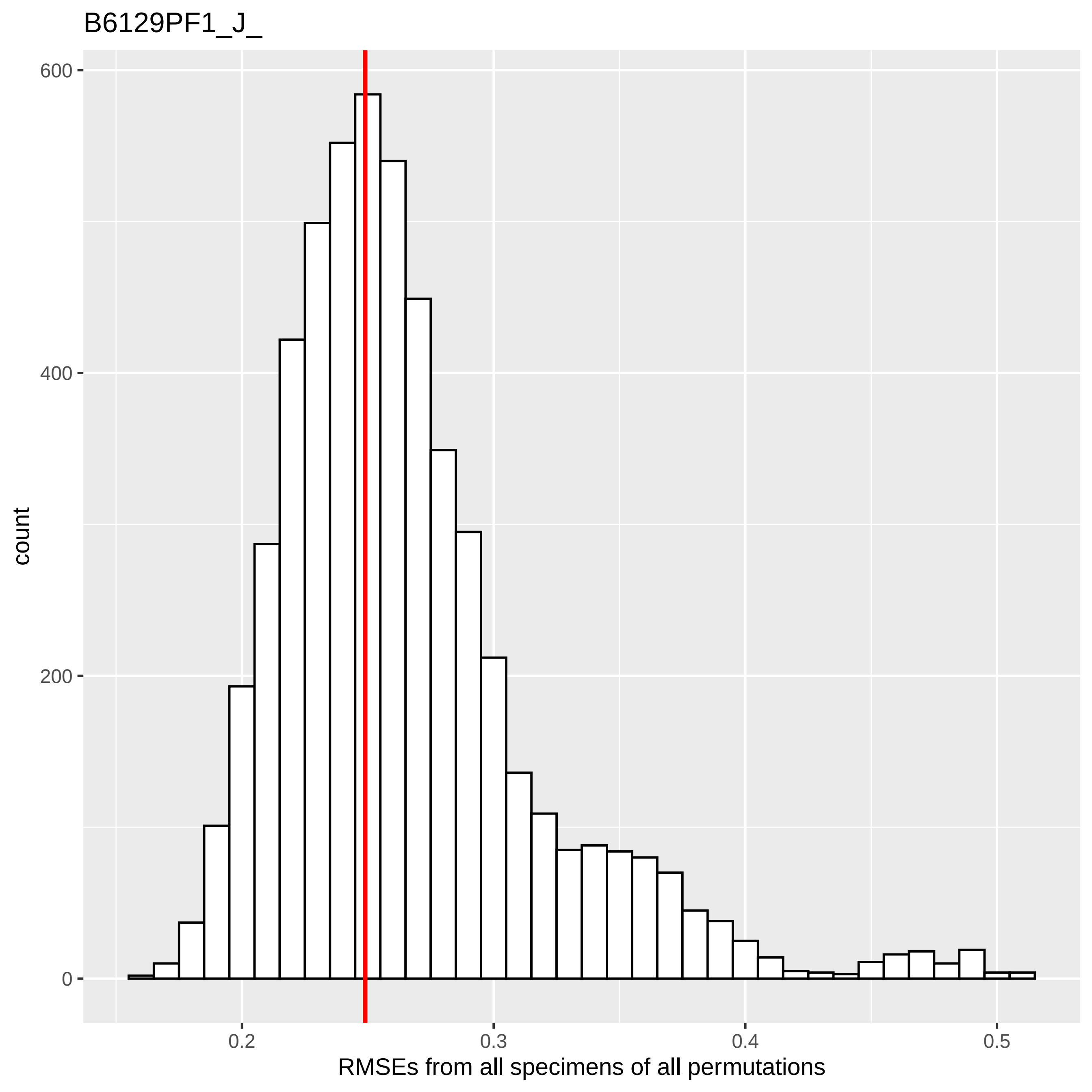
B6129PF1_J_
600
400
200
0
count
0.2
0.3
0.4
0.5
RMSEs from all specimens of all permutations

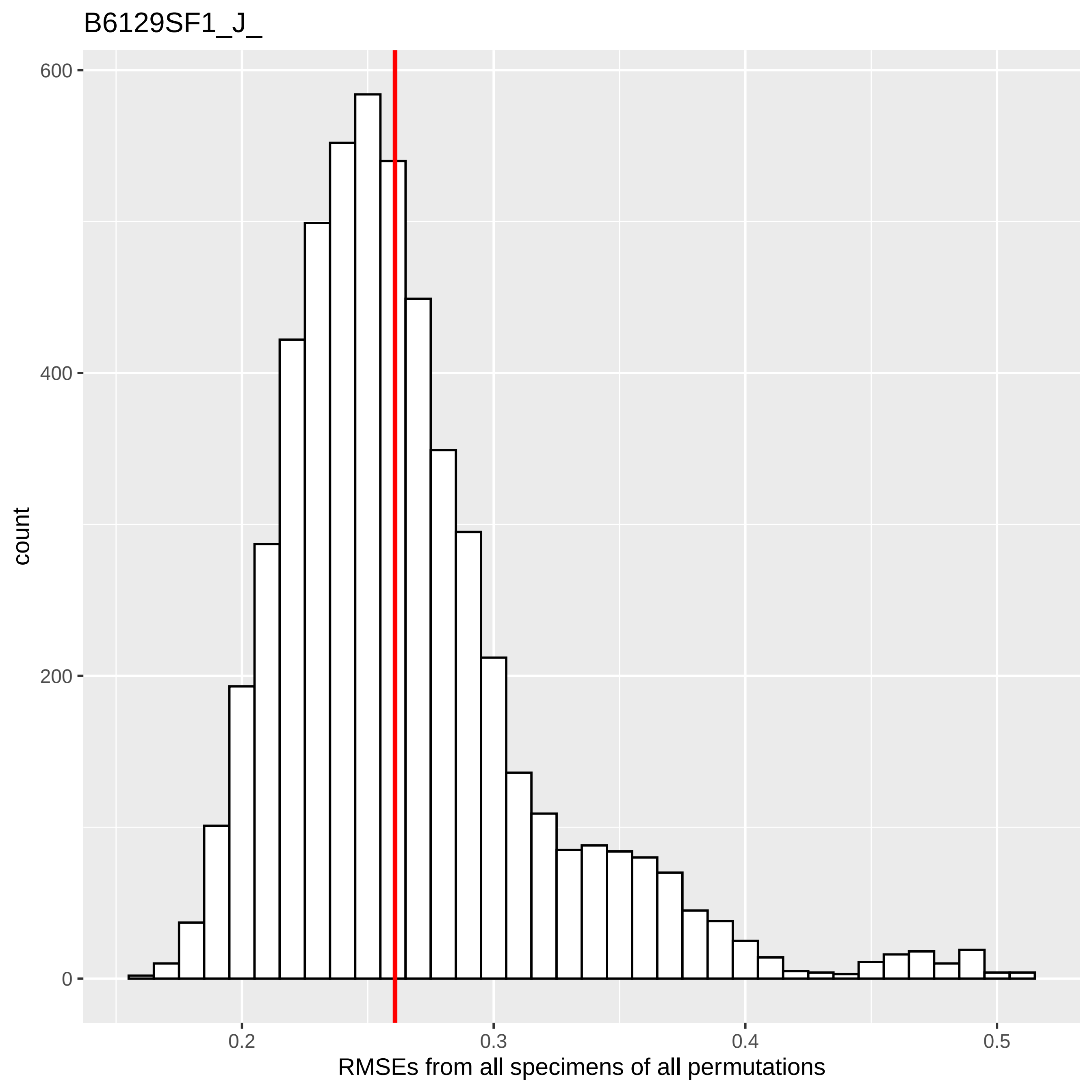

B6129SF1_J_
600
400
200
0
count
0.2
0.3
0.4
0.5
RMSEs from all specimens of all permutations

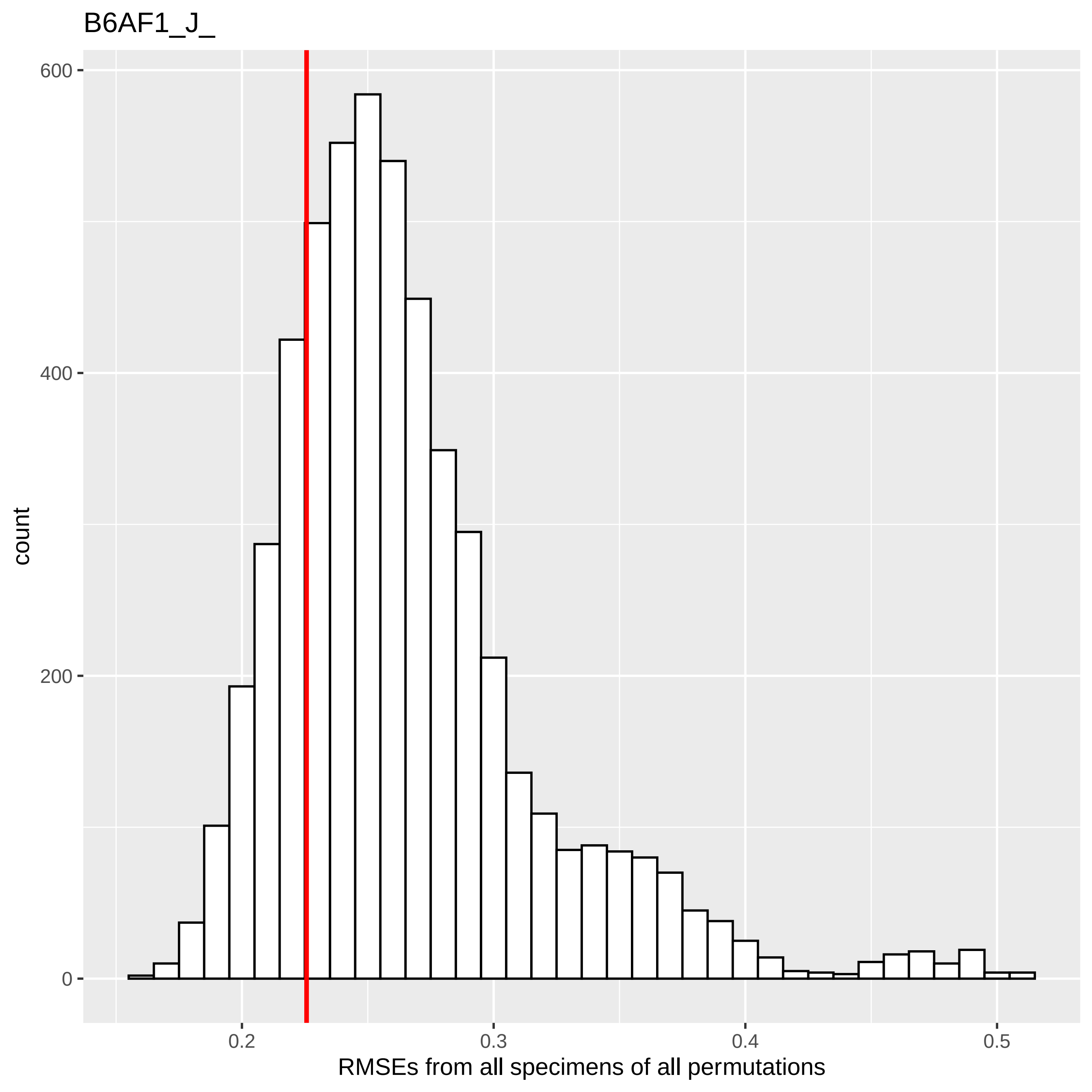
B6AF1_J_
count
600
400
200
0
0.2
0.3
0.4
0.5
RMSEs from all specimens of all permutations

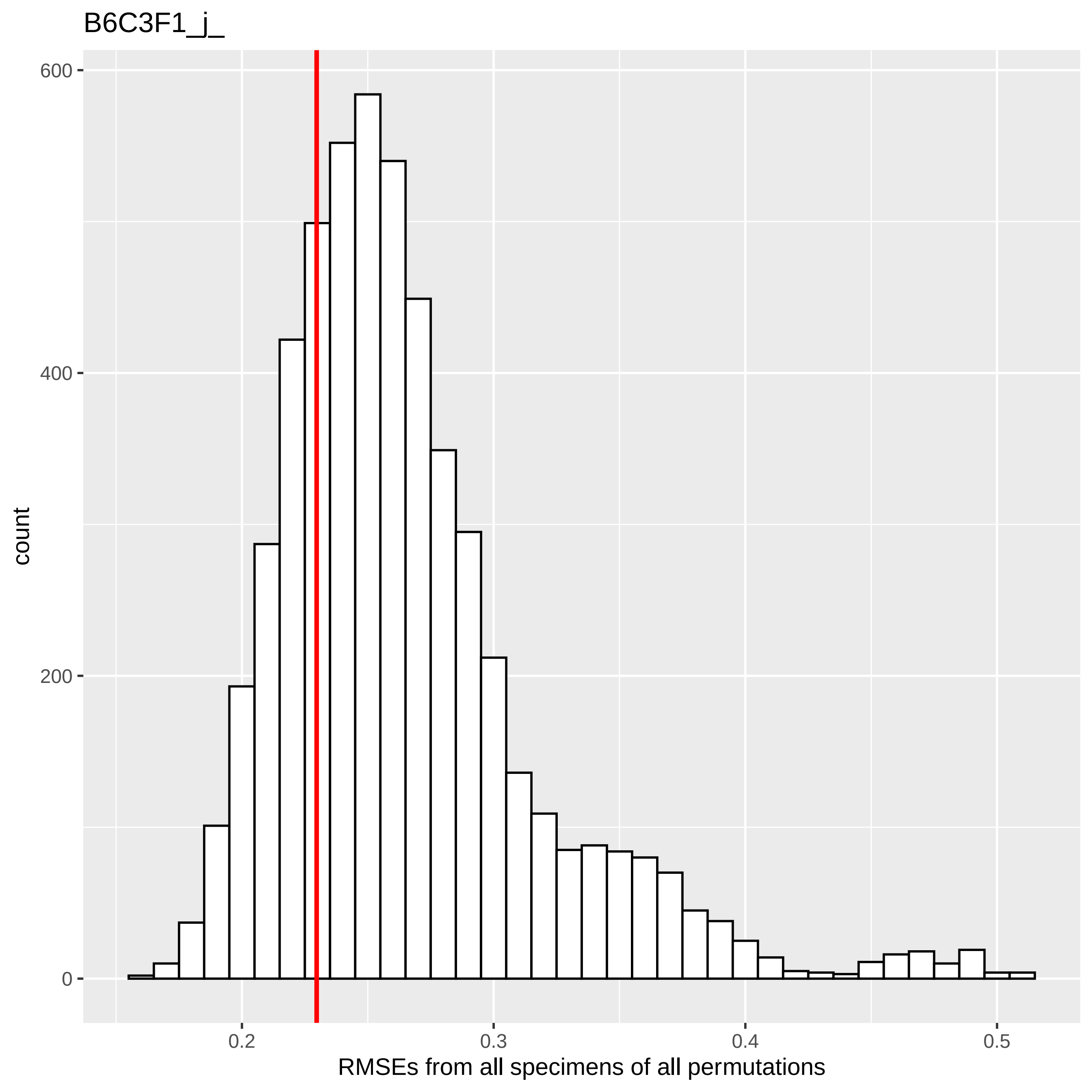
B6C3F1_j_
600
400
200
0
count
0.2
0.3
0.4
0.5
RMSEs from all specimens of all permutations

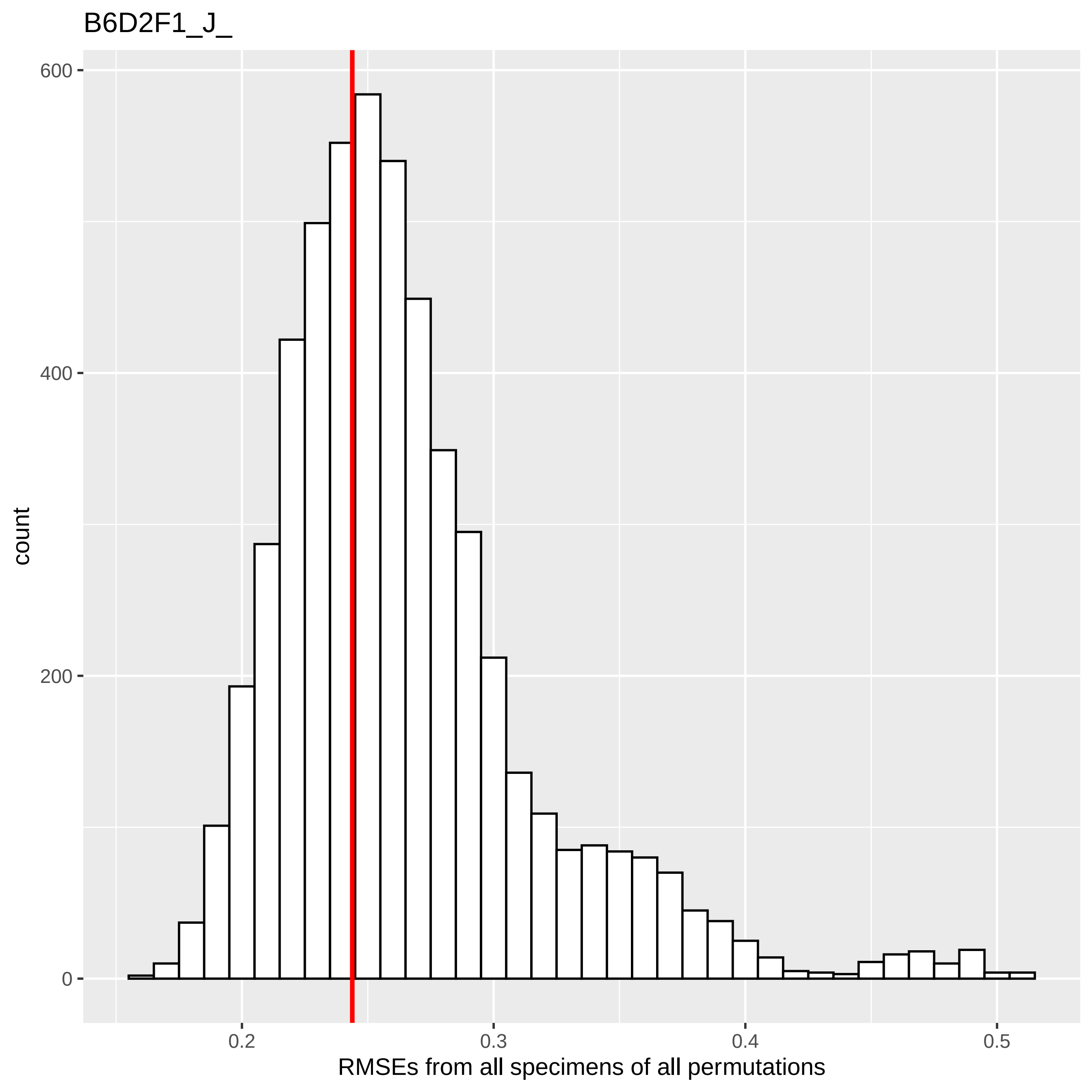
B6D2F1_J_
600
400
200
0
count
0.2
0.3
0.4
0.5
RMSEs from all specimens of all permutations

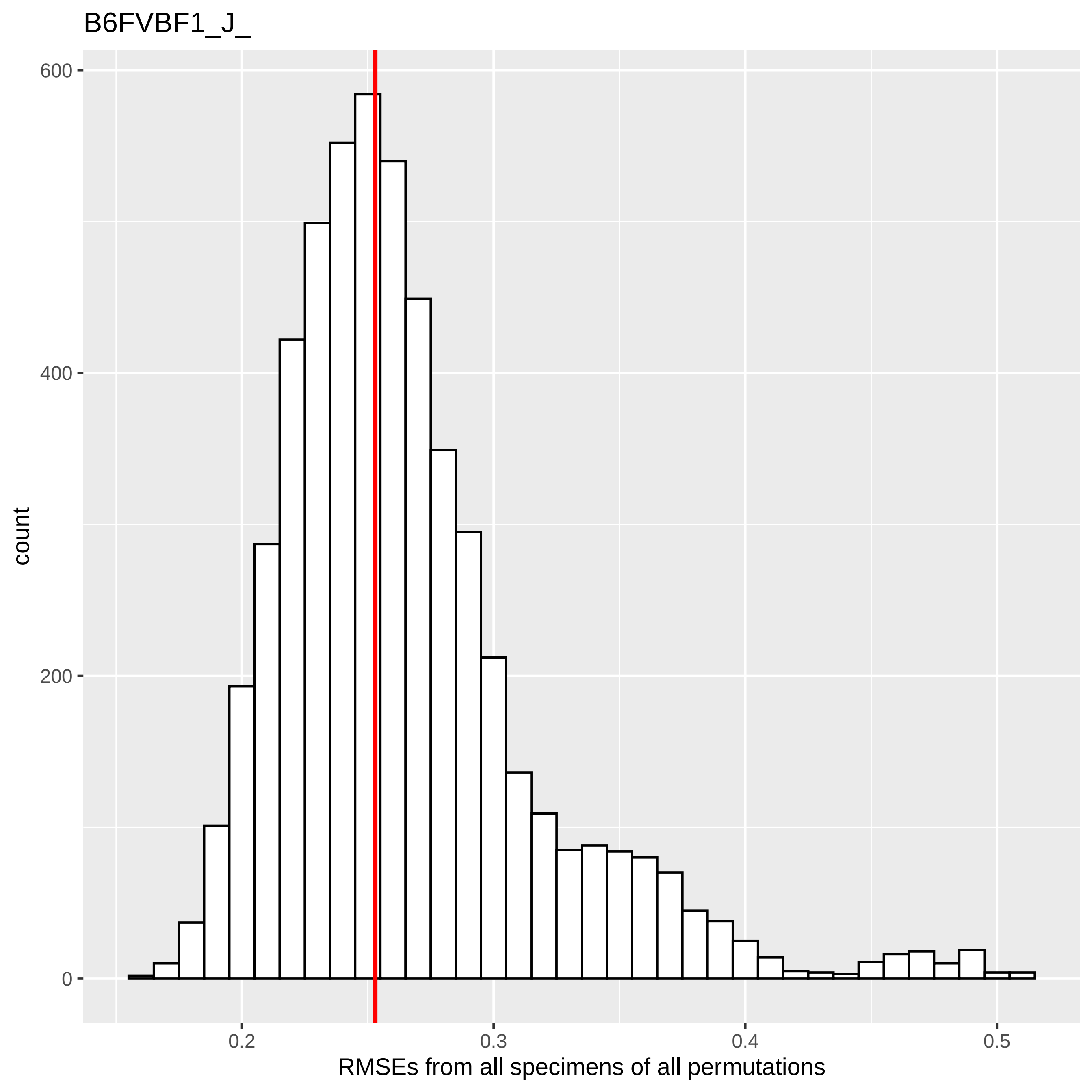
B6FVBF1_J_
count
RMSEs from all specimens of all permutations
0
200
400
600
0.2
0.3
0.4
0.5

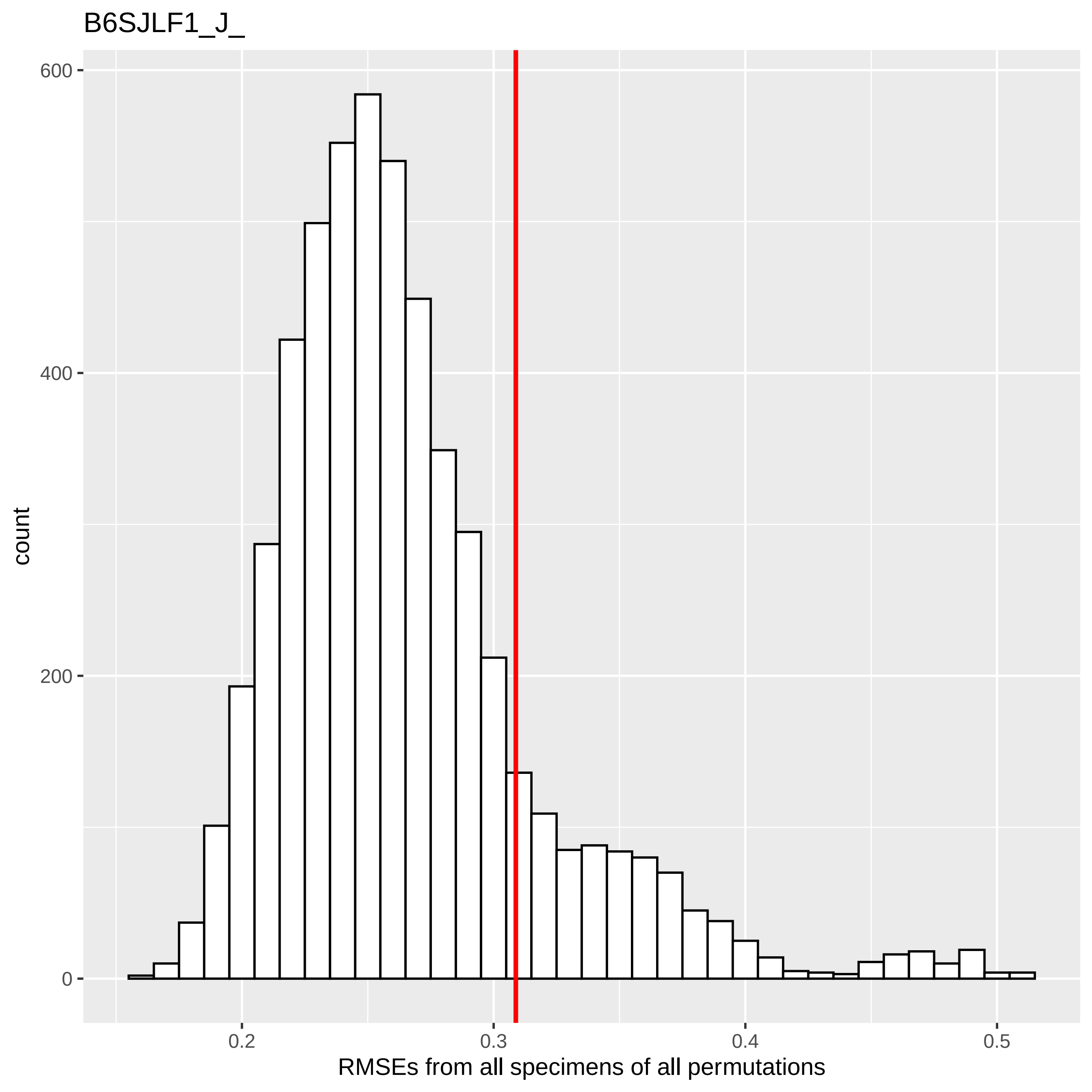
B6SJLF1_J_
count
600
400
200
0
0.2
0.3
0.4
0.5
RMSEs from all specimens of all permutations

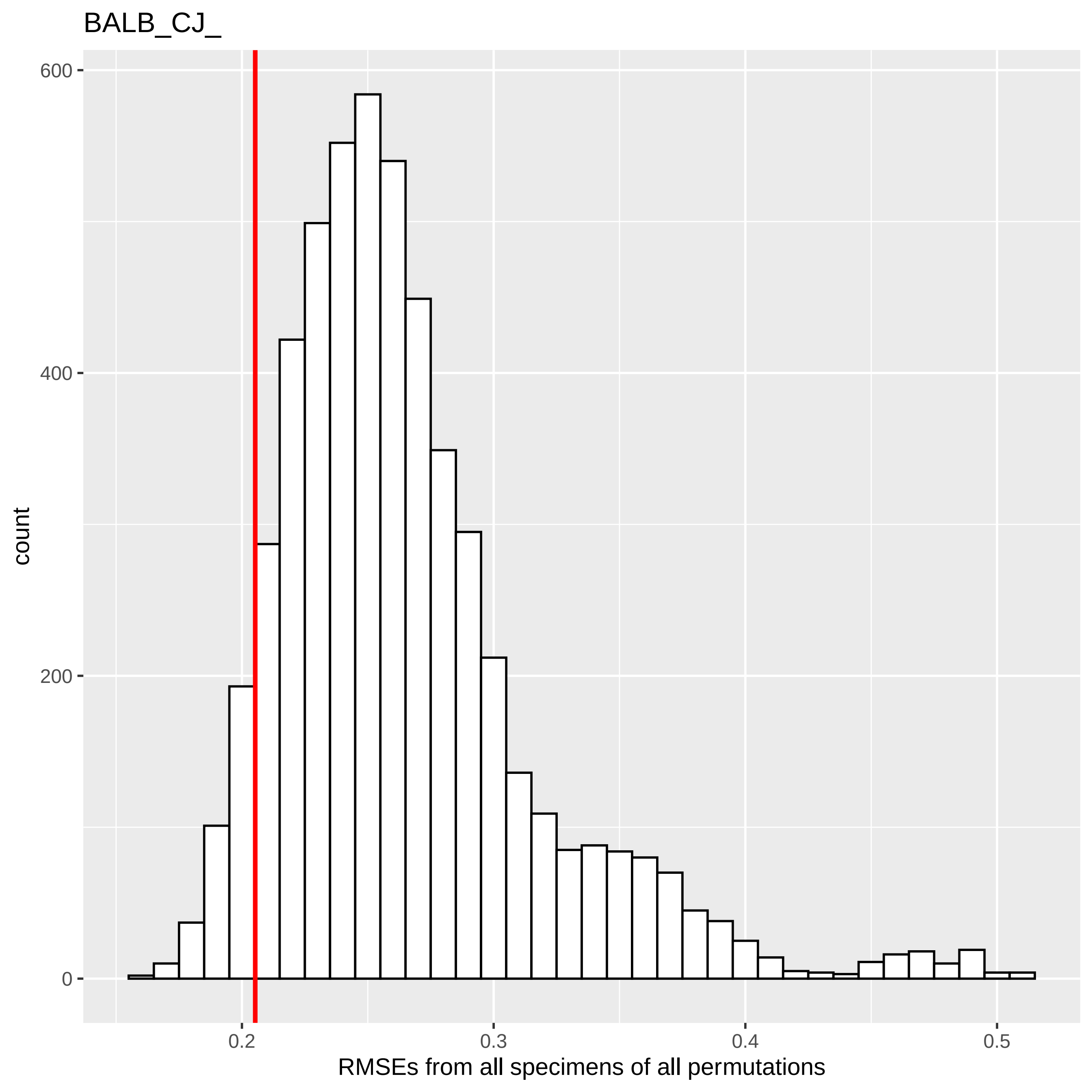
BALB_CJ_
600
400
200
0
count
0.2
0.3
0.4
0.5
RMSEs from all specimens of all permutations

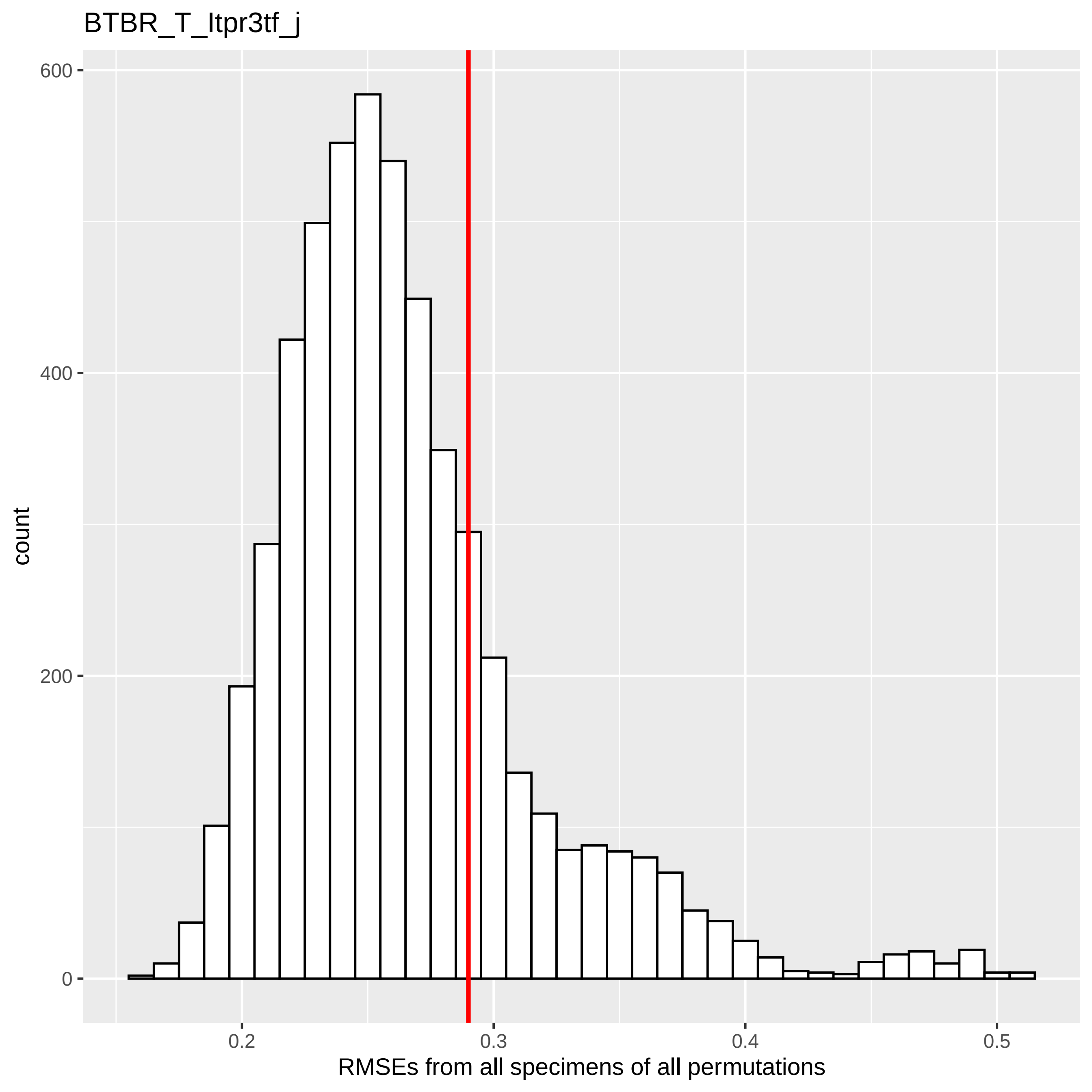

BTBR_T_Itpr3tf_j
count
600
400
200
0
0.2
0.3
0.4
0.5
RMSEs from all specimens of all permutations

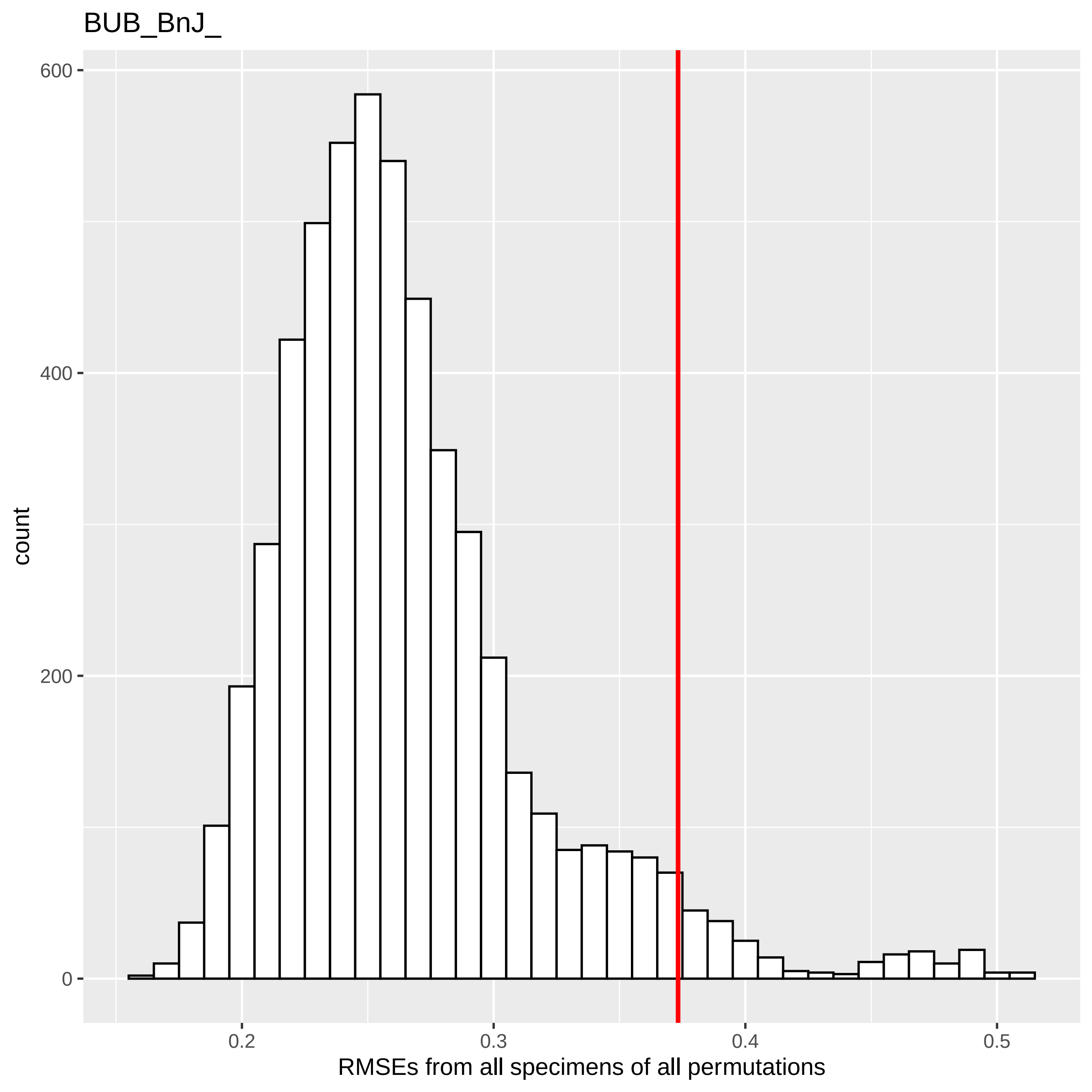

BUB_BnJ_
600
400
200
0
count
0.2
0.3
0.4
0.5
RMSEs from all specimens of all permutations

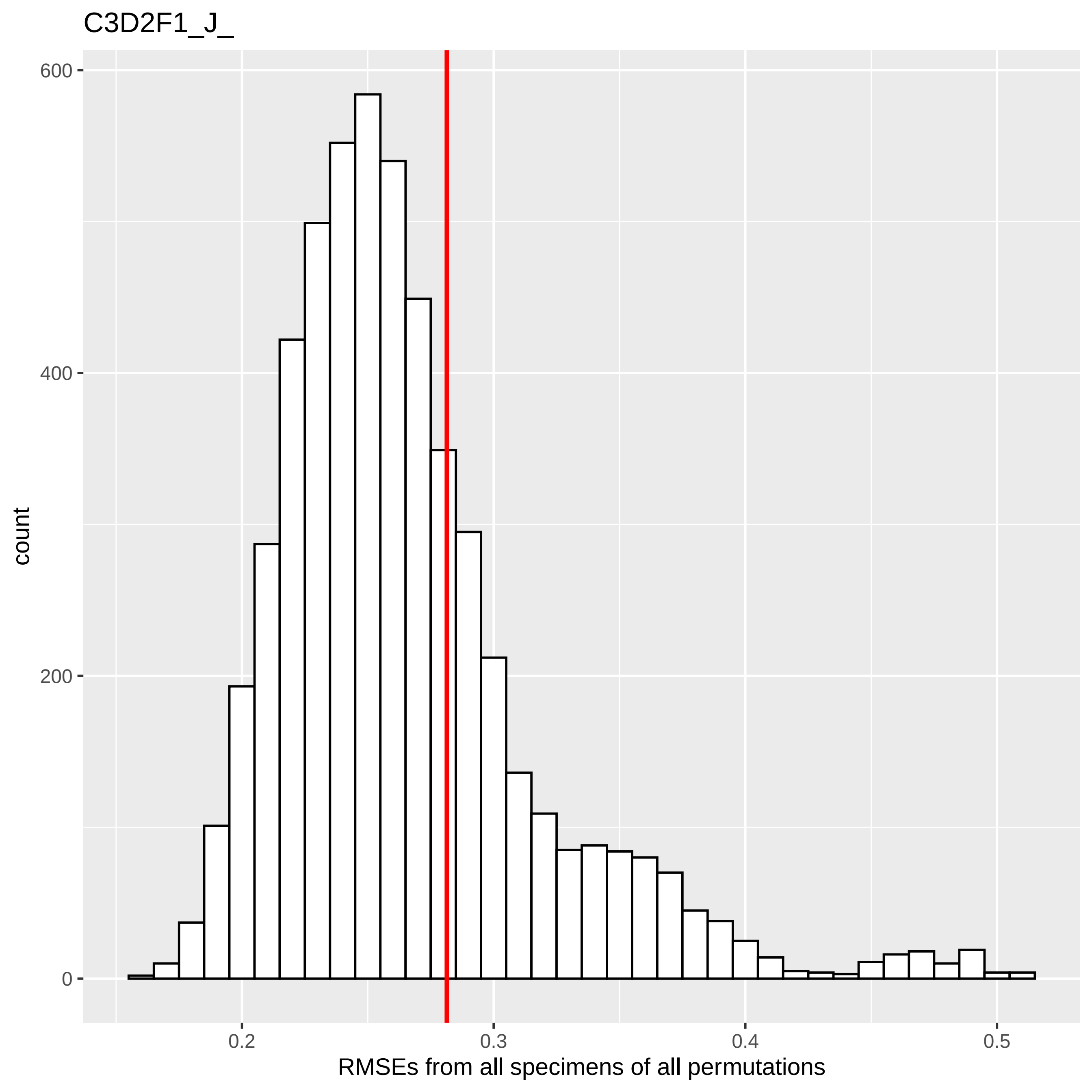
C3D2F1_J_
count
600
400
200
0
0.2
0.3
0.4
0.5
RMSEs from all specimens of all permutations

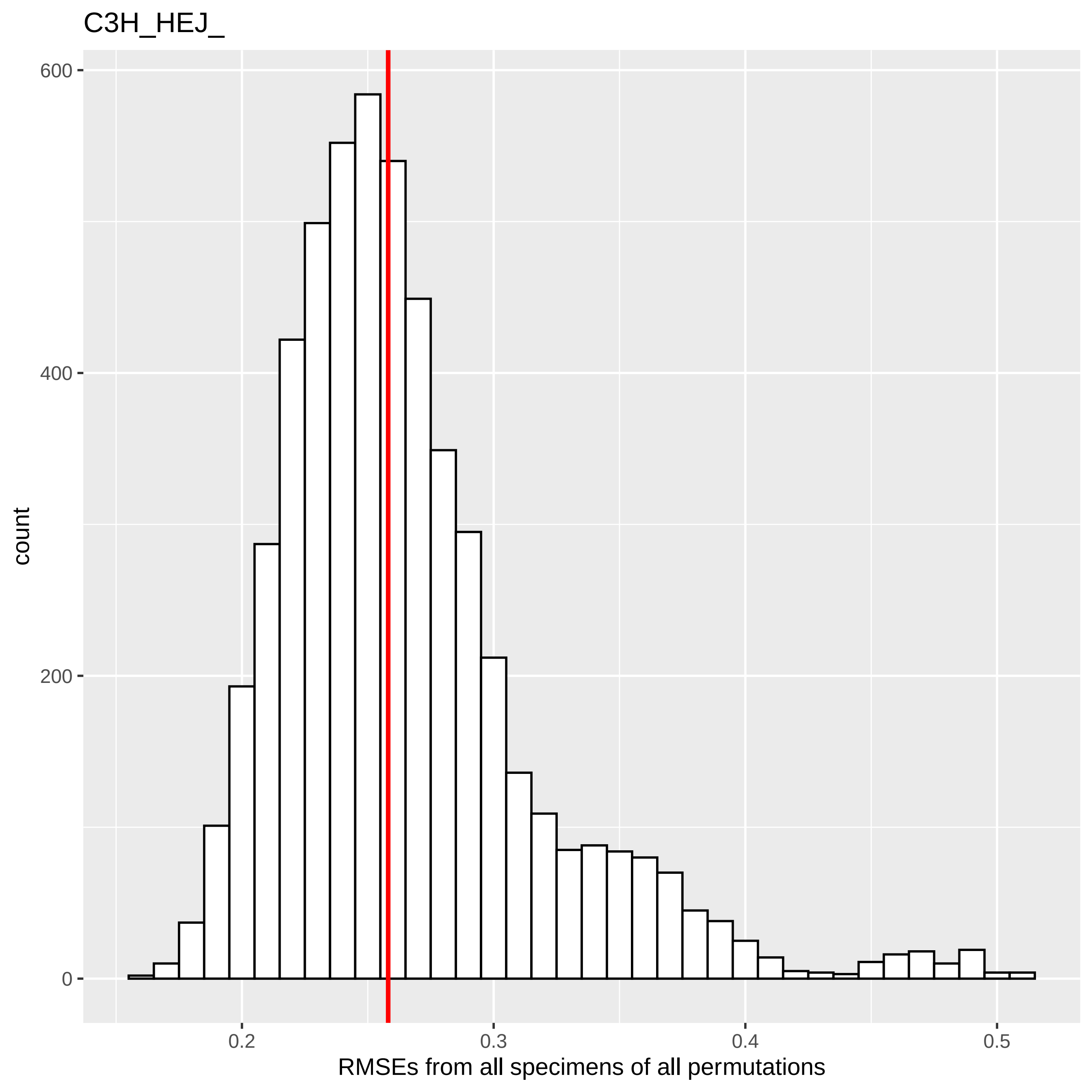

C3H_HEJ_
count
600
400
200
0
0.2
0.3
0.4
0.5
RMSEs from all specimens of all permutations

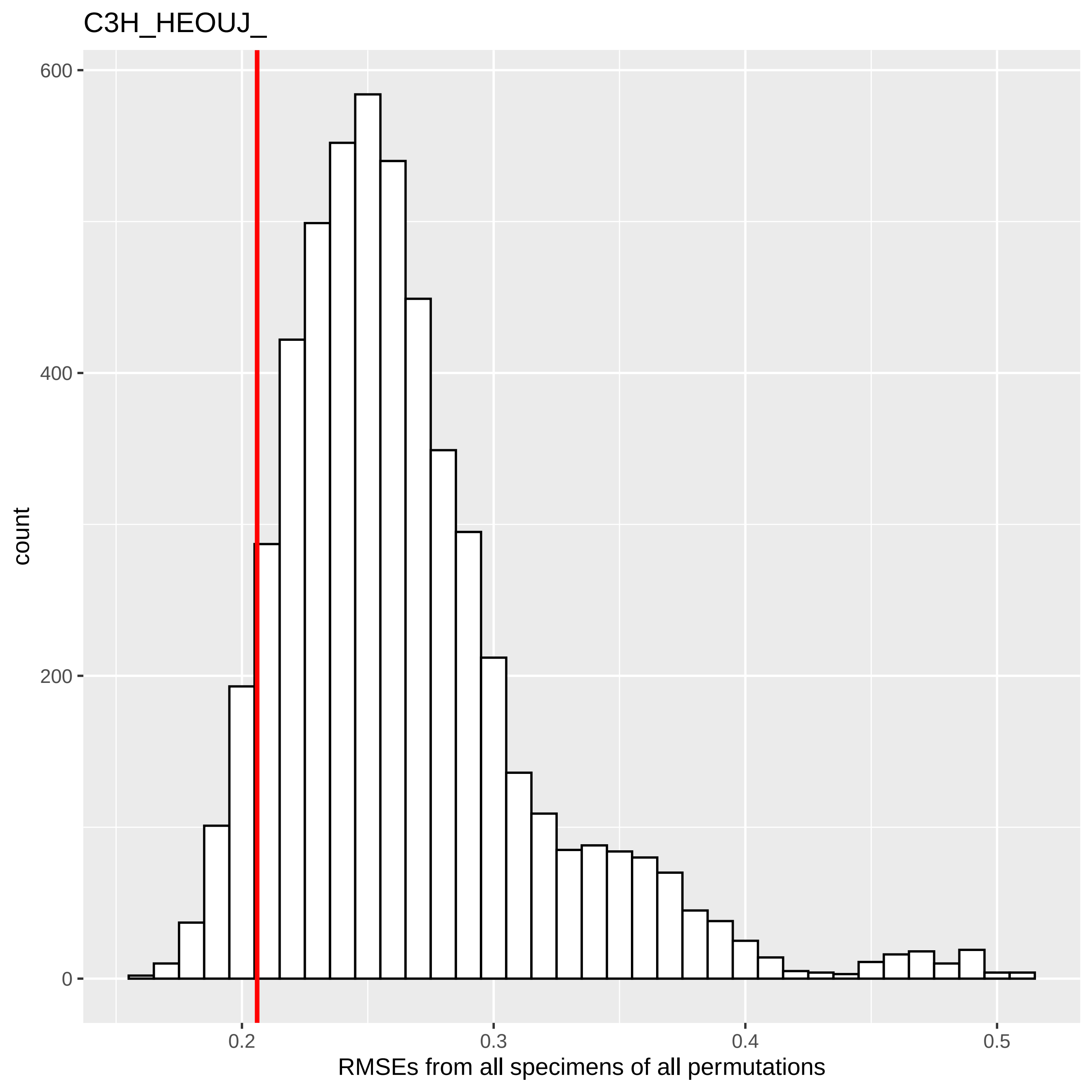
C3H_HEOUJ_
count
600
400
200
0
0.2
0.3
0.4
0.5
RMSEs from all specimens of all permutations

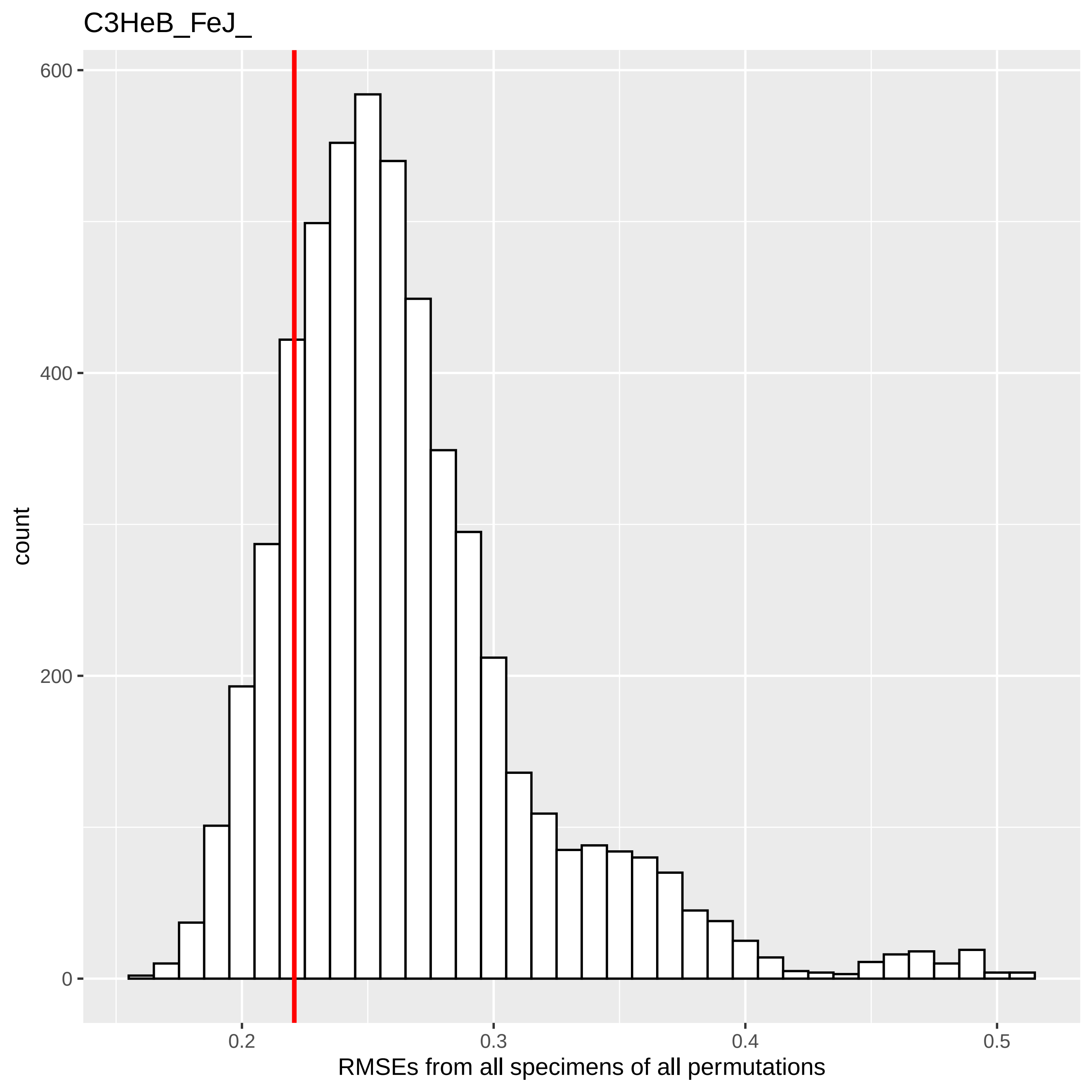
C3HeB_FeJ_
count
600
400
200
0
0.2
0.3
0.4
0.5
RMSEs from all specimens of all permutations

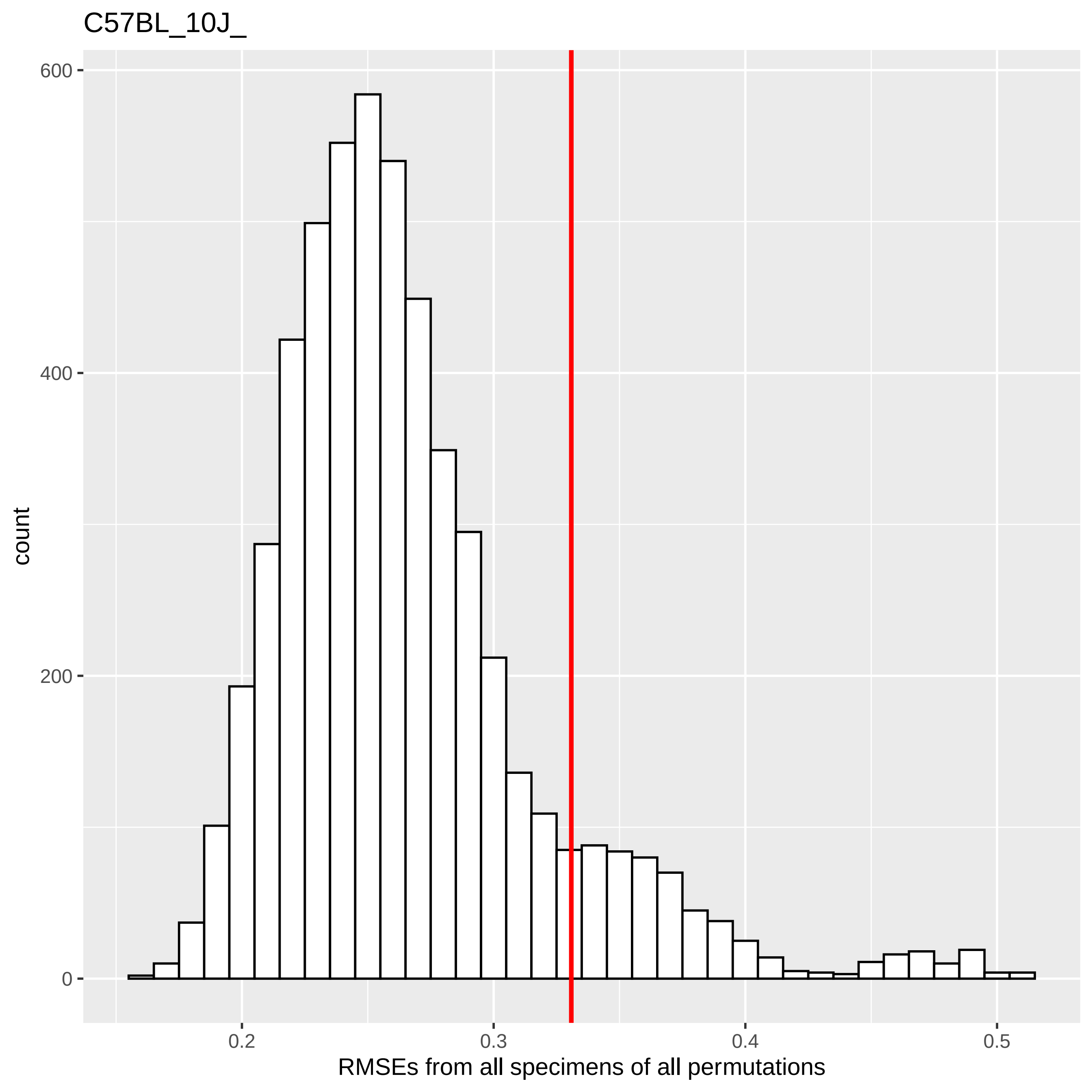
C57BL_10J_
600
400
200
0
count
0.2
0.3
0.4
0.5
RMSEs from all specimens of all permutations

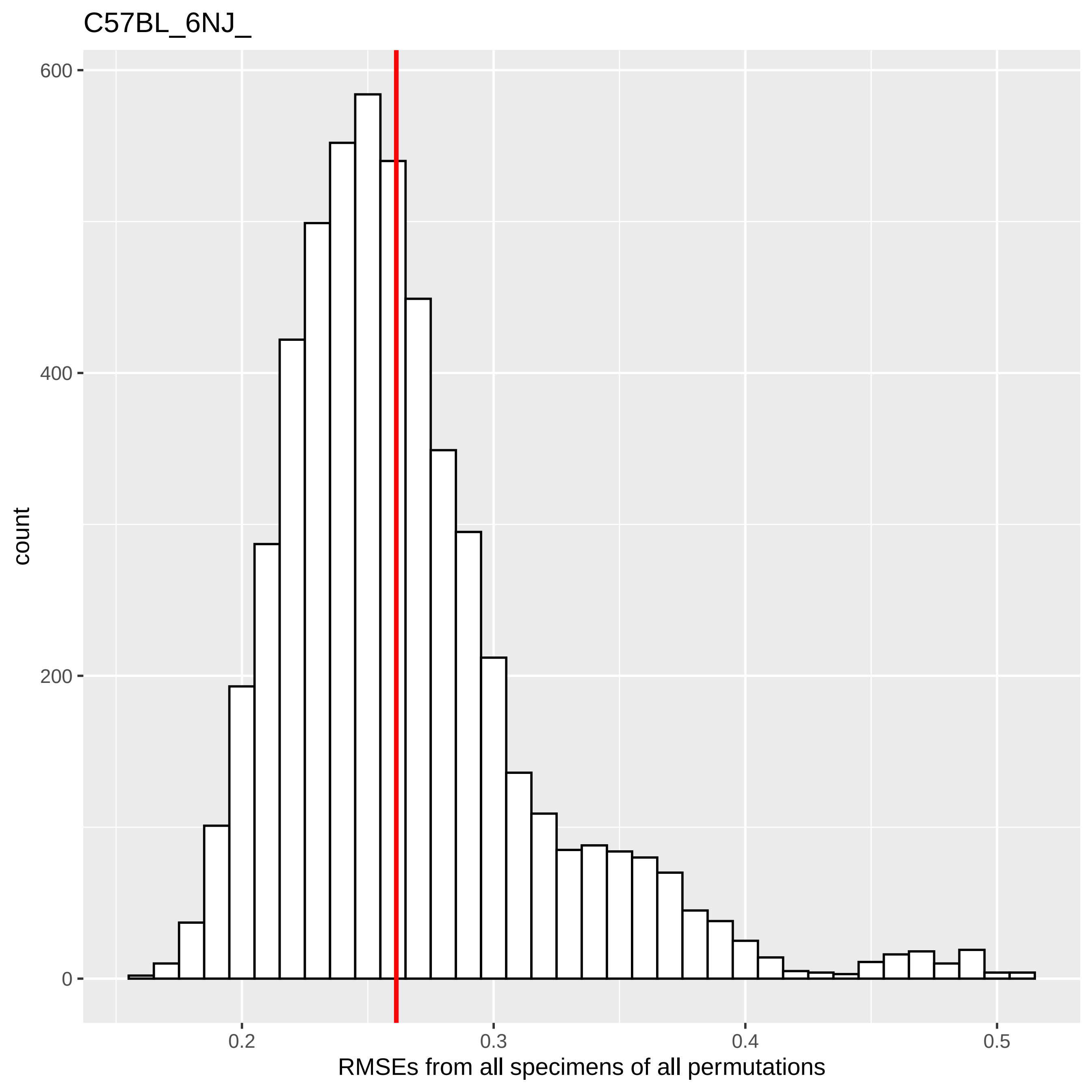
C57BL_6NJ_
600
400
200
0
count
0.2
0.3
0.4
0.5
RMSEs from all specimens of all permutations

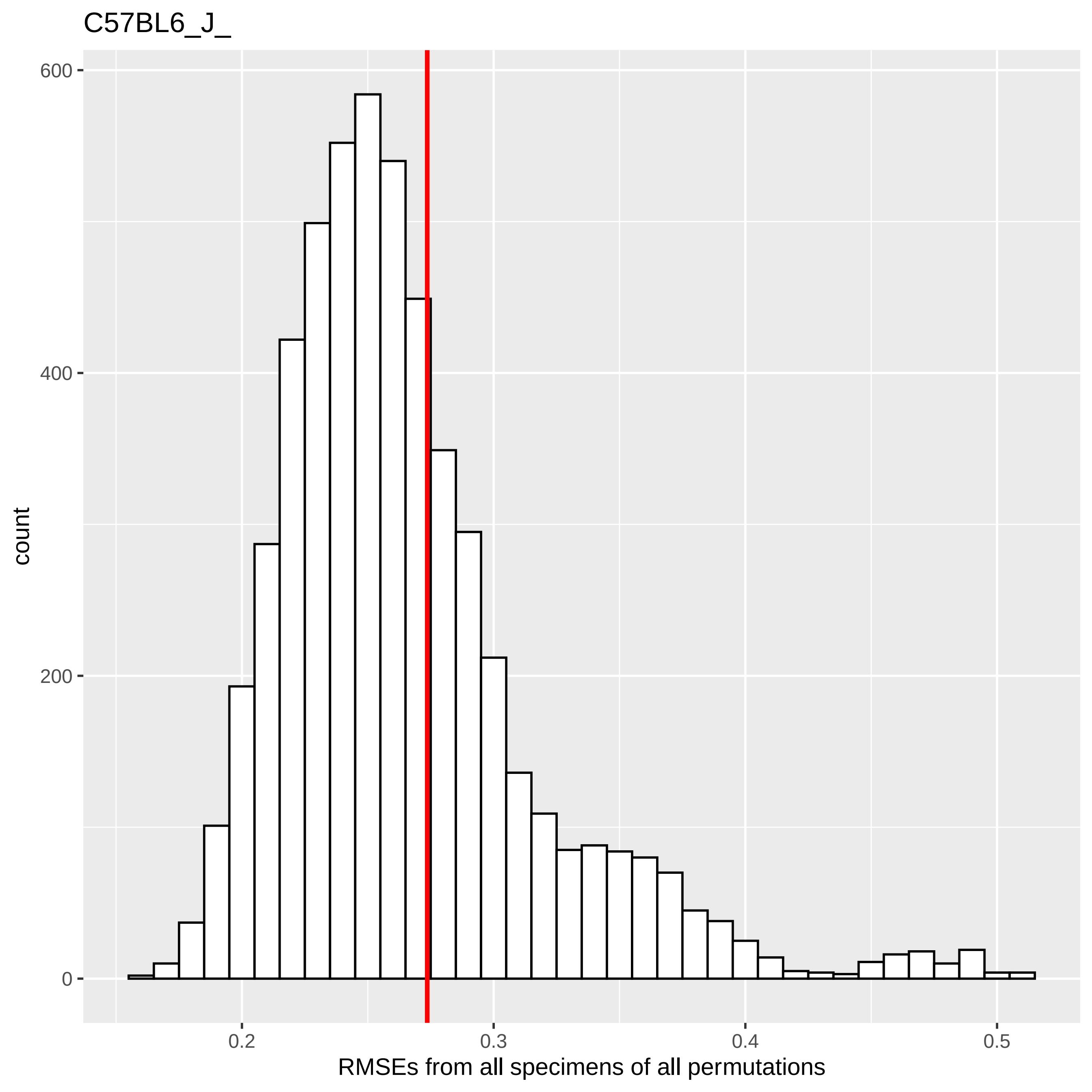
C57BL6_J_
count
600
400
200
0
0.2
0.3
0.4
0.5
RMSEs from all specimens of all permutations

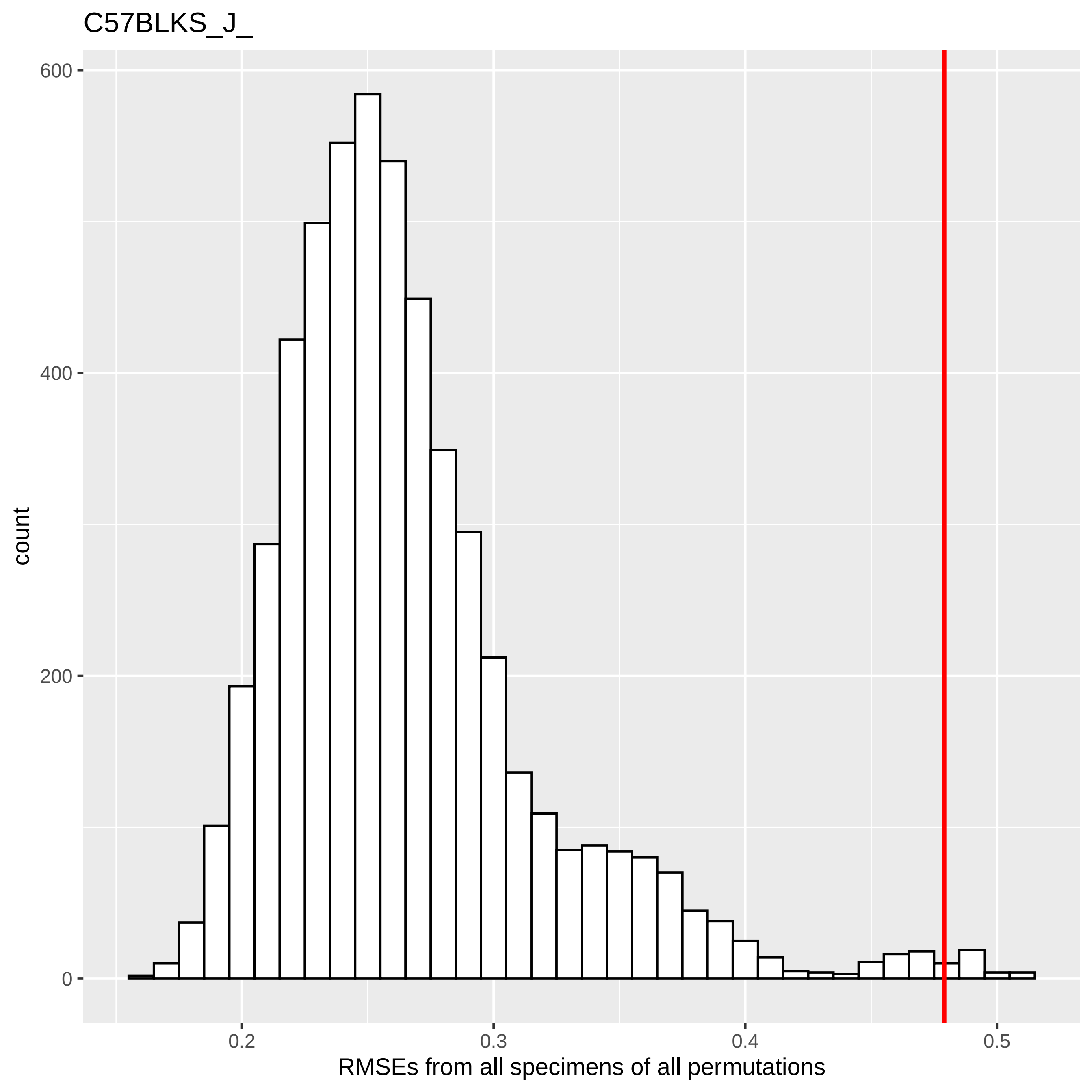
C57BLKS_J_
count
600
400
200
0
0.2
0.3
0.4
0.5
RMSEs from all specimens of all permutations

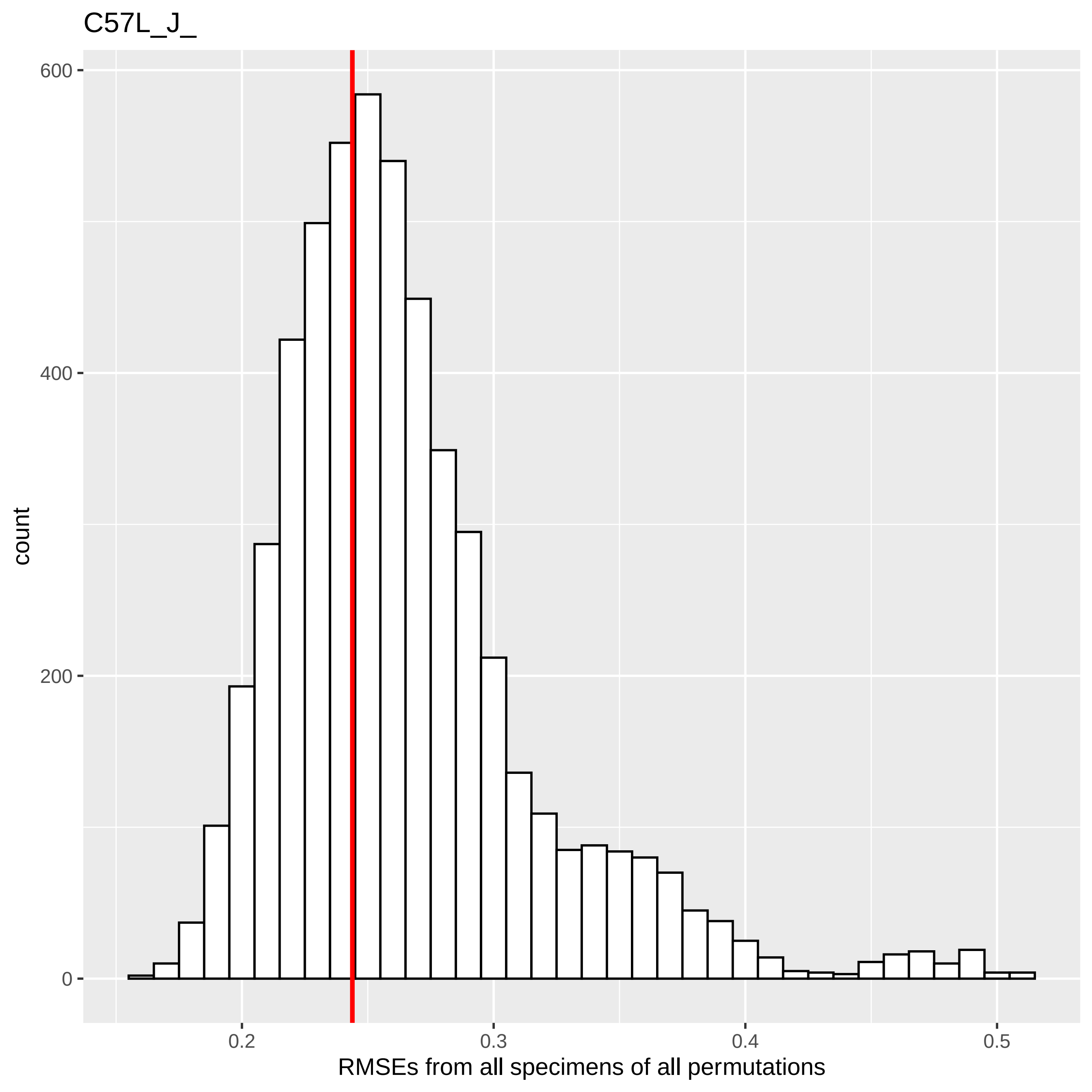
C57L_J_
600
400
200
0
count
0.2
0.3
0.4
0.5
RMSEs from all specimens of all permutations

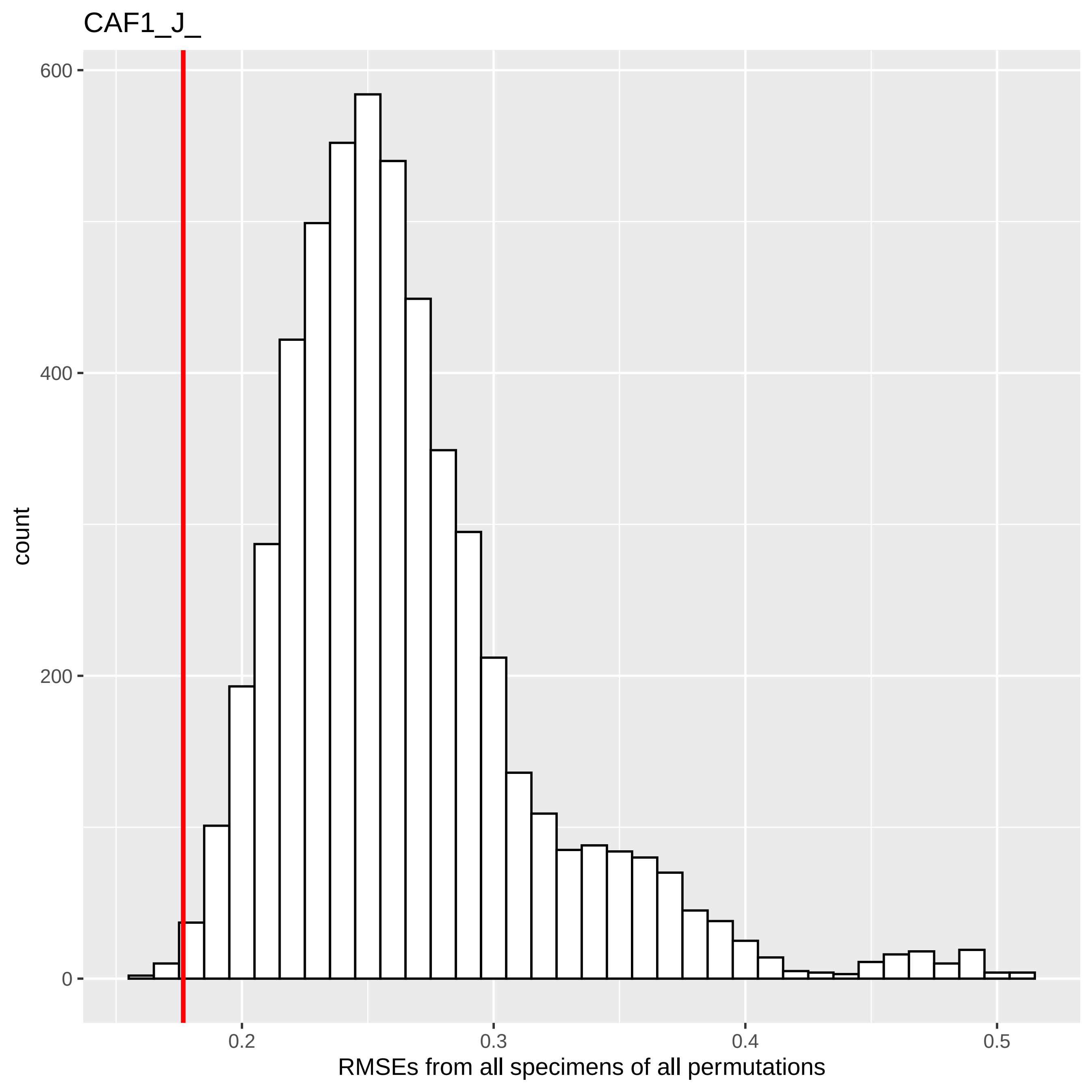
CAF1_J_
count
600
400
200
0
0.2
0.3
0.4
0.5
RMSEs from all specimens of all permutations

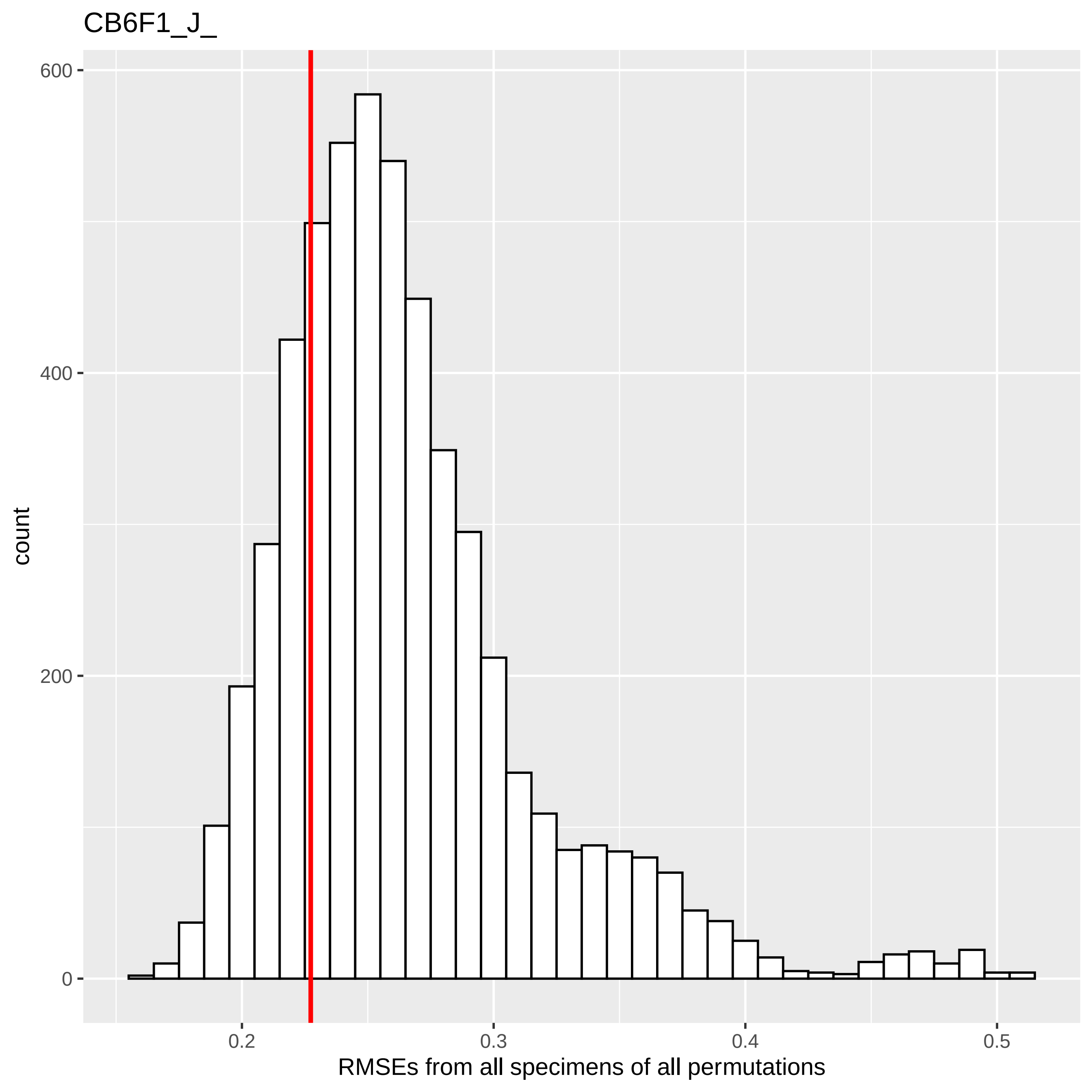
CB6F1_J_
600
400
200
0
count
0.2
0.3
0.4
0.5
RMSEs from all specimens of all permutations

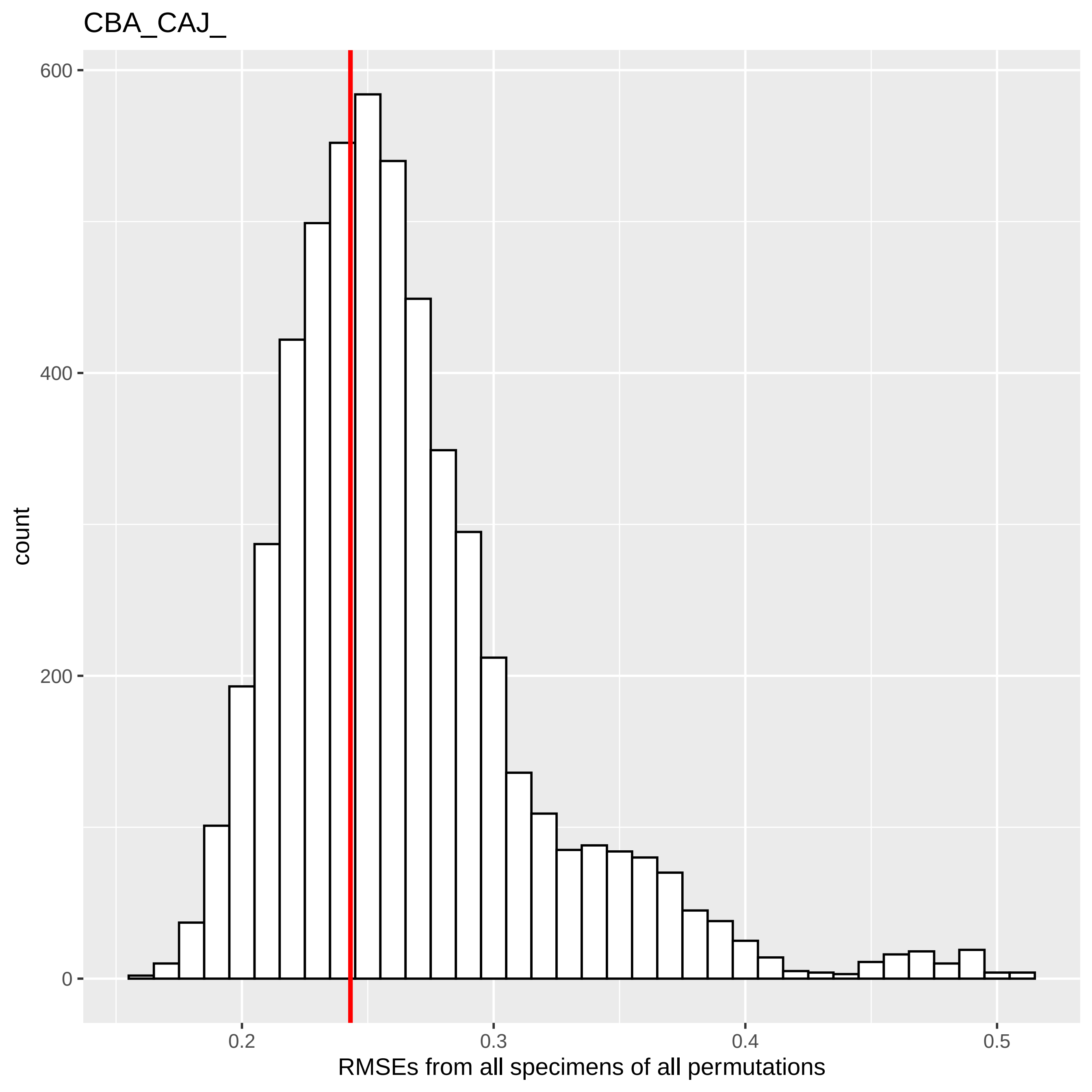

CBA_CAJ_
count
RMSEs from all specimens of all permutations
0
200
400
600
0.2
0.3
0.4
0.5

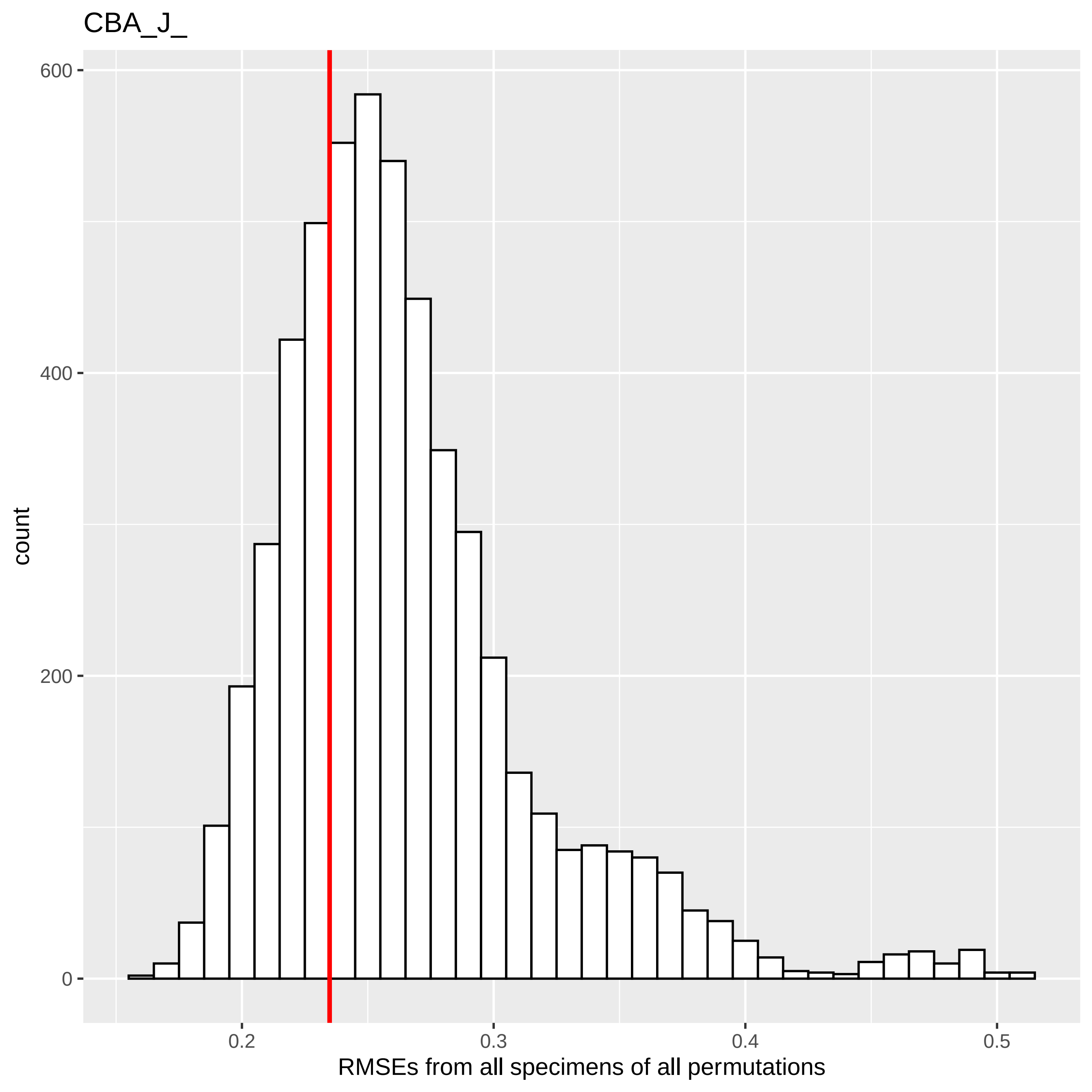

CBA_J_
count
600
400
200
0
0.2
0.3
0.4
0.5
RMSEs from all specimens of all permutations

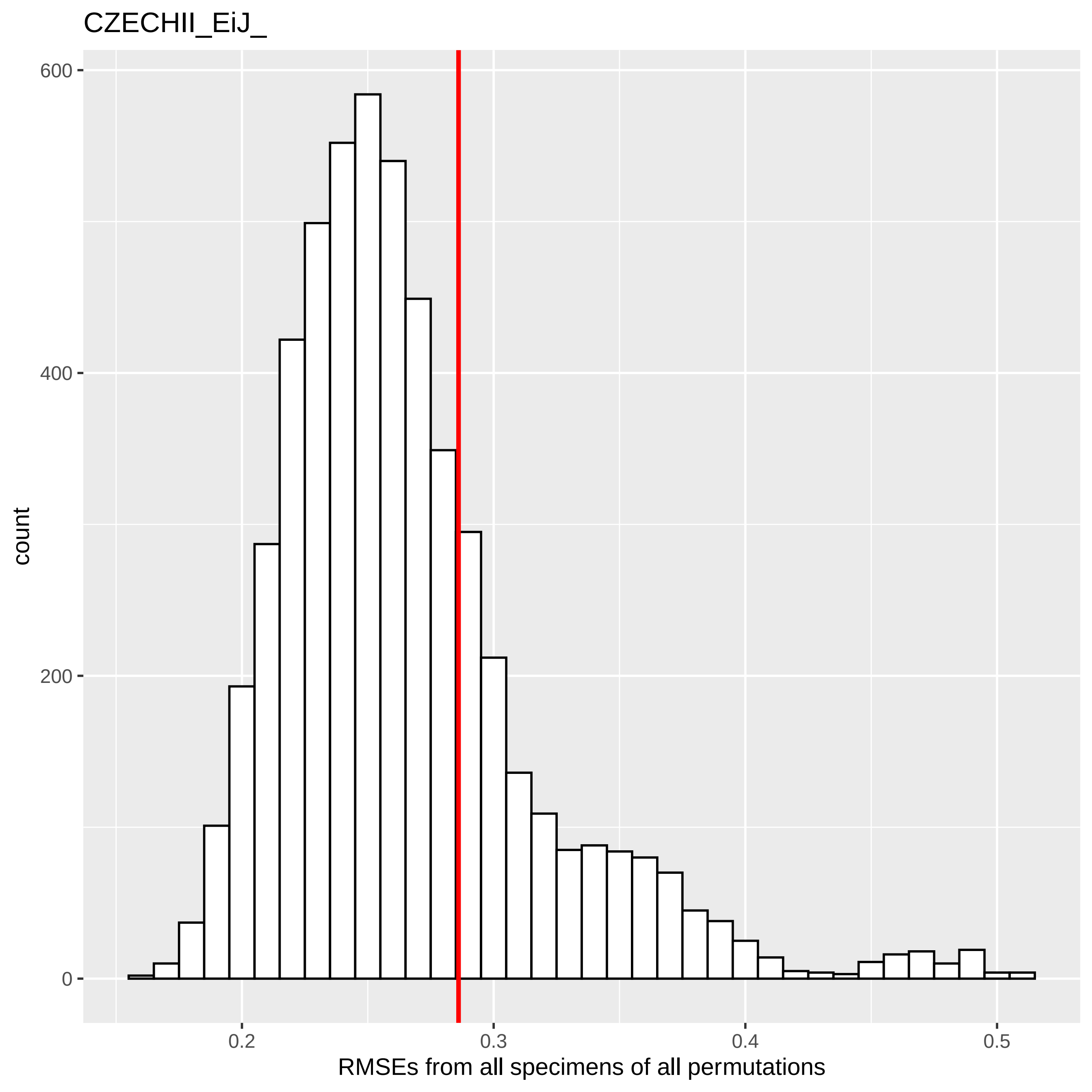
CZECHII_EiJ_
count
600
400
200
0
0.2
0.3
0.4
0.5
RMSEs from all specimens of all permutations

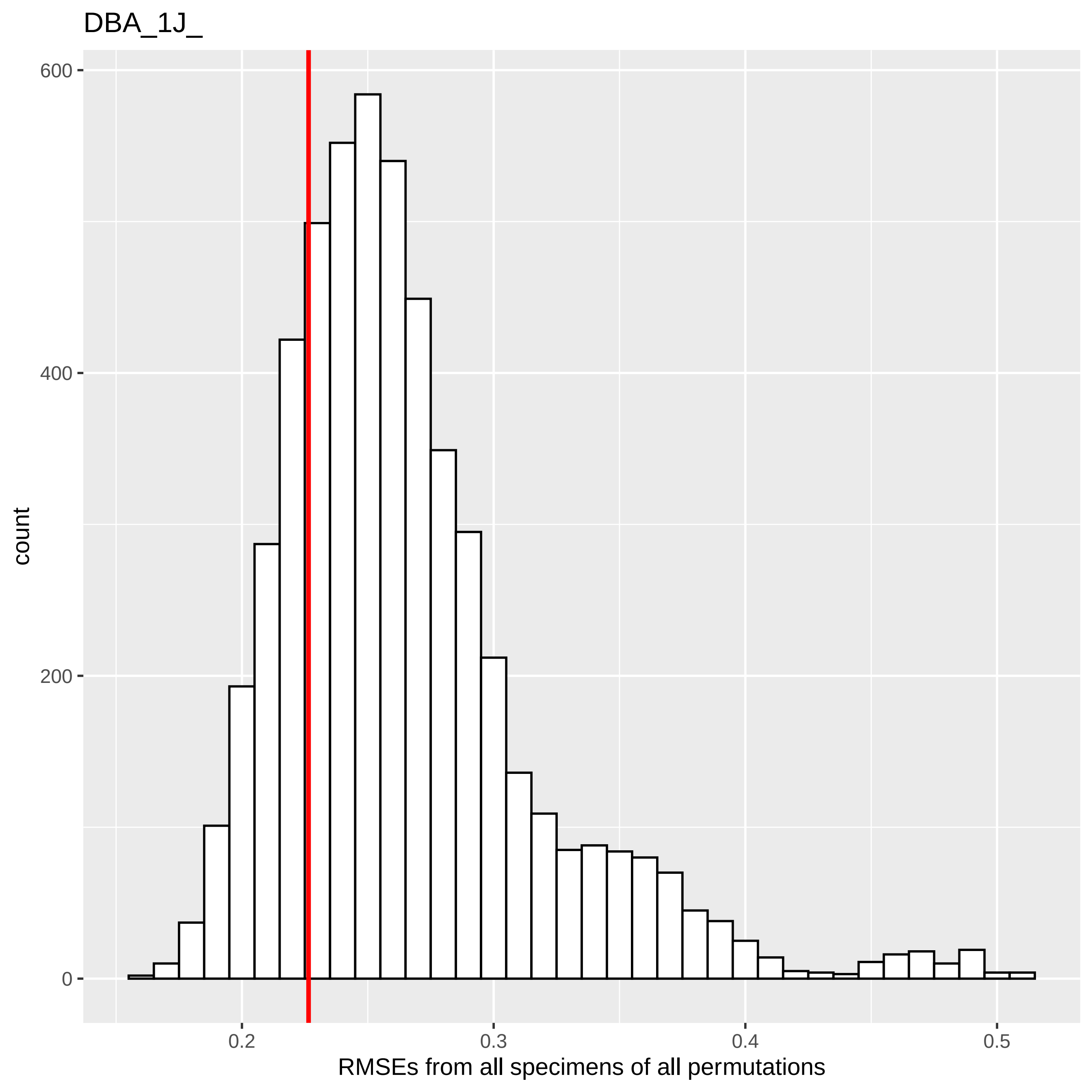

DBA_1J_
count
600
400
200
0
0.2
0.3
0.4
0.5
RMSEs from all specimens of all permutations

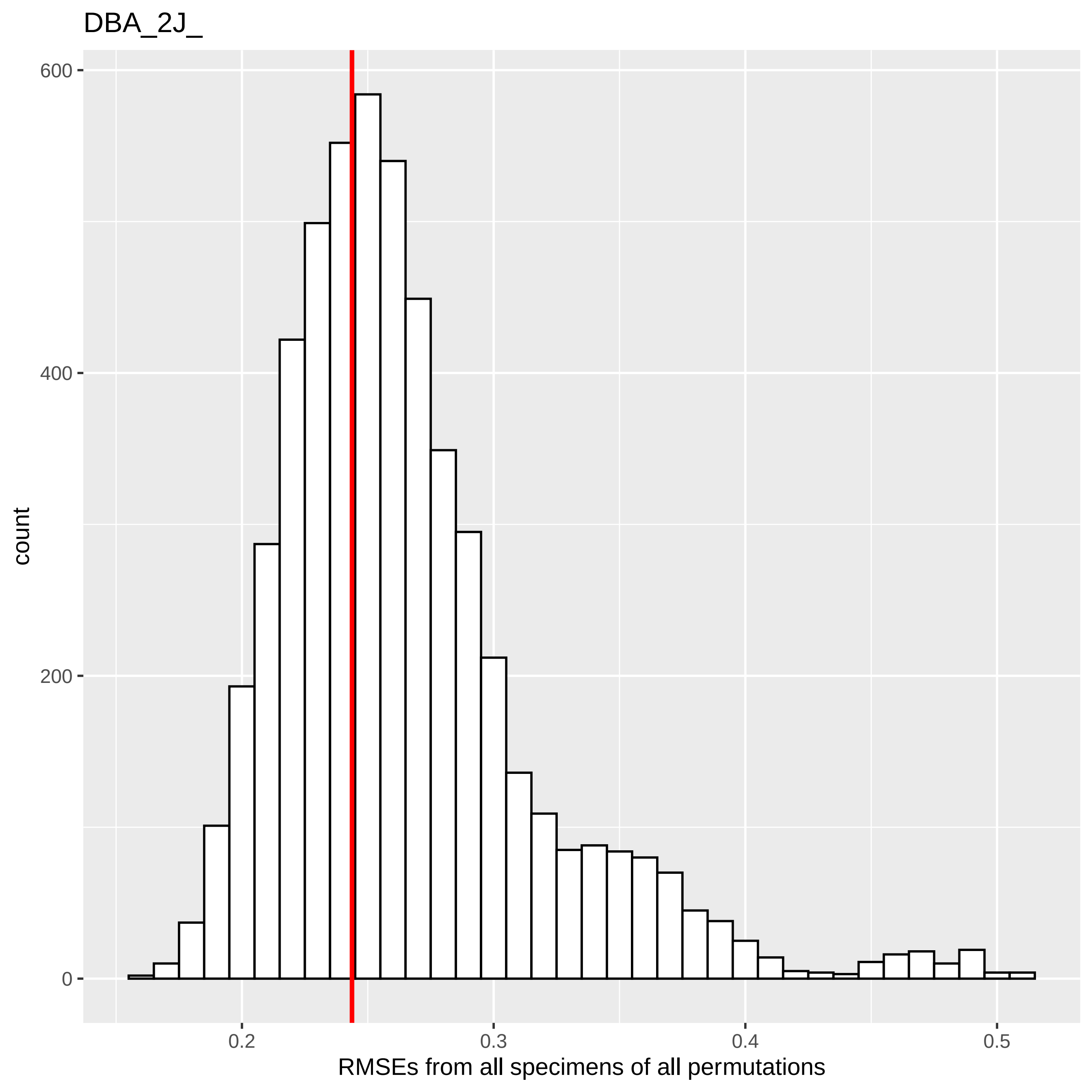
DBA_2J_
600
400
200
0
count
0.2
0.3
0.4
0.5
RMSEs from all specimens of all permutations

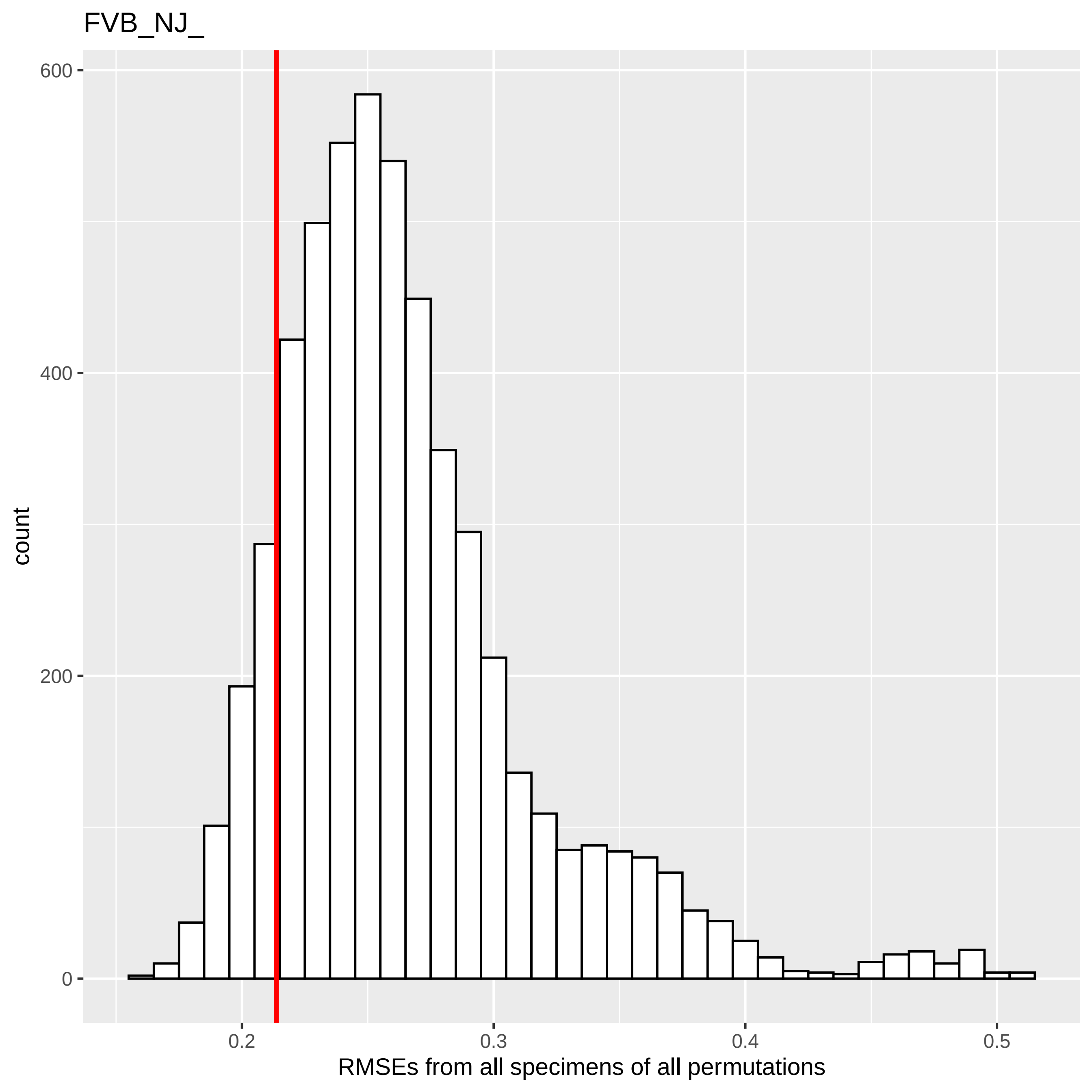
FVB_NJ_
count
600
400
200
0
0.2
0.3
0.4
0.5
RMSEs from all specimens of all permutations

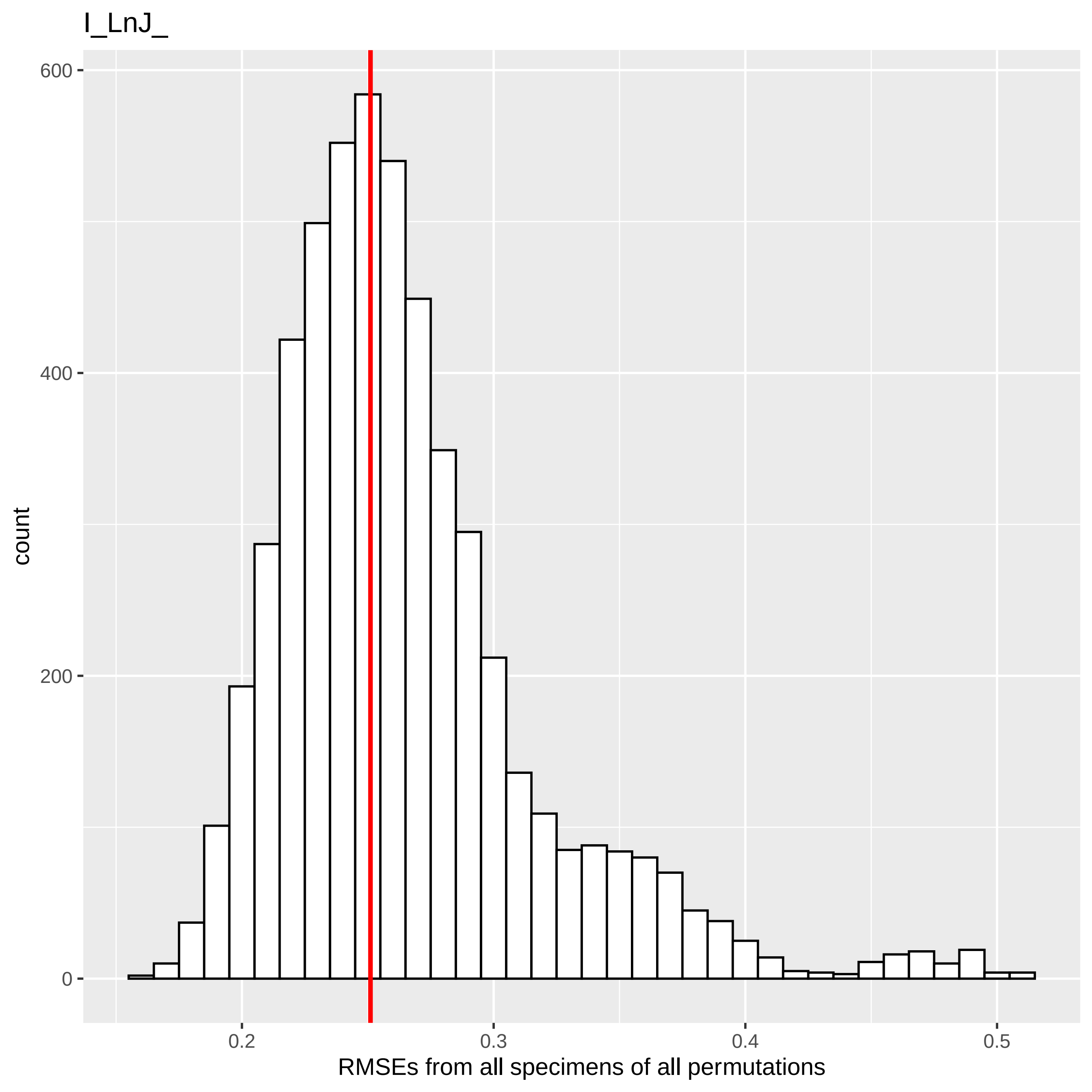
I_LnJ_
600
400
200
0
count
0.2
0.3
0.4
0.5
RMSEs from all specimens of all permutations

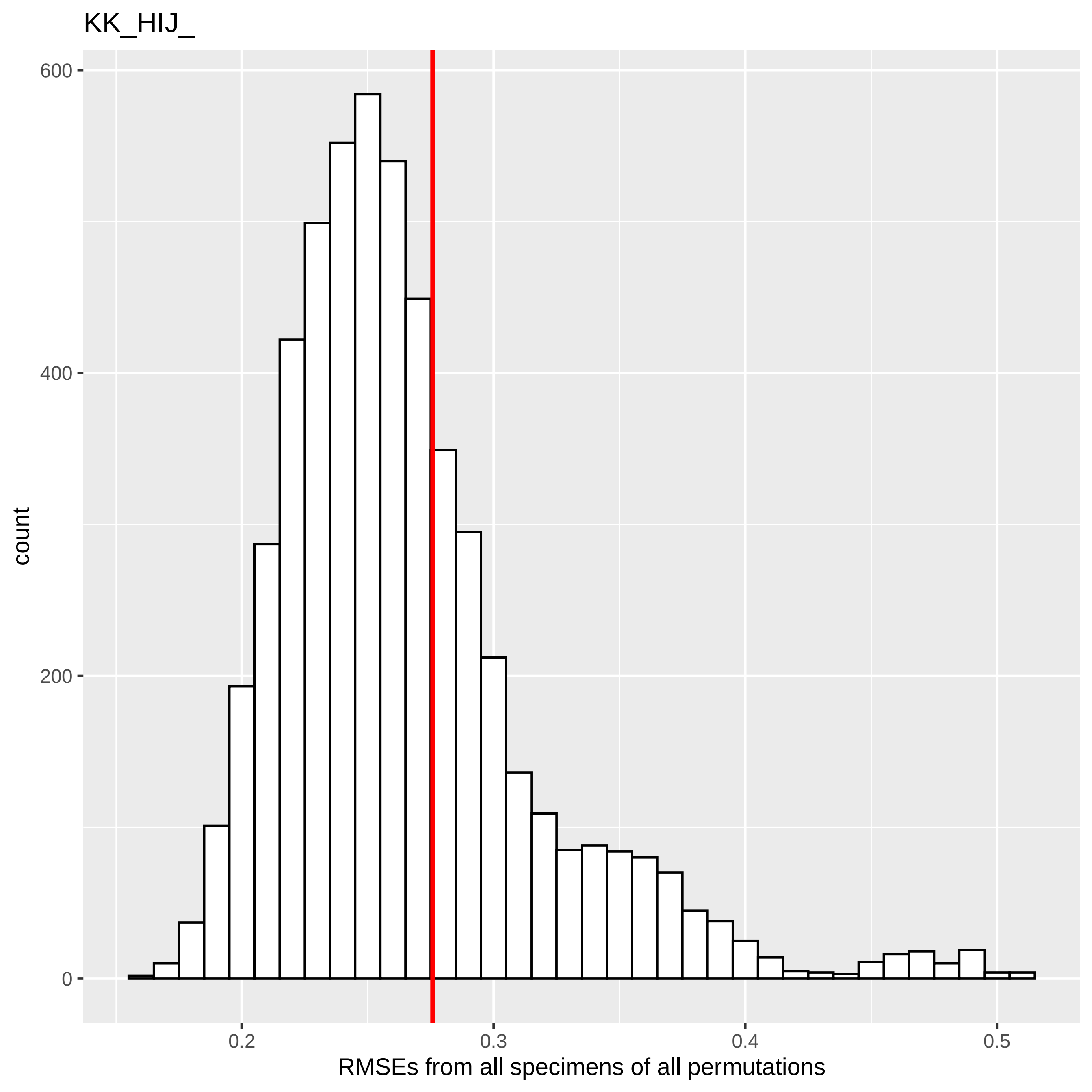

KK_HIJ_
count
600
400
200
0
0.2
0.3
0.4
0.5
RMSEs from all specimens of all permutations

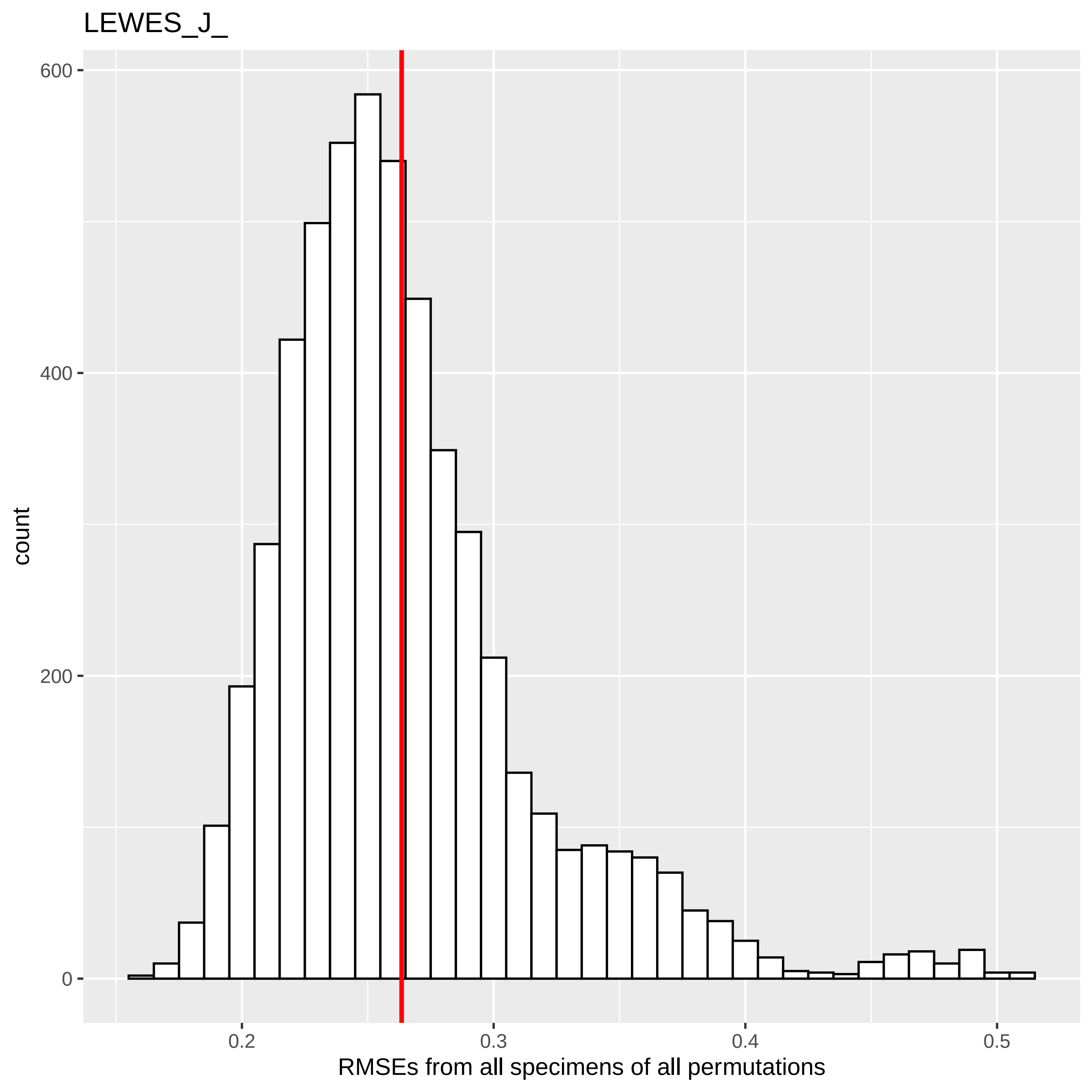

LEWES_J_
600
400
200
0
count
0.2
0.3
0.4
0.5
RMSEs from all specimens of all permutations

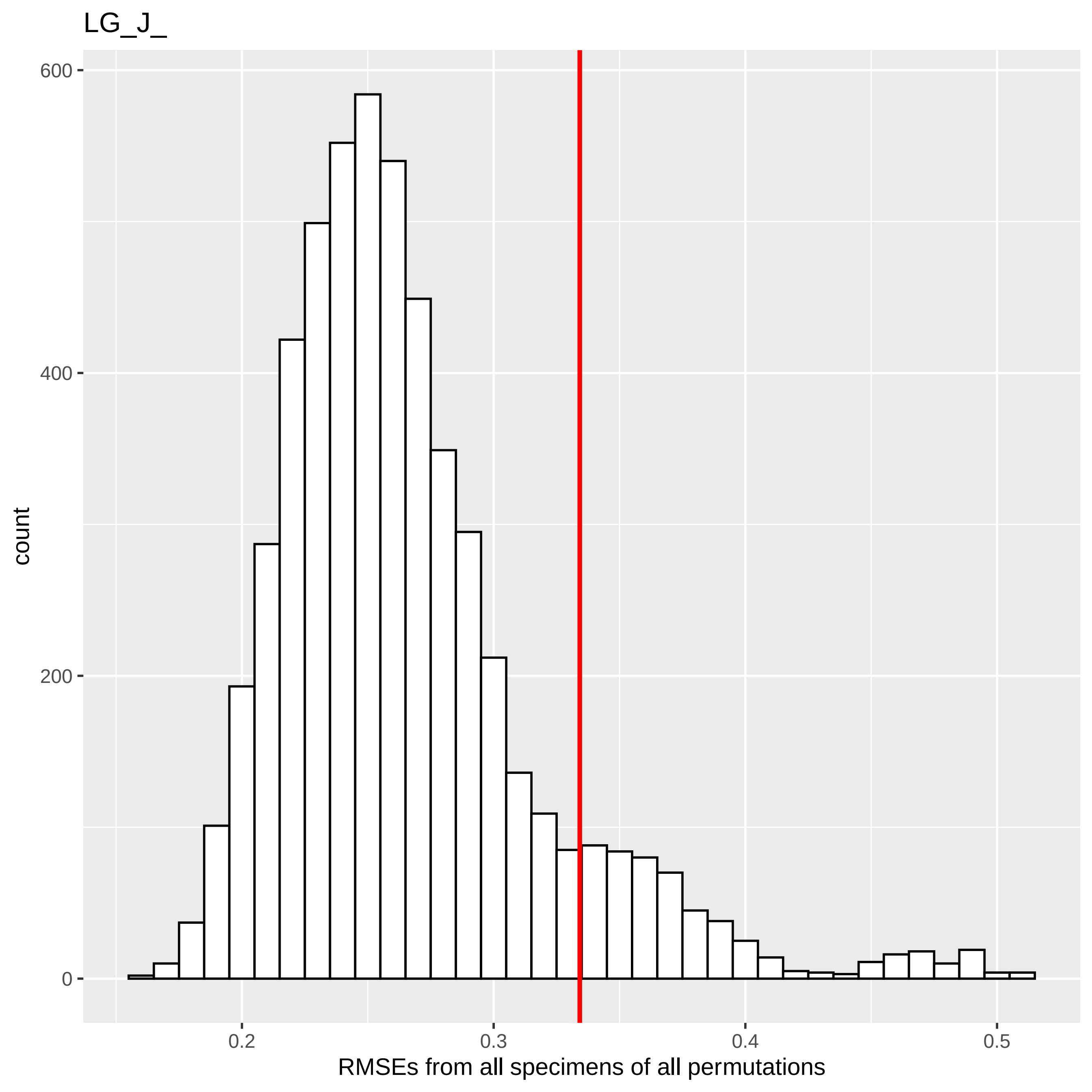
LG_J_
count
RMSEs from all specimens of all permutations
0
200
400
600
0.2
0.3
0.4
0.5

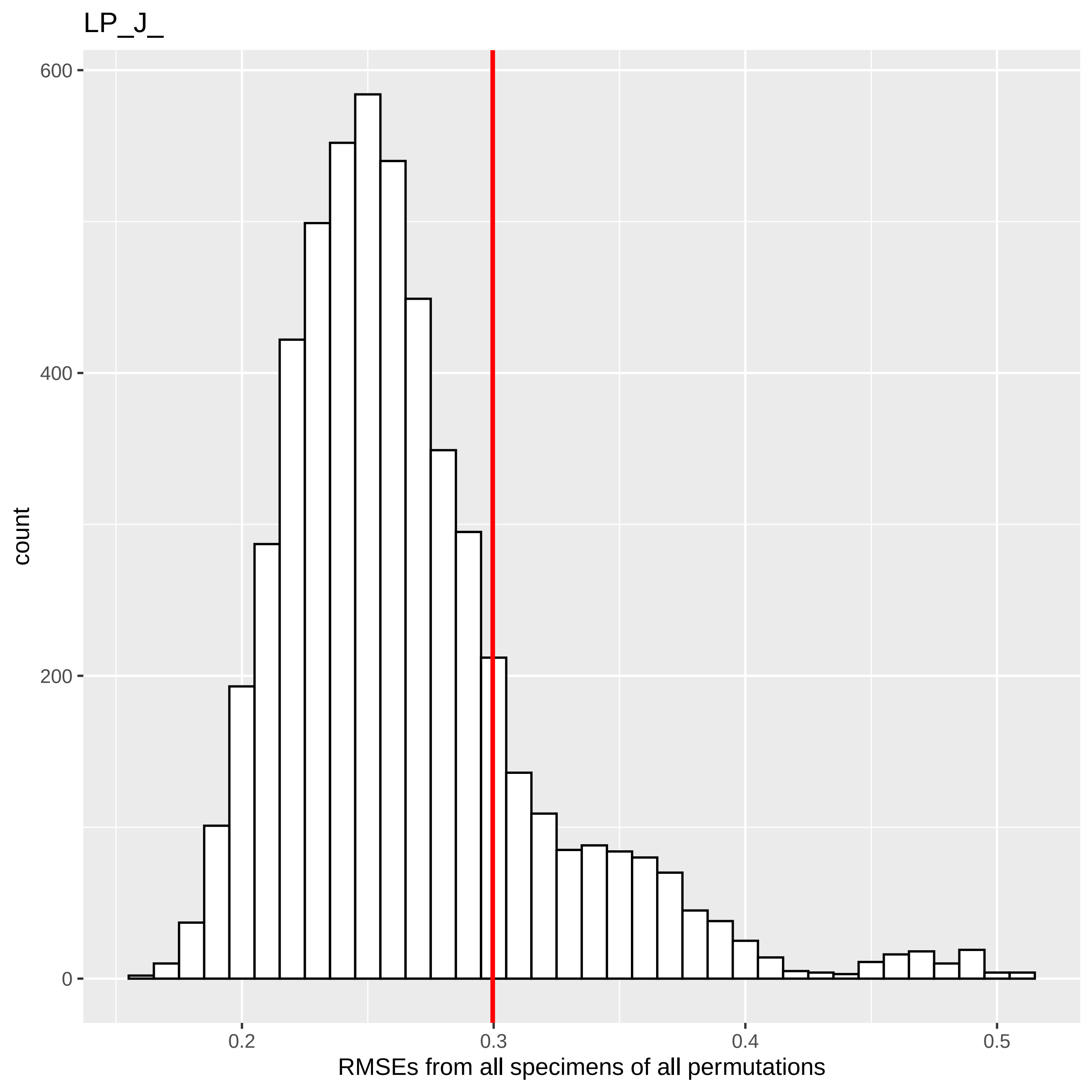

LP_J_
count
600
400
200
0
0.2
0.3
0.4
0.5
RMSEs from all specimens of all permutations

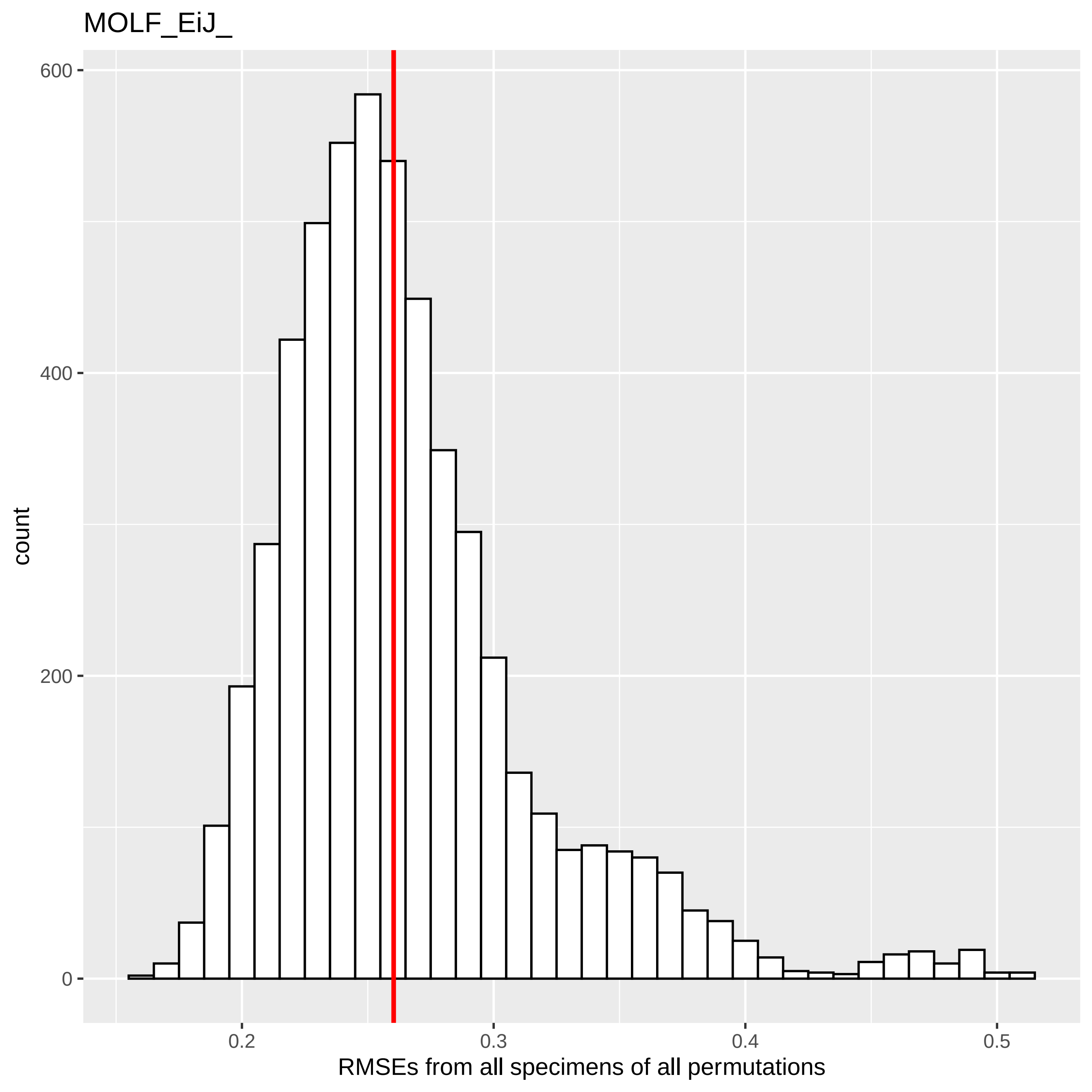
MOLF_EiJ_
count
600
400
200
0
0.2
0.3
0.4
0.5
RMSEs from all specimens of all permutations

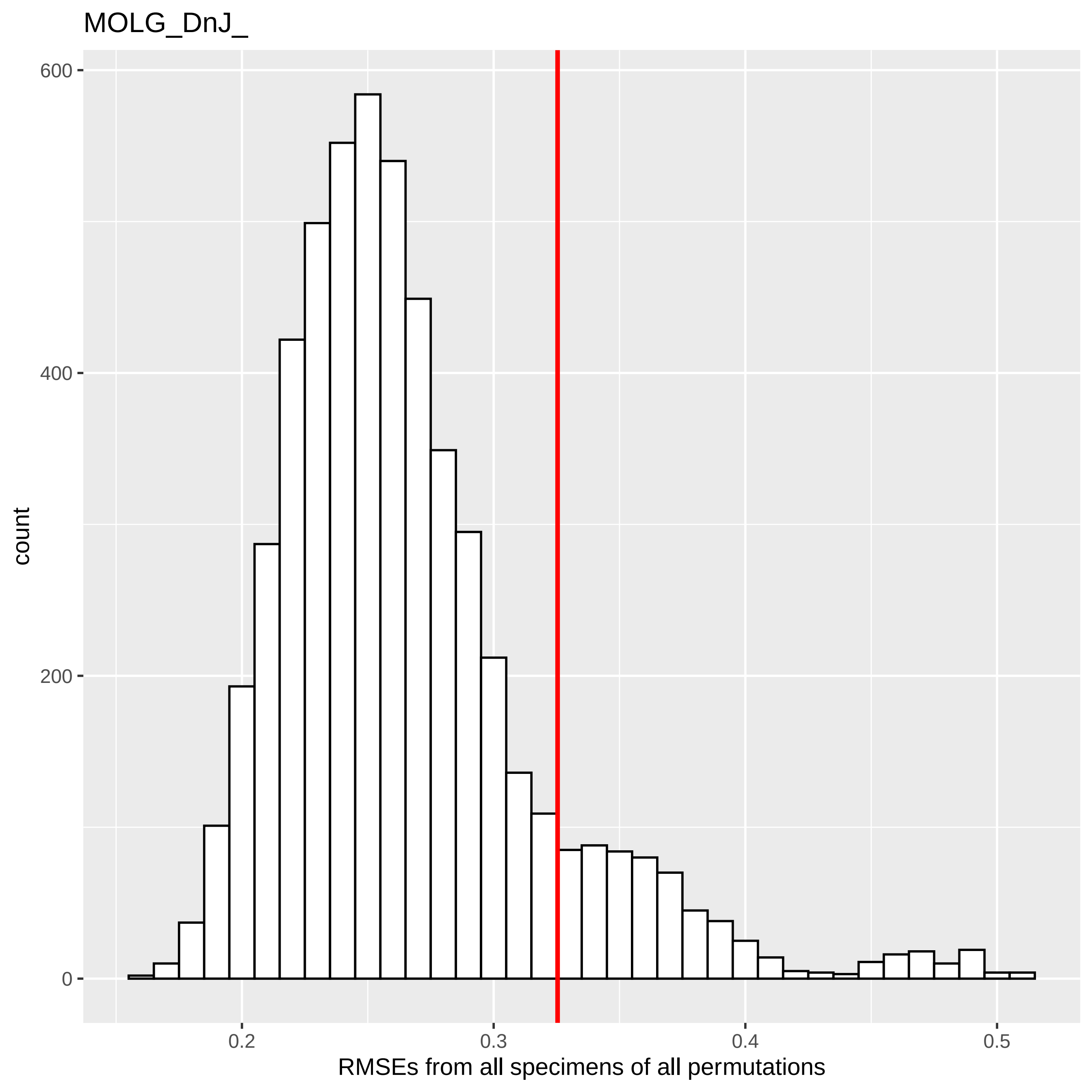
MOLG_DnJ_
600
400
200
0
count
0.2
0.3
0.4
0.5
RMSEs from all specimens of all permutations

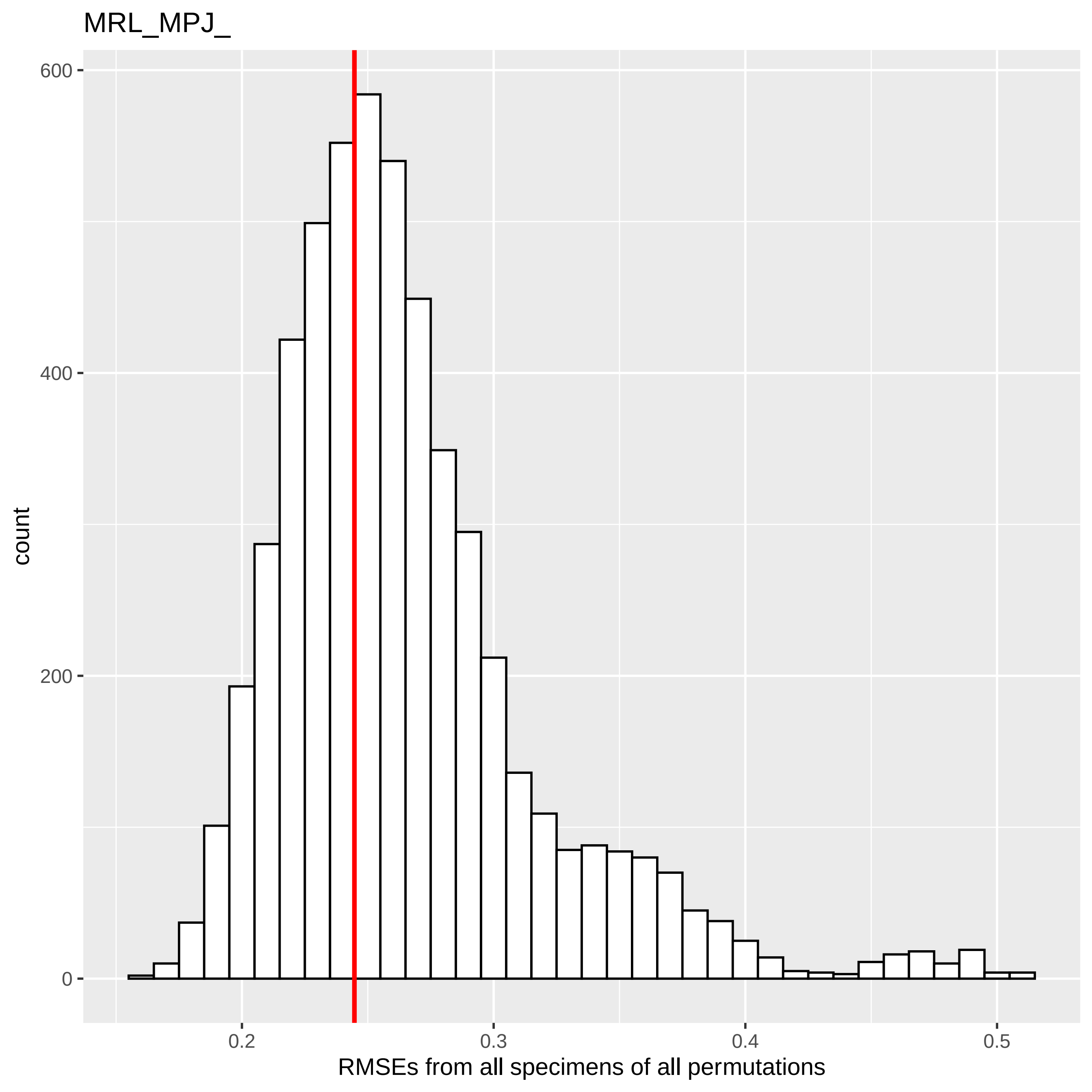
MRL_MPJ_
count
600
400
200
0
0.2
0.3
0.4
0.5
RMSEs from all specimens of all permutations

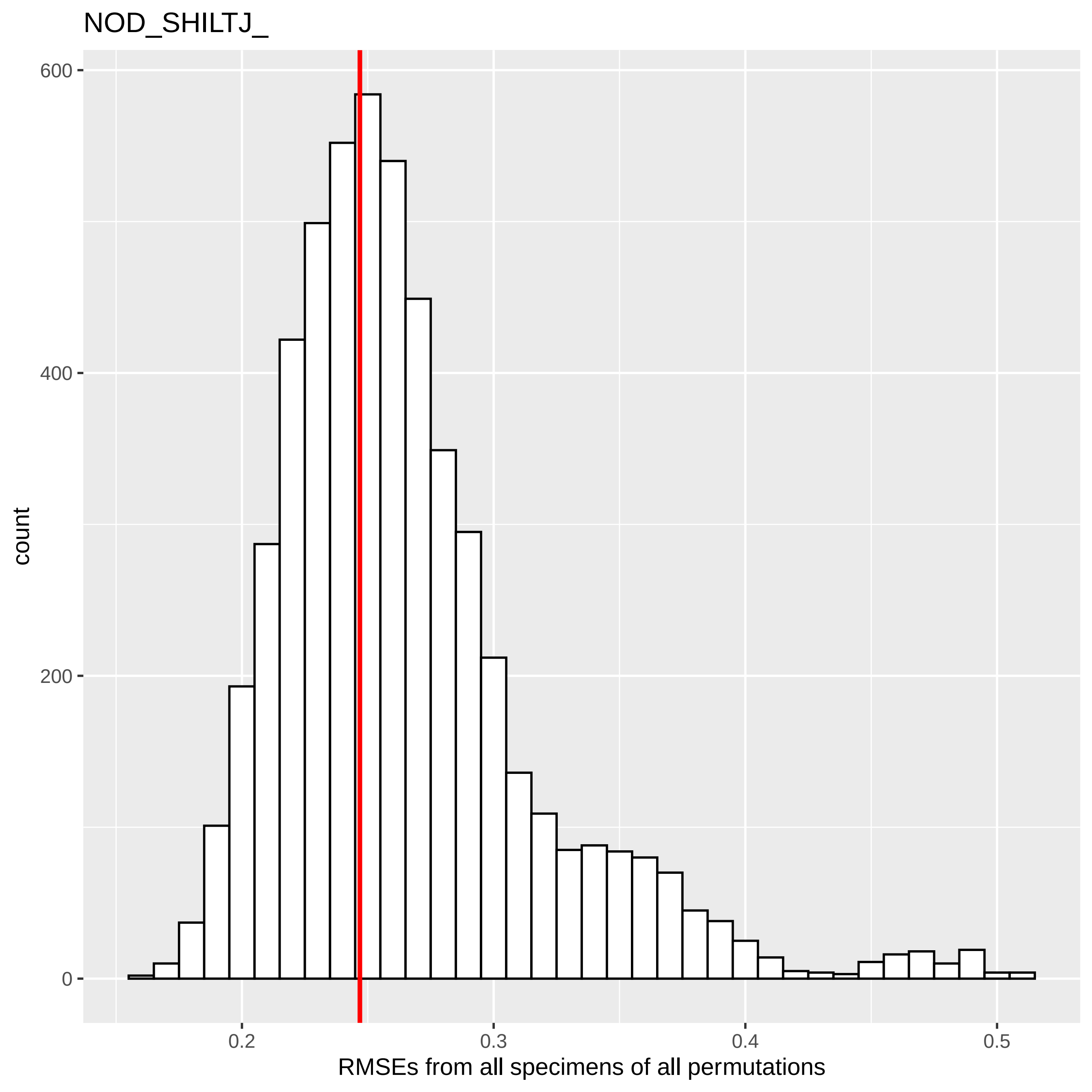

NOD_SHILTJ_
600
400
200
0
count
0.2
0.3
0.4
0.5
RMSEs from all specimens of all permutations

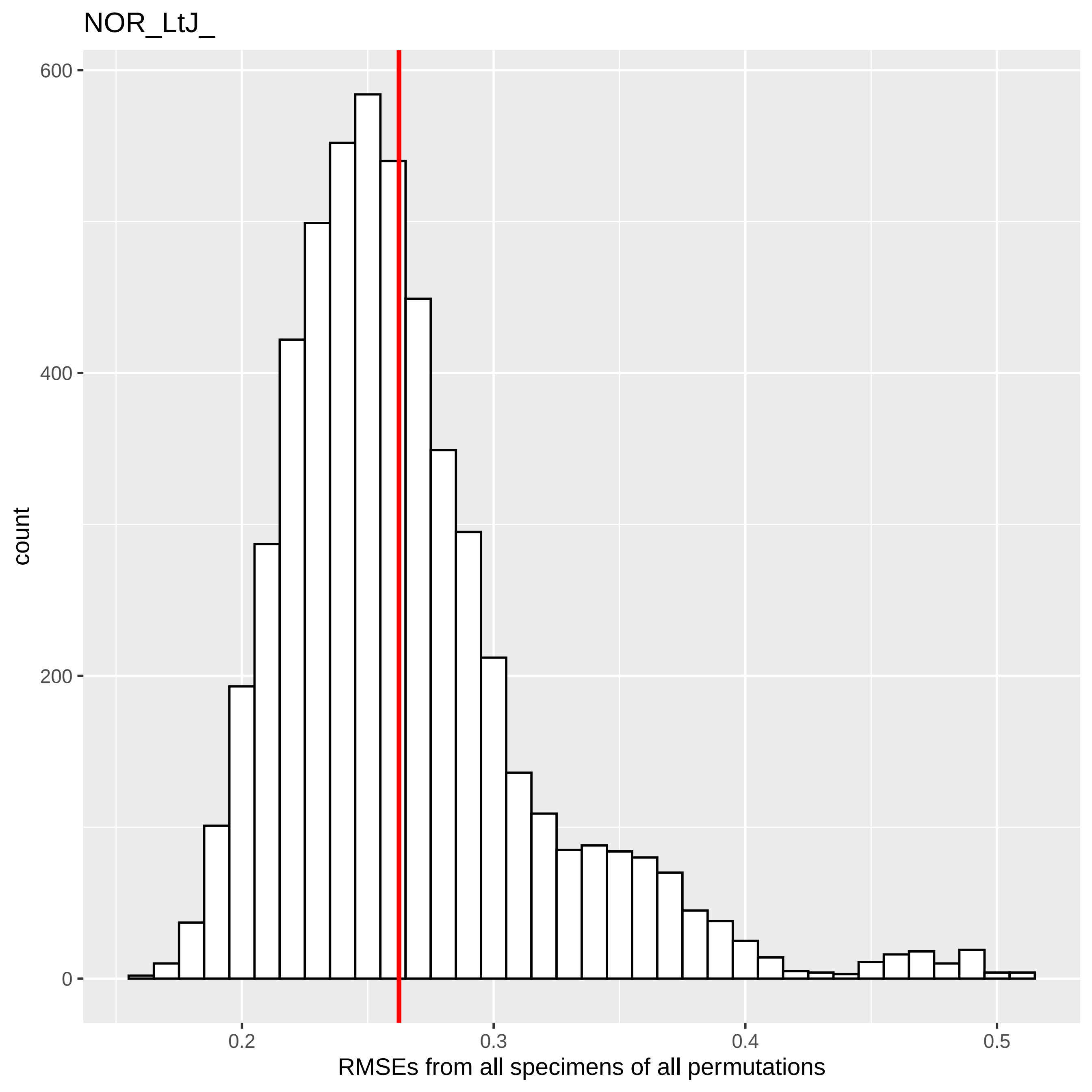

NOR_LtJ_
600
400
200
0
count
0.2
0.3
0.4
0.5
RMSEs from all specimens of all permutations

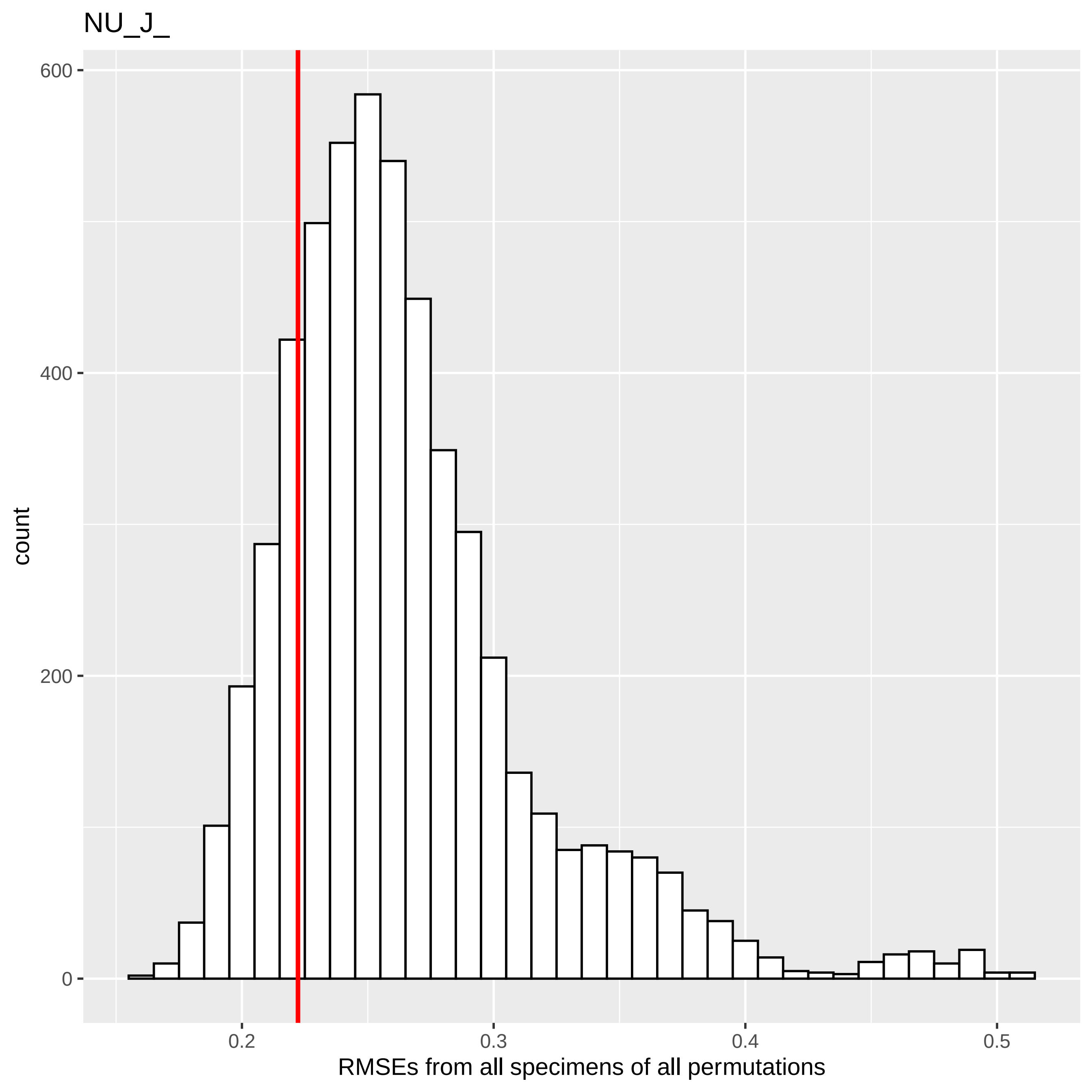
NU_J_
600
400
200
0
count
0.2
0.3
0.4
0.5
RMSEs from all specimens of all permutations

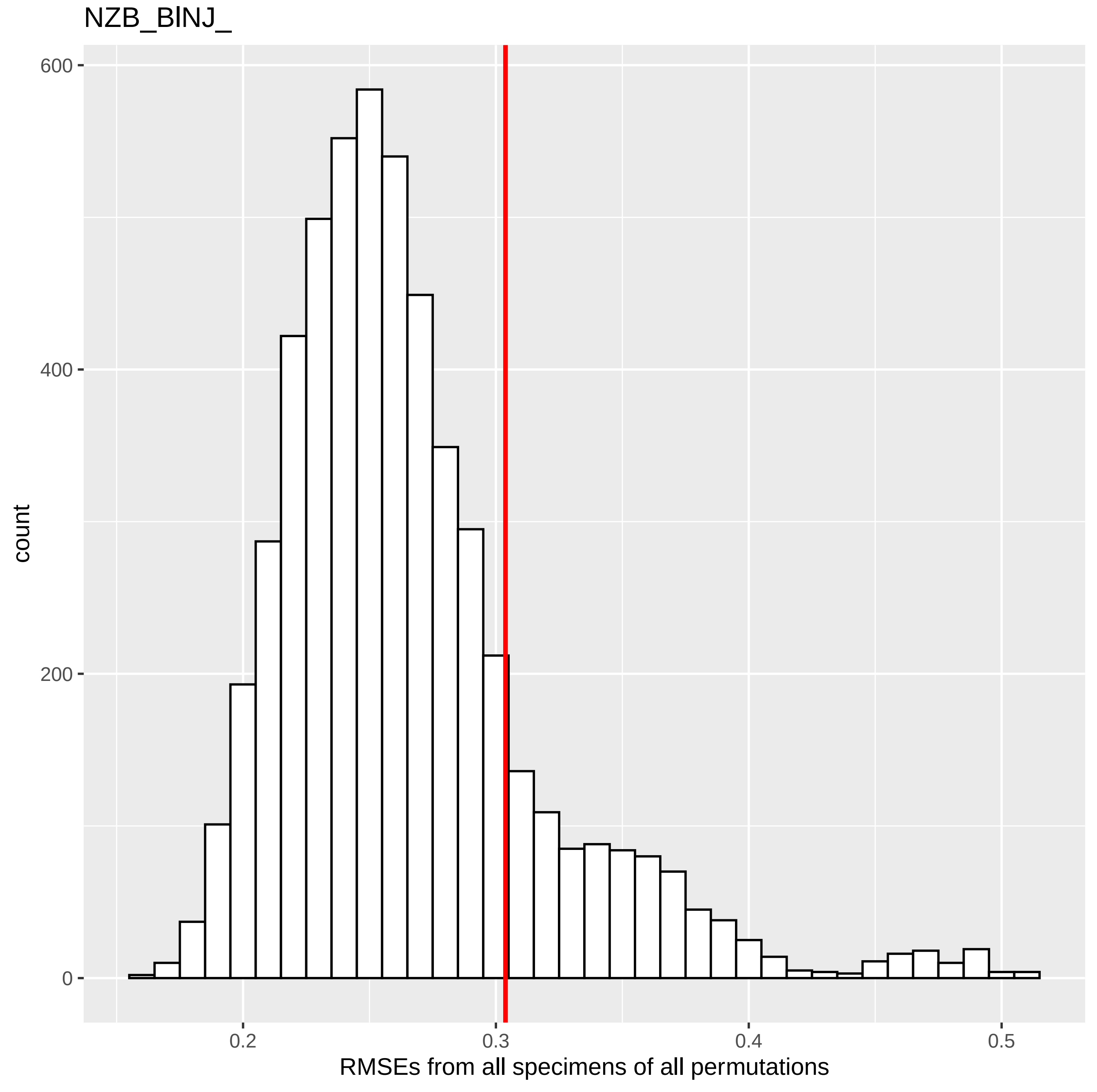

NZB_BlNJ_
600
400
200
0
count
0.2
0.3
0.4
0.5
RMSEs from all specimens of all permutations

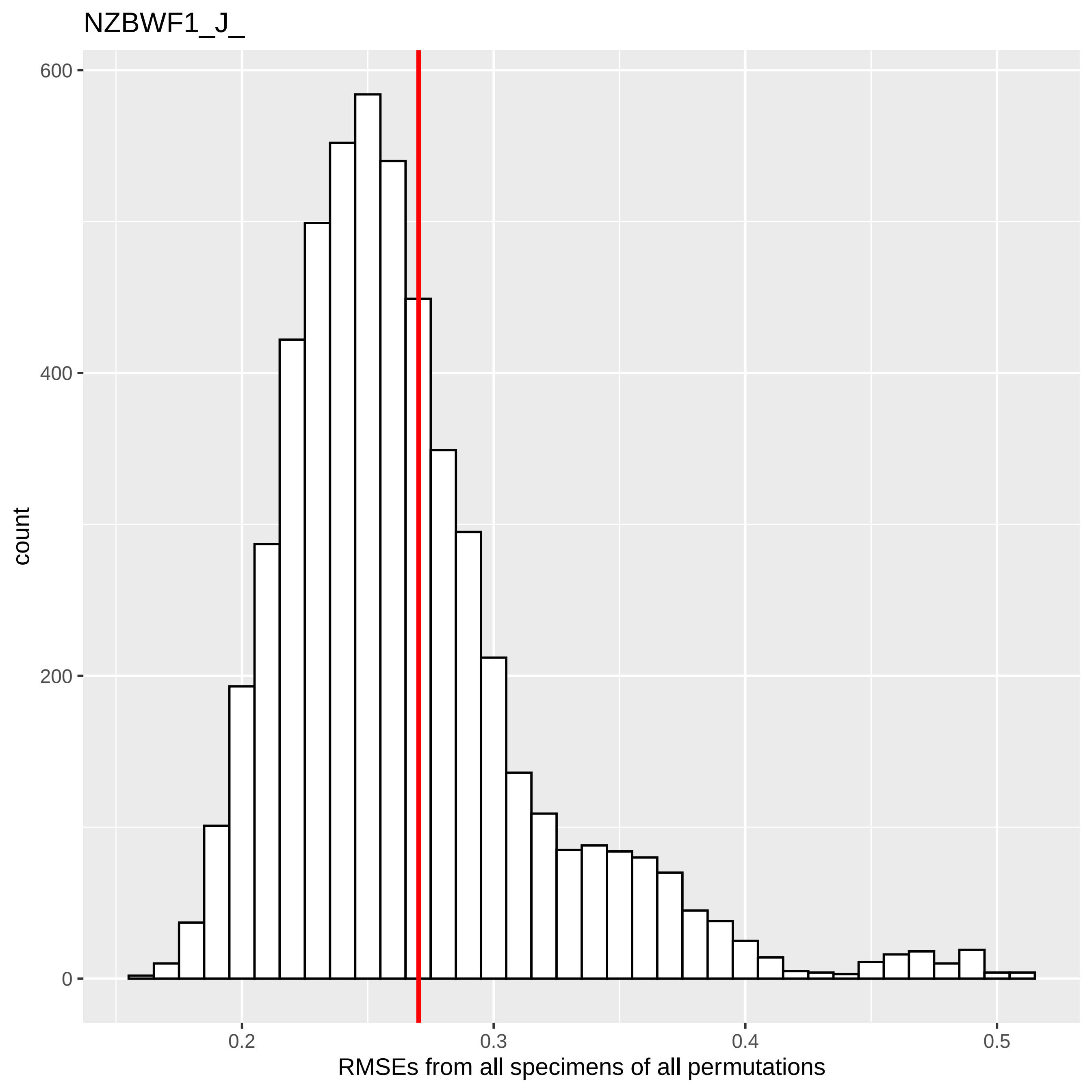

NZBWF1_J_
count
600
400
200
0
0.2
0.3
0.4
0.5
RMSEs from all specimens of all permutations

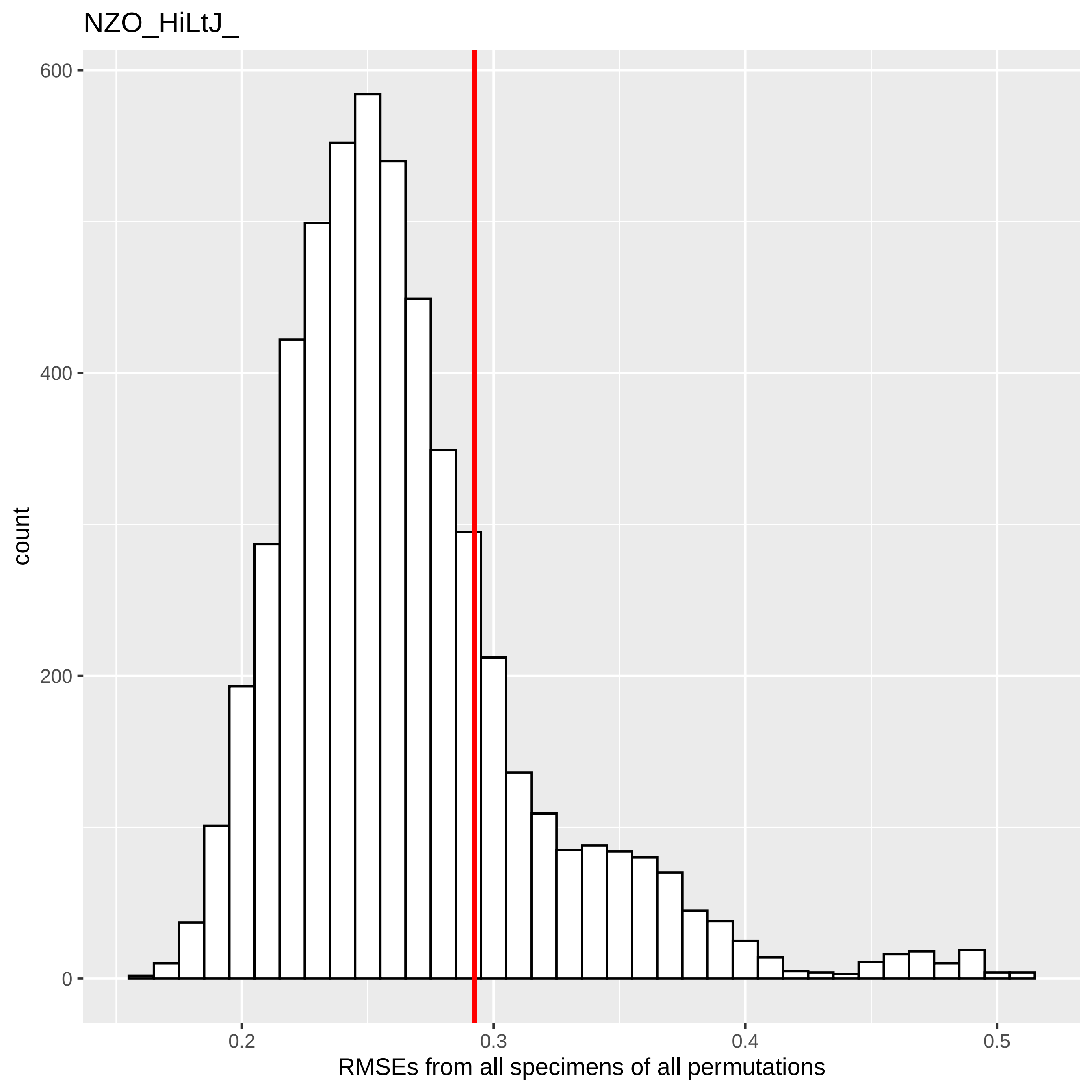
NZO_HiLtJ_
600
400
200
0
count
0.2
0.3
0.4
0.5
RMSEs from all specimens of all permutations

NZW_LACJ_

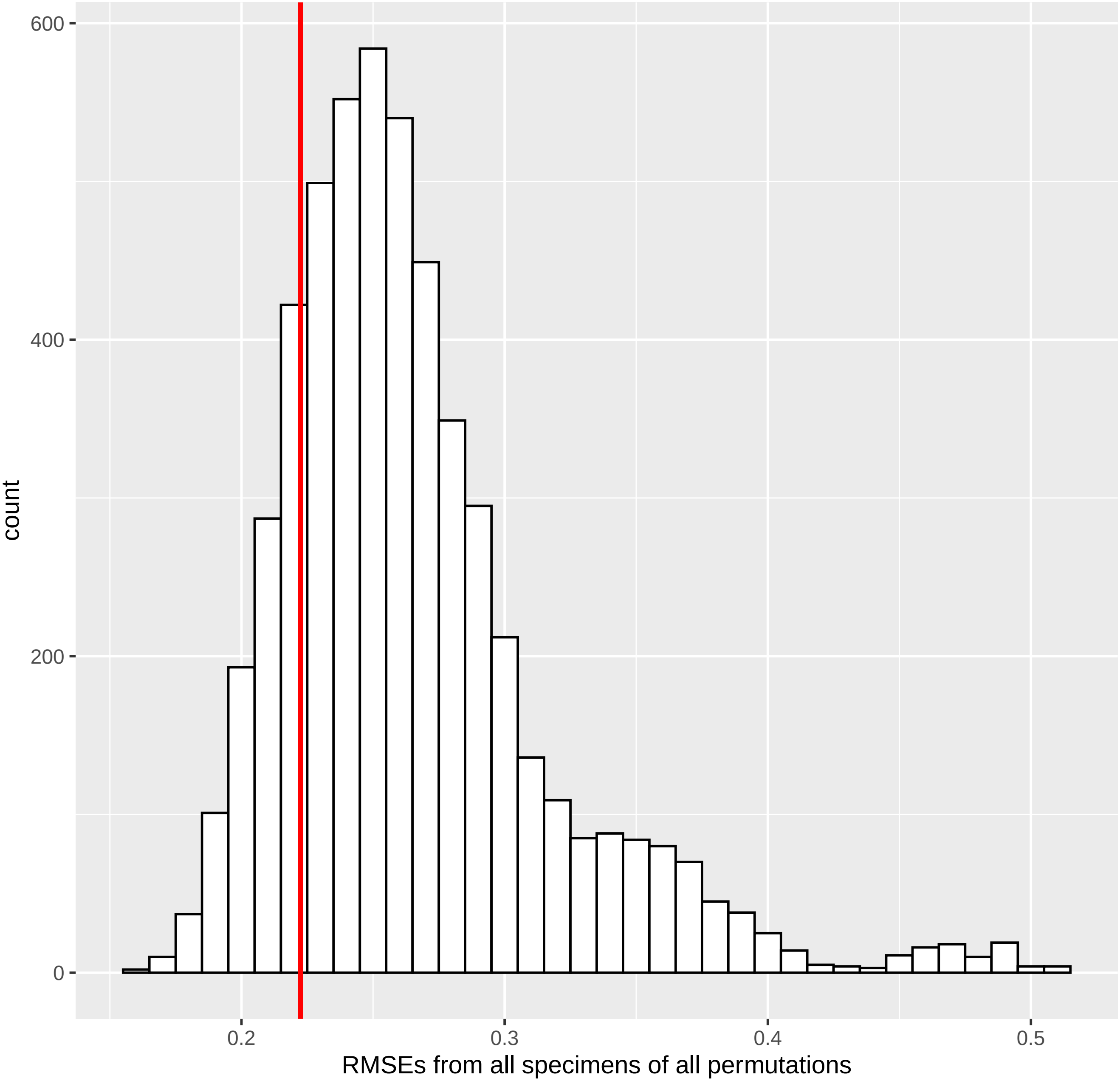

600
400
200
0
count
0.2
0.3
0.4
0.5
RMSEs from all specimens of all permutations

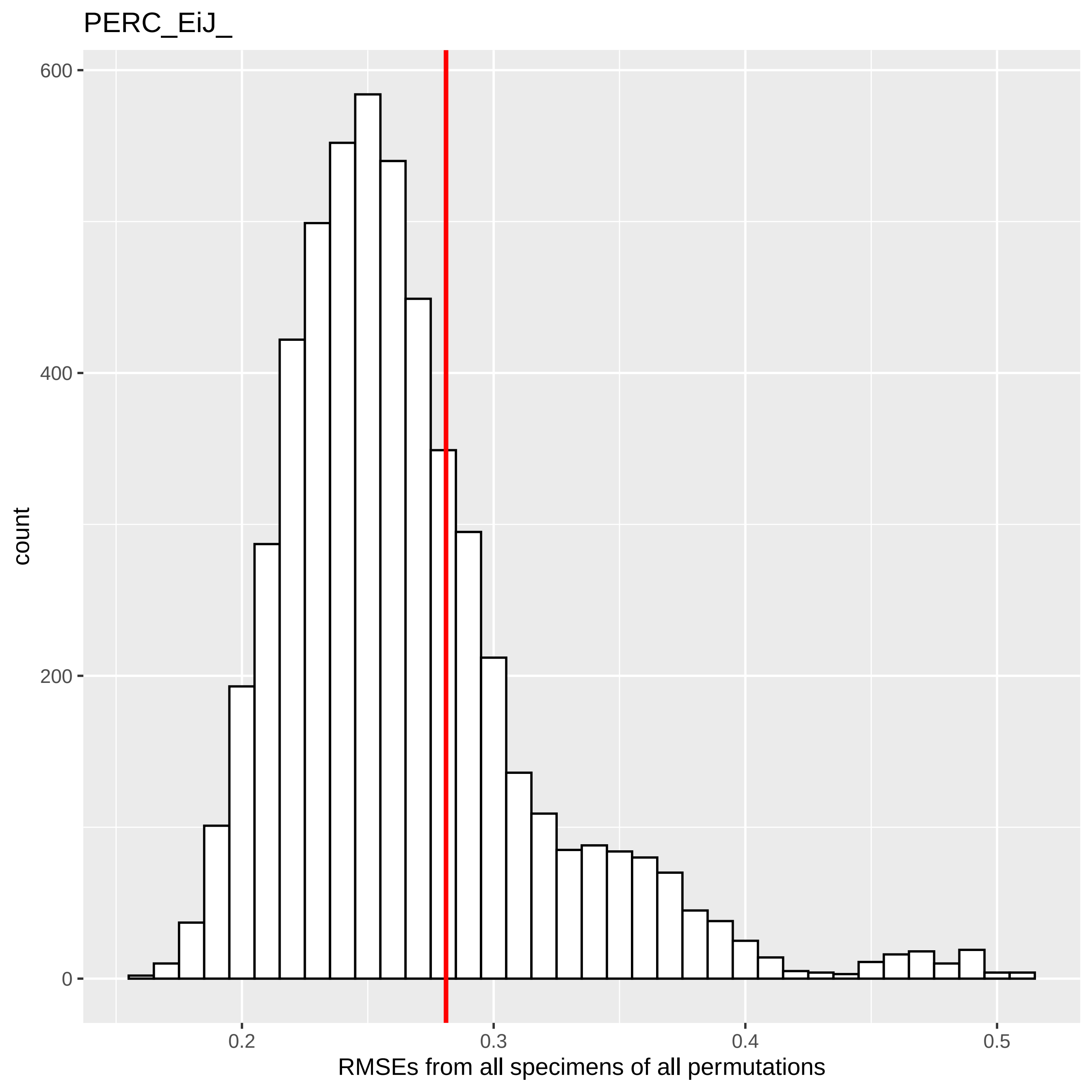
PERC_EiJ_
count
600
400
200
0
0.2
0.3
0.4
0.5
RMSEs from all specimens of all permutations

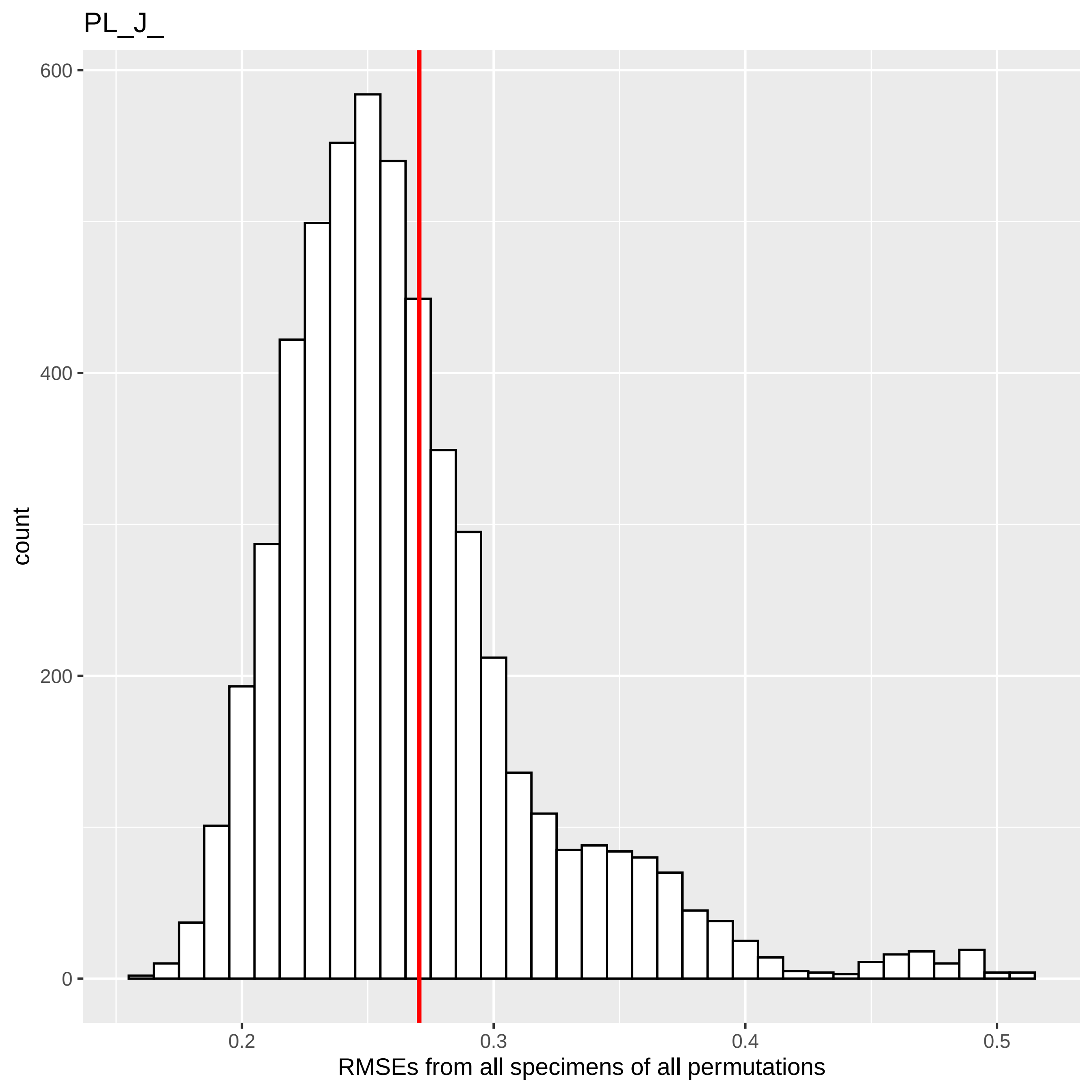

PL_J_
600
400
200
0
count
0.2
0.3
0.4
0.5
RMSEs from all specimens of all permutations

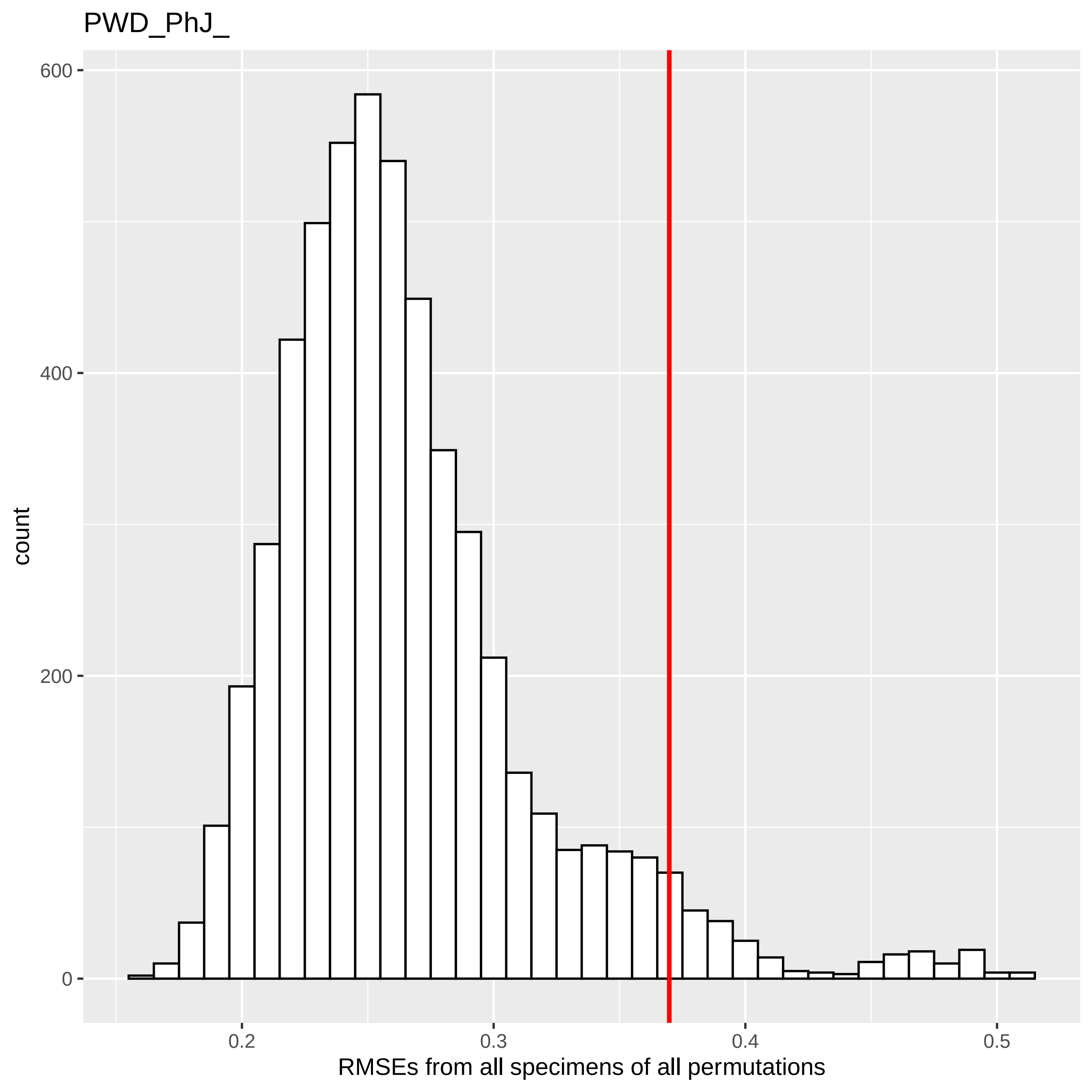

PWD_PhJ_
count
600
400
200
0
0.2
0.3
0.4
0.5
RMSEs from all specimens of all permutations

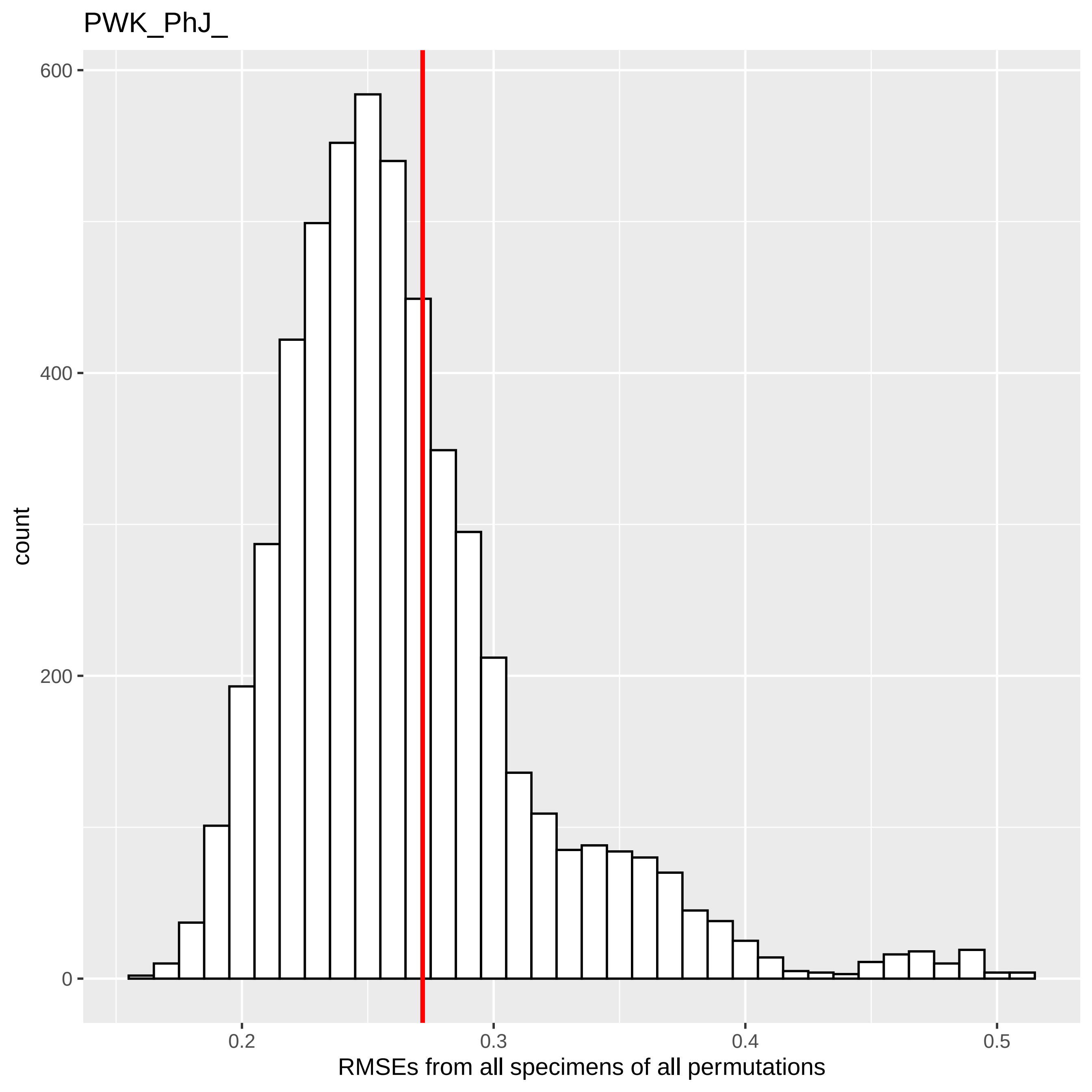
PWK_PhJ_
600
400
200
0
count
0.2
0.3
0.4
0.5
RMSEs from all specimens of all permutations

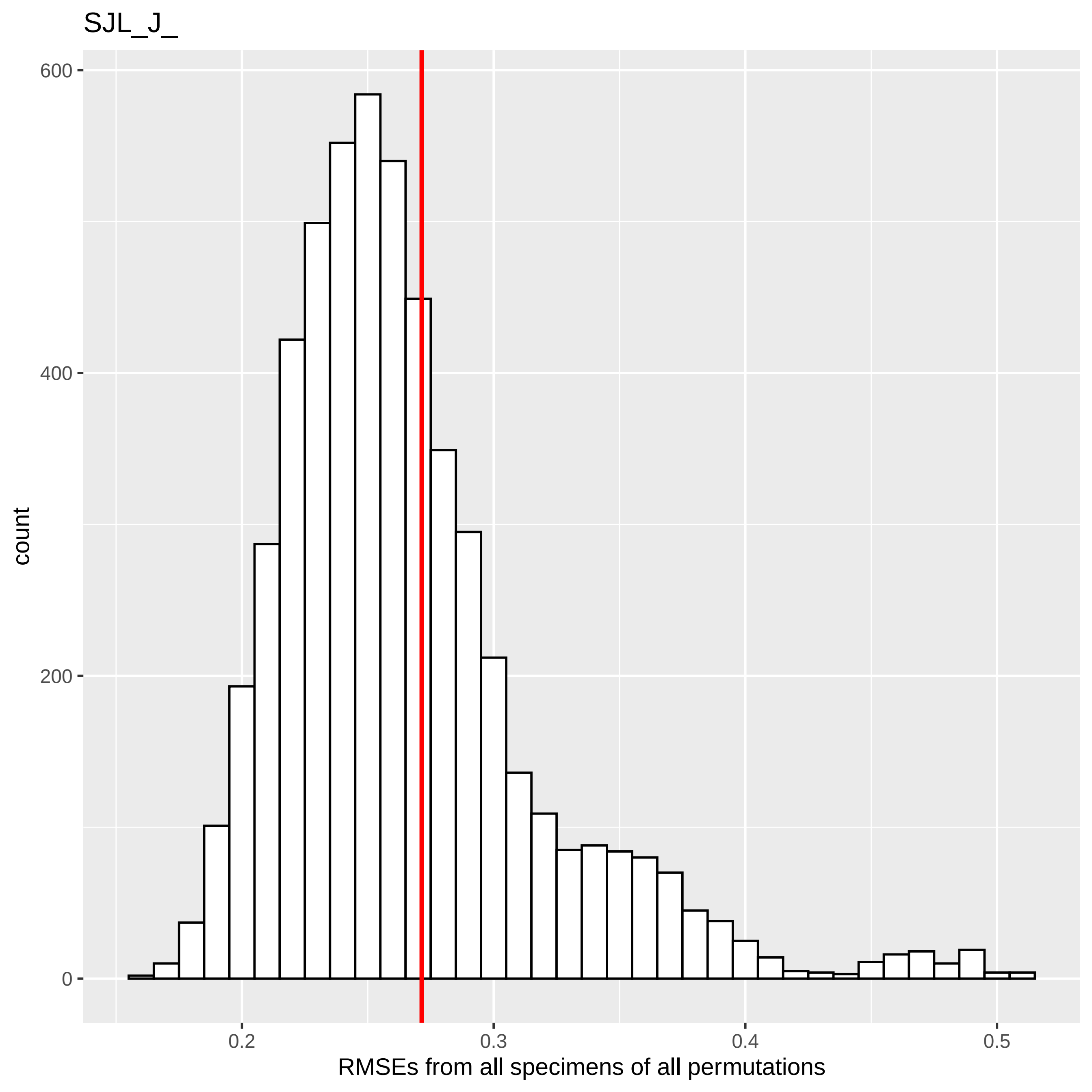
SJL_J_
count
600
400
200
0
0.2
0.3
0.4
0.5
RMSEs from all specimens of all permutations

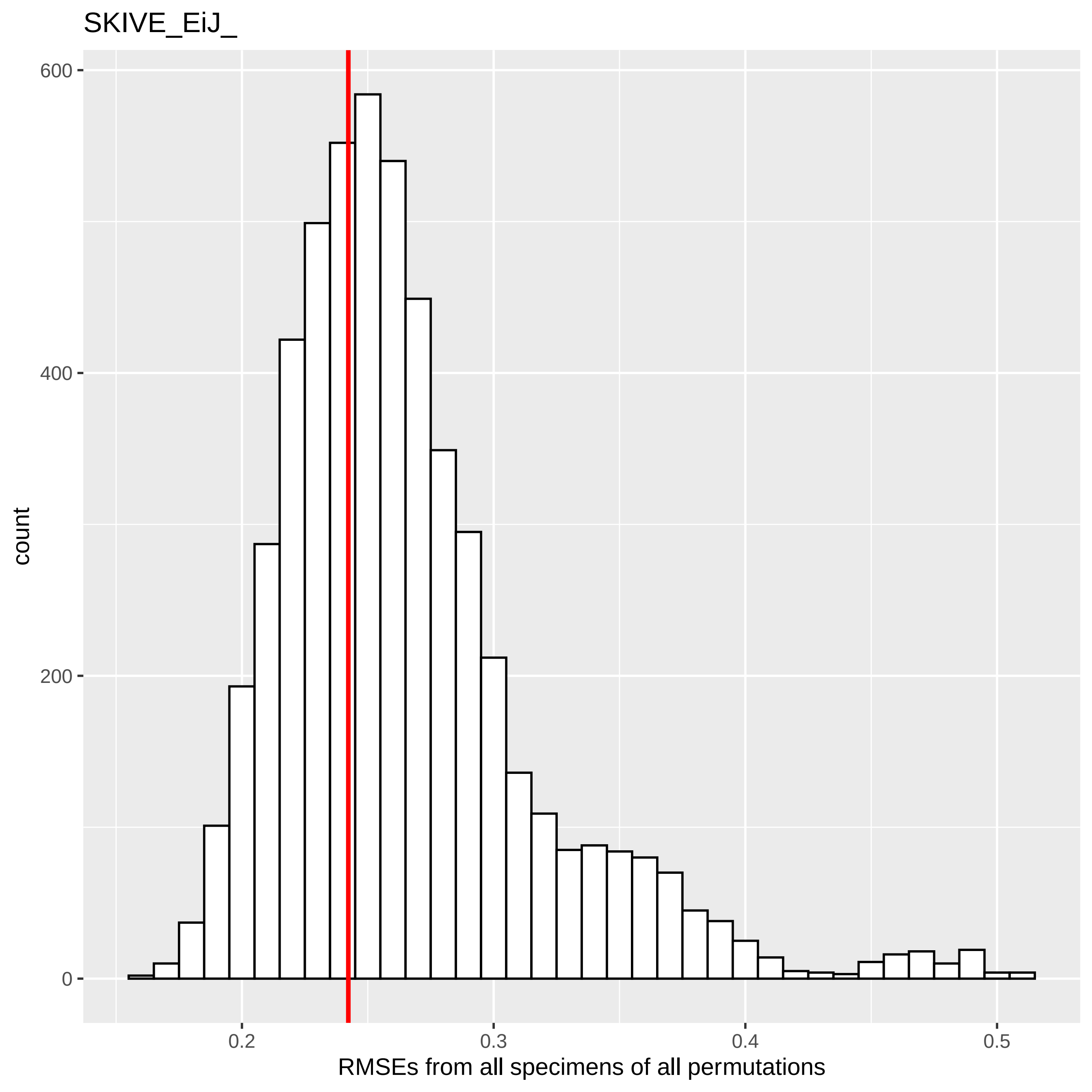

SKIVE_EiJ_
600
400
200
0
count
0.2
0.3
0.4
0.5
RMSEs from all specimens of all permutations

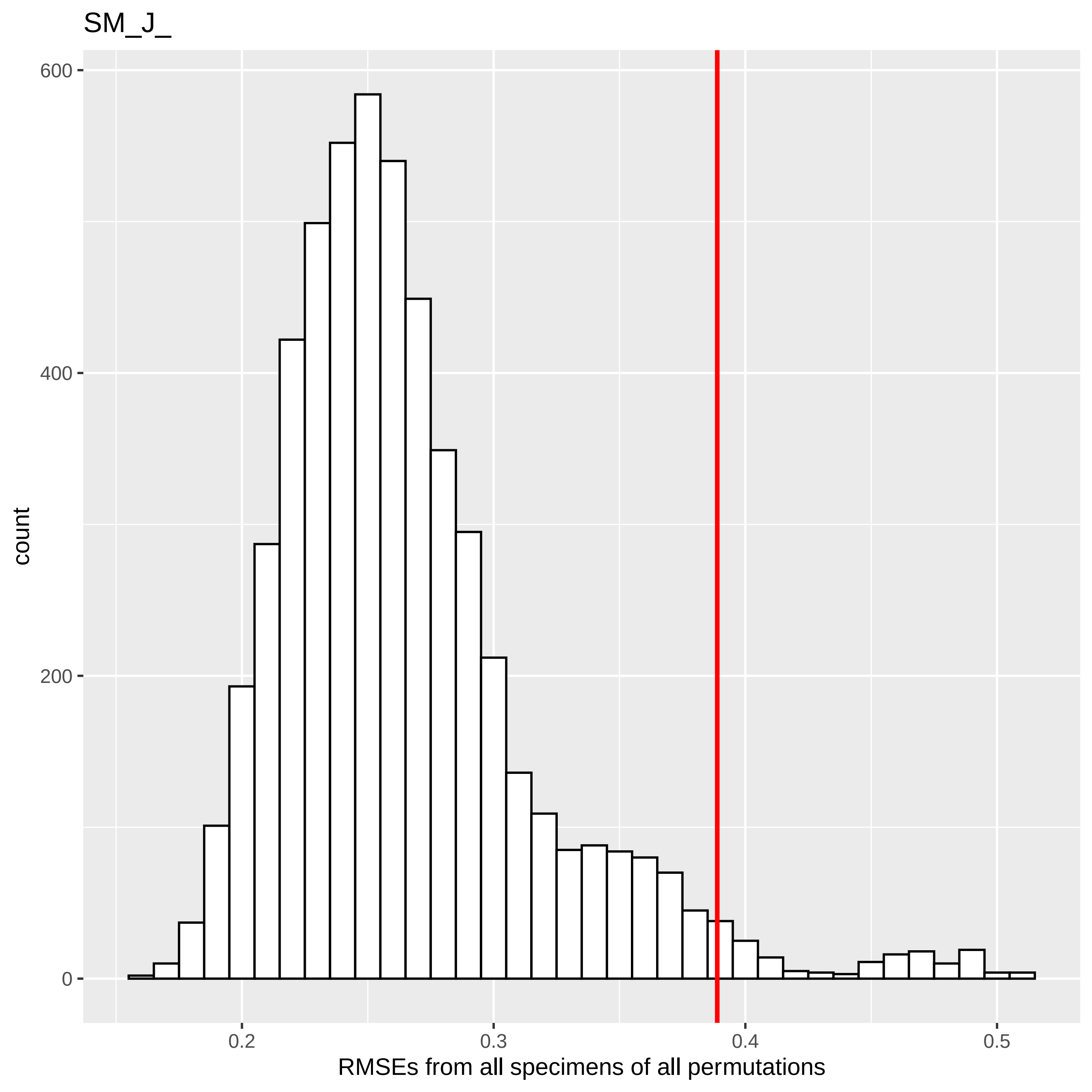
SM_J_
count
600
400
200
0
0.2
0.3
0.4
0.5
RMSEs from all specimens of all permutations

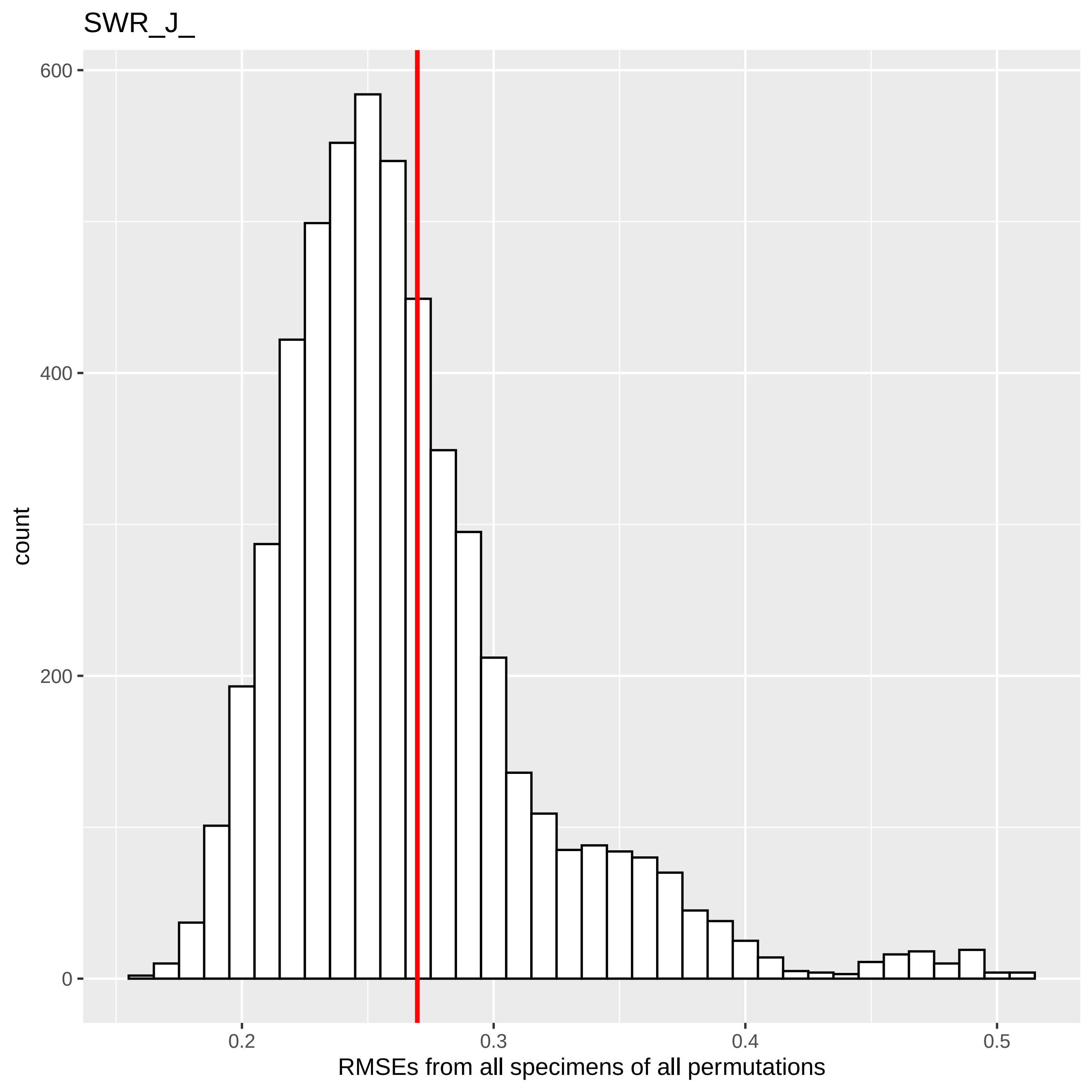
SWR_J_
count
600
400
200
0
0.2
0.3
0.4
0.5
RMSEs from all specimens of all permutations

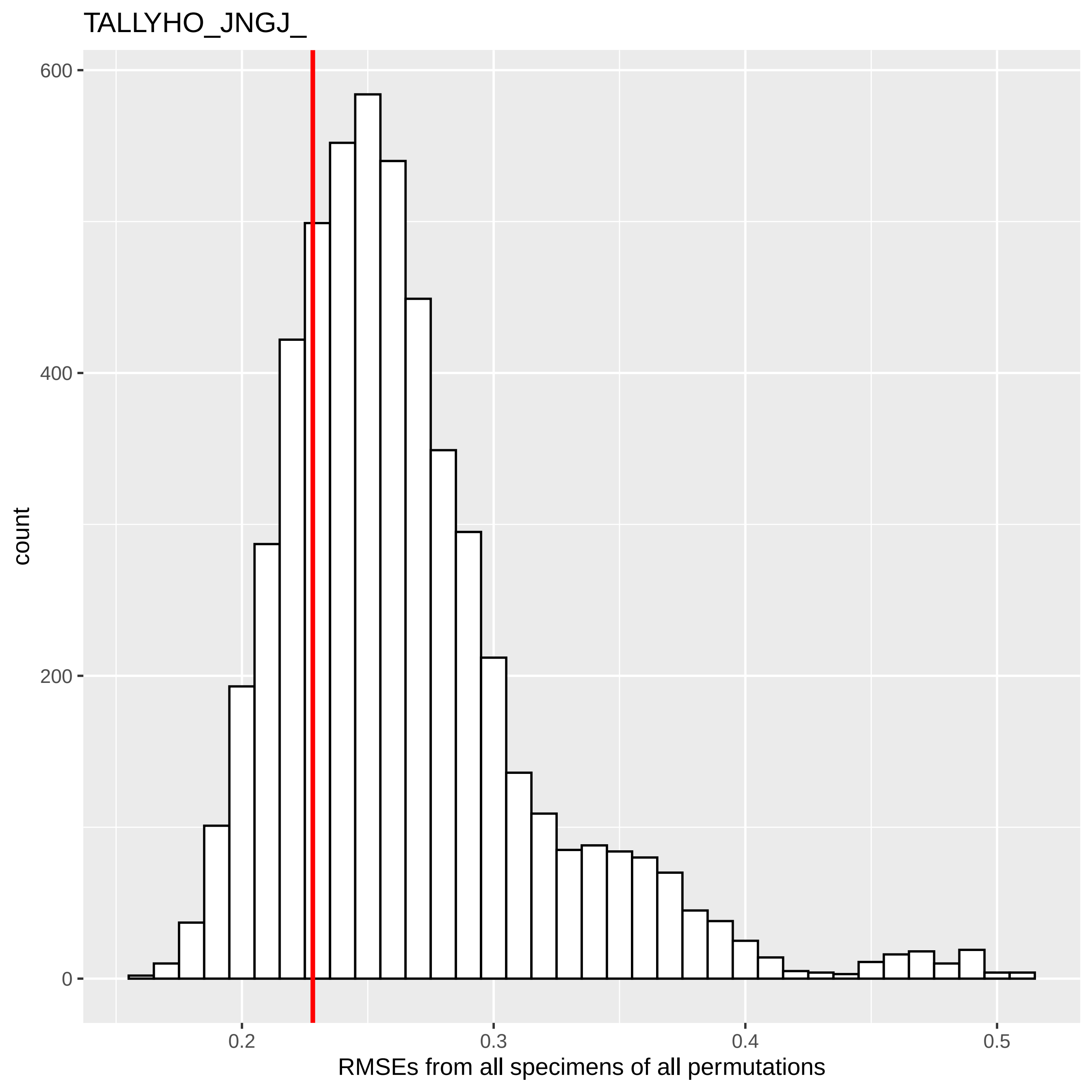
TALLYHO_JNGJ_
count
600
400
200
0
0.2
0.3
0.4
0.5
RMSEs from all specimens of all permutations
