## Supplemental Figure 7 for "Automated Landmarking via Multiple Templates"

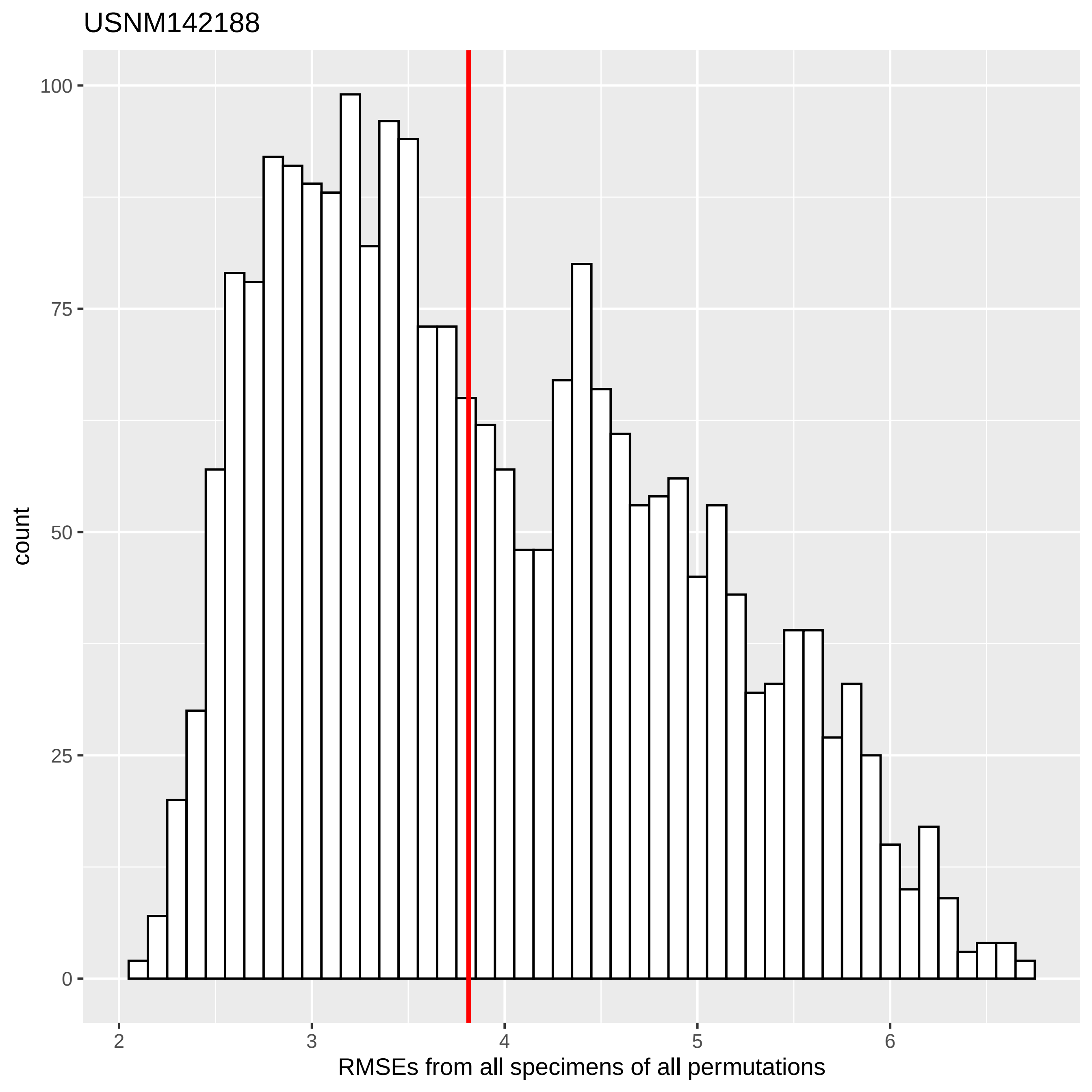
USNM142188
count
100
75
50
25
0
2
3
4
5
6
RMSEs from all specimens of all permutations

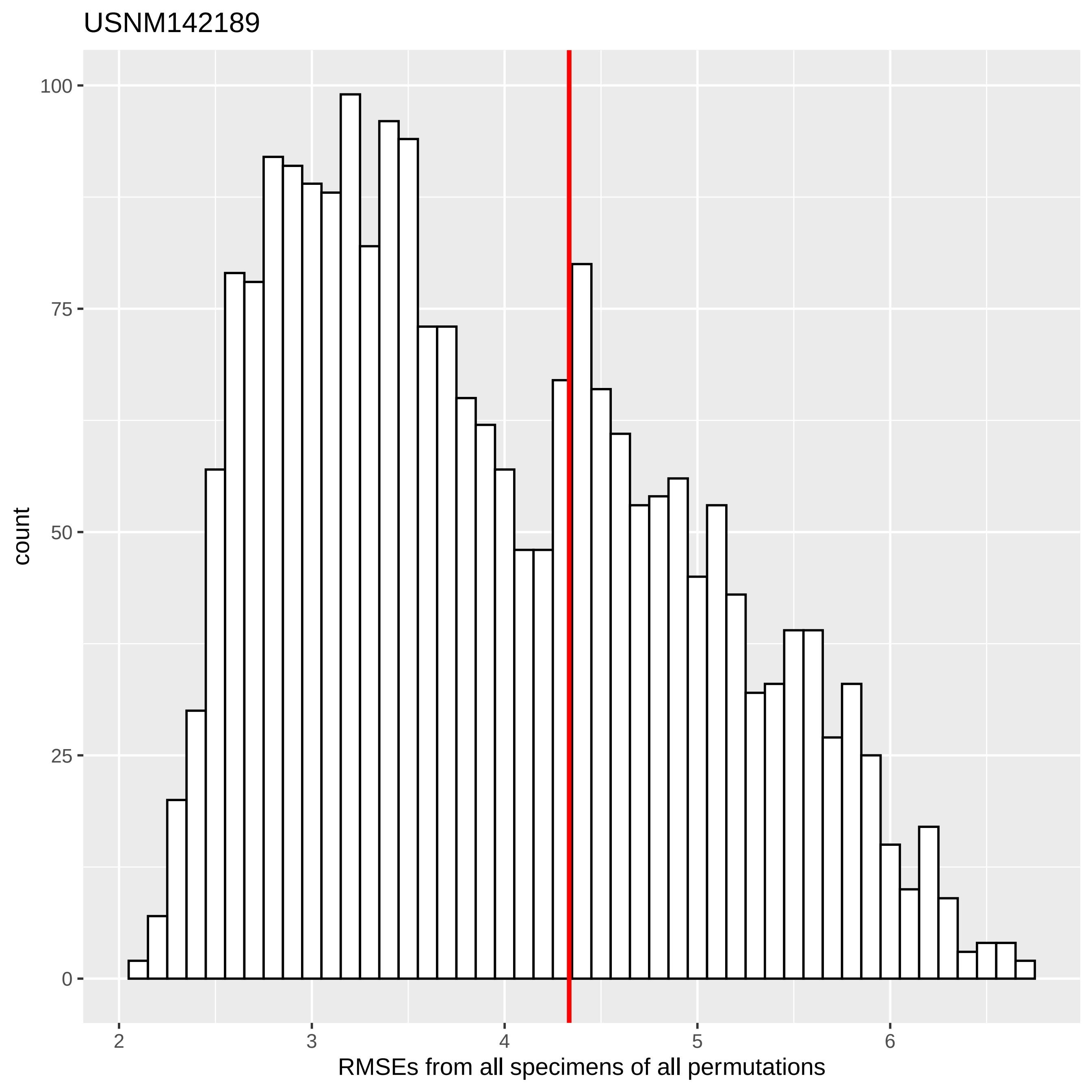

USNM142189
count
100
75
50
25
0
2
3
4
5
6
RMSEs from all specimens of all permutations

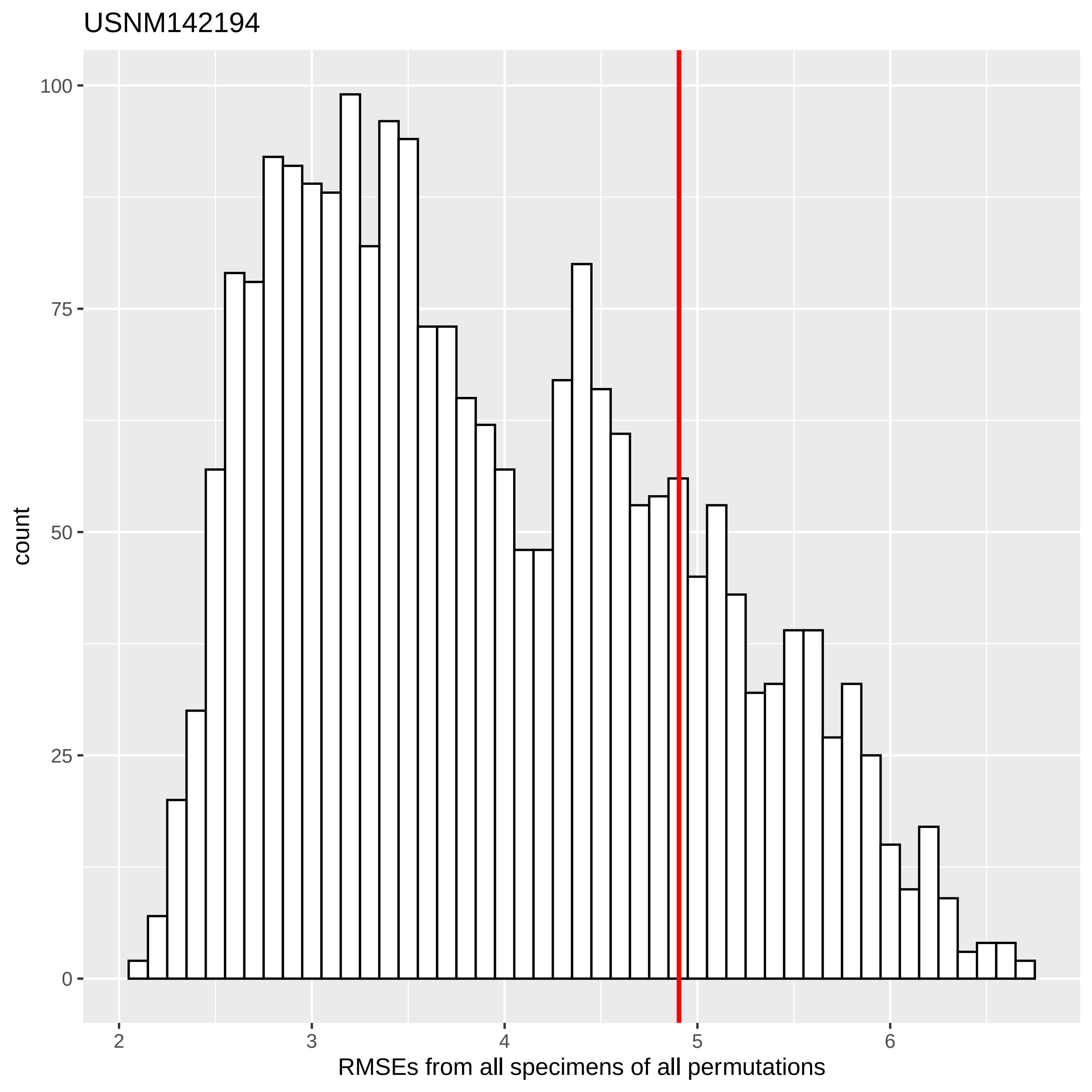
USNM142194
count
100
75
50
25
0
2
3
4
5
6
RMSEs from all specimens of all permutations

USNM145300

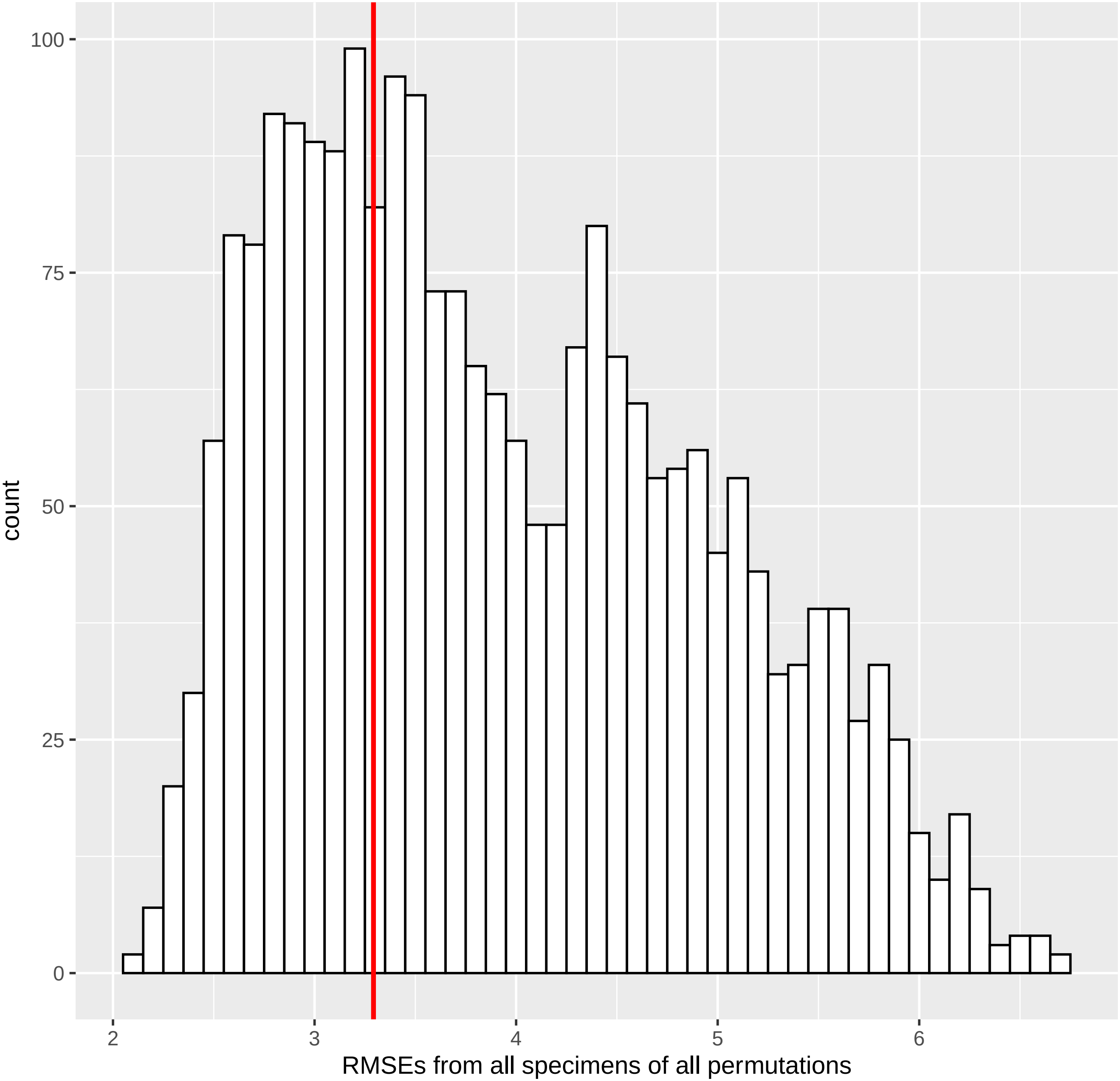

100
75
50
25
0
count
2
3
4
5
6
RMSEs from all specimens of all permutations

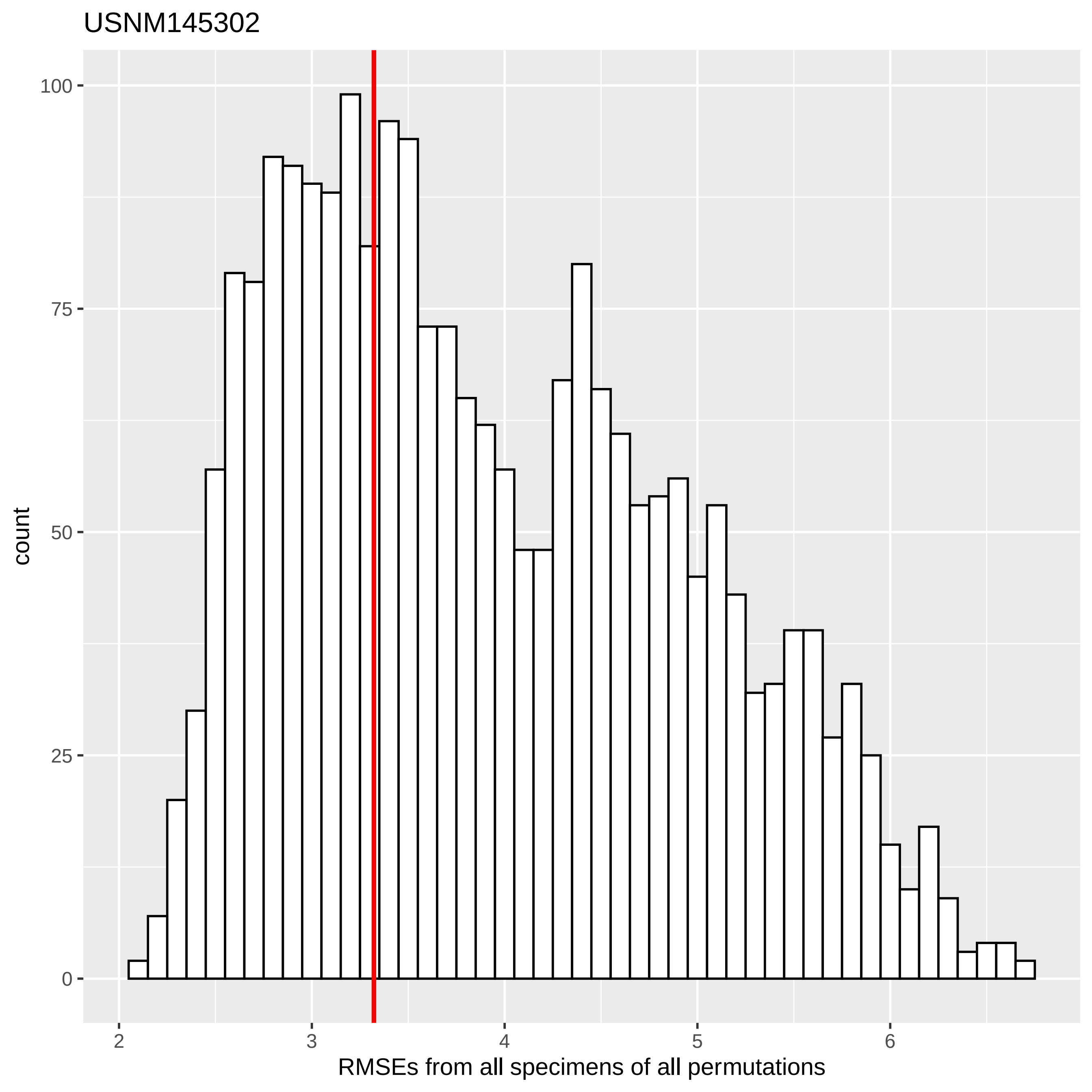

USNM145302
count
100
75
50
25
0
2
3
4
5
6
RMSEs from all specimens of all permutations

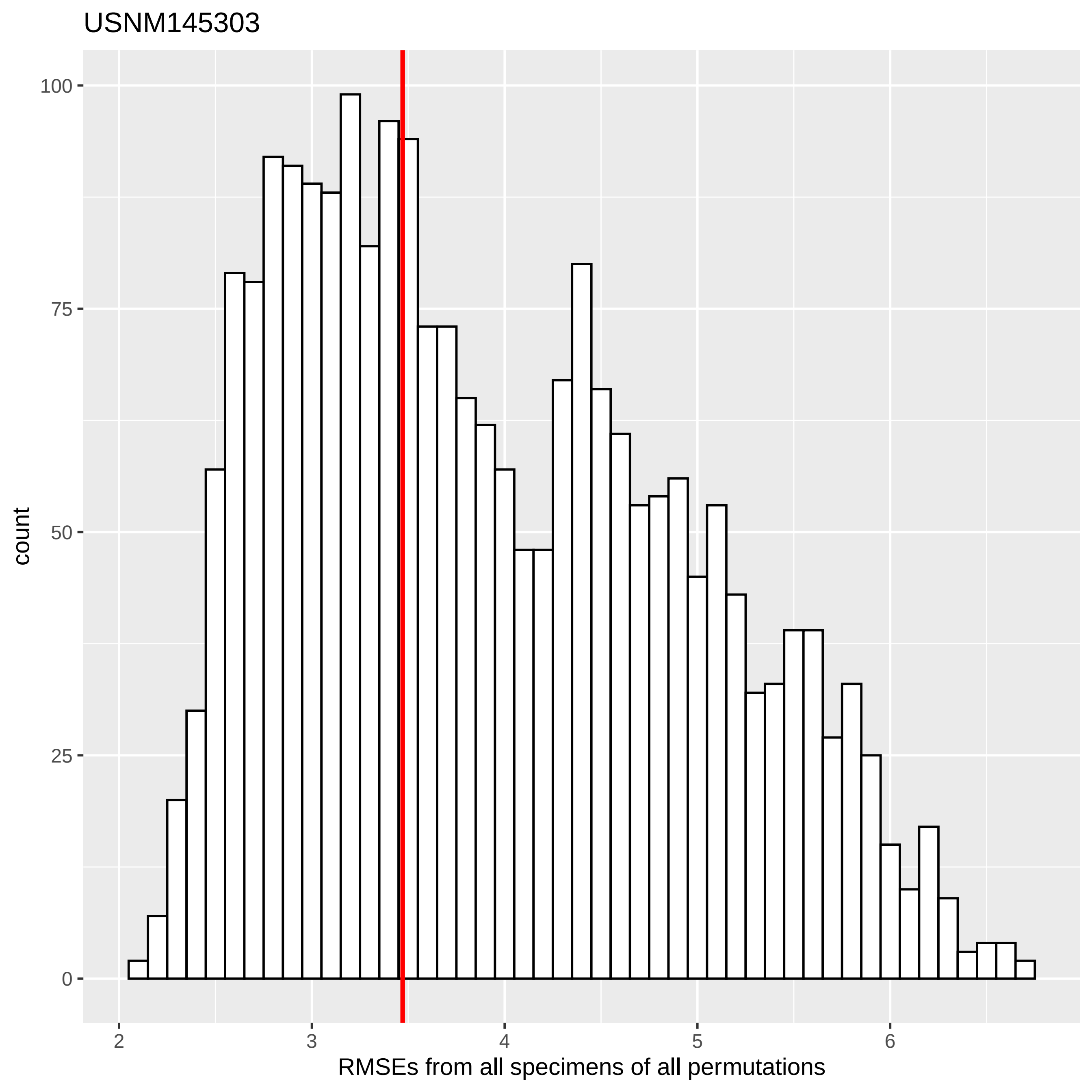
USNM145303
count
100
75
50
25
0
2
3
4
5
6
RMSEs from all specimens of all permutations

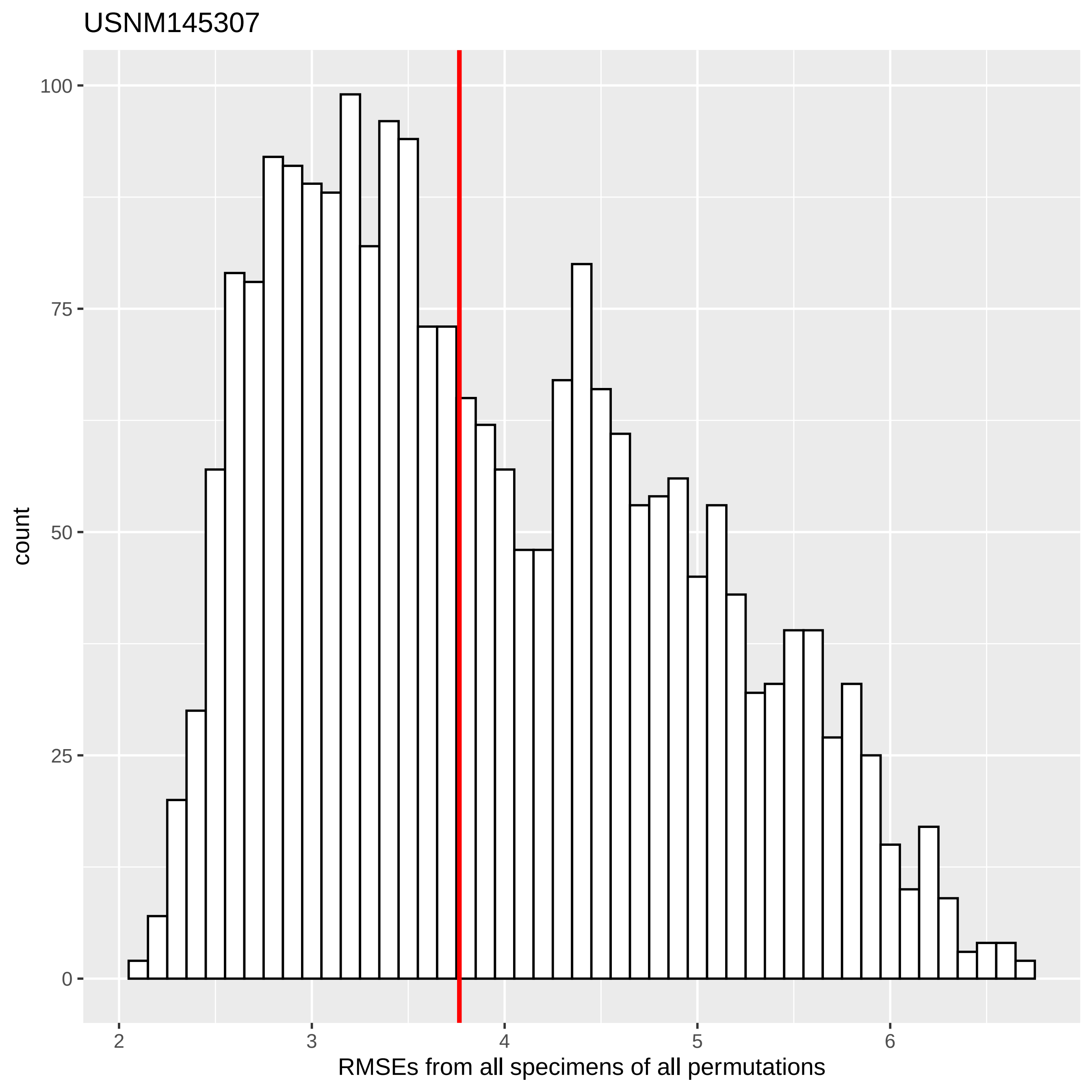

USNM145307
count
100
75
50
25
0
2
3
4
5
6
RMSEs from all specimens of all permutations

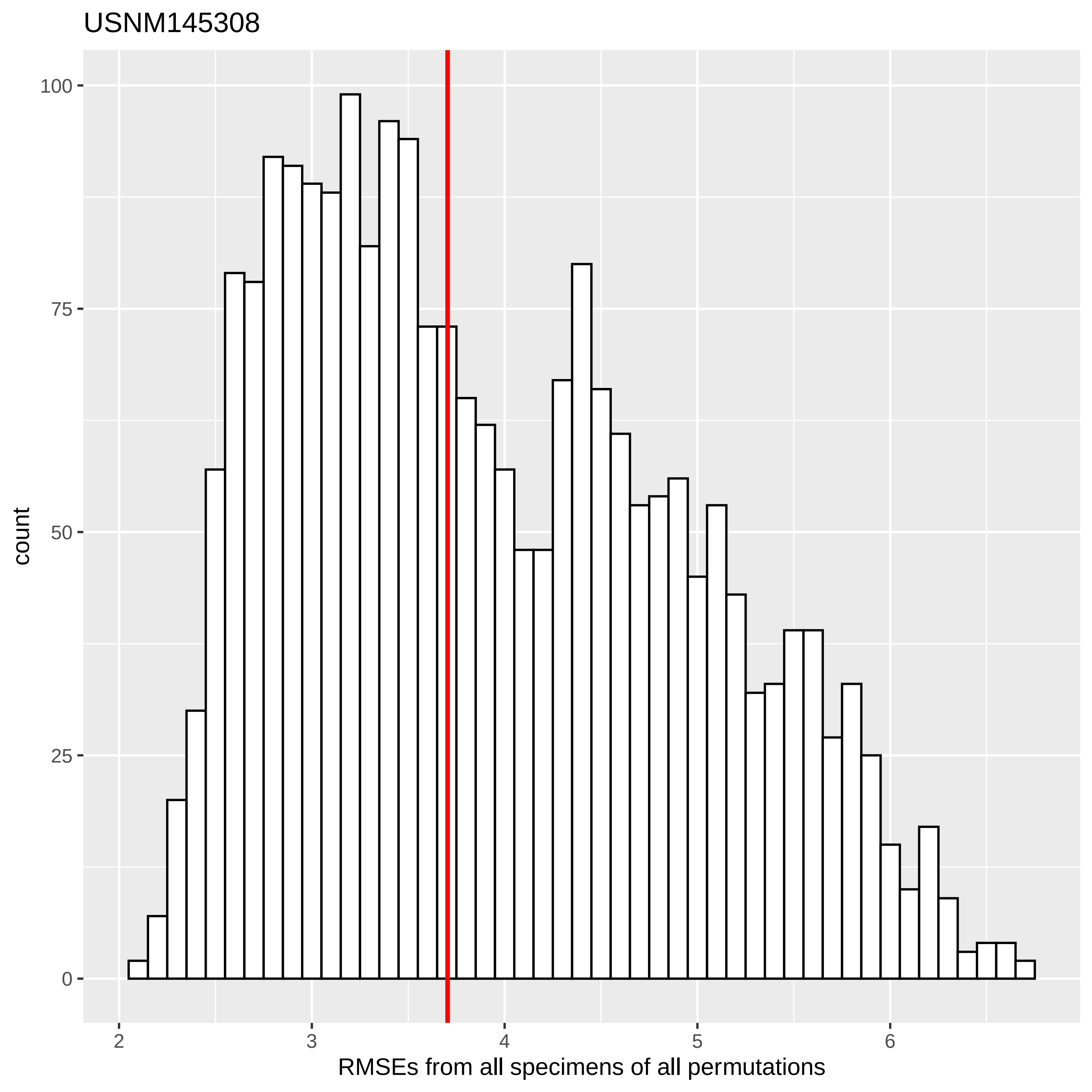

USNM145308
count
100
75
50
25
0
2
3
4
5
6
RMSEs from all specimens of all permutations

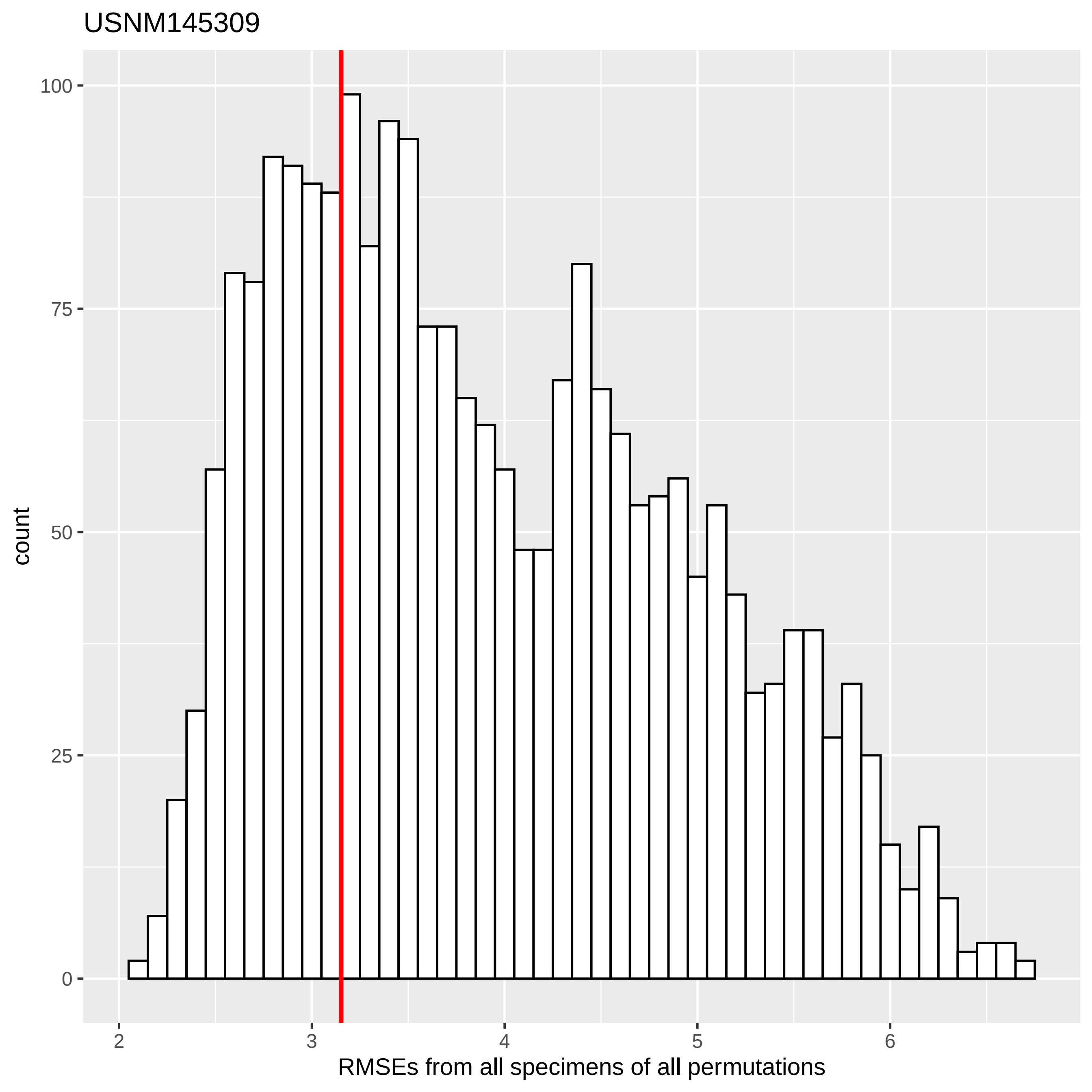
USNM145309
count
100
75
50
25
0
2
3
4
5
6
RMSEs from all specimens of all permutations

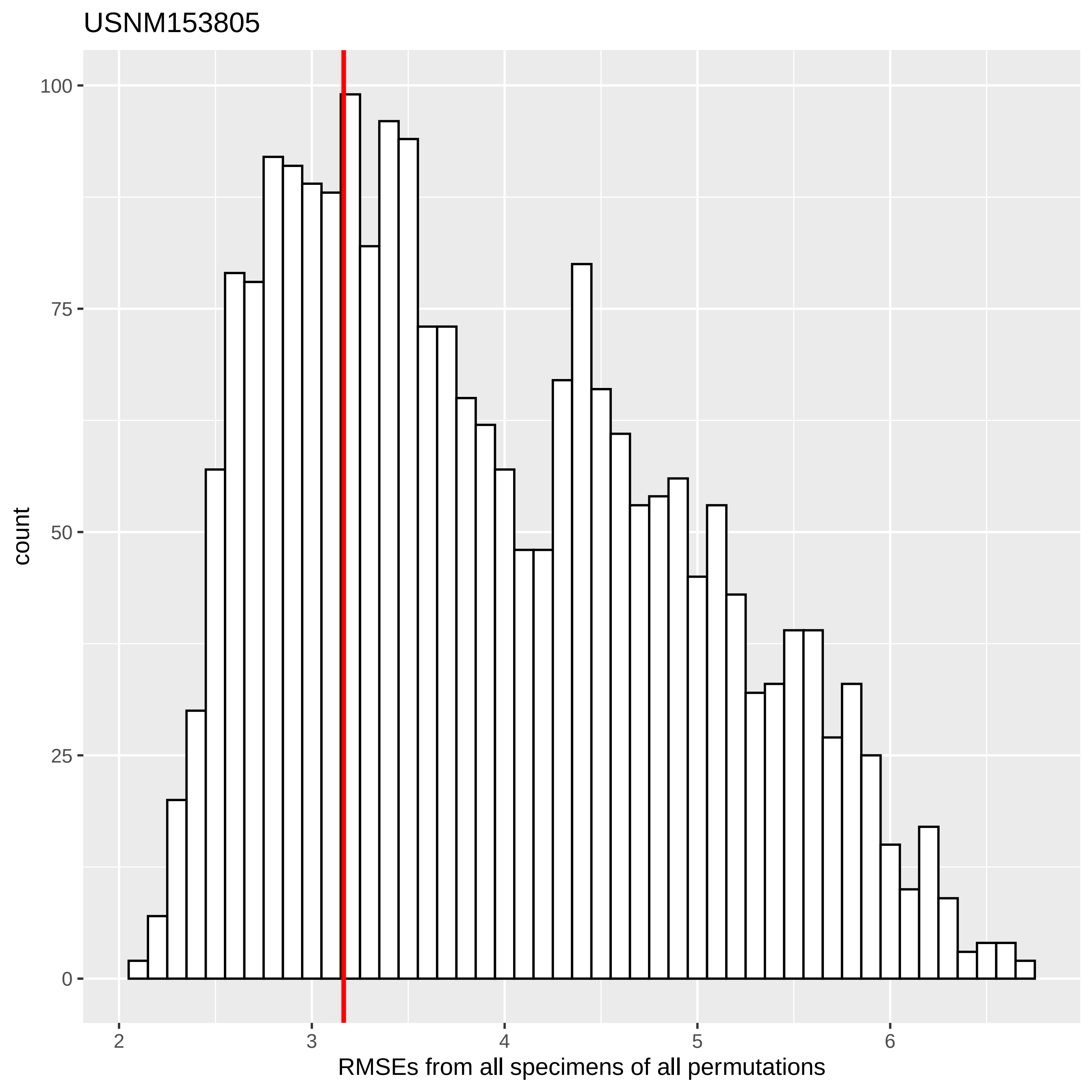

USNM153805
count
100
75
50
25
0
2
3
4
5
6
RMSEs from all specimens of all permutations

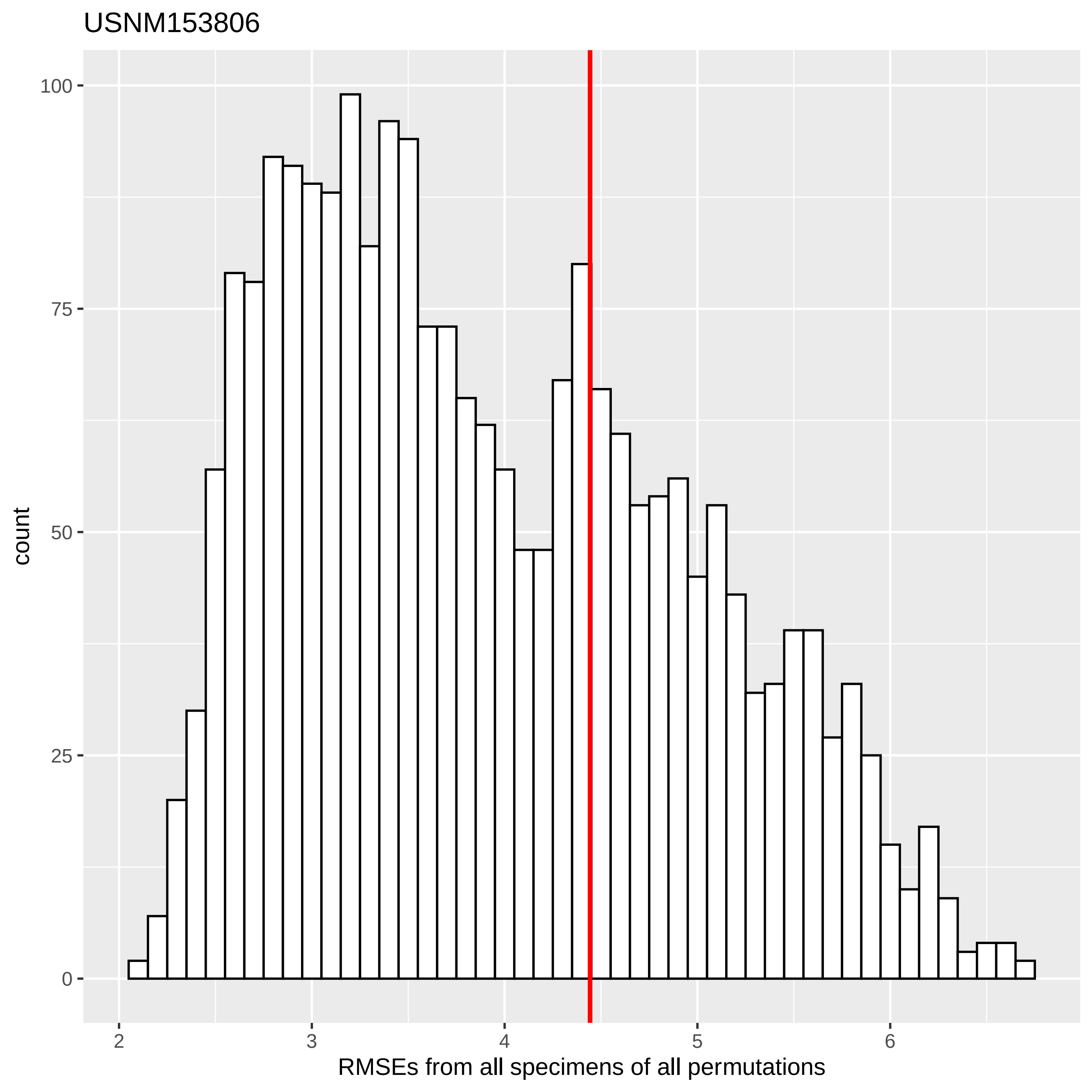
USNM153806
count
100
75
50
25
0
2
3
4
5
6
RMSEs from all specimens of all permutations

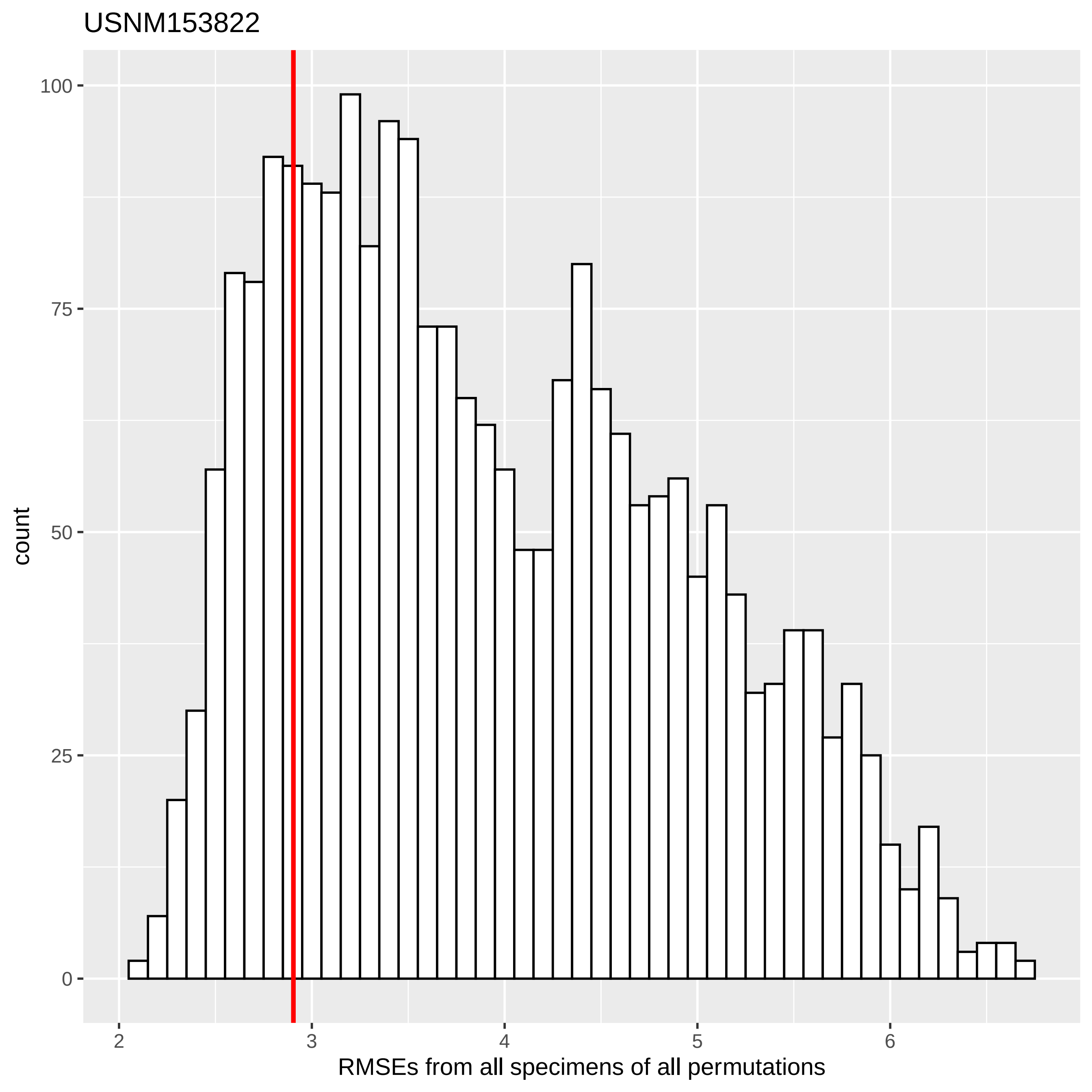

USNM153822
count
100
75
50
25
0
2
3
4
5
6
RMSEs from all specimens of all permutations

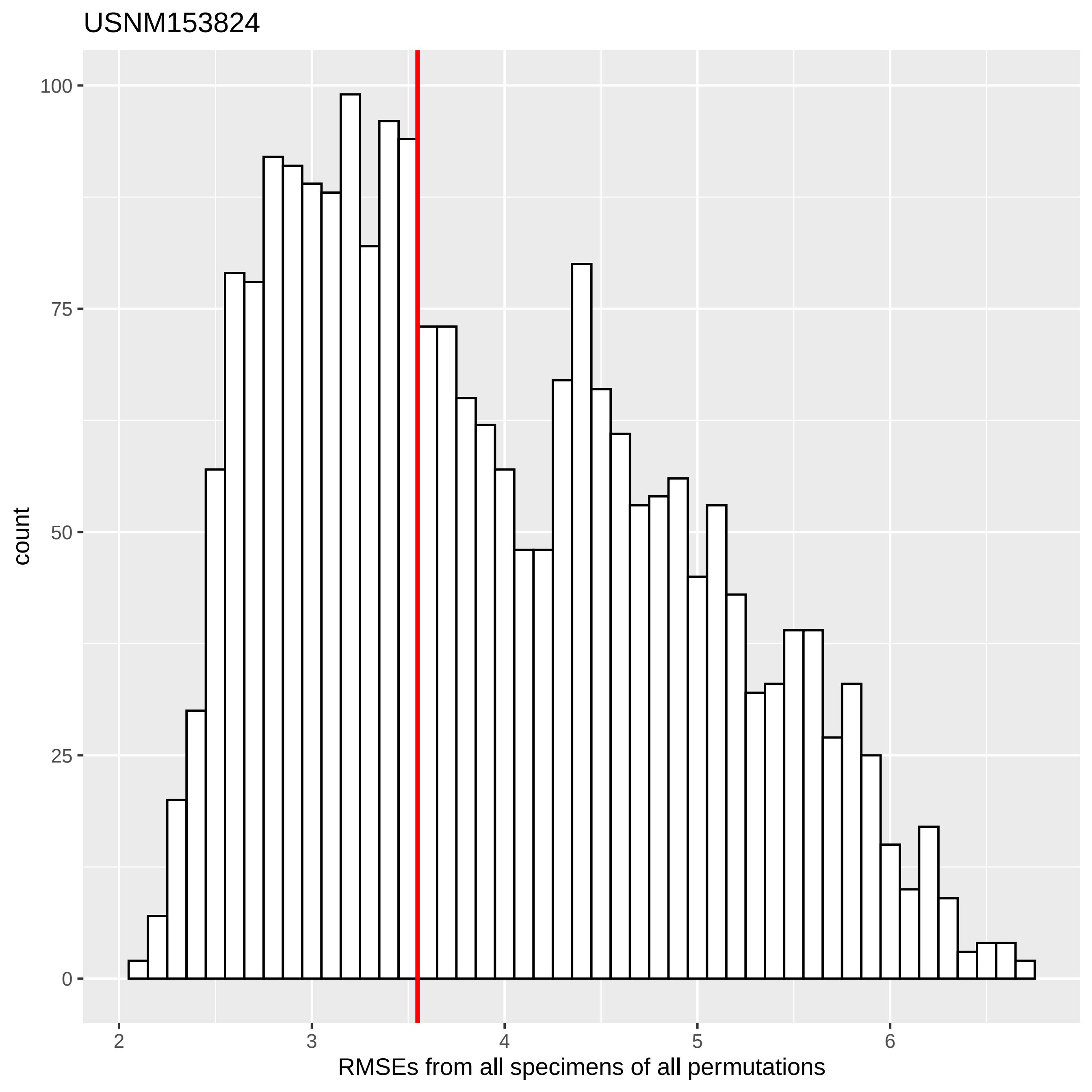

USNM153824
count
100
75
50
25
0
2
3
4
5
6
RMSEs from all specimens of all permutations

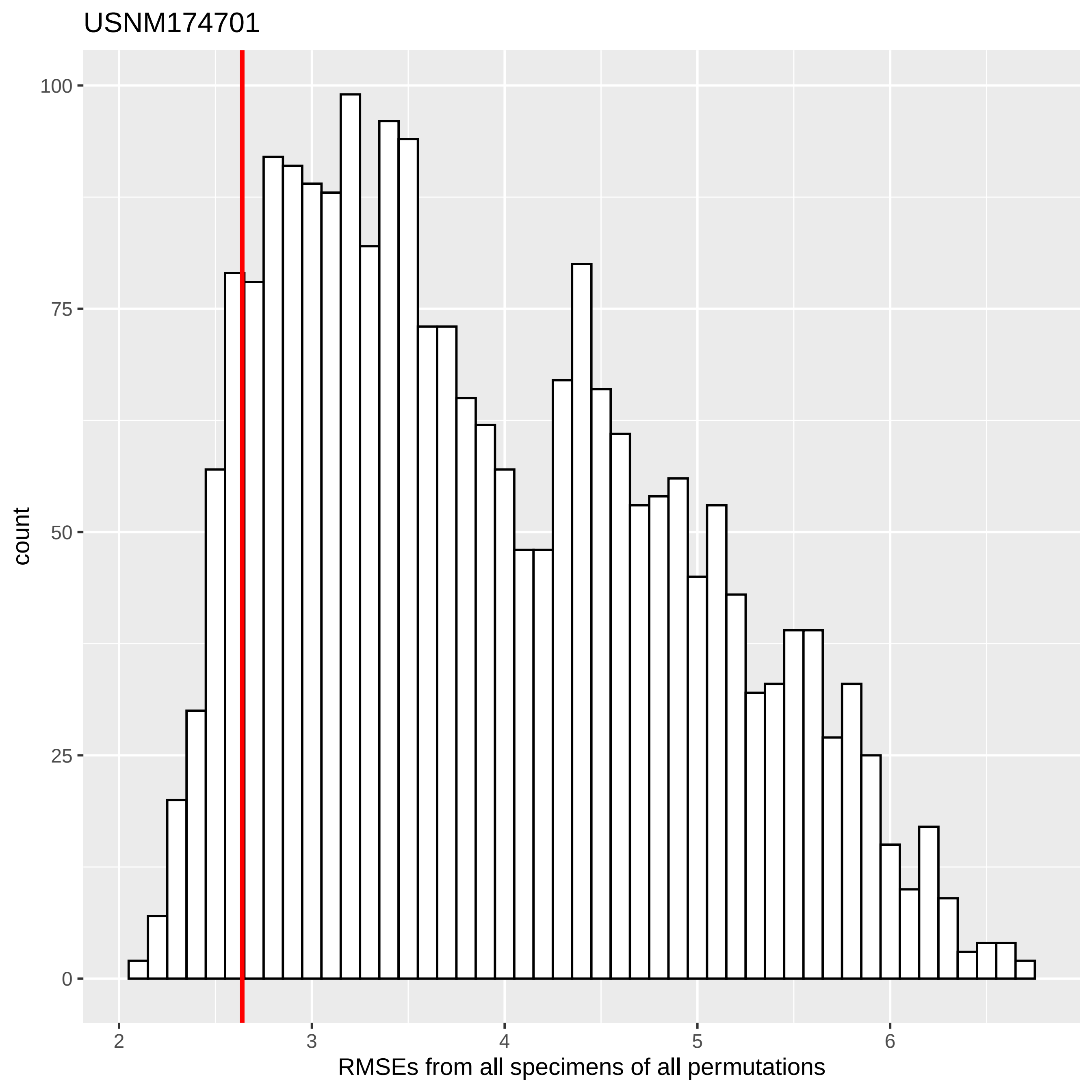
USNM174701
count
RMSEs from all specimens of all permutations
0
25
50
75
100
2
3
4
5
6

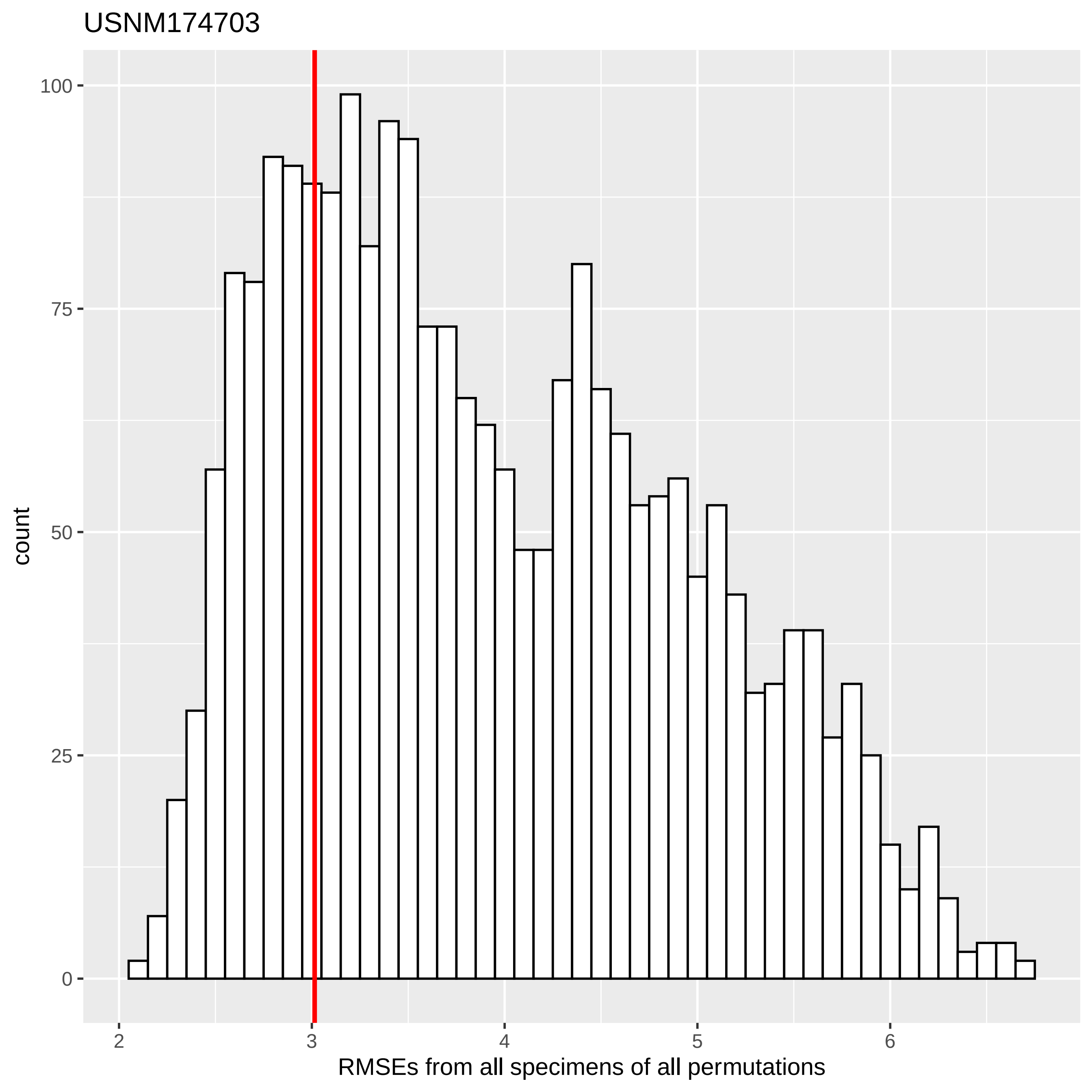
USNM174703
count
RMSEs from all specimens of all permutations
0
25
50
75
100
2
3
4
5
6

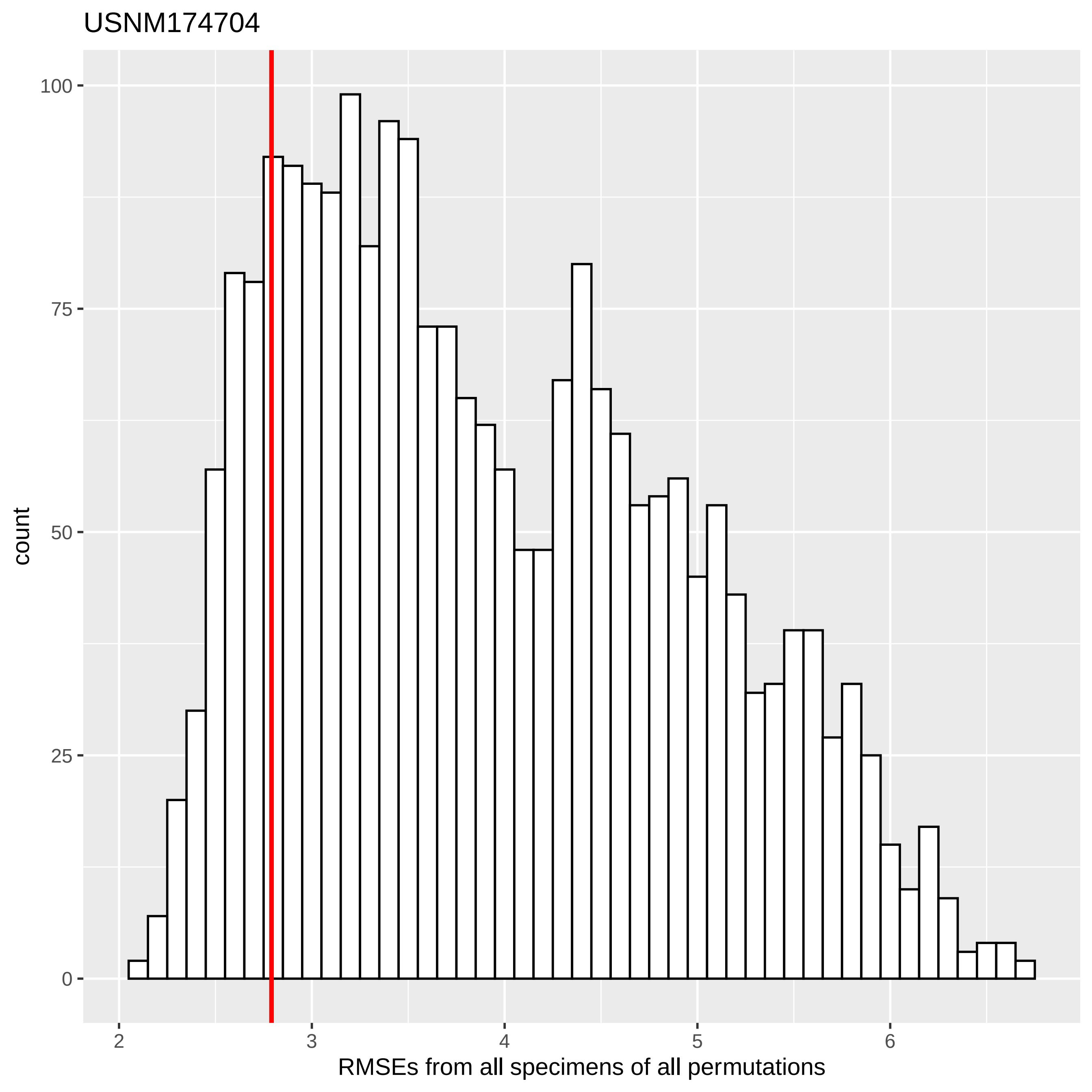
USNM174704
count
RMSEs from all specimens of all permutations
0
25
50
75
100
2
3
4
5
6

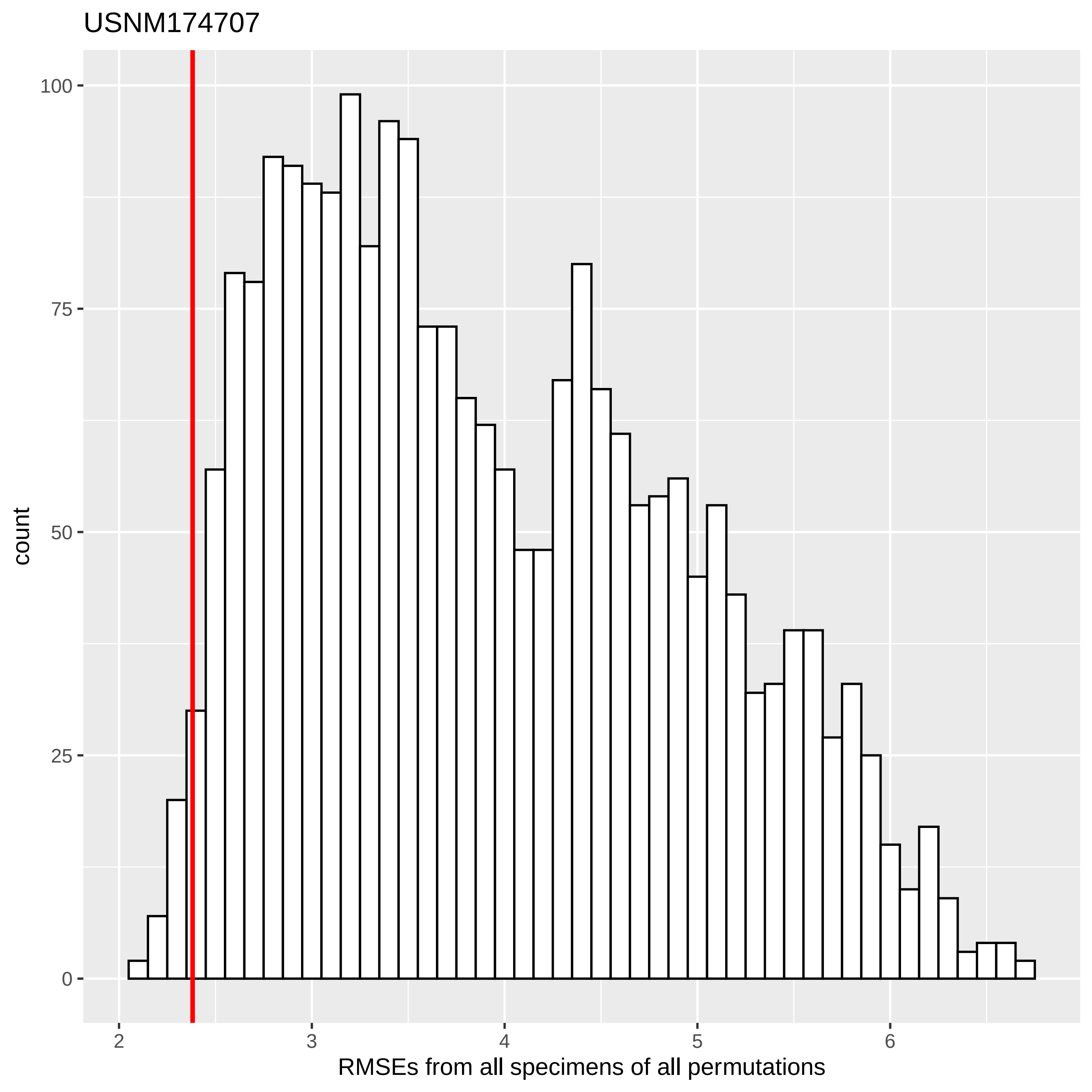

USNM174707
count
100
75
50
25
0
2
3
4
5
6
RMSEs from all specimens of all permutations

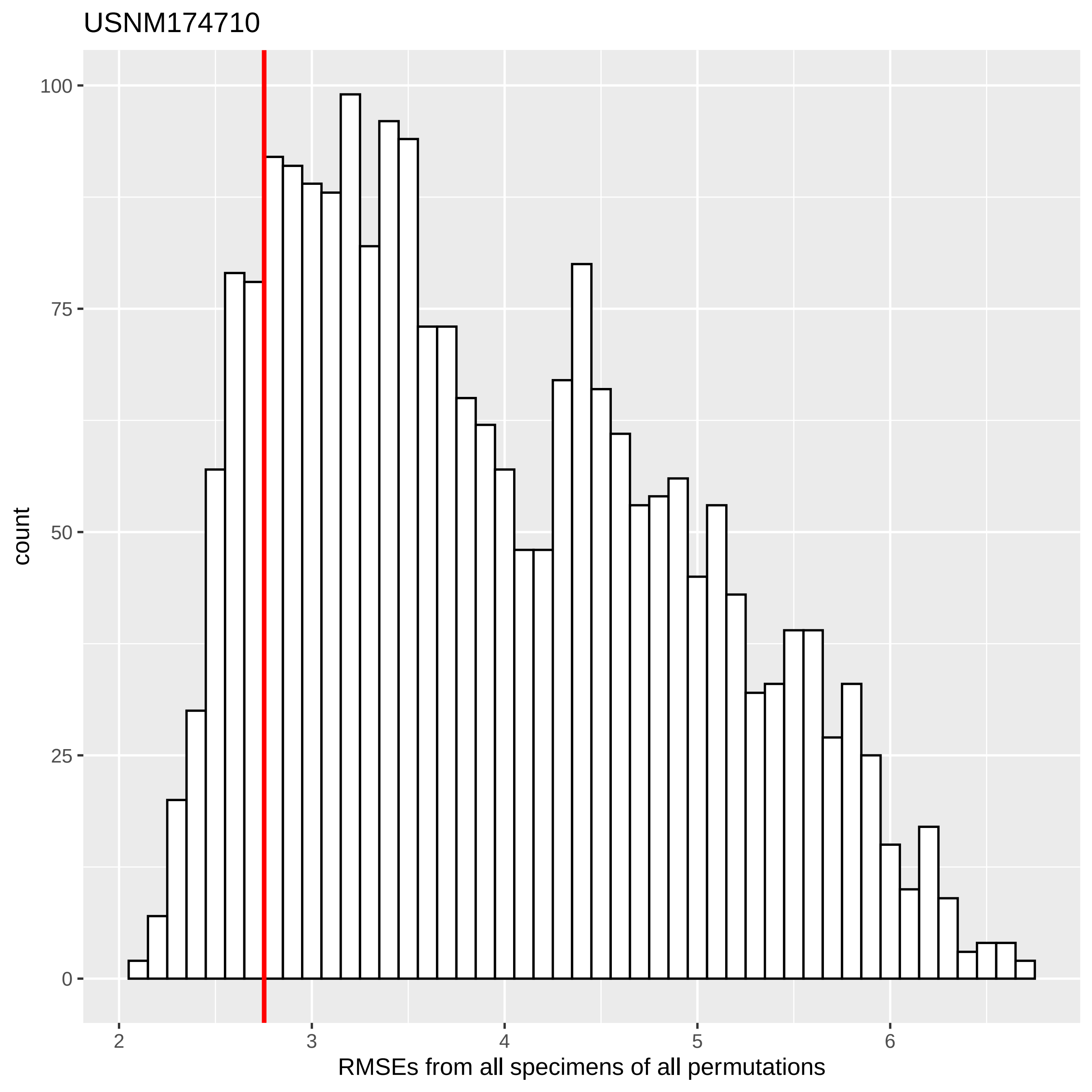

USNM174710
count
100
75
50
25
0
2
3
4
5
6
RMSEs from all specimens of all permutations

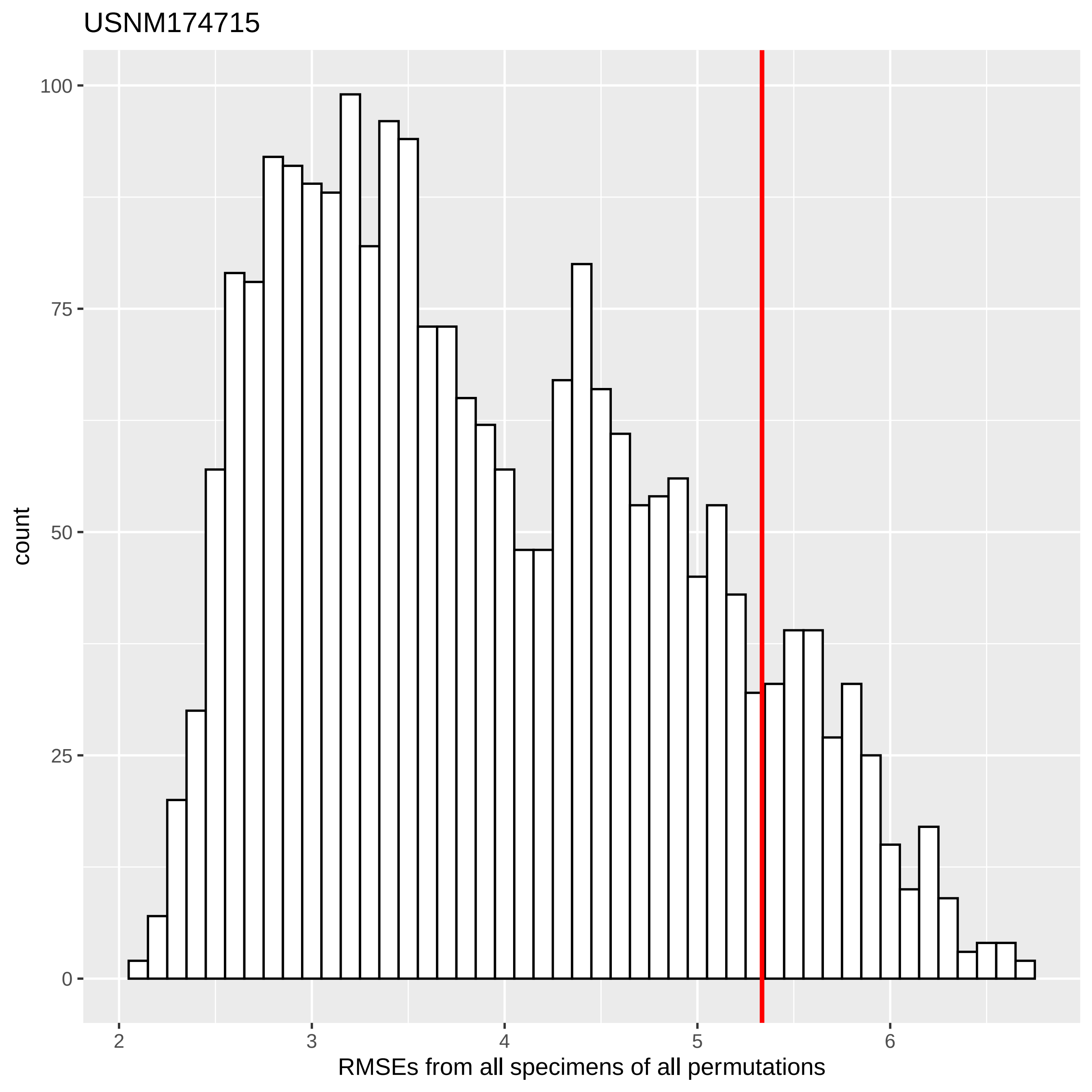

USNM174715
count
100
75
50
25
0
2
3
4
5
6
RMSEs from all specimens of all permutations

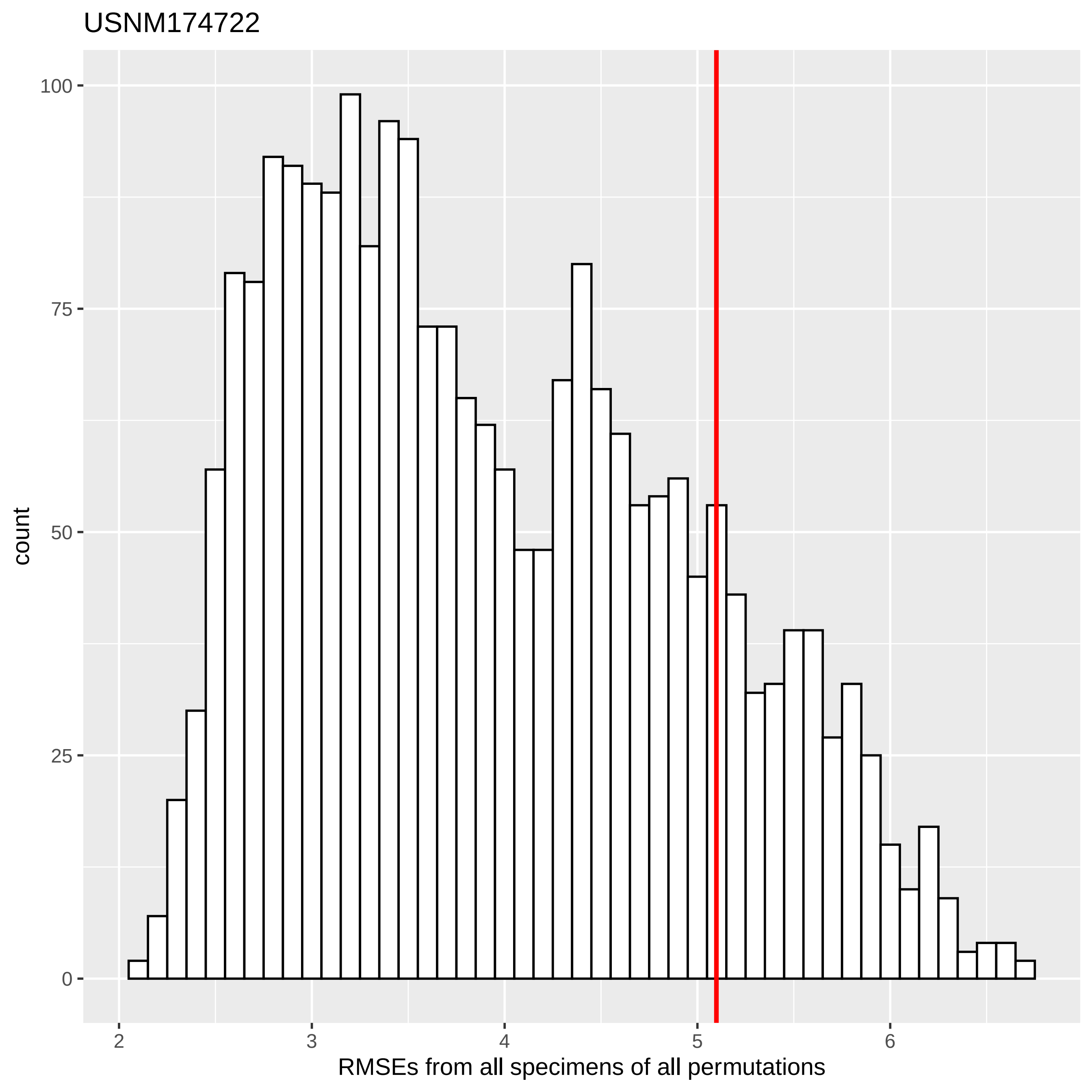

USNM174722
count
100
75
50
25
0
2
3
4
5
6
RMSEs from all specimens of all permutations

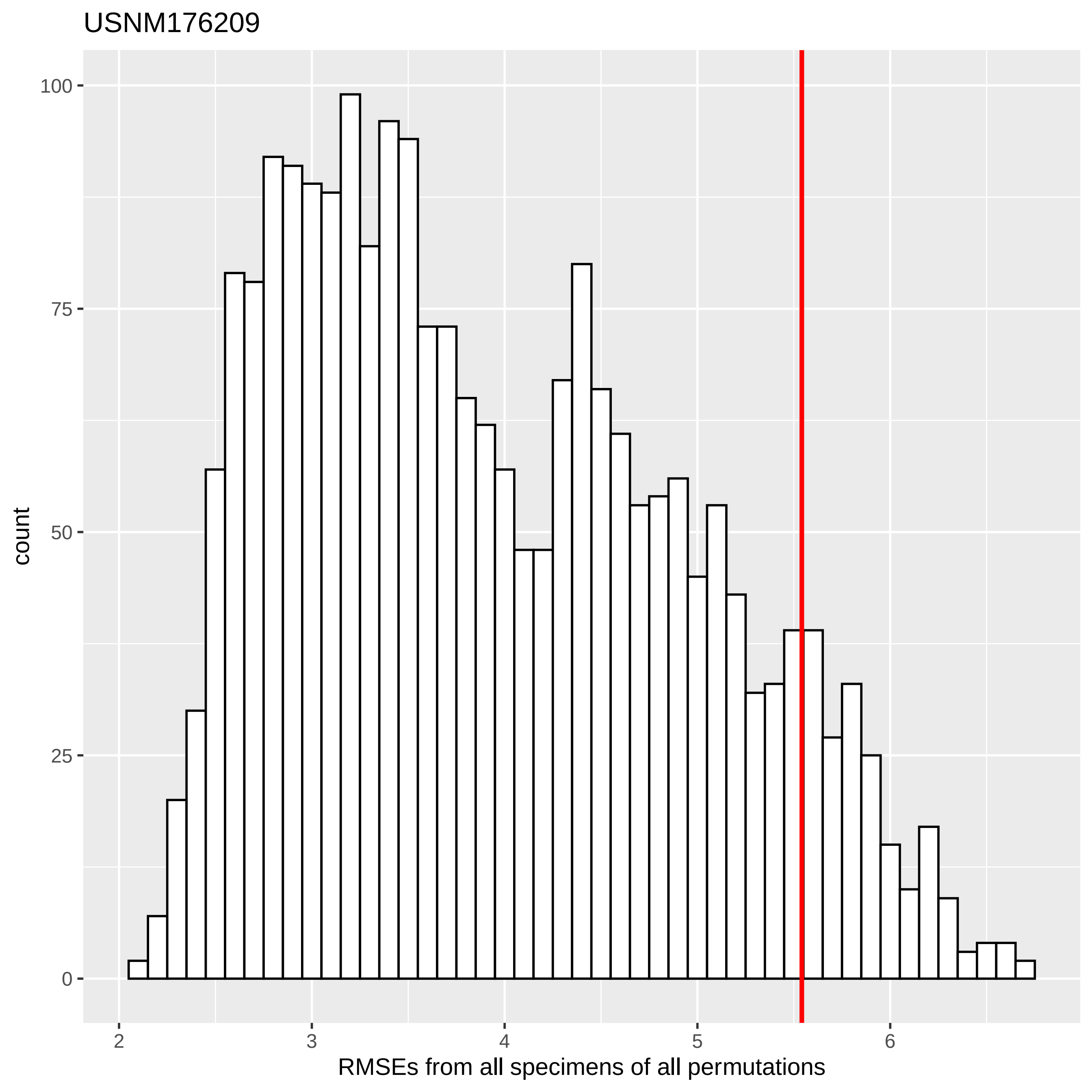

USNM176209
count
100
75
50
25
0
2
3
4
5
6
RMSEs from all specimens of all permutations

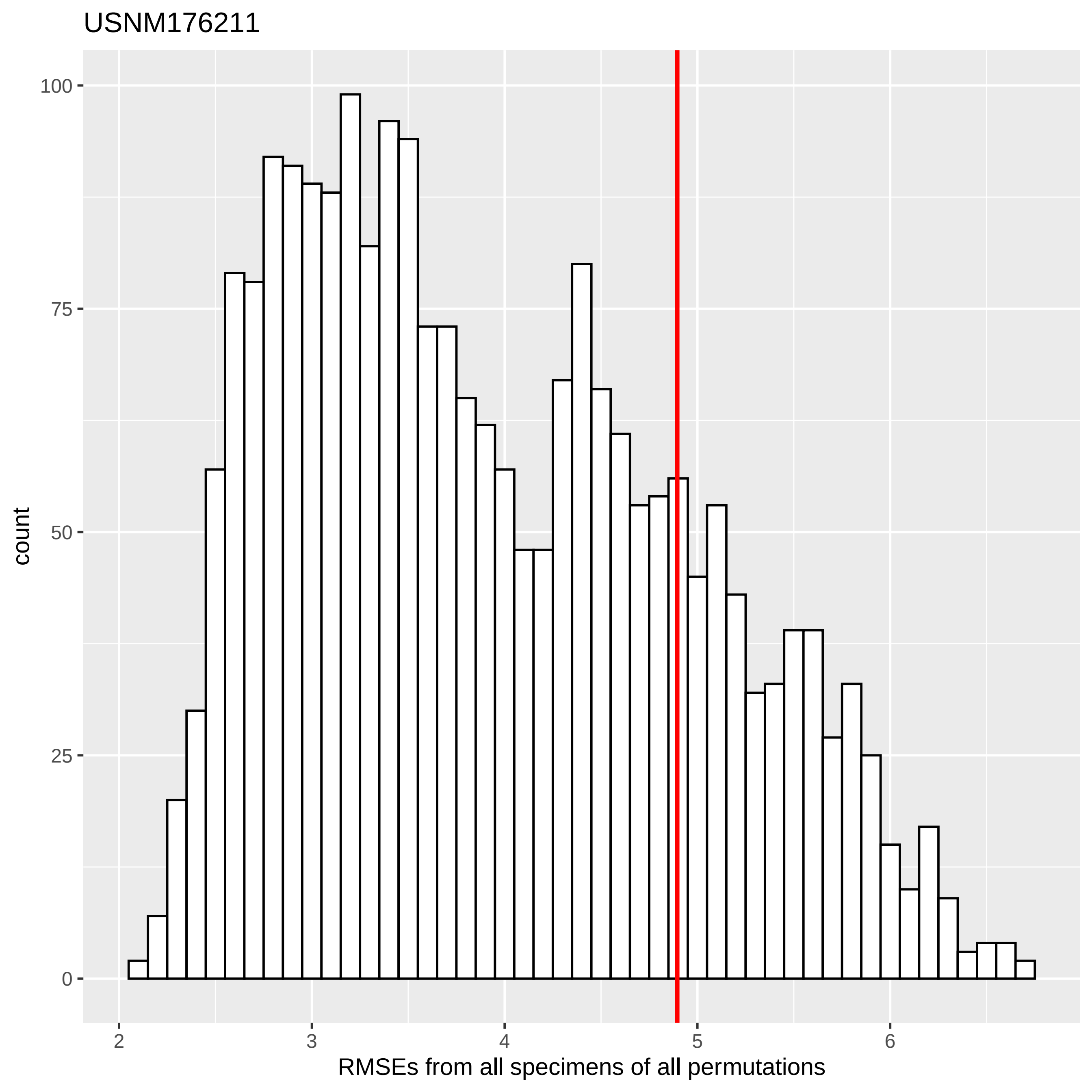
USNM176211
count
RMSEs from all specimens of all permutations
100
75
50
25
0
2
3
4
5
6

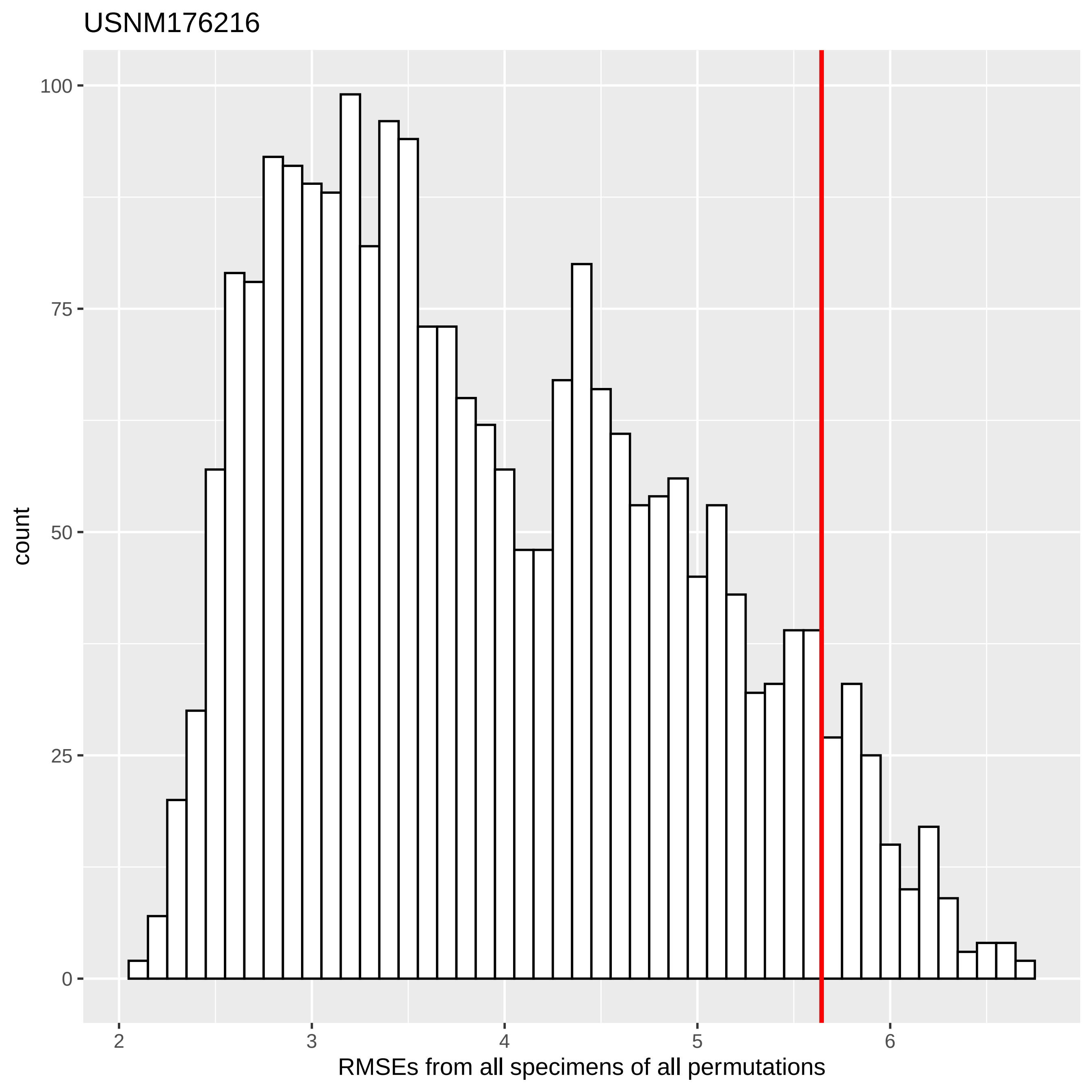
USNM176216
count
100
75
50
25
0
2
3
4
5
6
RMSEs from all specimens of all permutations

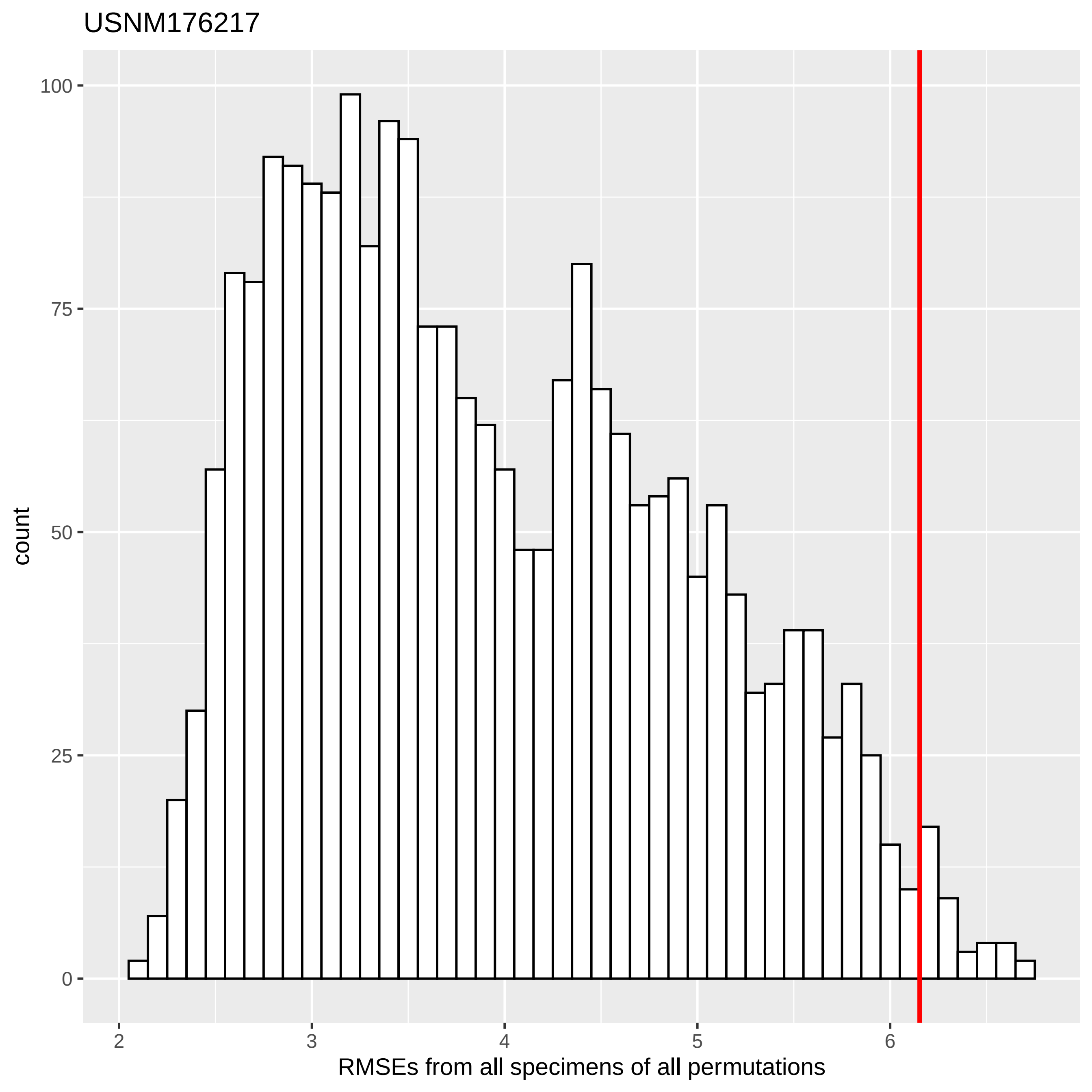

USNM176217
count
100
75
50
25
0
2
3
4
5
6
RMSEs from all specimens of all permutations

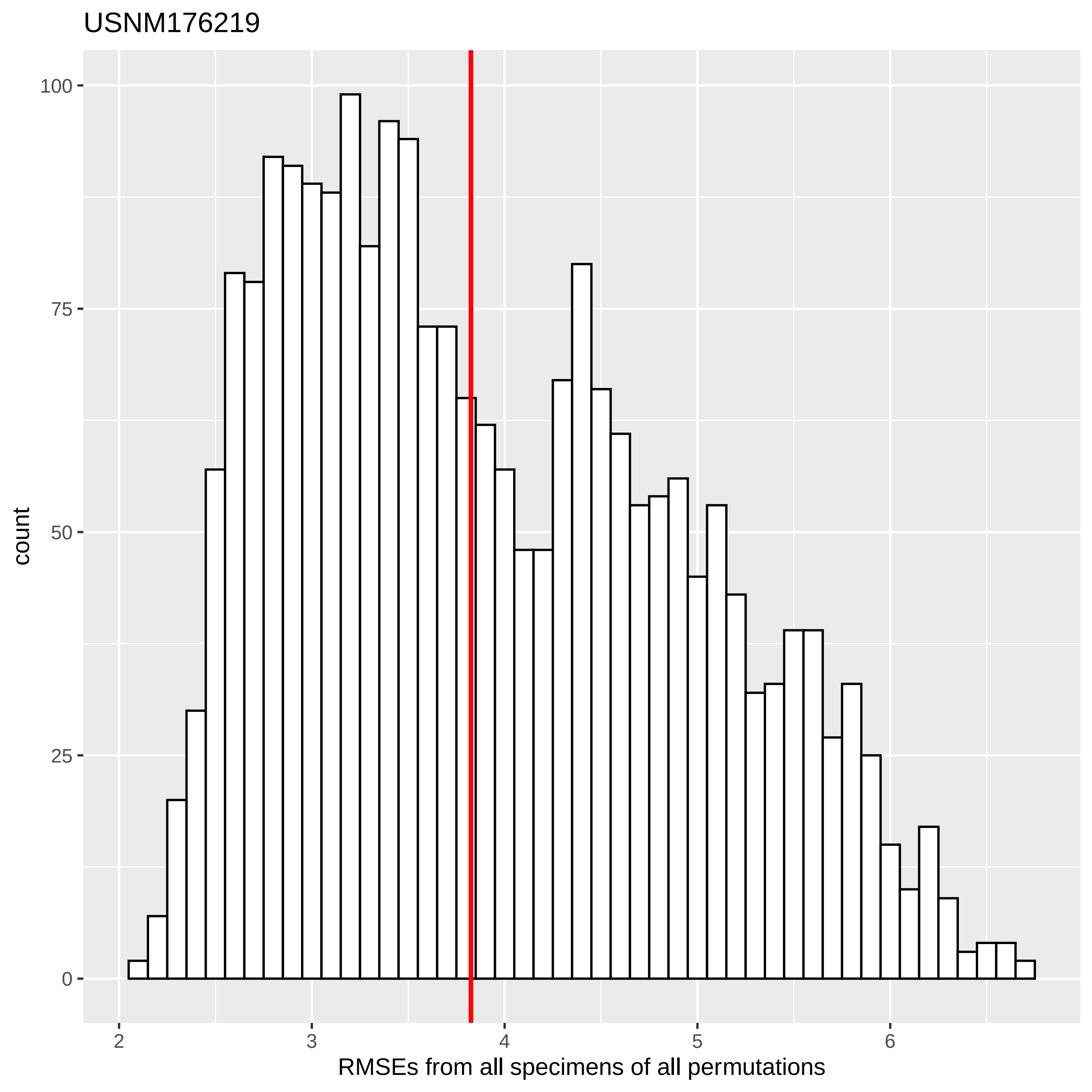

USNM176219
100
75
50
25
0
count
2
3
4
5
6
RMSEs from all specimens of all permutations

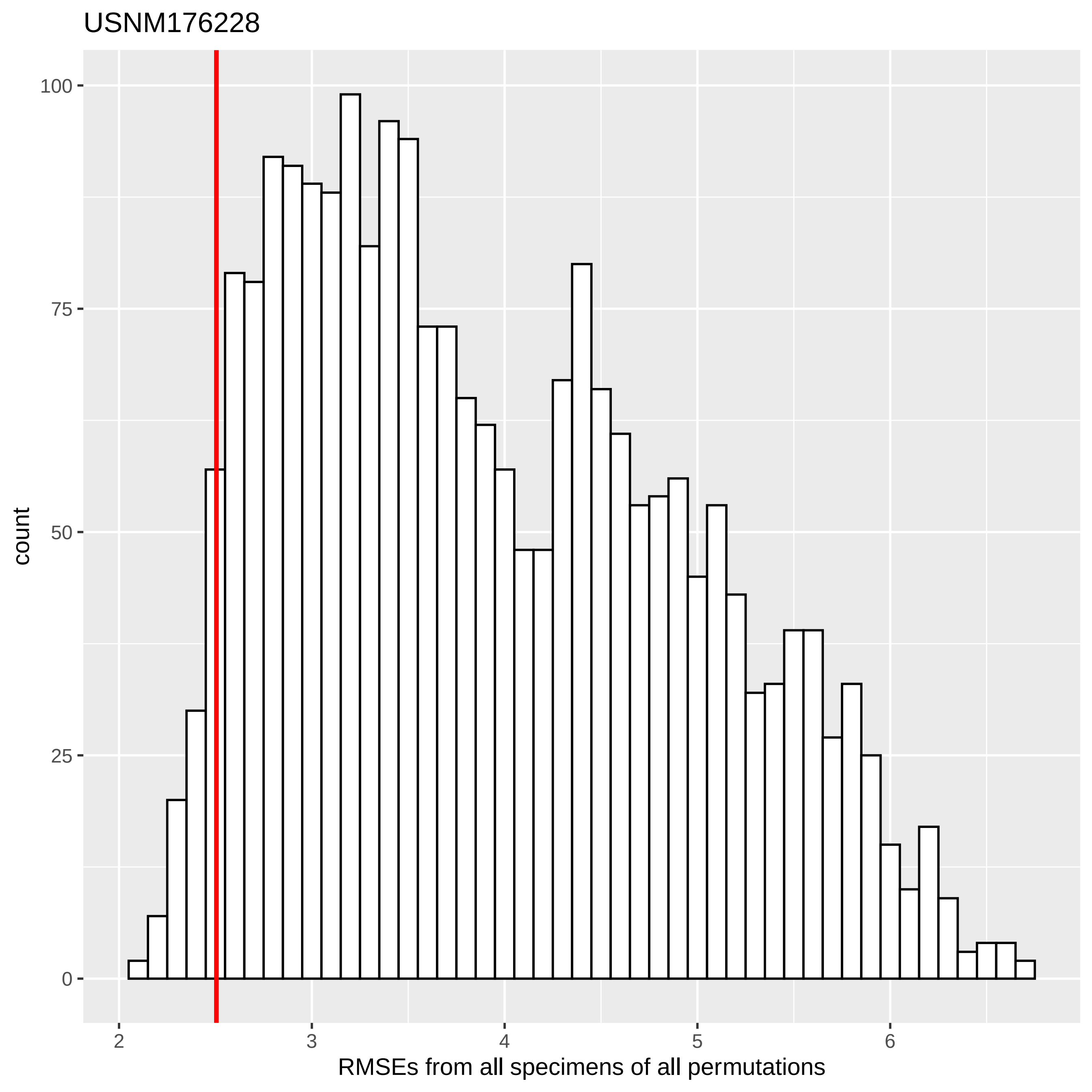
USNM176228
count
100
75
50
25
0
2
3
4
5
6
RMSEs from all specimens of all permutations

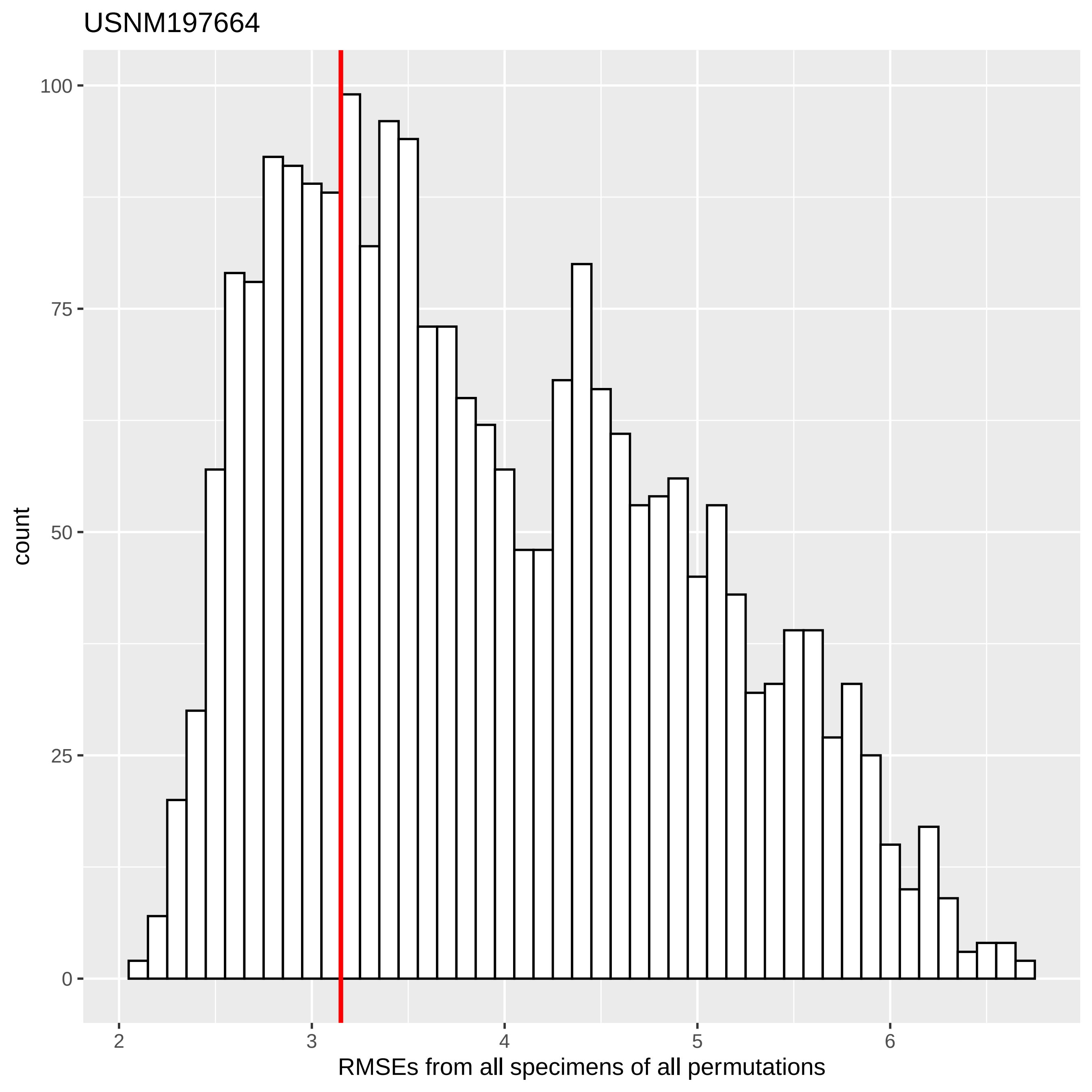

USNM197664
count
100
75
50
25
0
2
3
4
5
6
RMSEs from all specimens of all permutations

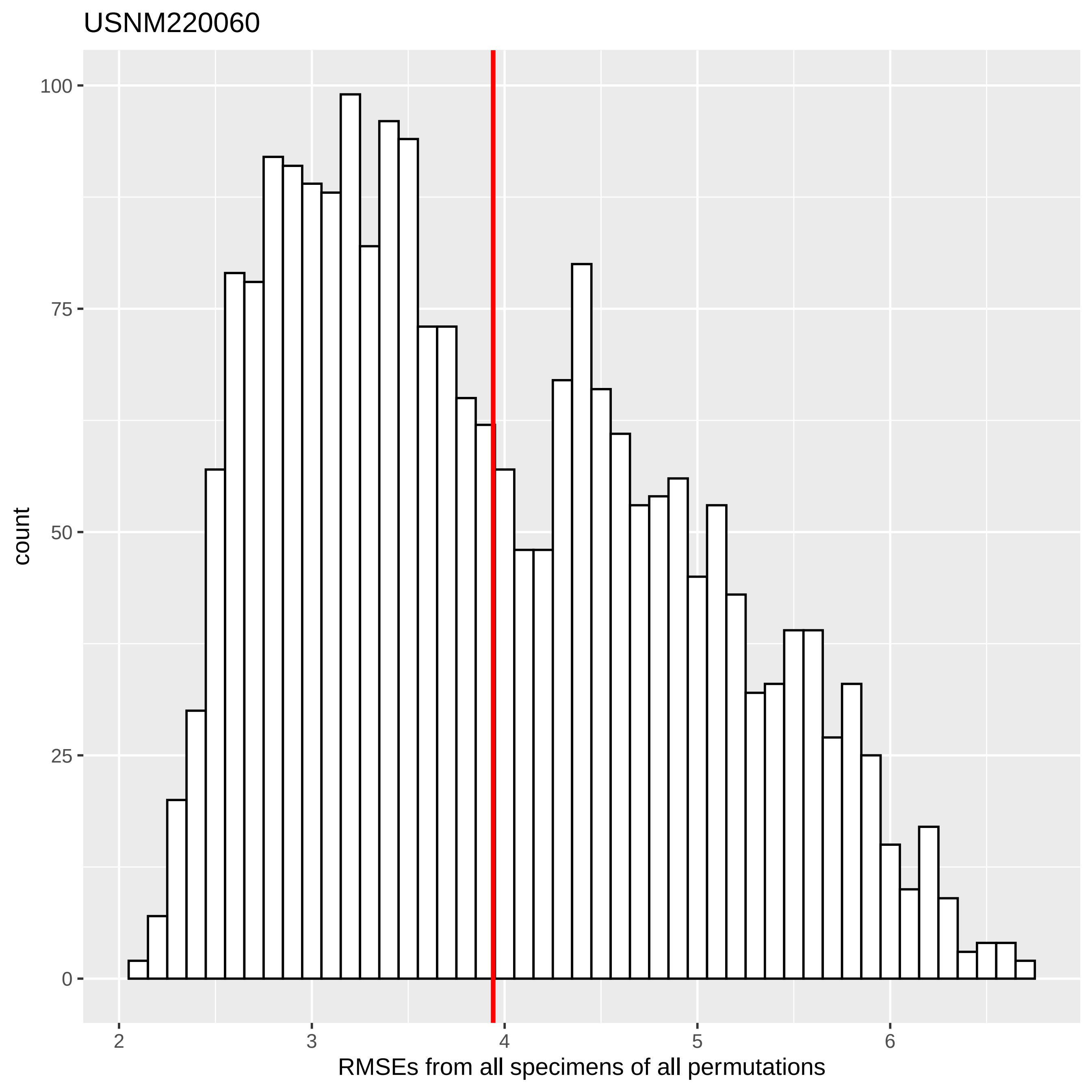

USNM220060
count
100
75
50
25
0
2
3
4
5
6
RMSEs from all specimens of all permutations

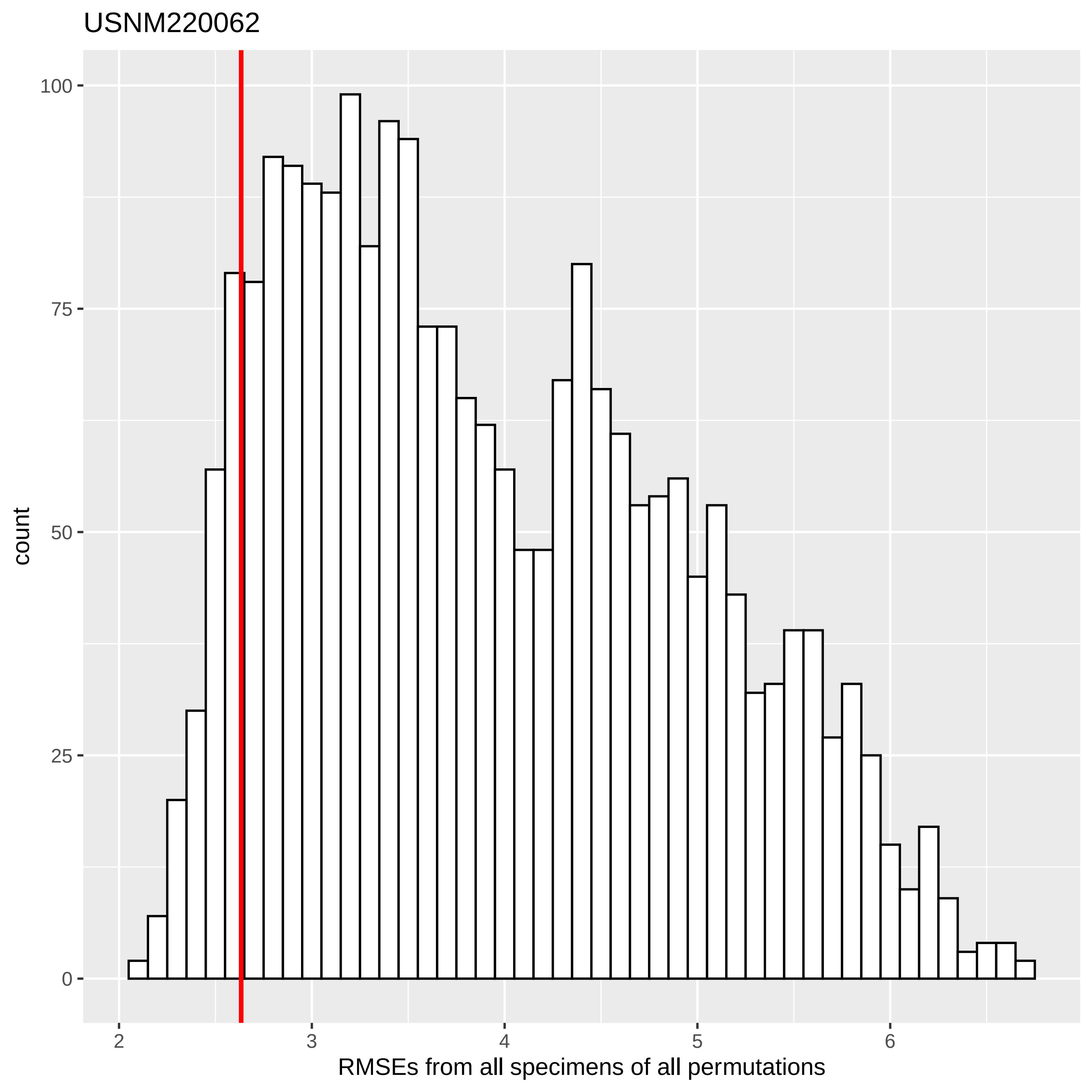
USNM220062
count
100
75
50
25
0
2
3
4
5
6
RMSEs from all specimens of all permutations

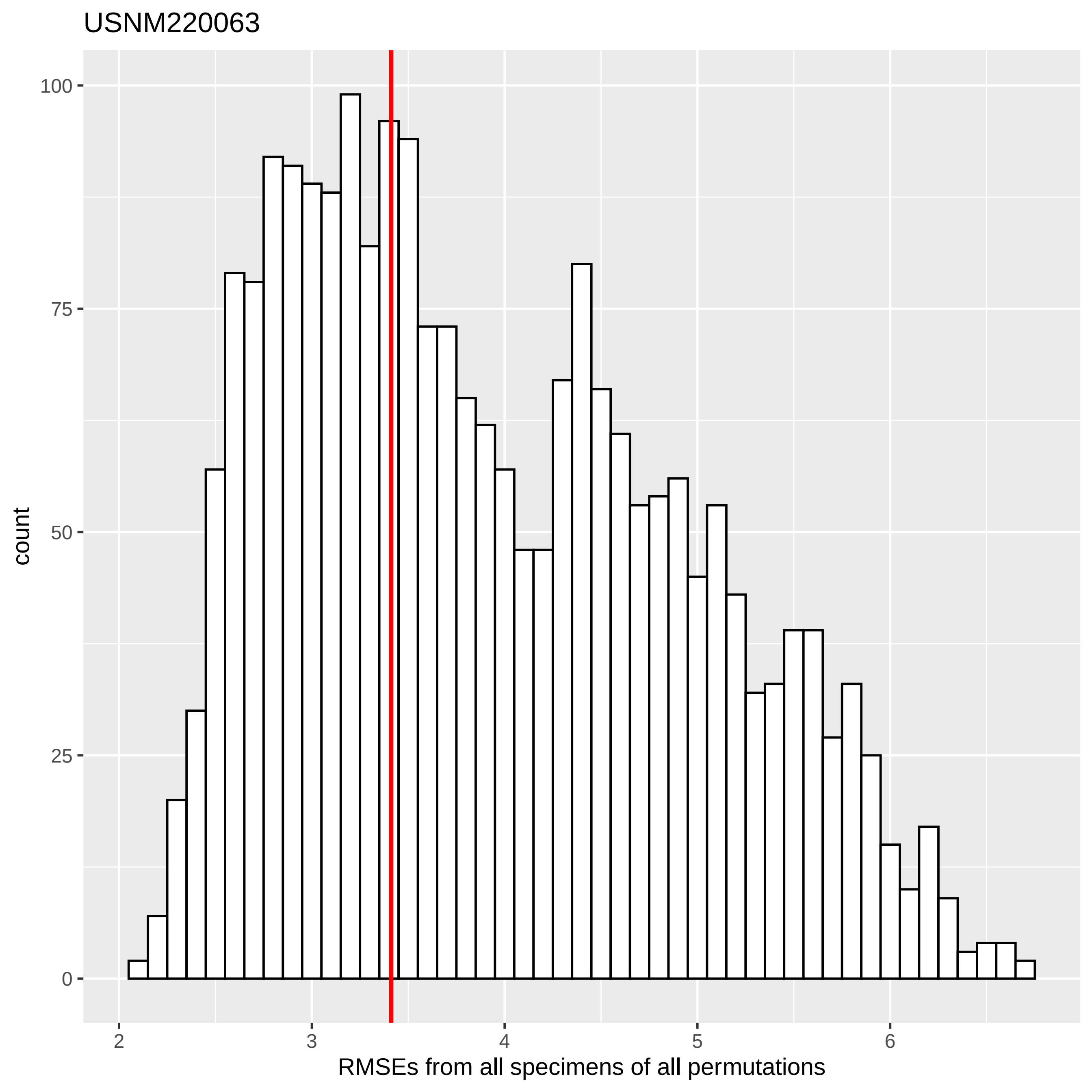
USNM220063
count
100
75
50
25
0
2
3
4
5
6
RMSEs from all specimens of all permutations

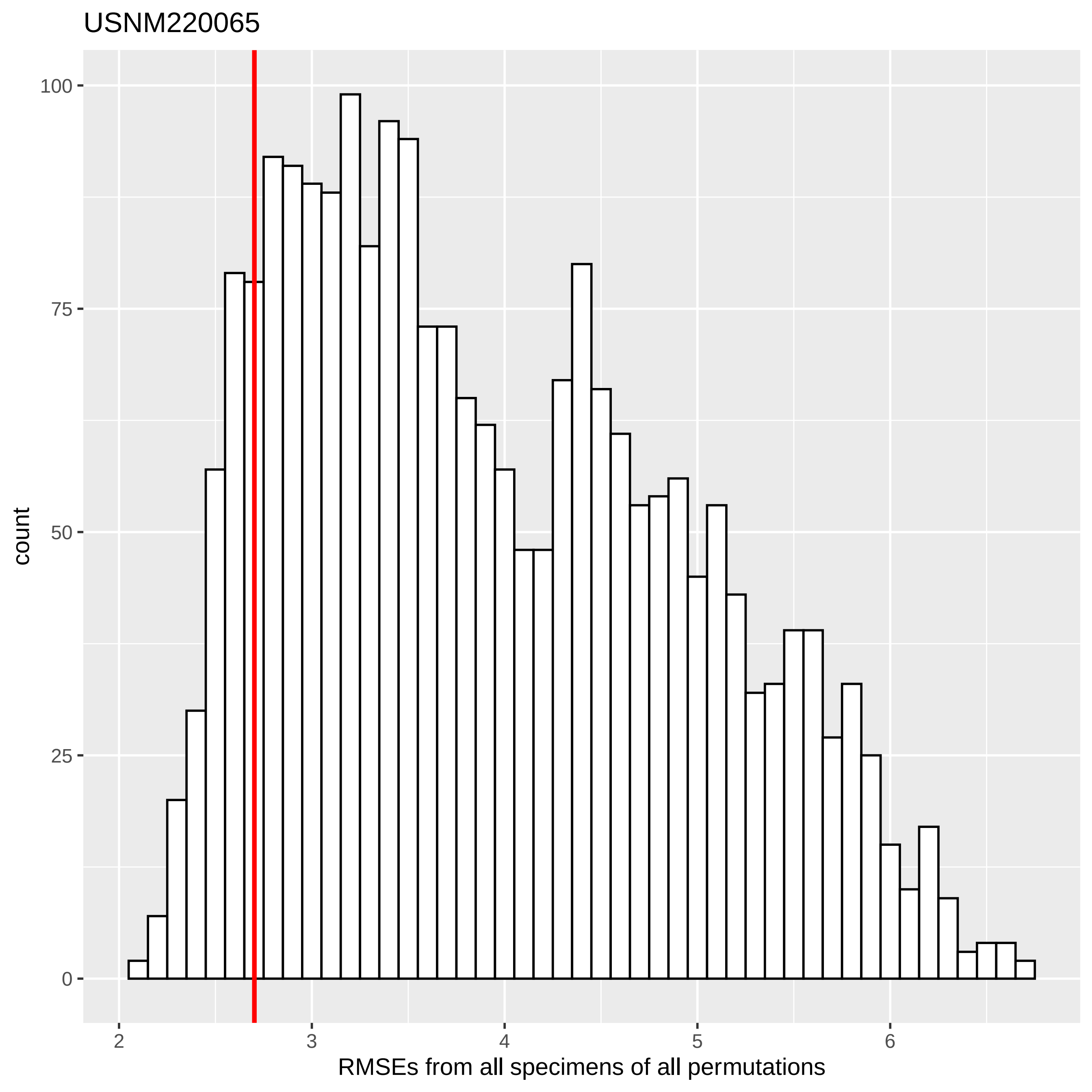

USNM220065
count
100
75
50
25
0
2
3
4
5
6
RMSEs from all specimens of all permutations

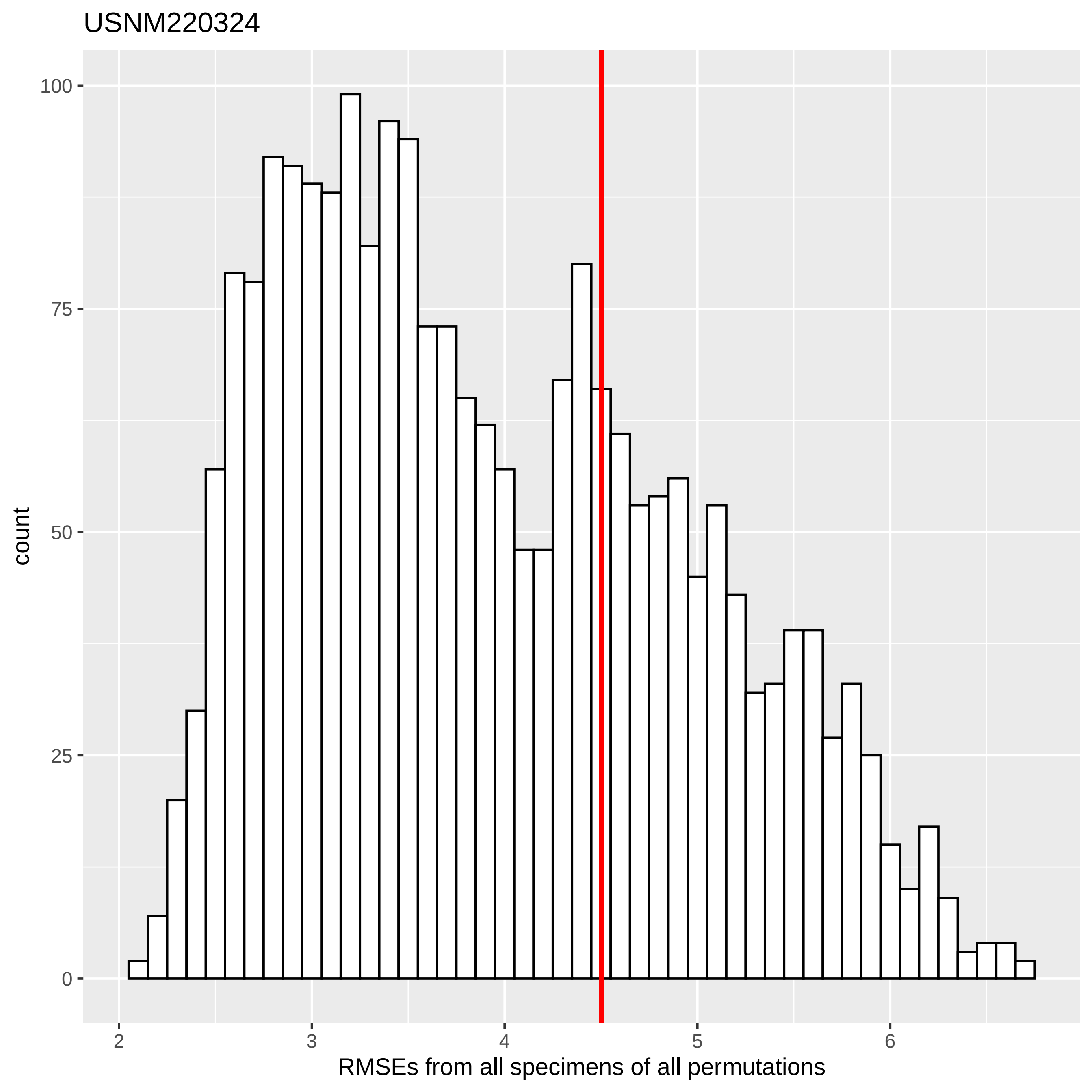
USNM220324
count
RMSEs from all specimens of all permutations
100
75
50
25
0
2
3
4
5
6

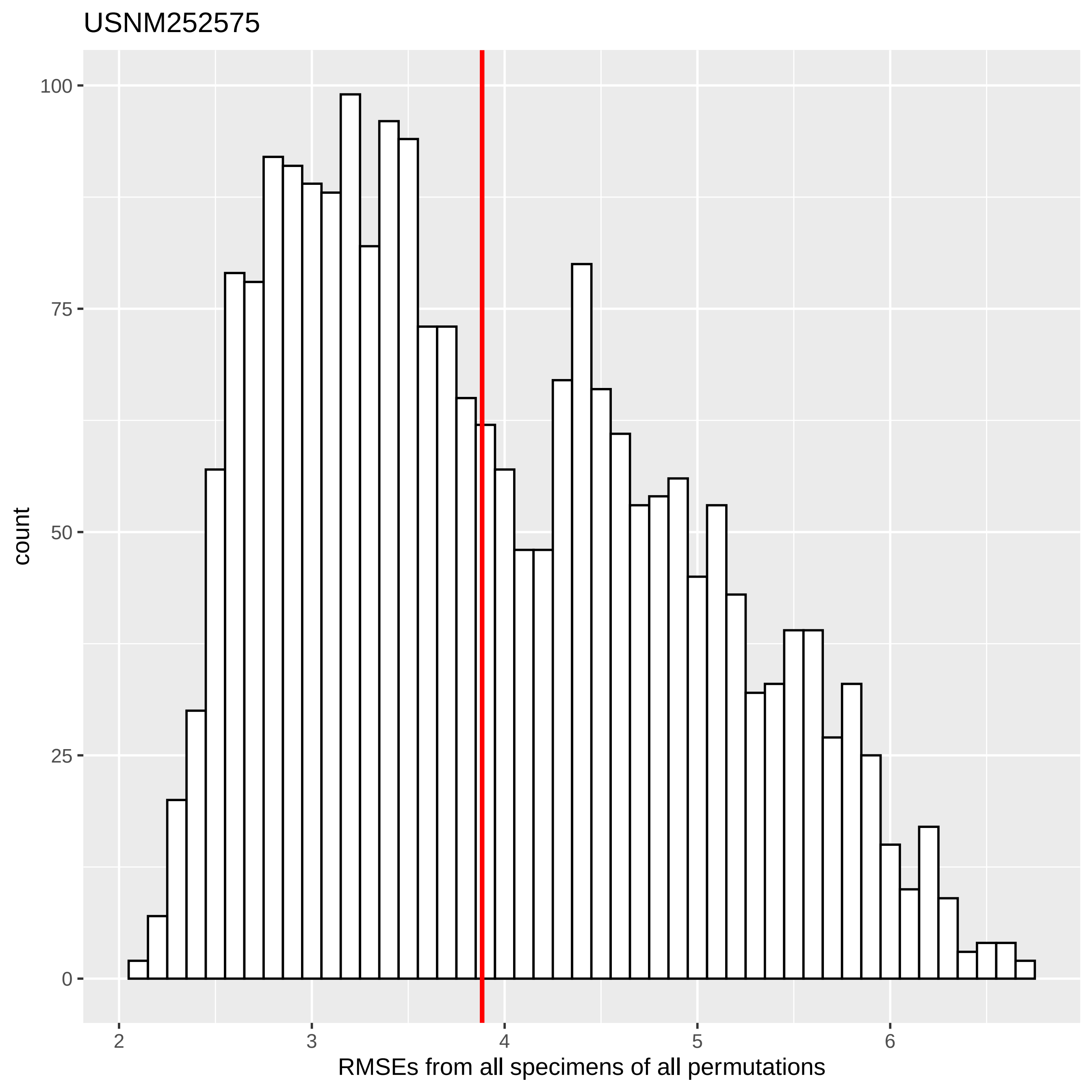

USNM252575
count
100
75
50
25
0
2
3
4
5
6
RMSEs from all specimens of all permutations

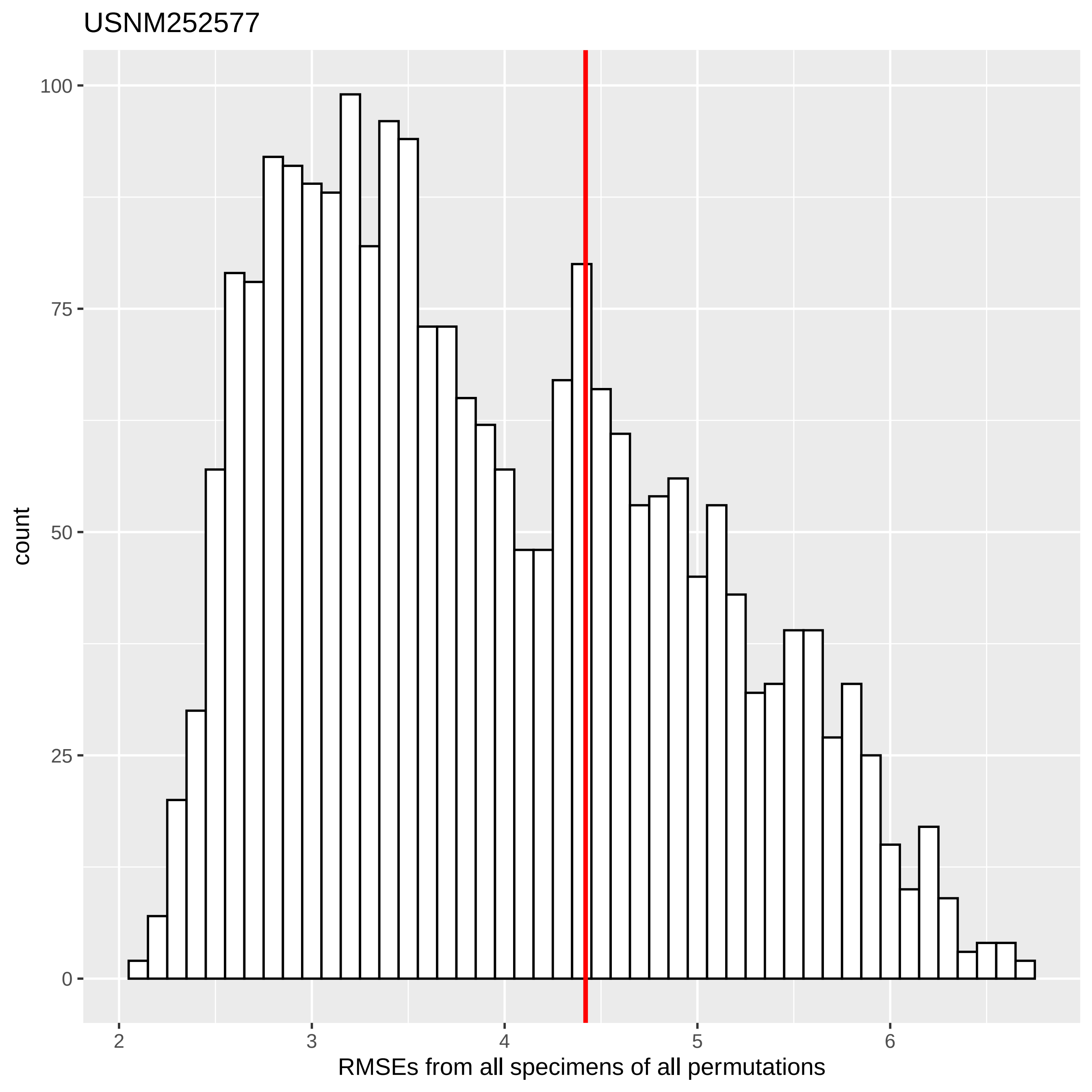
USNM252577
count
100
75
50
25
0
2
3
4
5
6
RMSEs from all specimens of all permutations

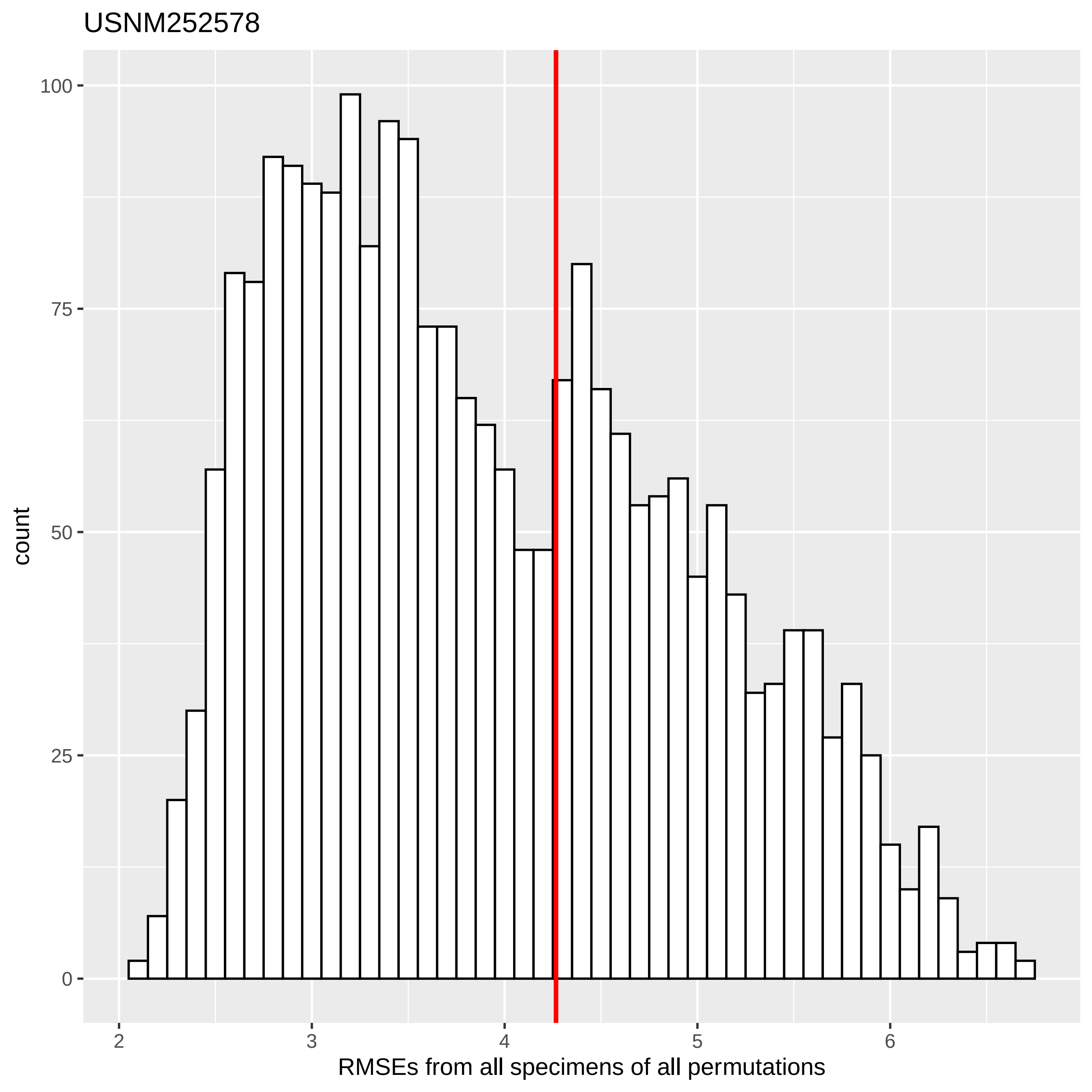

USNM252578
count
100
75
50
25
0
2
3
4
5
6
RMSEs from all specimens of all permutations

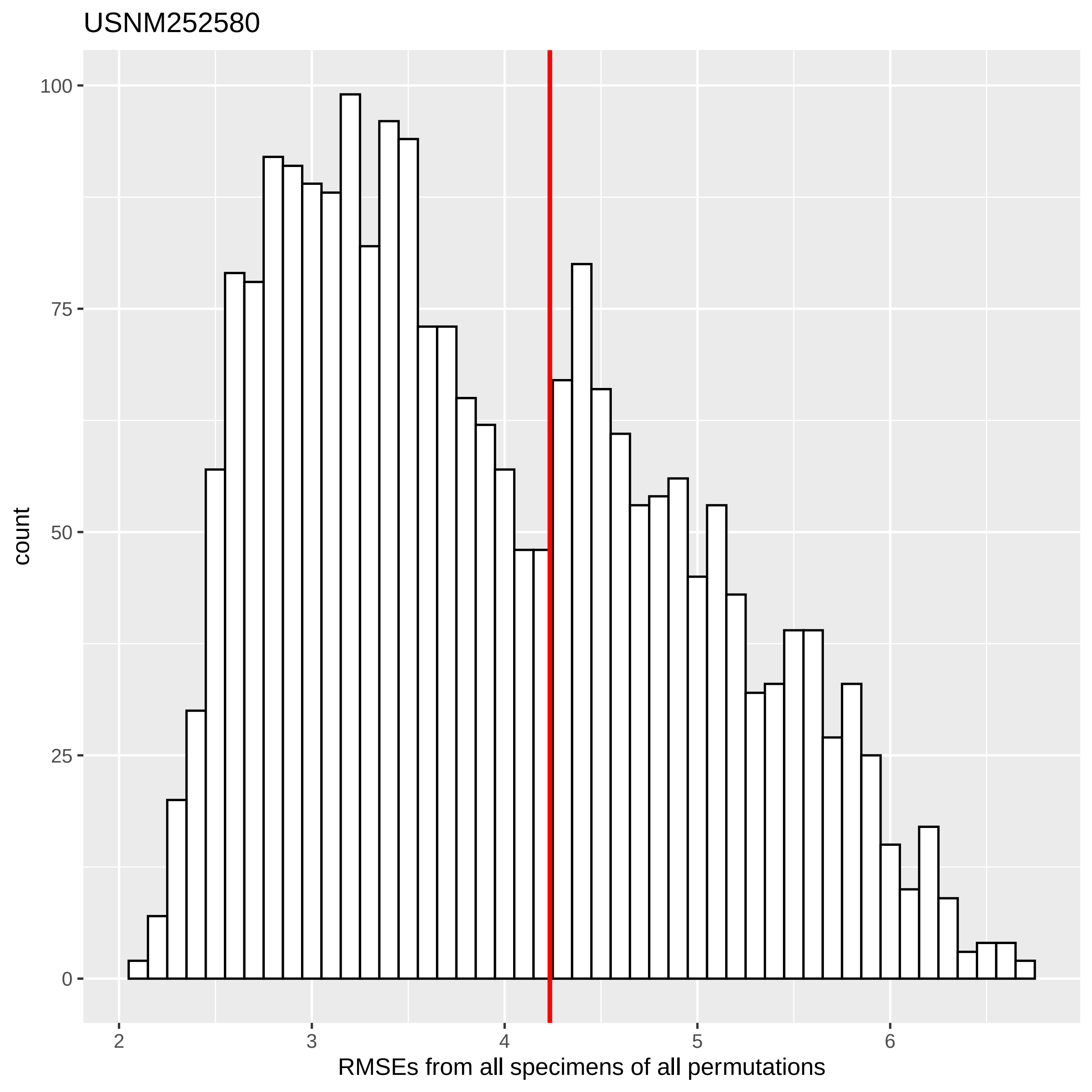

USNM252580
count
100
75
50
25
0
2
3
4
5
6
RMSEs from all specimens of all permutations

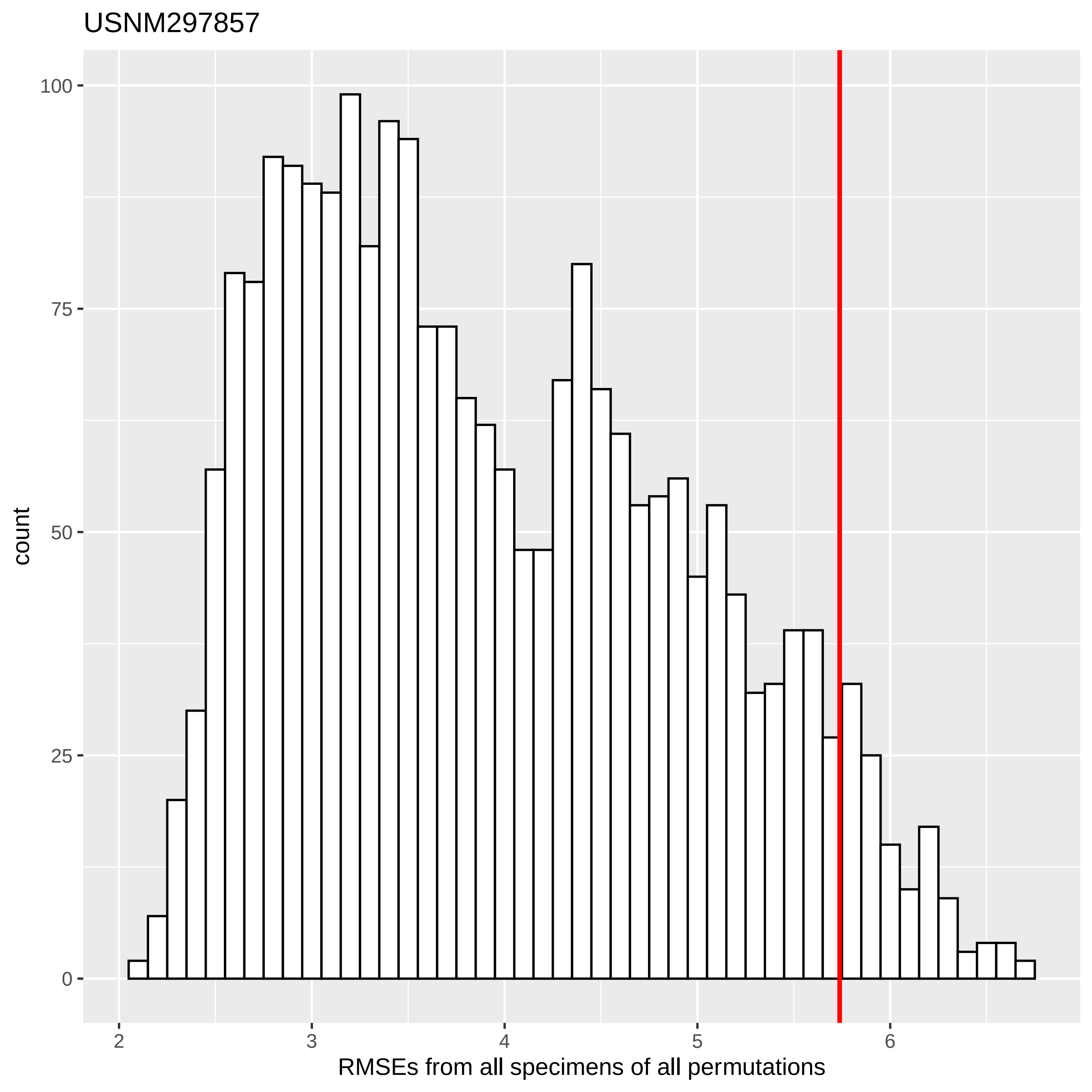

USNM297857
count
100
75
50
25
0
2
3
4
5
6
RMSEs from all specimens of all permutations

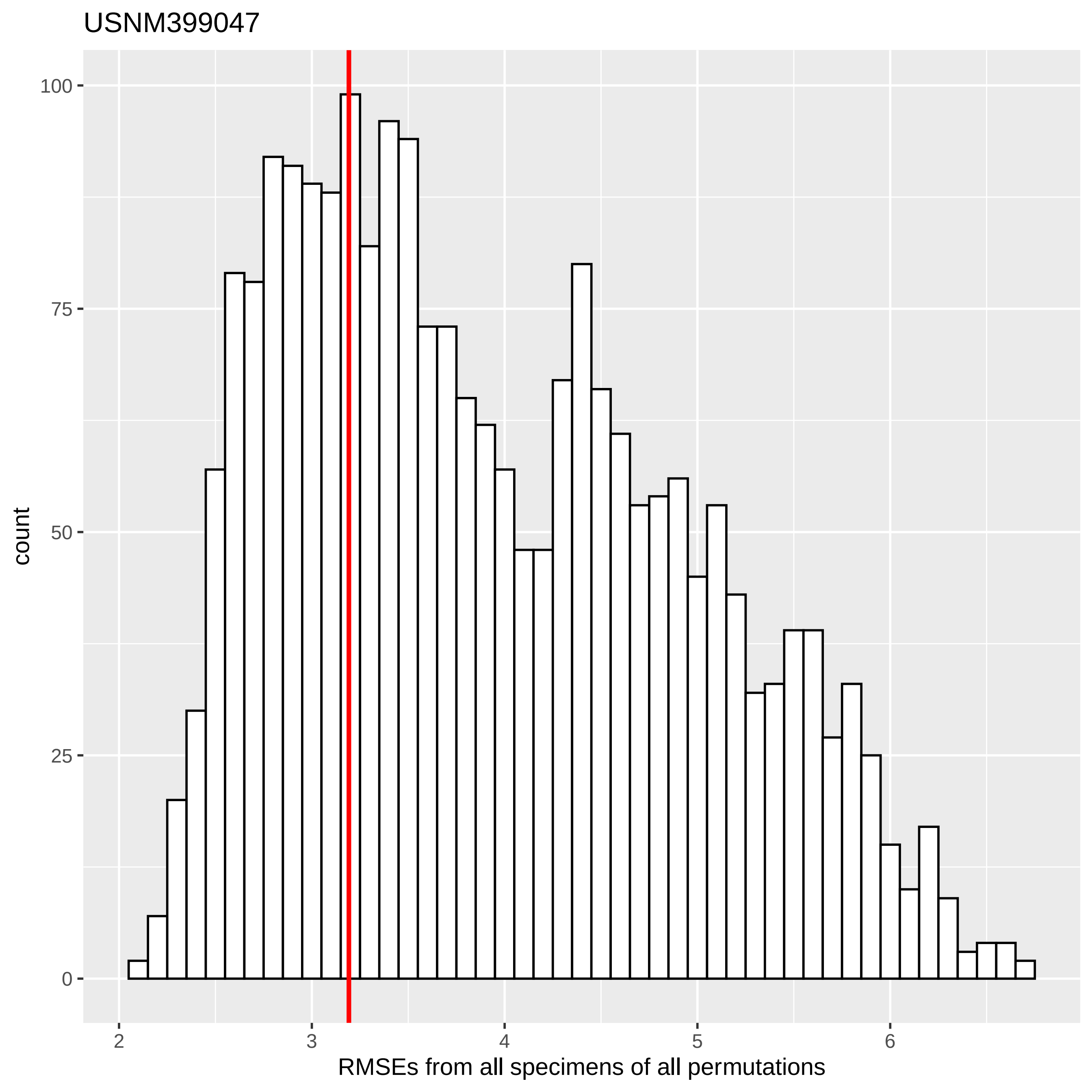
USNM399047
count
100
75
50
25
0
2
3
4
5
6
RMSEs from all specimens of all permutations

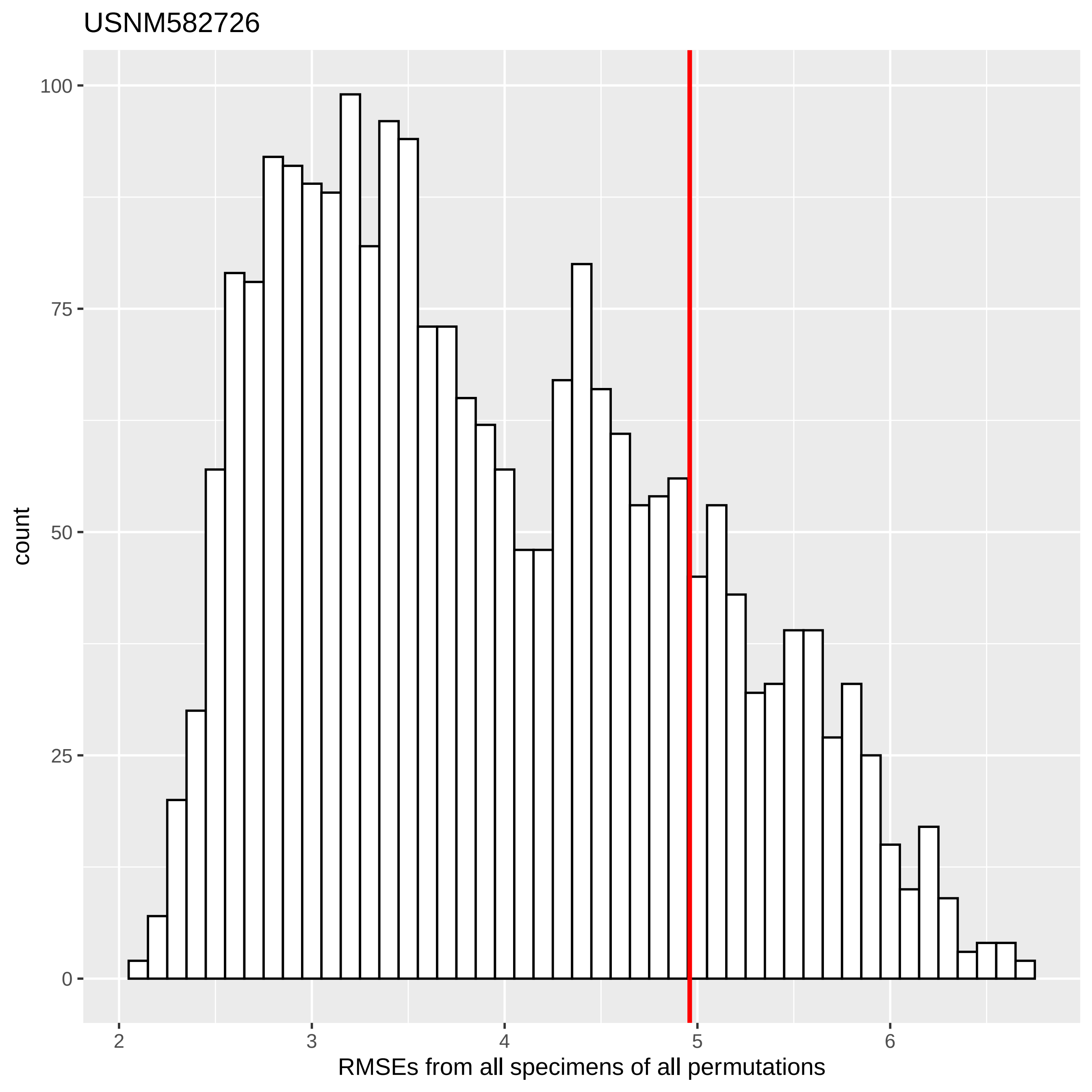
USNM582726
count
100
75
50
25
0
2
3
4
5
6
RMSEs from all specimens of all permutations

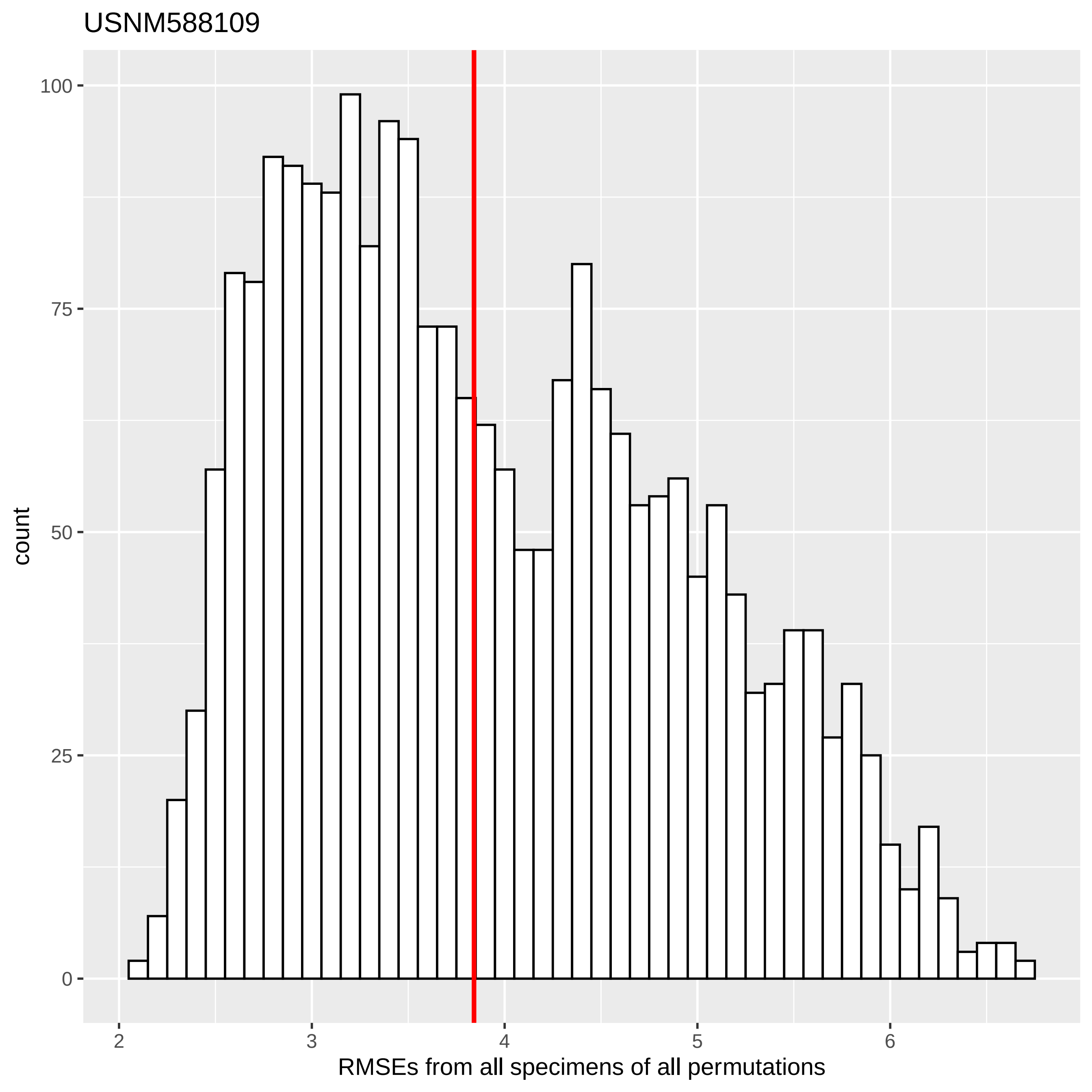
USNM588109
count
100
75
50
25
0
2
3
4
5
6
RMSEs from all specimens of all permutations

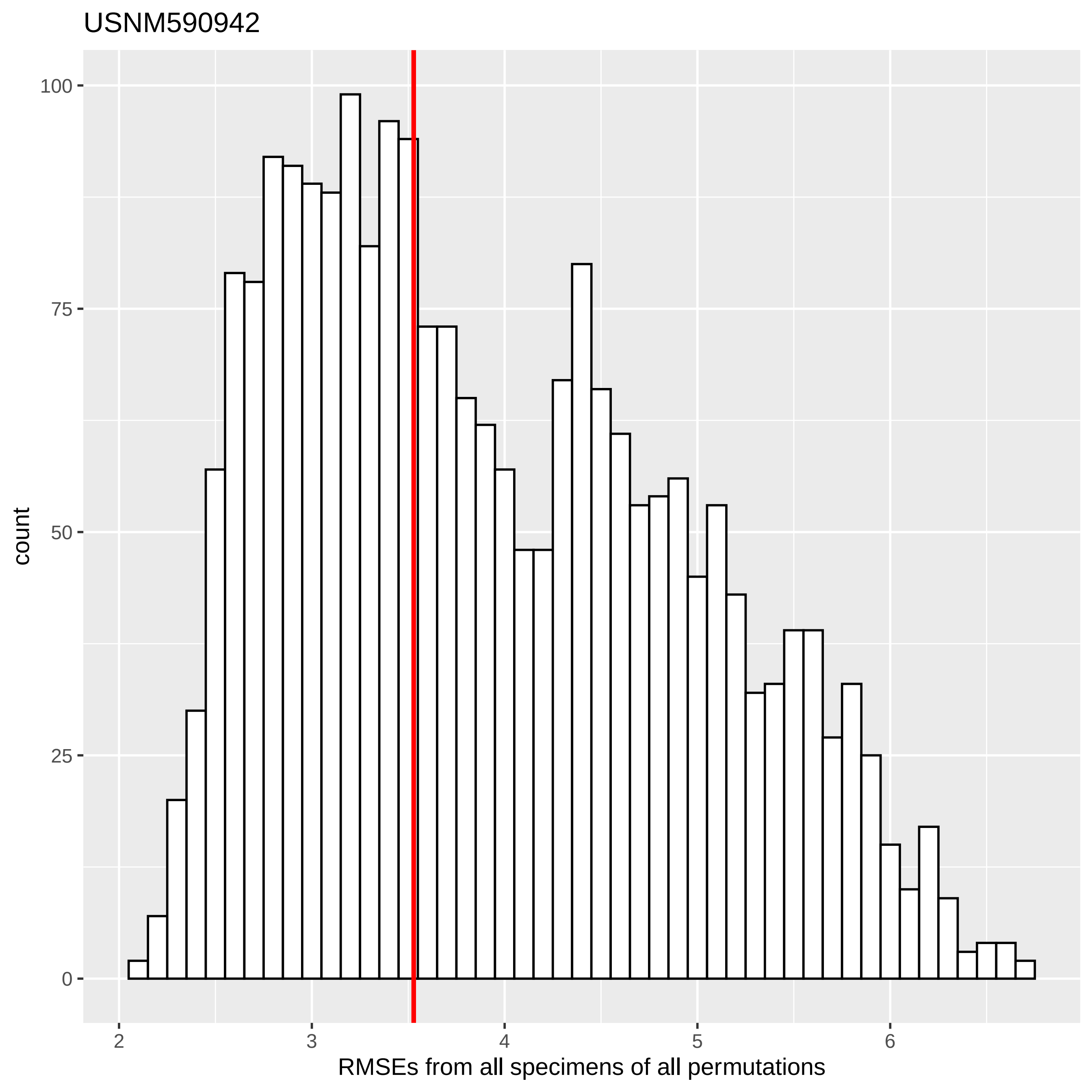

USNM590942
count
100
75
50
25
0
2
3
4
5
6
RMSEs from all specimens of all permutations

USNM590947

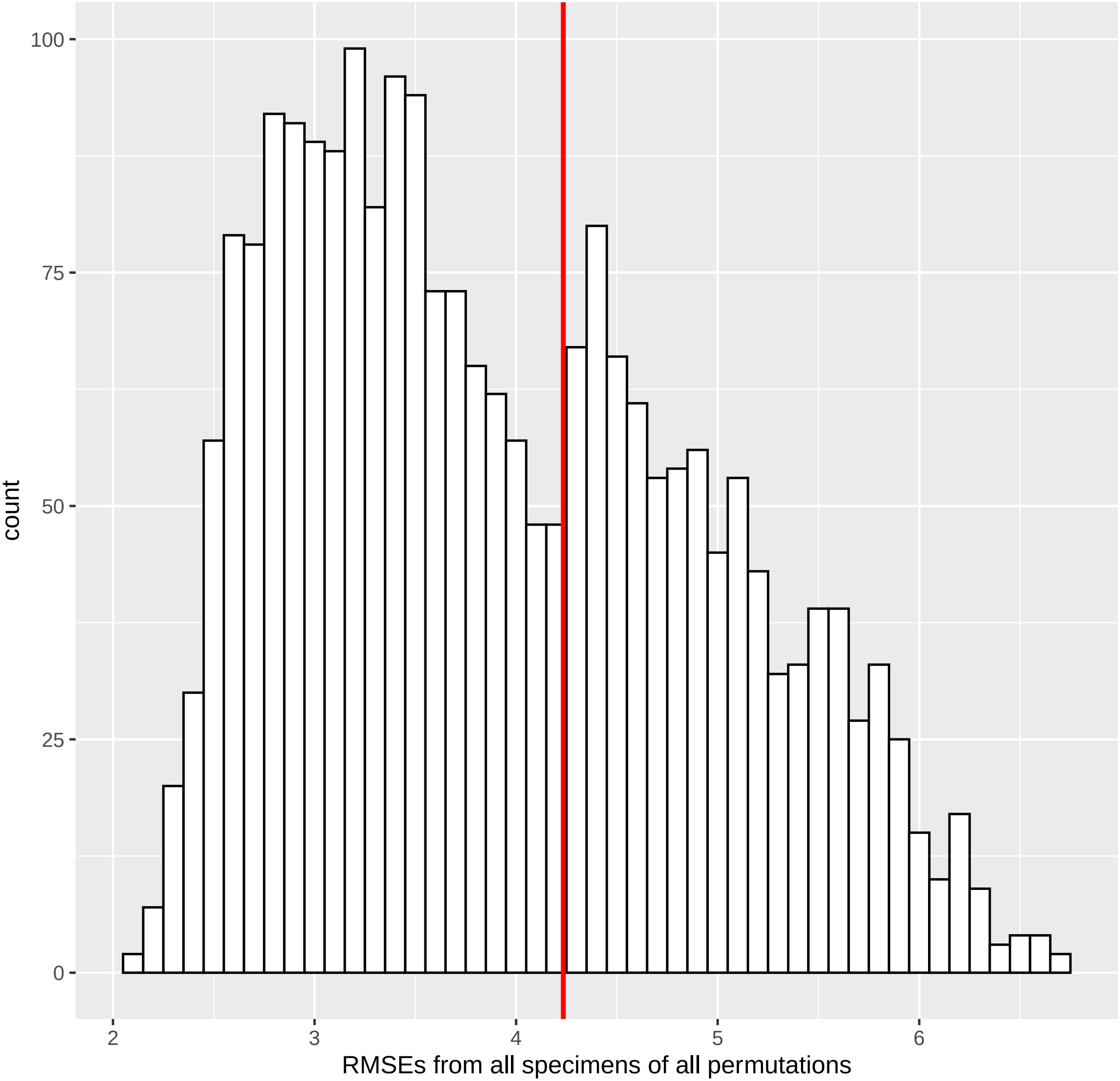

100
75
50
25
0
count
2
3
4
5
6
RMSEs from all specimens of all permutations

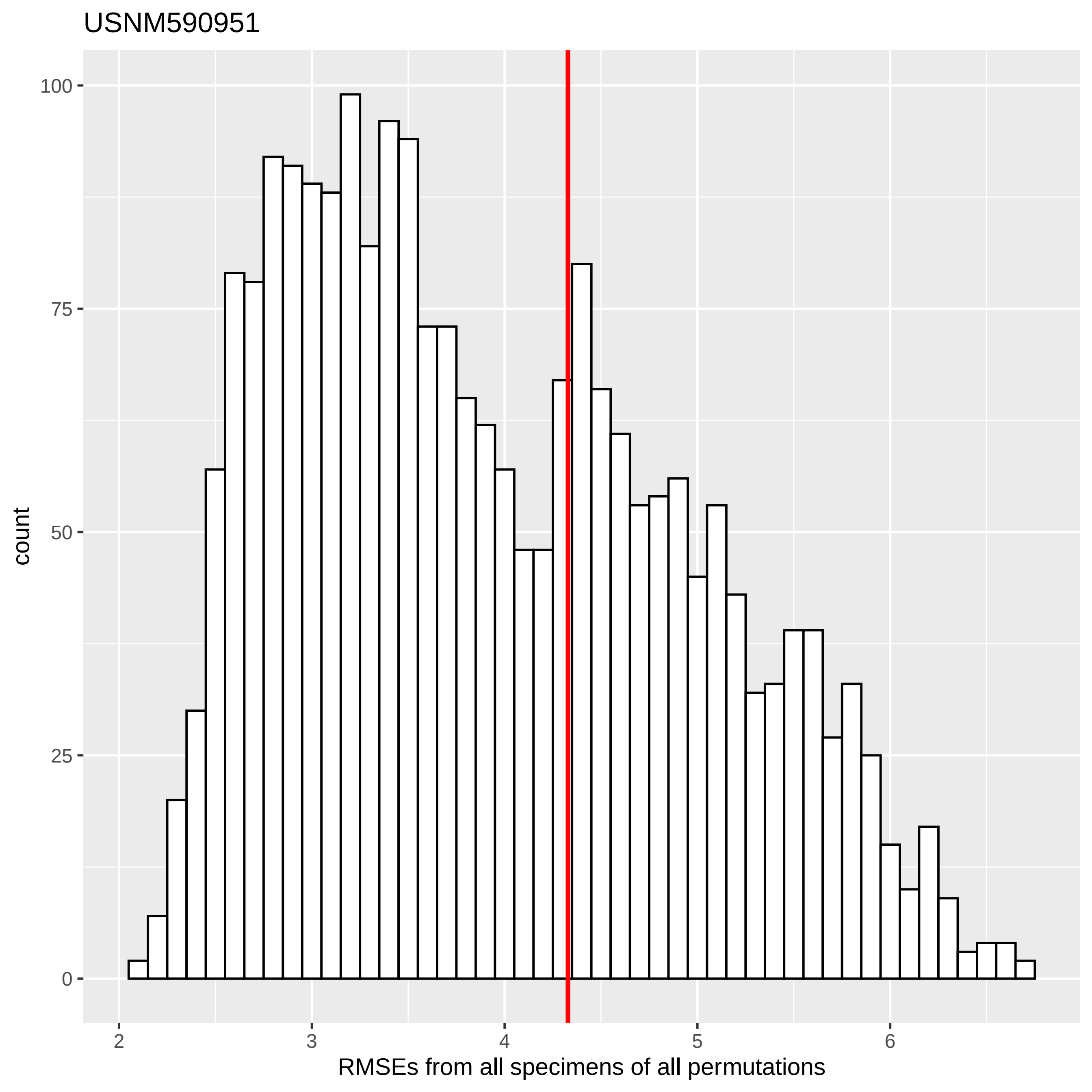
USNM590951
count
100
75
50
25
0
2
3
4
5
6
RMSEs from all specimens of all permutations

USNM590954

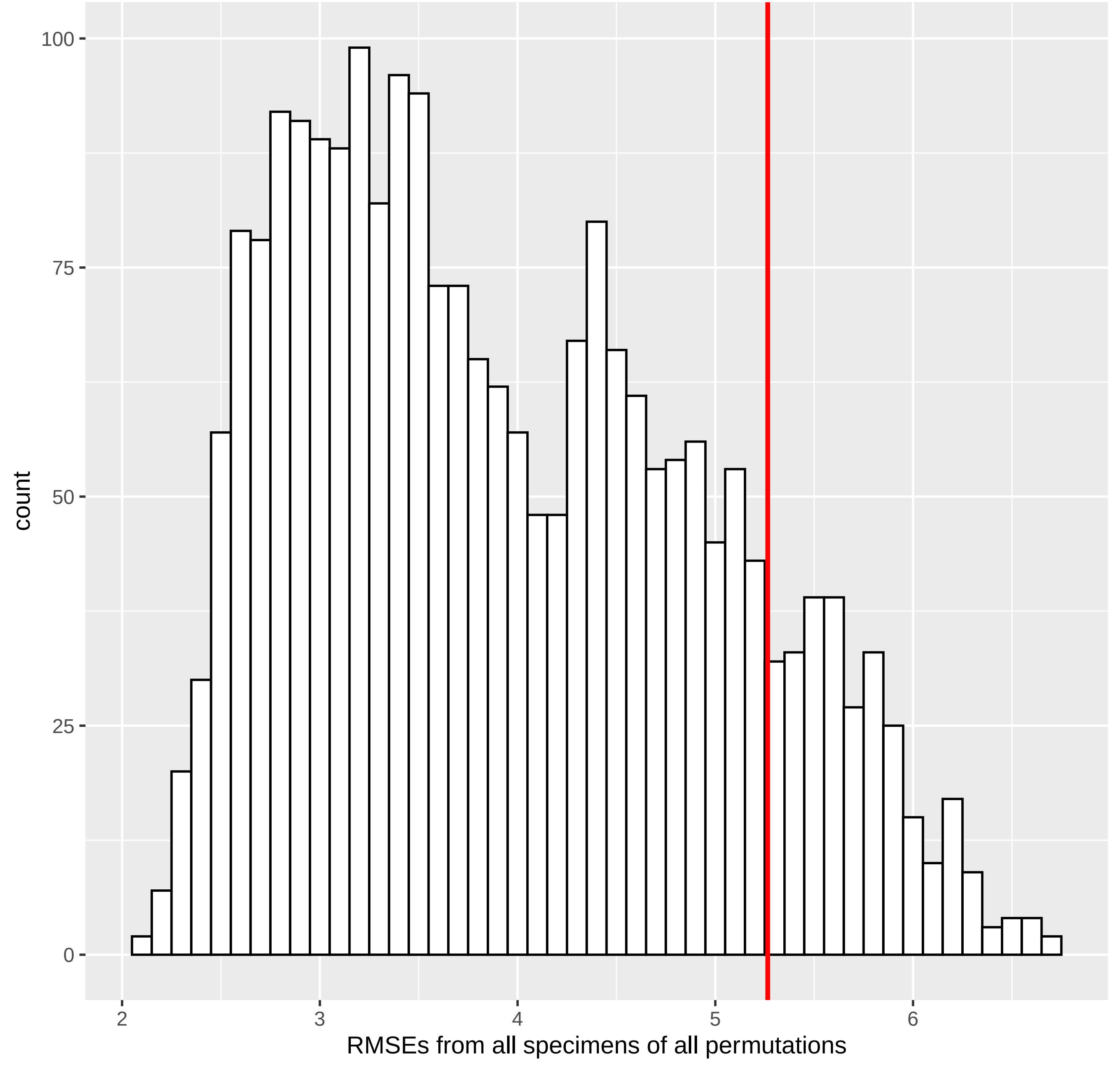

100
75
50
25
0
count
2
3
4
5
6
RMSEs from all specimens of all permutations

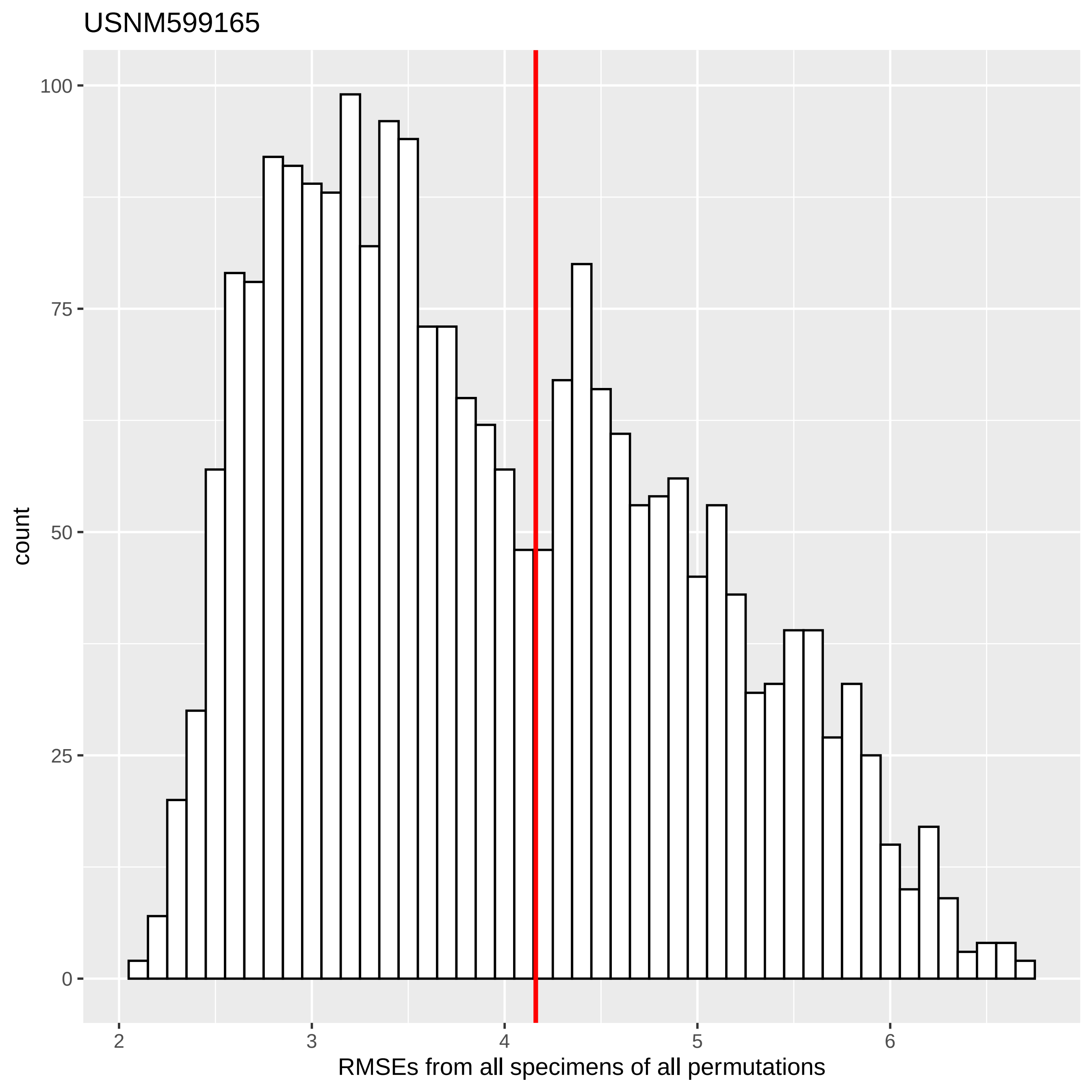

USNM599165
count
100
75
50
25
0
2
3
4
5
6
RMSEs from all specimens of all permutations

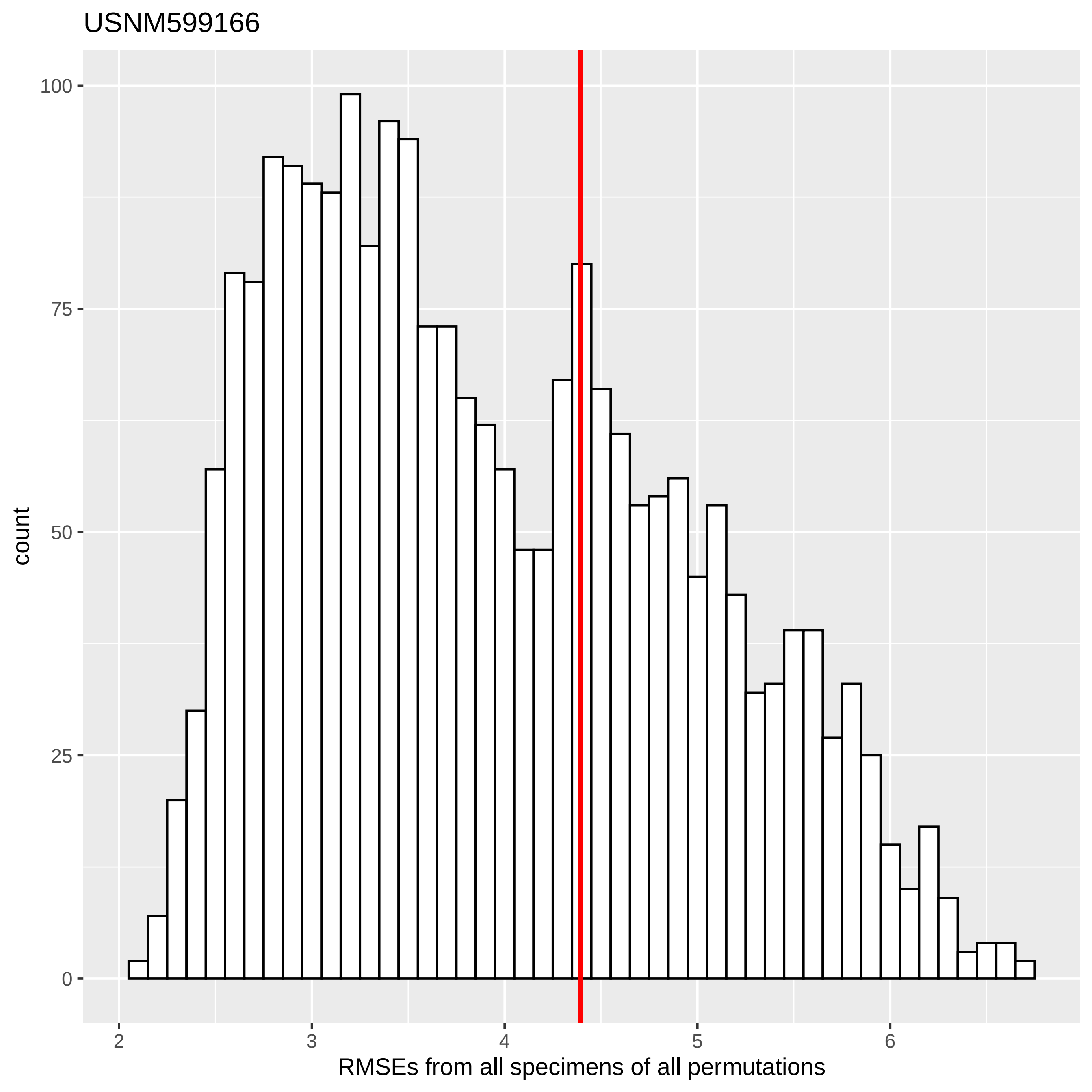
USNM599166
count
100
75
50
25
0
2
3
4
5
6
RMSEs from all specimens of all permutations
