## Supplemental Figure 8 for "Automated Landmarking via Multiple Templates"

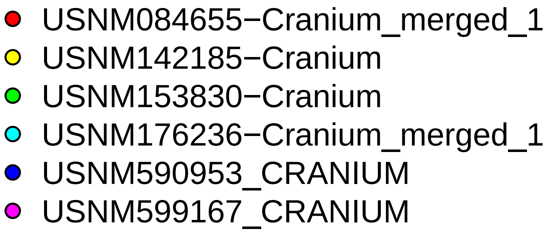
USNM084655−Cranium_merged_1
USNM142185−Cranium
USNM153830−Cranium
USNM176236−Cranium_merged_1
USNM590953_CRANIUM
USNM599167_CRANIUM

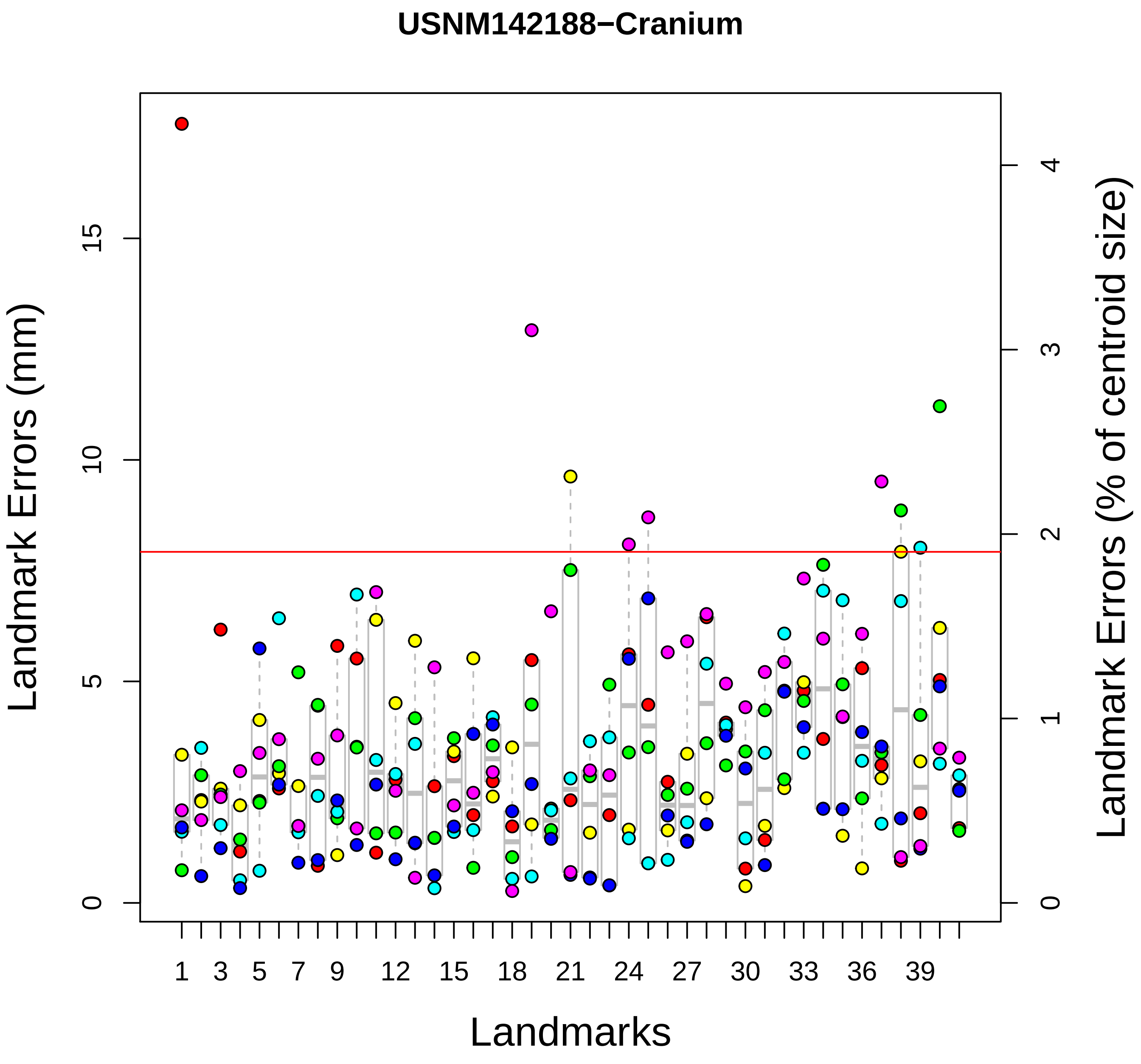

USNM142188−Cranium
Landmark Errors (mm)
Landmark Errors (% of centroid size)
Landmarks
0
5
10
15
0
1
2
3
4
1
3
5
7
9
12
15
18
21
24
27
30
33
36
39

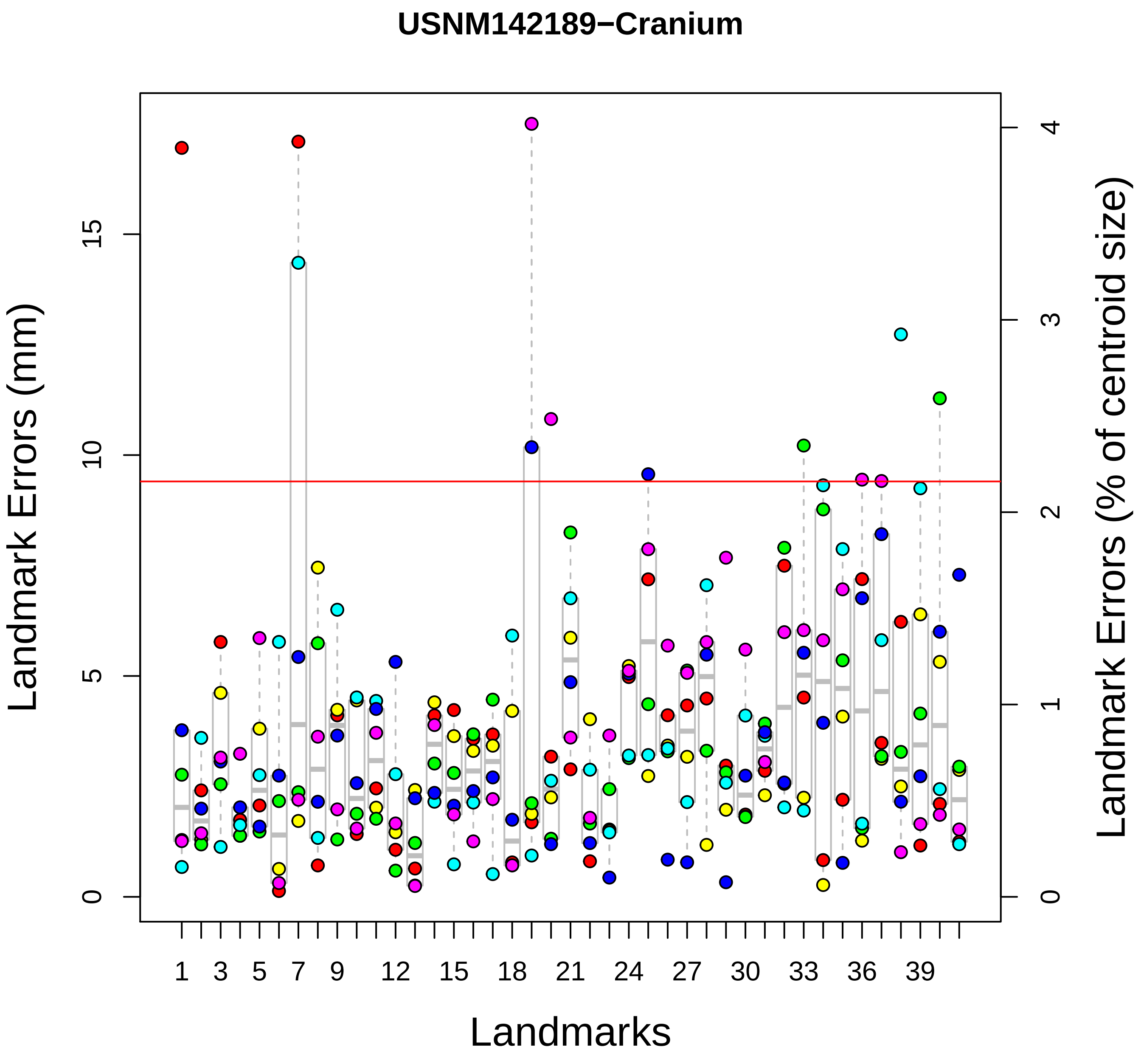

USNM142189−Cranium
Landmark Errors (mm)
Landmark Errors (% of centroid size)
Landmarks
0
5
10
15
0
1
2
3
4
1
3
5
7
9
12
15
18
21
24
27
30
33
36
39

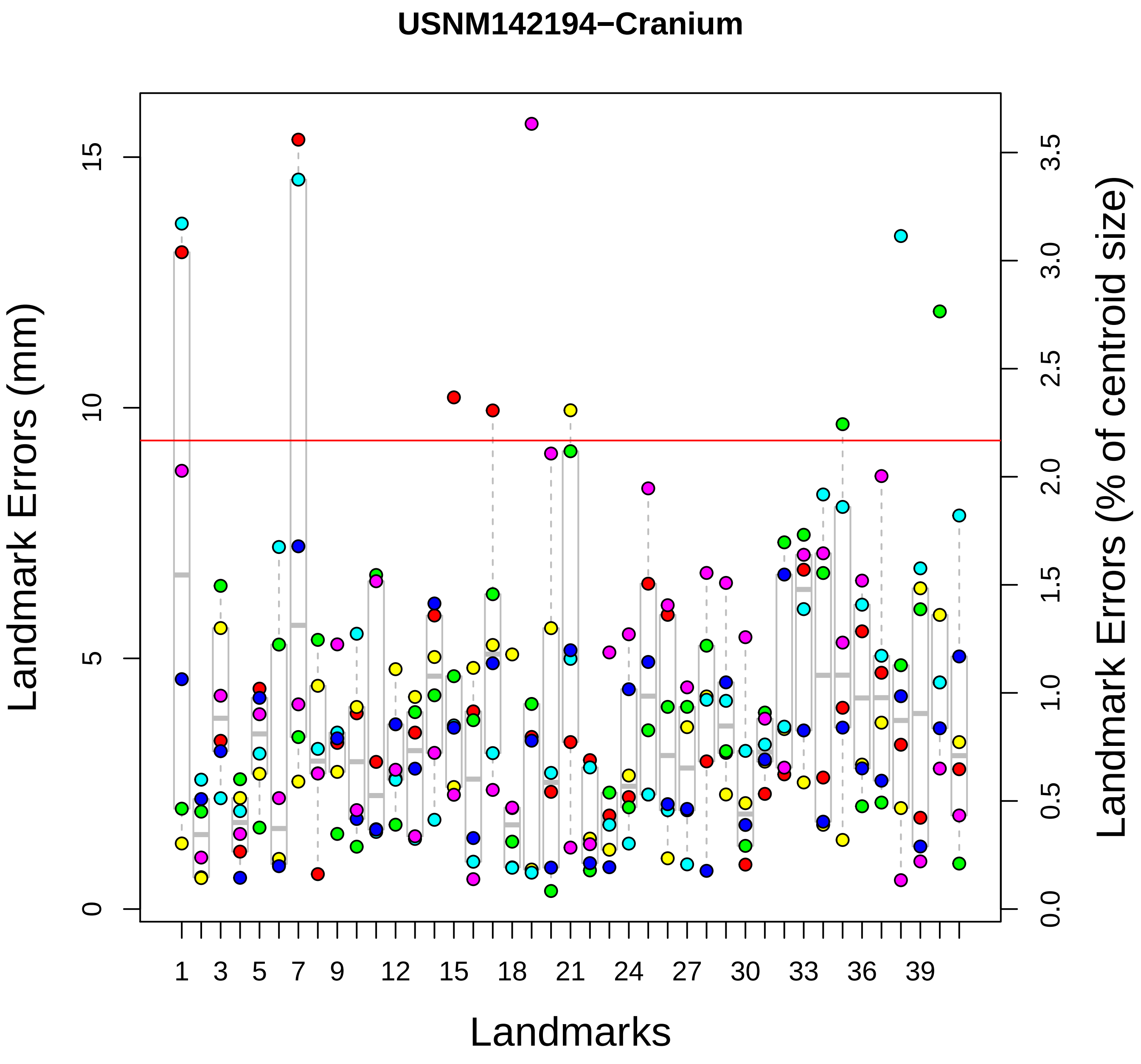
USNM142194−Cranium
Landmark Errors (mm)
Landmark Errors (% of centroid size)
Landmarks
0
5
10
15
0.0
0.5
1.0
1.5
2.0
2.5
3.0
3.5
1
3
5
7
9
12
15
18
21
24
27
30
33
36
39

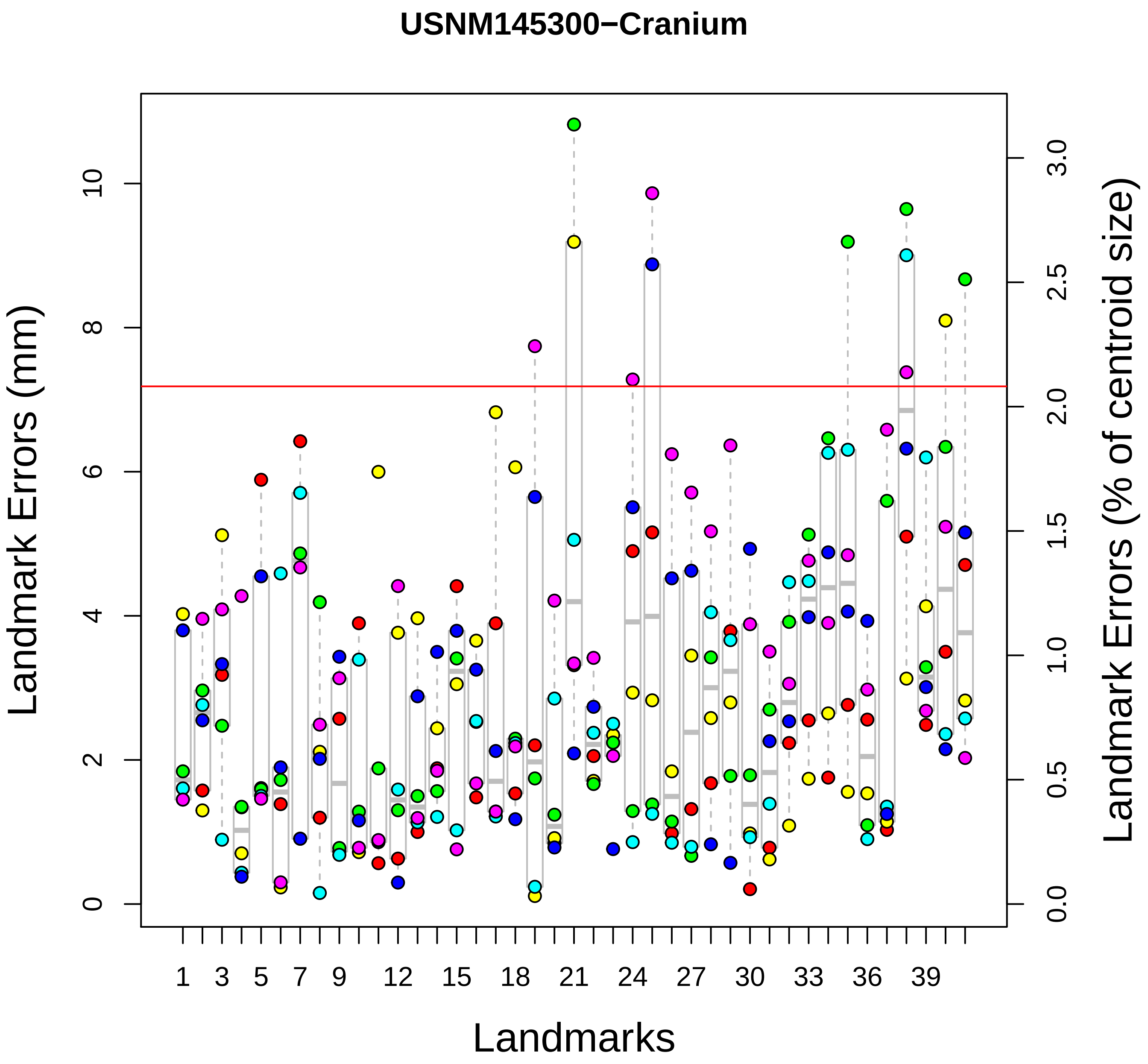

USNM145300−Cranium
Landmark Errors (mm)
Landmark Errors (% of centroid size)
Landmarks
0
2
4
6
8
10
0.0
0.5
1.0
1.5
2.0
2.5
3.0
1
3
5
7
9
12
15
18
21
24
27
30
33
36
39

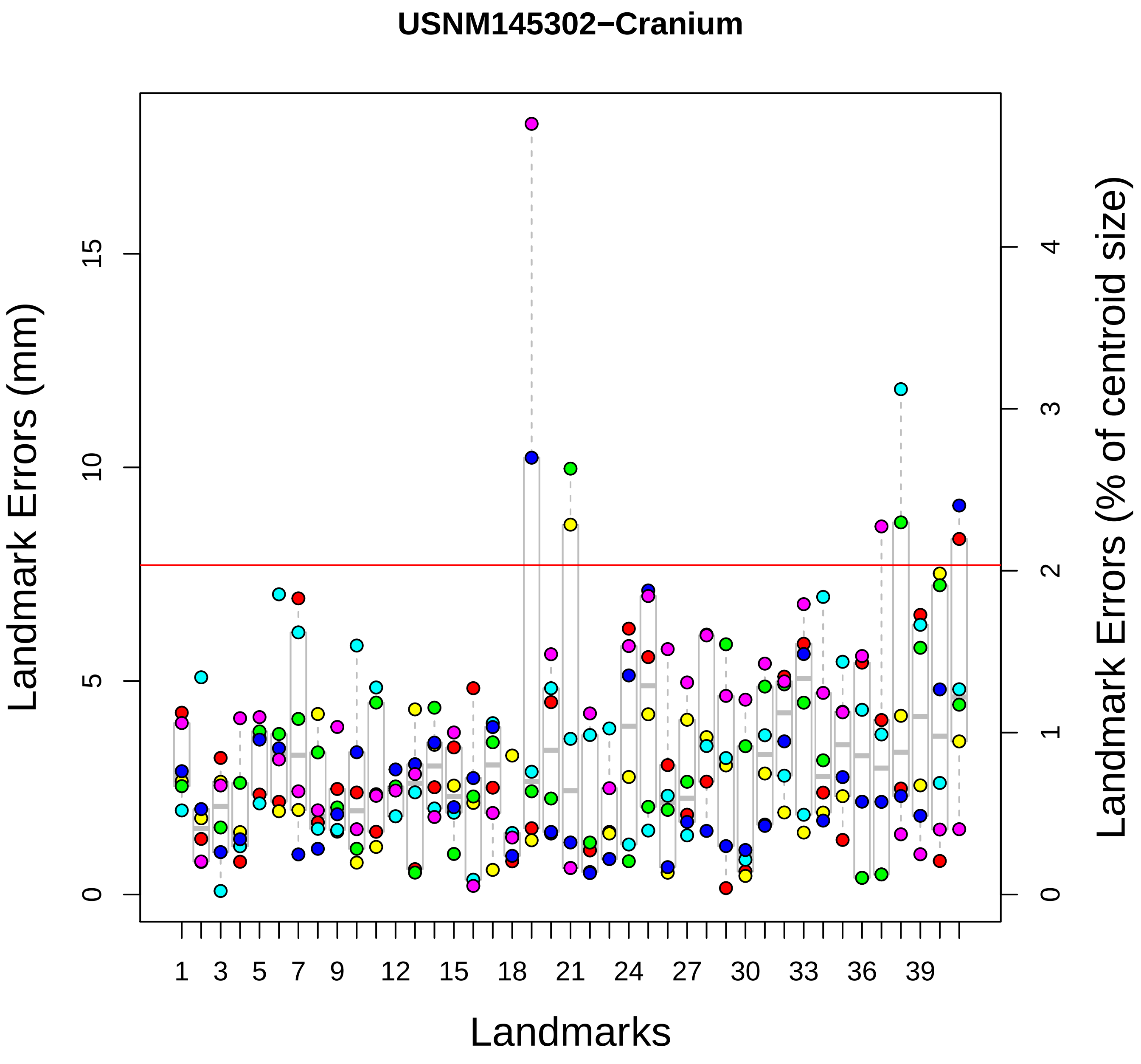
USNM145302−Cranium
Landmark Errors (mm)
Landmark Errors (% of centroid size)
Landmarks
0
5
10
15
0
1
2
3
4
1
3
5
7
9
12
15
18
21
24
27
30
33
36
39

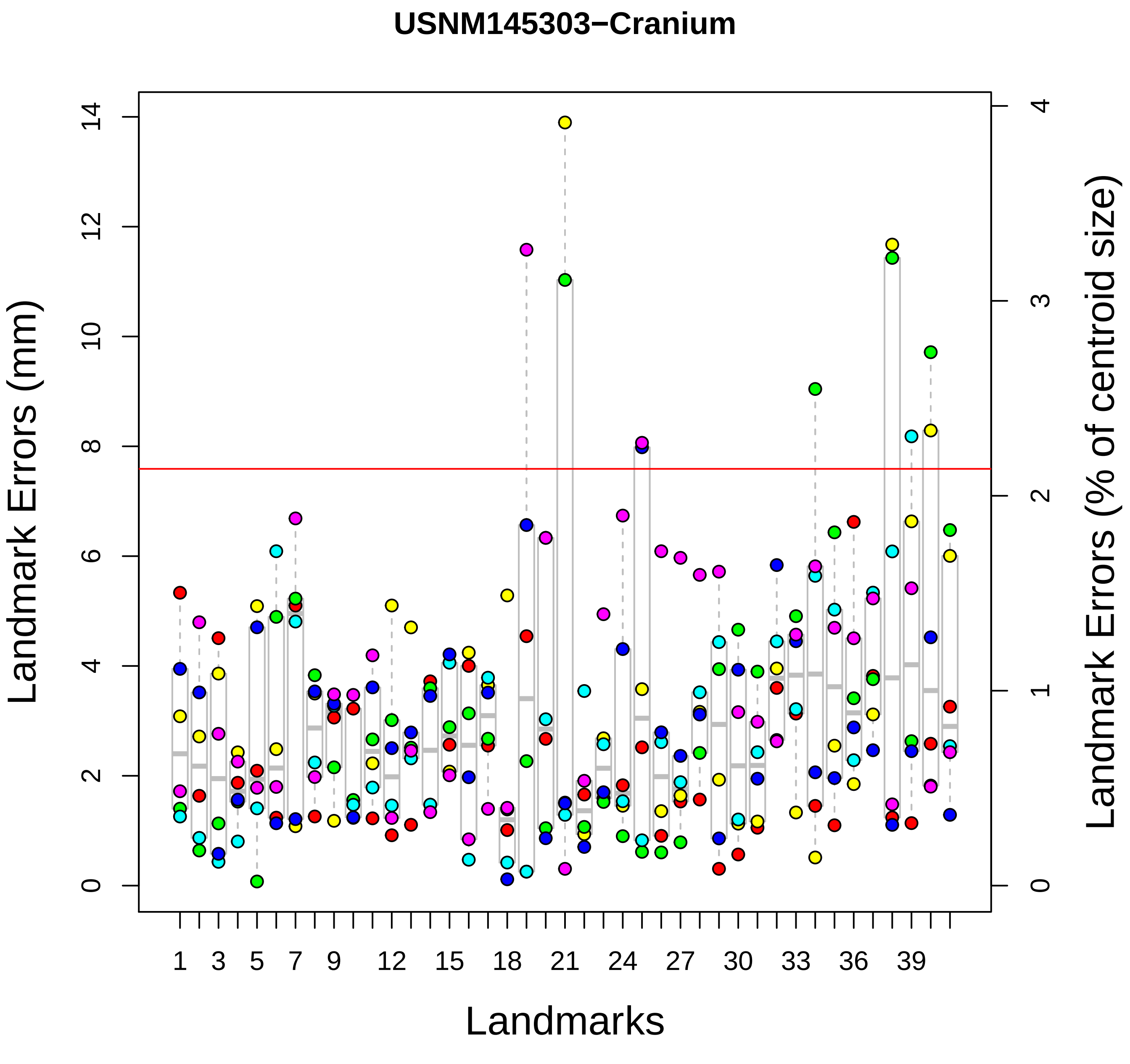
USNM145303−Cranium
Landmark Errors (mm)
Landmark Errors (% of centroid size)
Landmarks
0
2
4
6
8
10
12
14
0
1
2
3
4
1
3
5
7
9
12
15
18
21
24
27
30
33
36
39

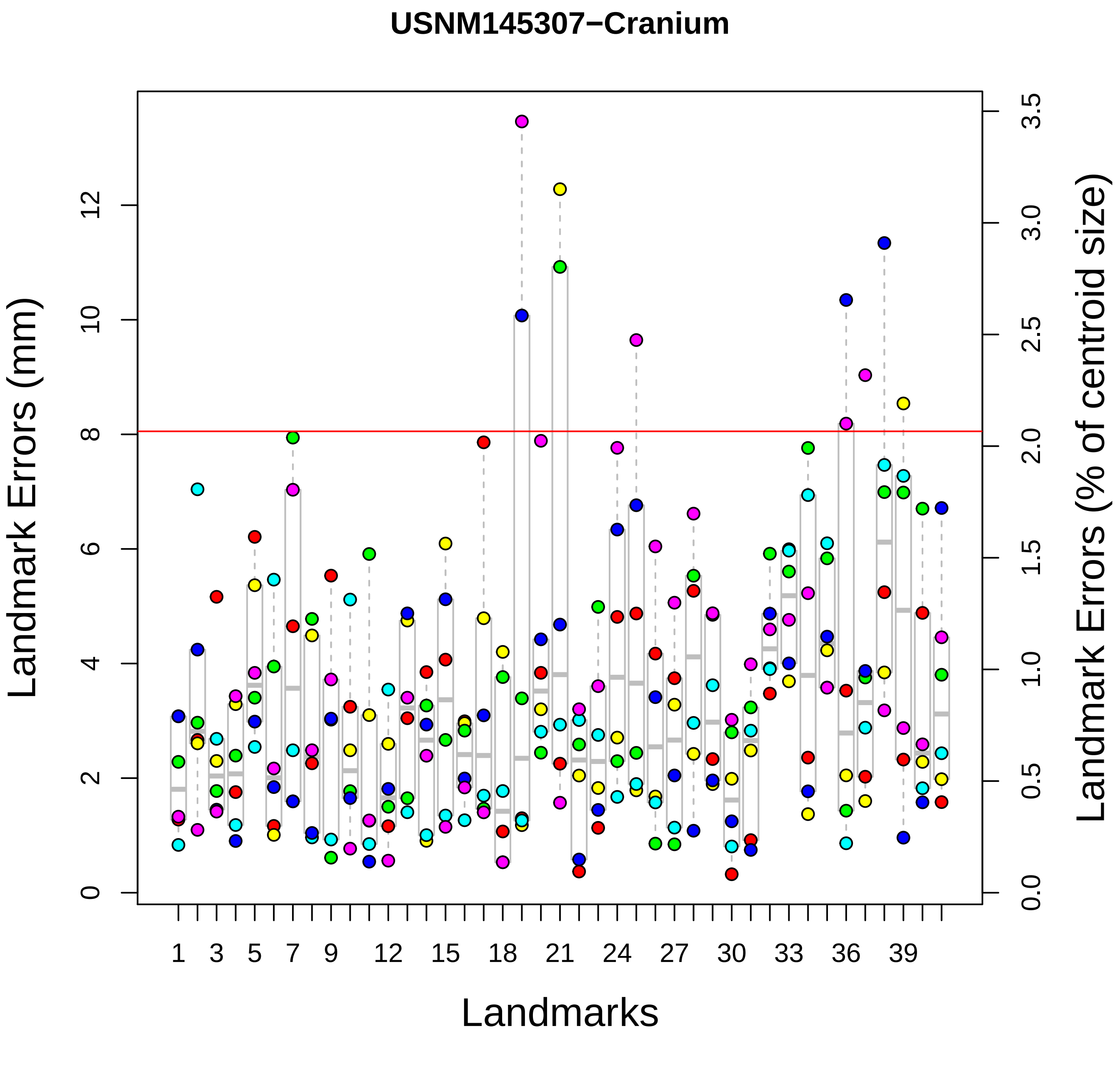

USNM145307−Cranium
Landmark Errors (mm)
Landmark Errors (% of centroid size)
Landmarks
0
2
4
6
8
10
12
0.0
0.5
1.0
1.5
2.0
2.5
3.0
3.5
1
3
5
7
9
12
15
18
21
24
27
30
33
36
39

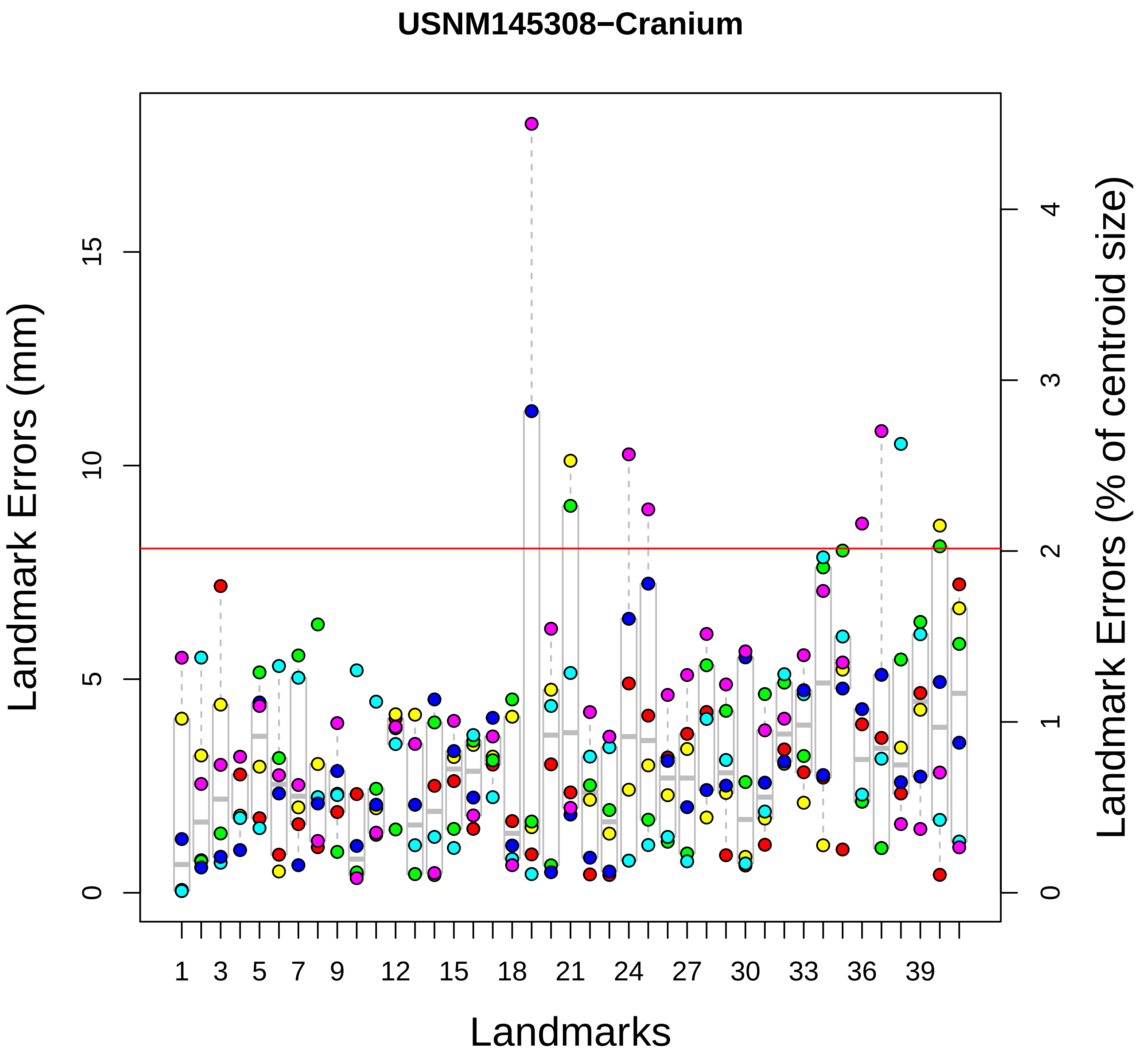

USNM145308−Cranium
Landmark Errors (mm)
Landmark Errors (% of centroid size)
Landmarks
0
5
10
15
0
1
2
3
4
1
3
5
7
9
12
15
18
21
24
27
30
33
36
39

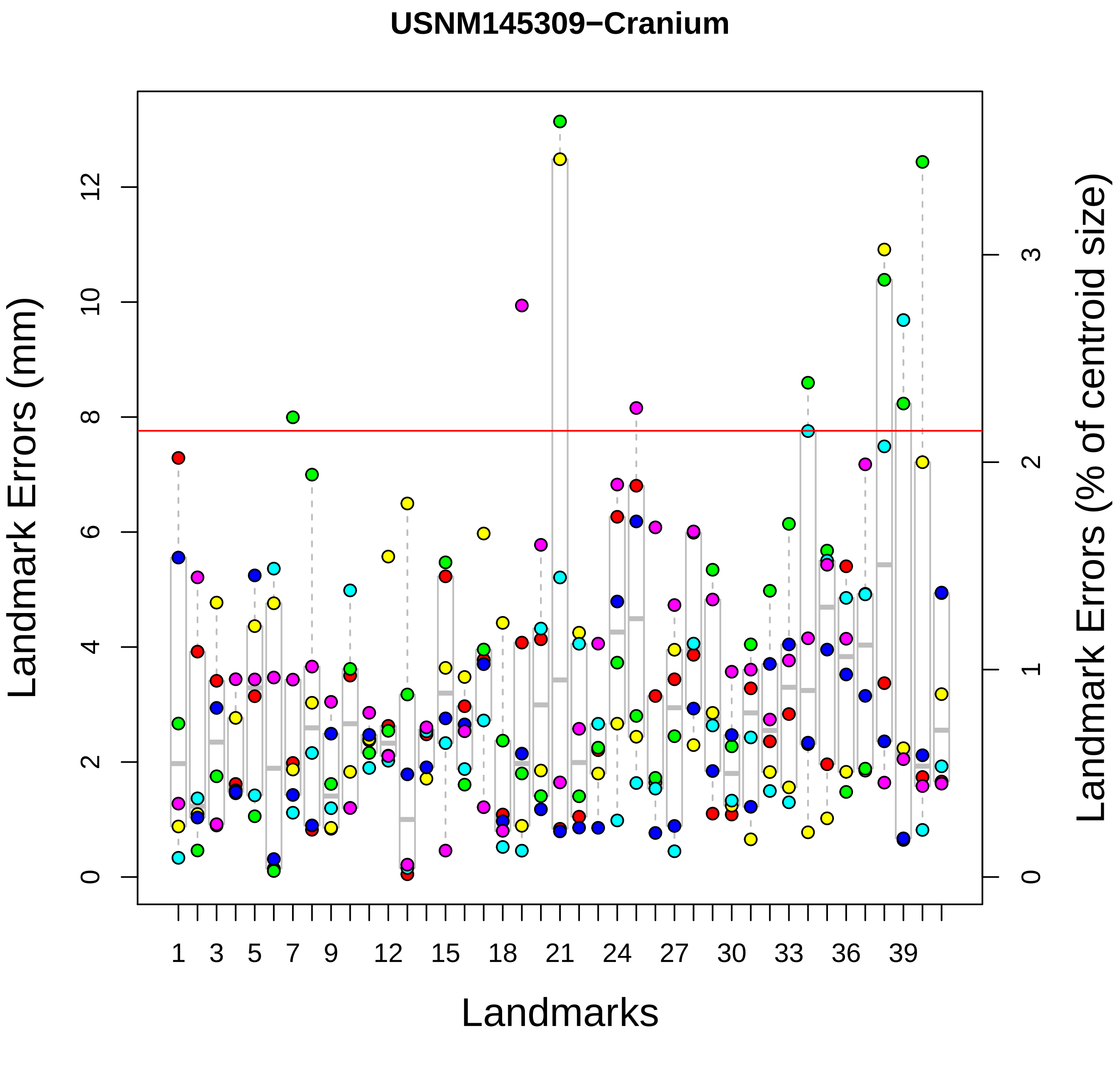

USNM145309−Cranium
Landmark Errors (mm)
Landmark Errors (% of centroid size)
Landmarks
0
2
4
6
8
10
12
0
1
2
3
1
3
5
7
9
12
15
18
21
24
27
30
33
36
39

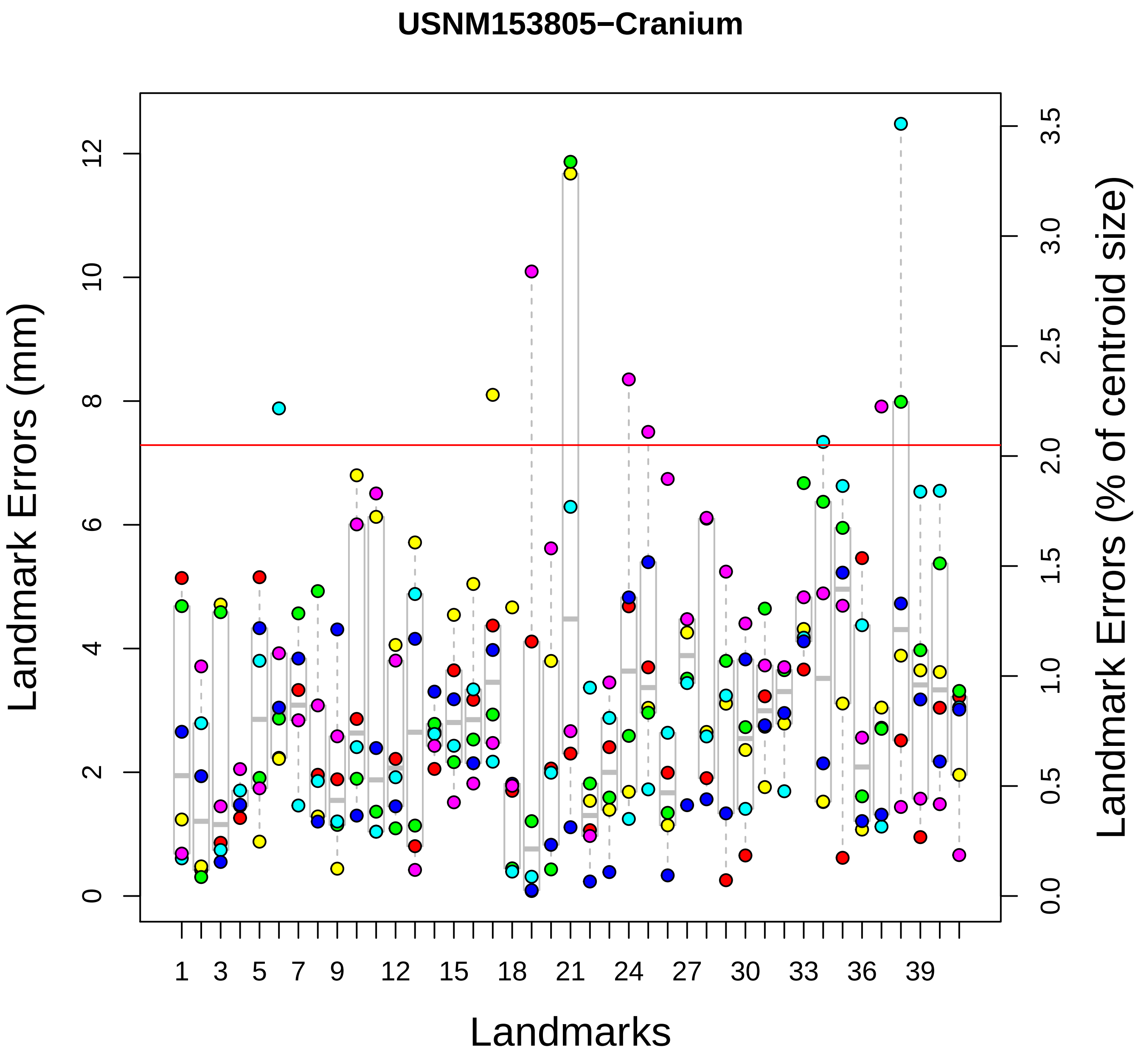
USNM153805−Cranium
Landmark Errors (mm)
Landmark Errors (% of centroid size)
Landmarks
0
2
4
6
8
10
12
0.0
0.5
1.0
1.5
2.0
2.5
3.0
3.5
1
3
5
7
9
12
15
18
21
24
27
30
33
36
39

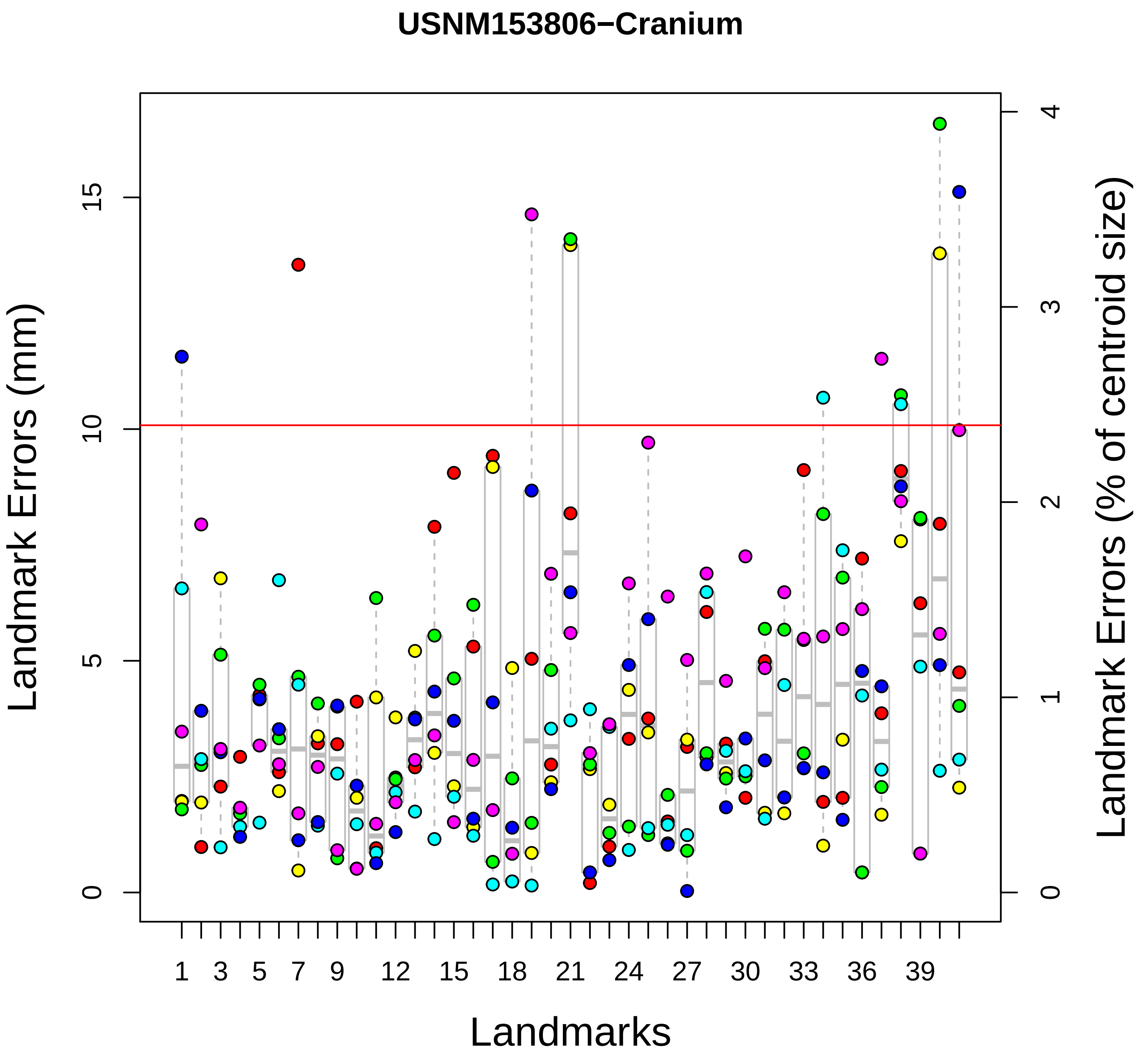
USNM153806−Cranium
Landmark Errors (mm)
Landmark Errors (% of centroid size)
Landmarks
0
5
10
15
0
1
2
3
4
1
3
5
7
9
12
15
18
21
24
27
30
33
36
39

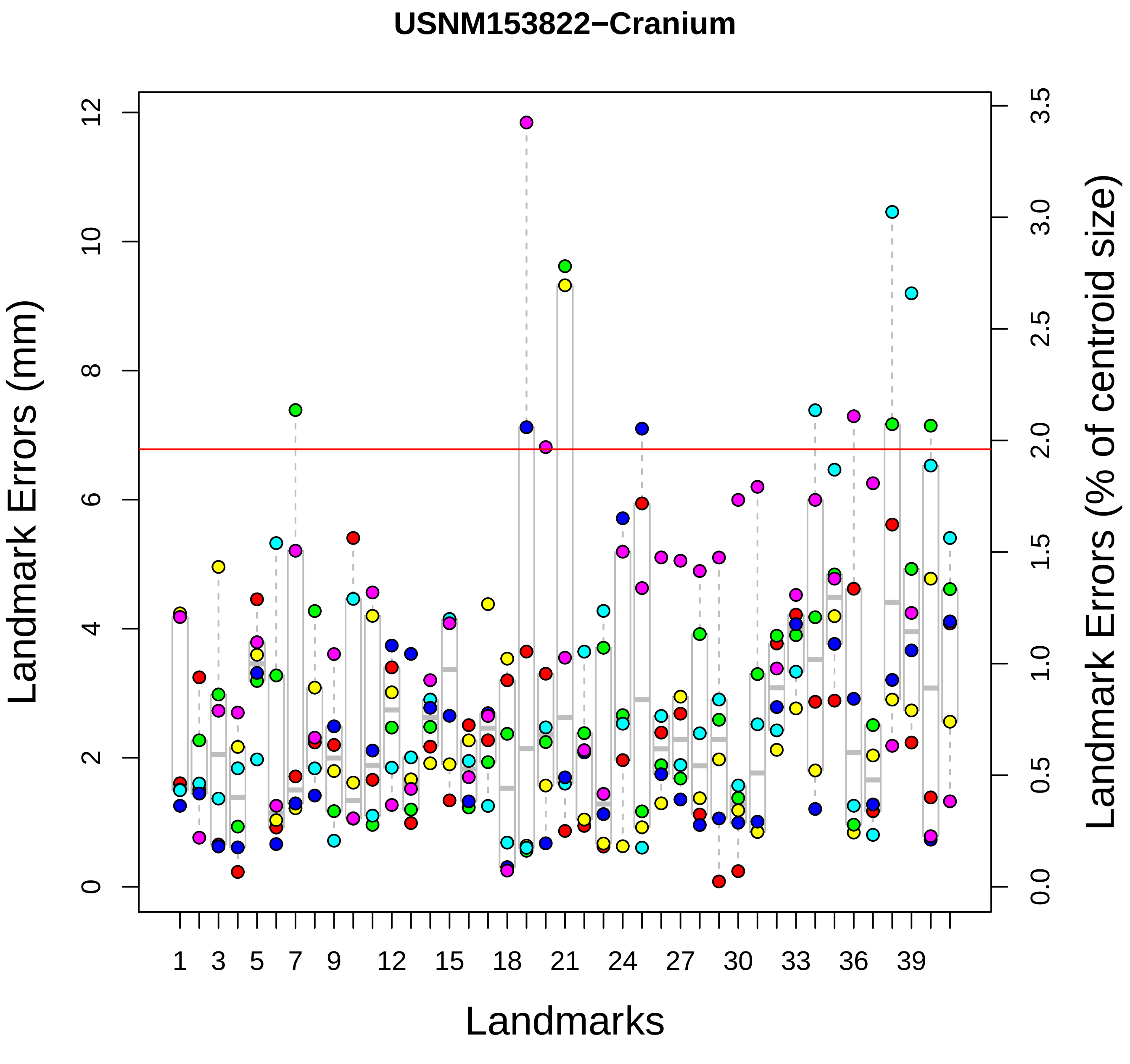
USNM153822−Cranium
Landmark Errors (mm)
Landmark Errors (% of centroid size)
Landmarks
0
2
4
6
8
10
12
0.0
0.5
1.0
1.5
2.0
2.5
3.0
3.5
1
3
5
7
9
12
15
18
21
24
27
30
33
36
39

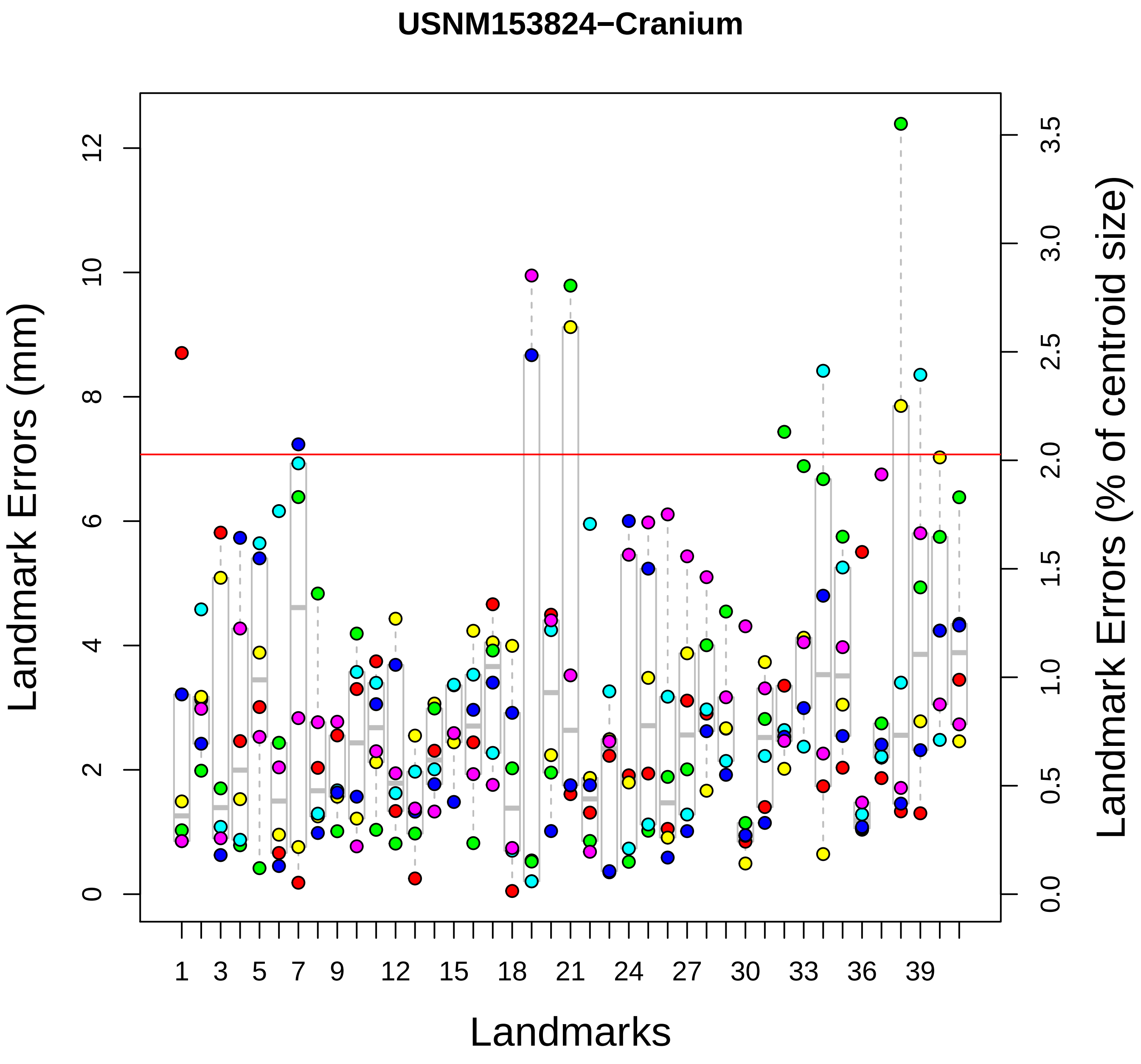

USNM153824−Cranium
Landmark Errors (mm)
Landmark Errors (% of centroid size)
Landmarks
0
2
4
6
8
10
12
0.0
0.5
1.0
1.5
2.0
2.5
3.0
3.5
1
3
5
7
9
12
15
18
21
24
27
30
33
36
39

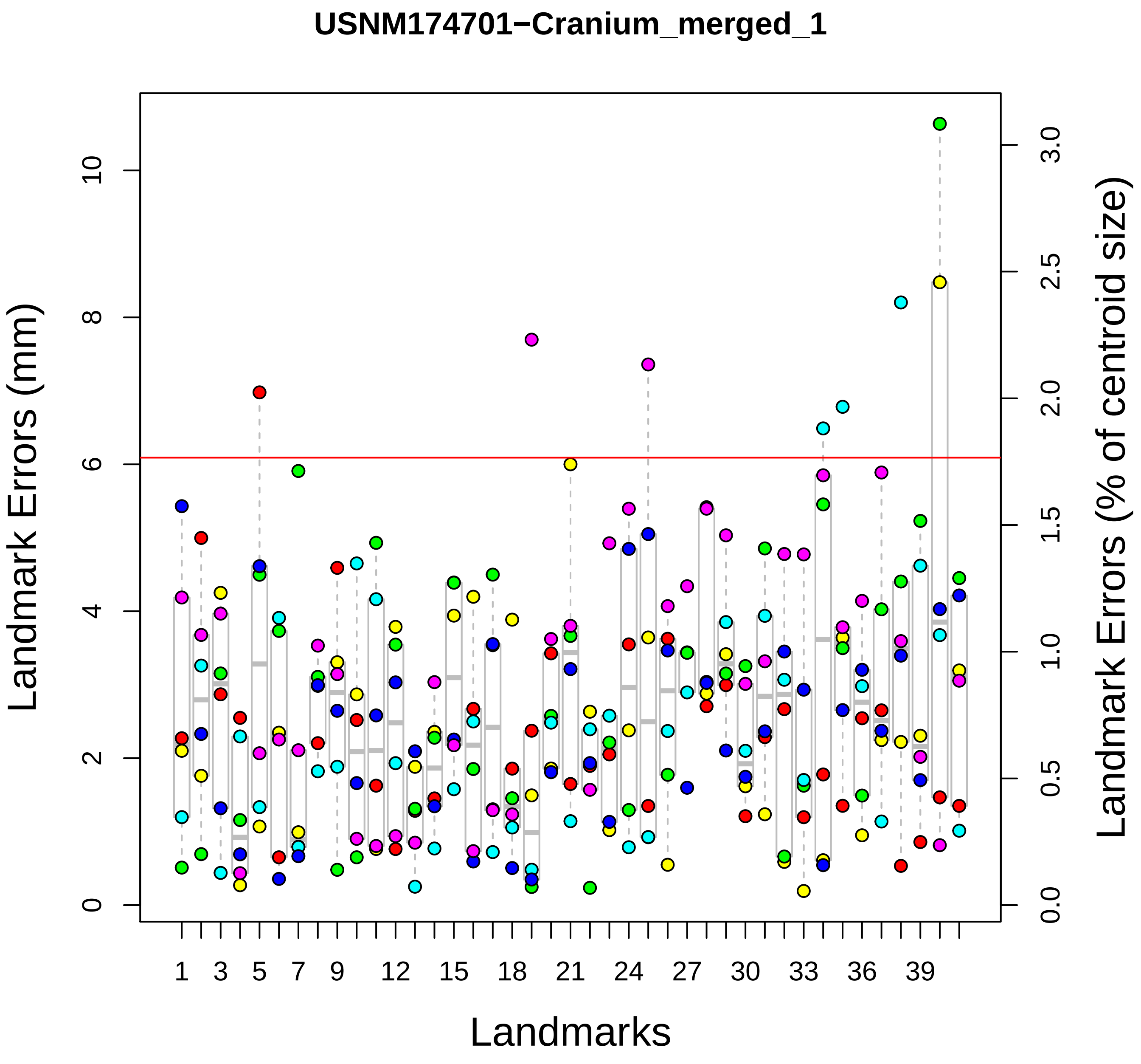
USNM174701−Cranium_merged_1
Landmark Errors (mm)
Landmark Errors (% of centroid size)
Landmarks
0
2
4
6
8
10
0.0
0.5
1.0
1.5
2.0
2.5
3.0
1
3
5
7
9
12
15
18
21
24
27
30
33
36
39

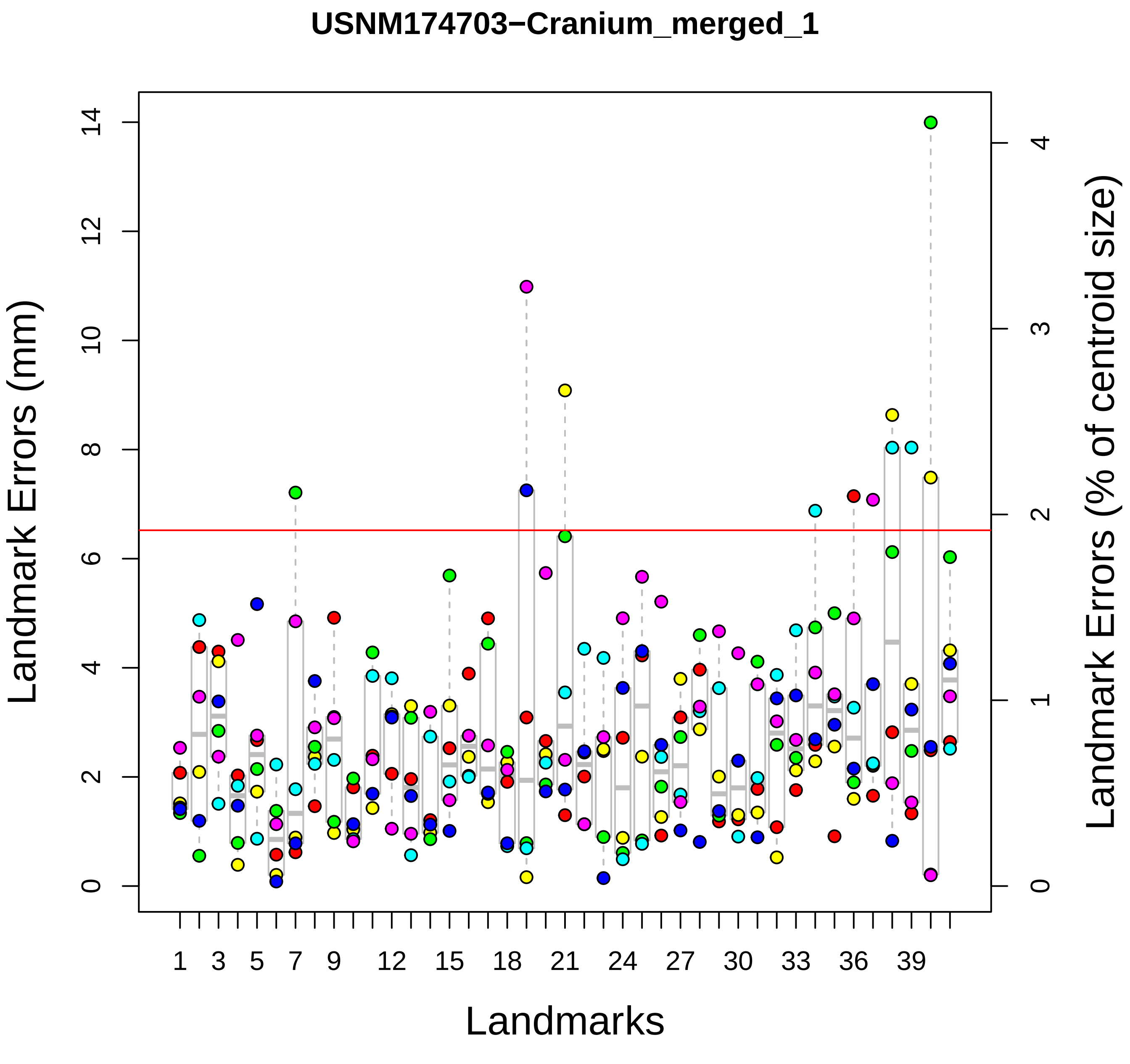

USNM174703−Cranium_merged_1
Landmark Errors (mm)
Landmark Errors (% of centroid size)
Landmarks
0
2
4
6
8
10
12
14
0
1
2
3
4
1
3
5
7
9
12
15
18
21
24
27
30
33
36
39

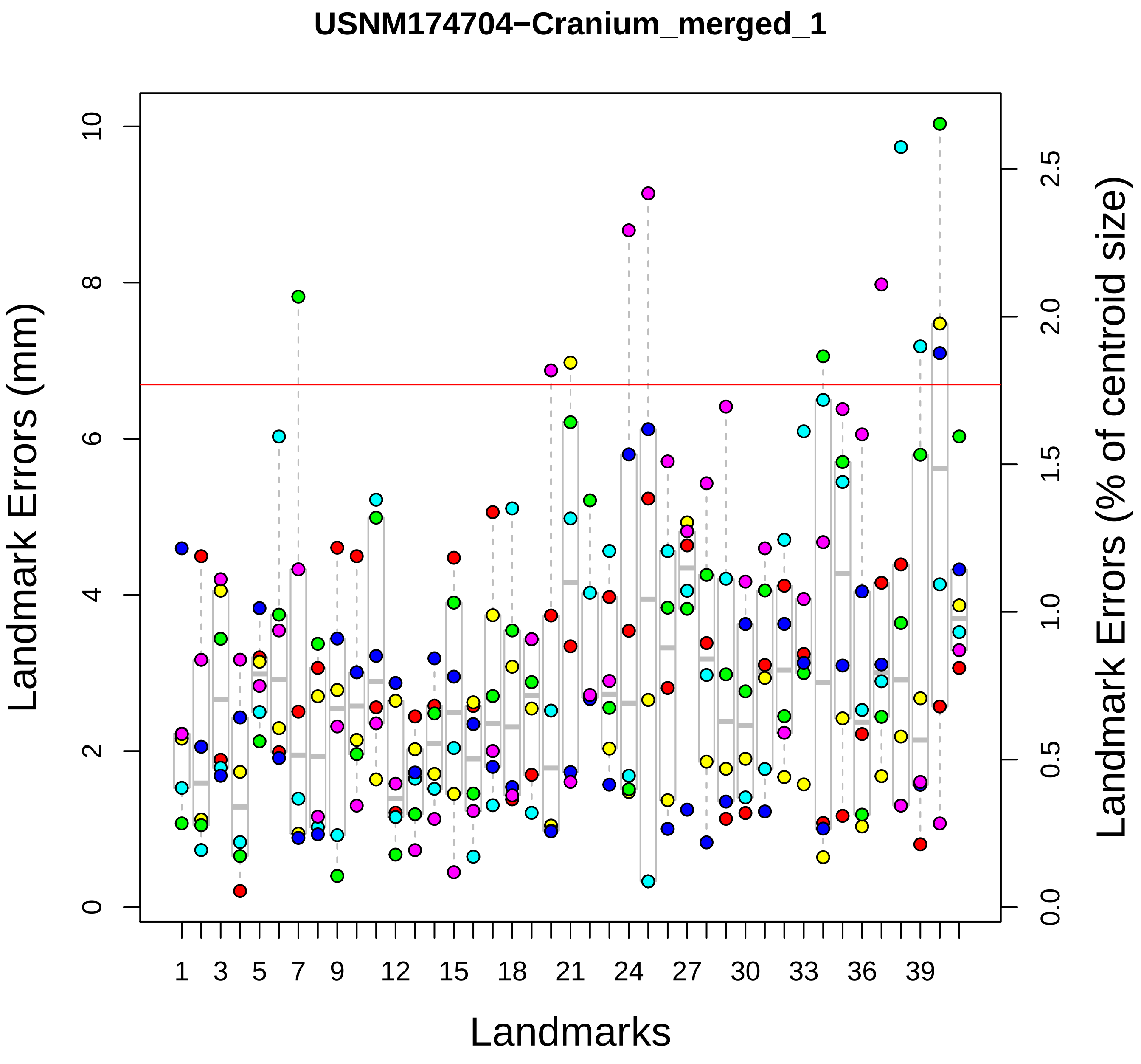

USNM174704−Cranium_merged_1
Landmark Errors (mm)
Landmark Errors (% of centroid size)
Landmarks
0
2
4
6
8
10
0.0
0.5
1.0
1.5
2.0
2.5
1
3
5
7
9
12
15
18
21
24
27
30
33
36
39

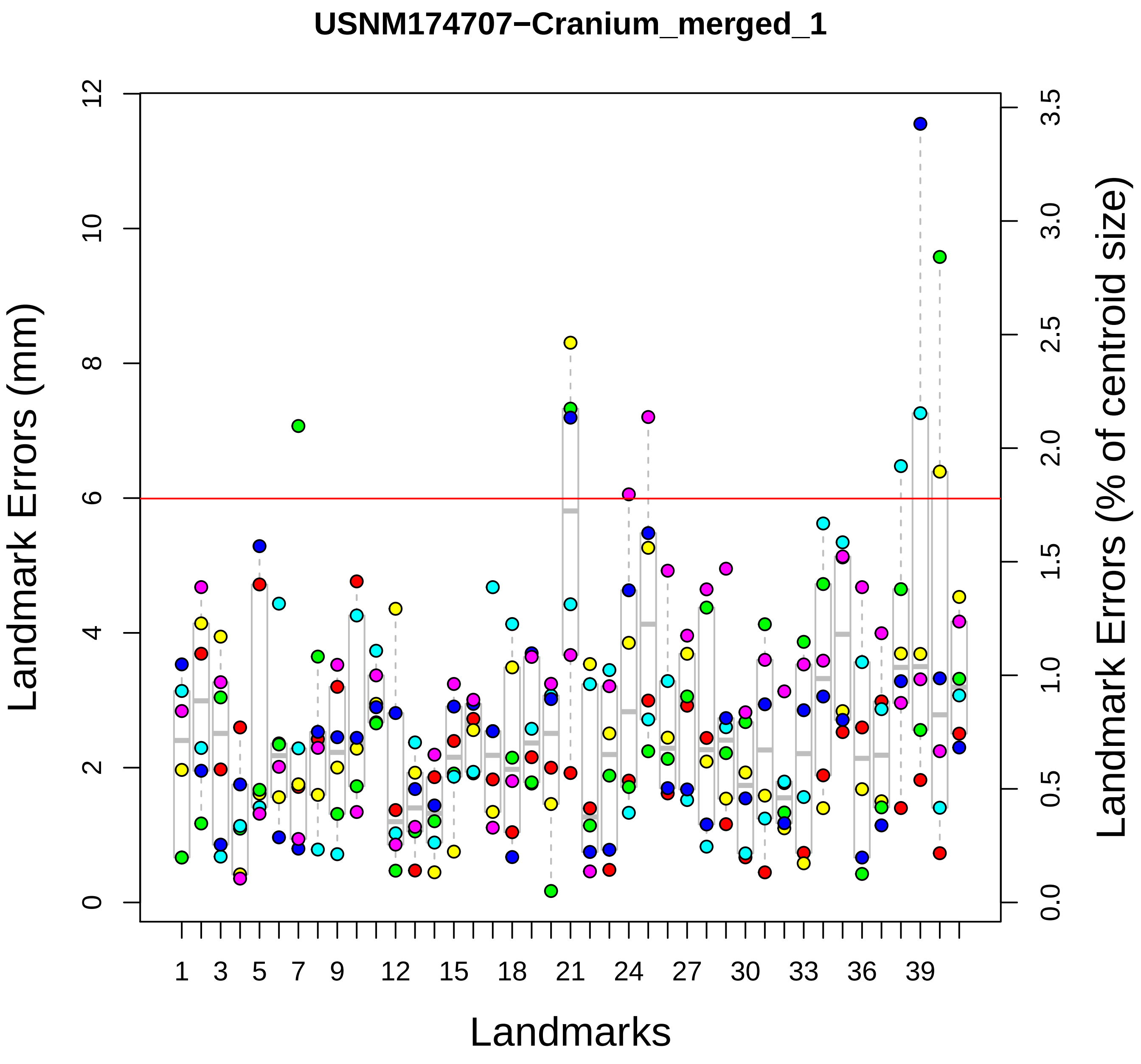
USNM174707−Cranium_merged_1
Landmark Errors (mm)
Landmark Errors (% of centroid size)
Landmarks
0
2
4
6
8
10
12
0.0
0.5
1.0
1.5
2.0
2.5
3.0
3.5
1
3
5
7
9
12
15
18
21
24
27
30
33
36
39

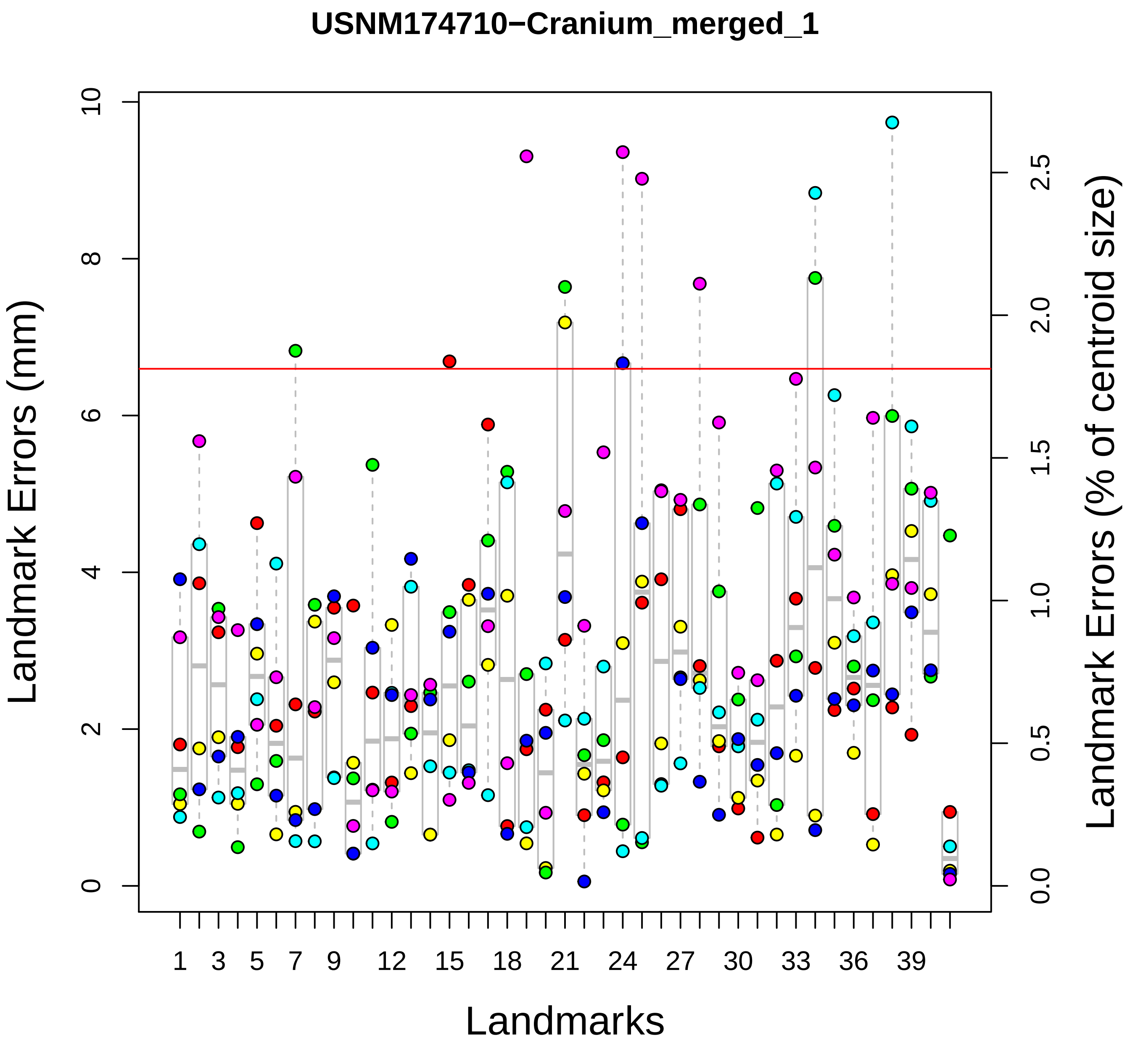

USNM174710−Cranium_merged_1
Landmark Errors (mm)
Landmark Errors (% of centroid size)
Landmarks
0
2
4
6
8
10
0.0
0.5
1.0
1.5
2.0
2.5
1
3
5
7
9
12
15
18
21
24
27
30
33
36
39

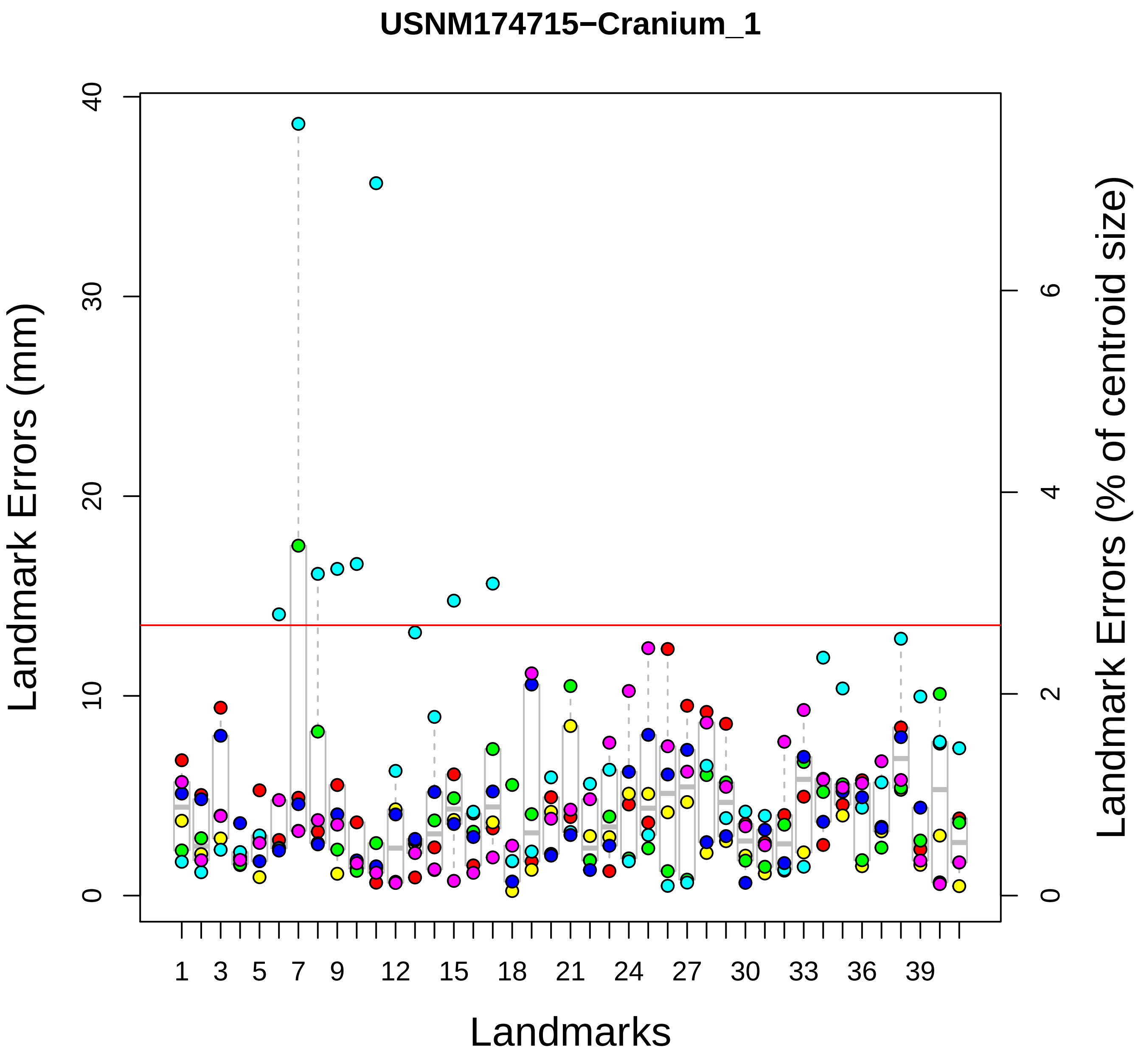

USNM174715−Cranium_1
Landmark Errors (mm)
Landmark Errors (% of centroid size)
Landmarks
0
10
20
30
40
0
2
4
6
1
3
5
7
9
12
15
18
21
24
27
30
33
36
39

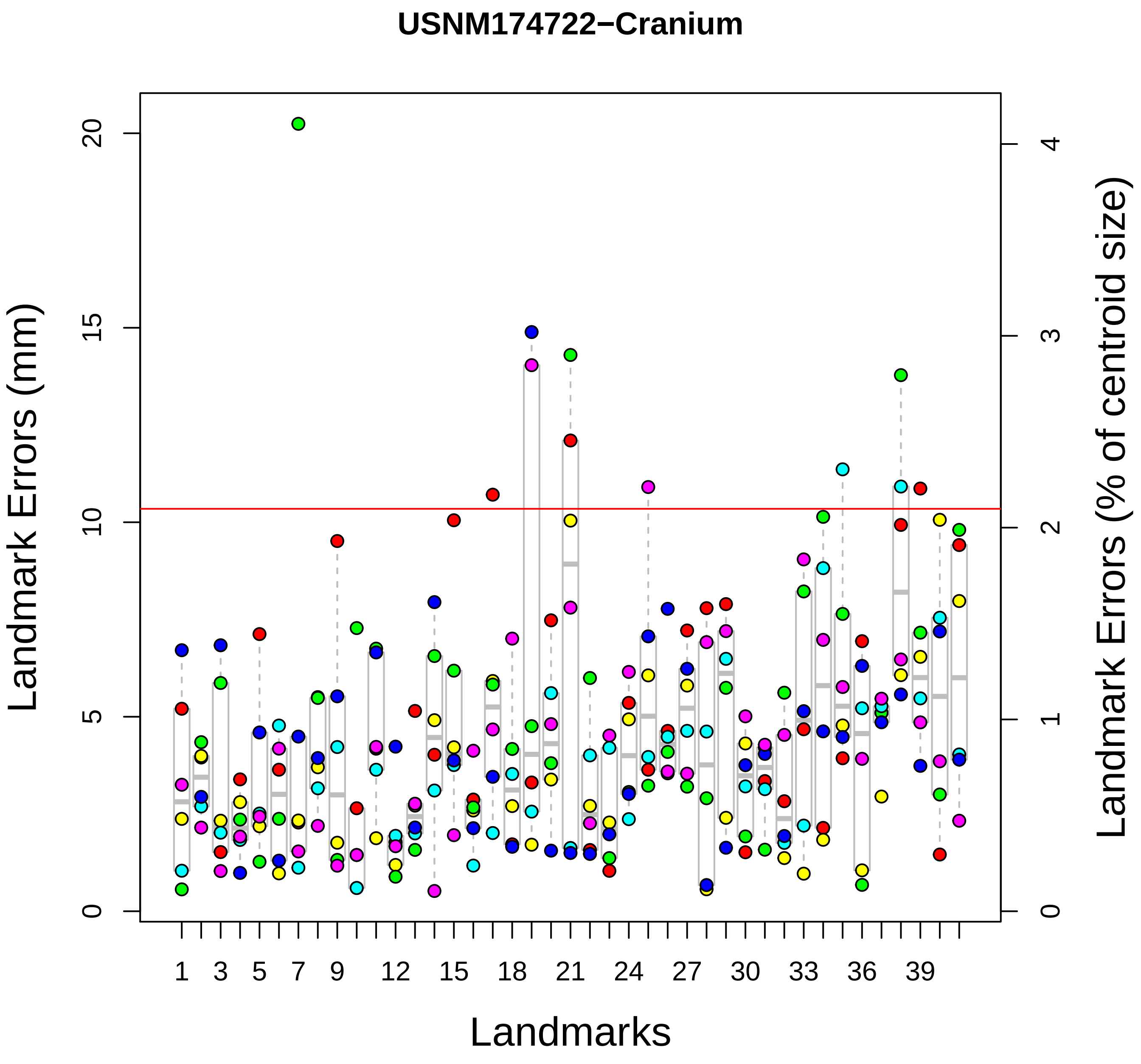

USNM174722−Cranium
Landmark Errors (mm)
Landmark Errors (% of centroid size)
Landmarks
0
5
10
15
20
0
1
2
3
4
1
3
5
7
9
12
15
18
21
24
27
30
33
36
39

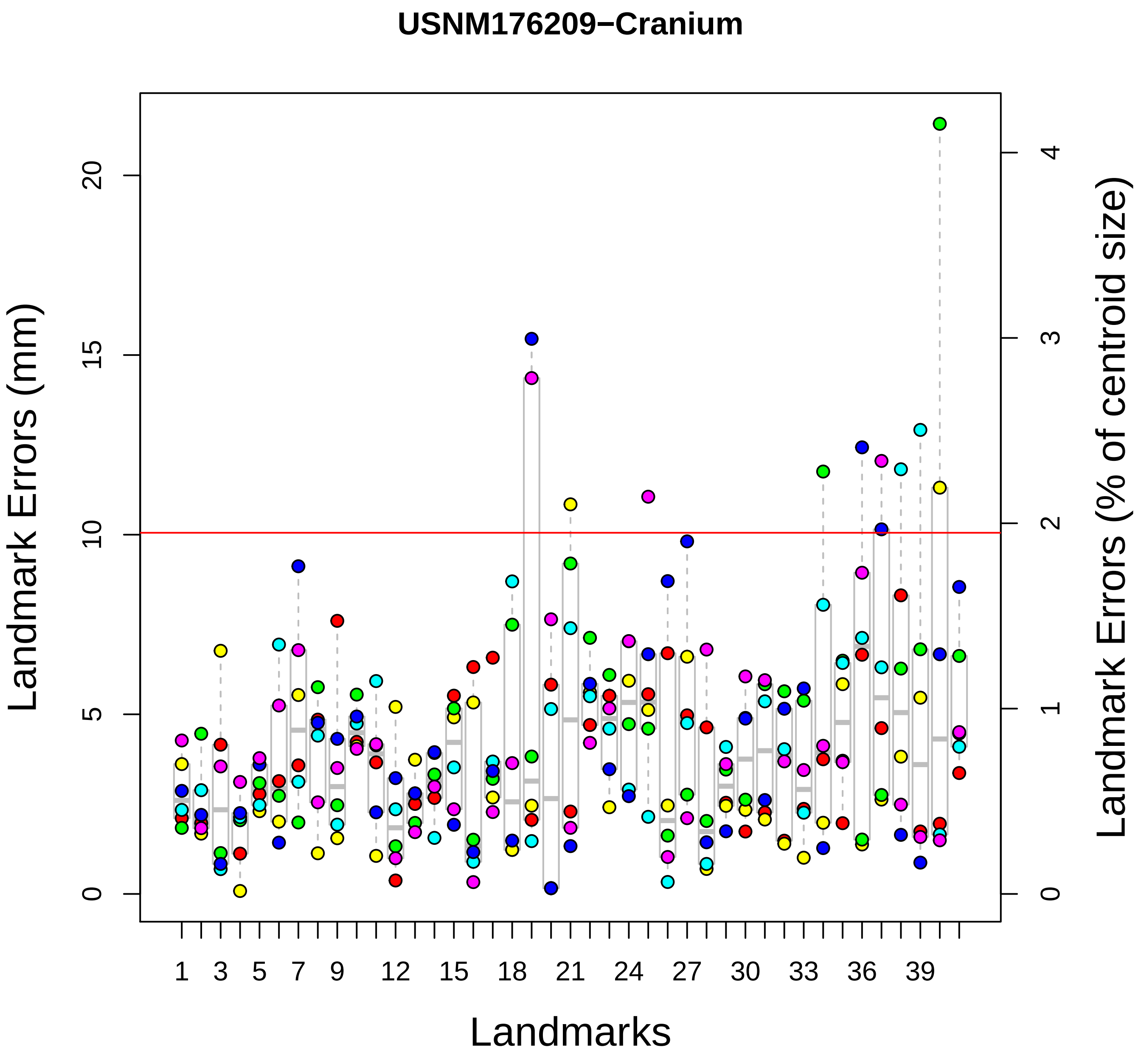

USNM176209−Cranium
Landmark Errors (mm)
Landmark Errors (% of centroid size)
Landmarks
0
5
10
15
20
0
1
2
3
4
1
3
5
7
9
12
15
18
21
24
27
30
33
36
39

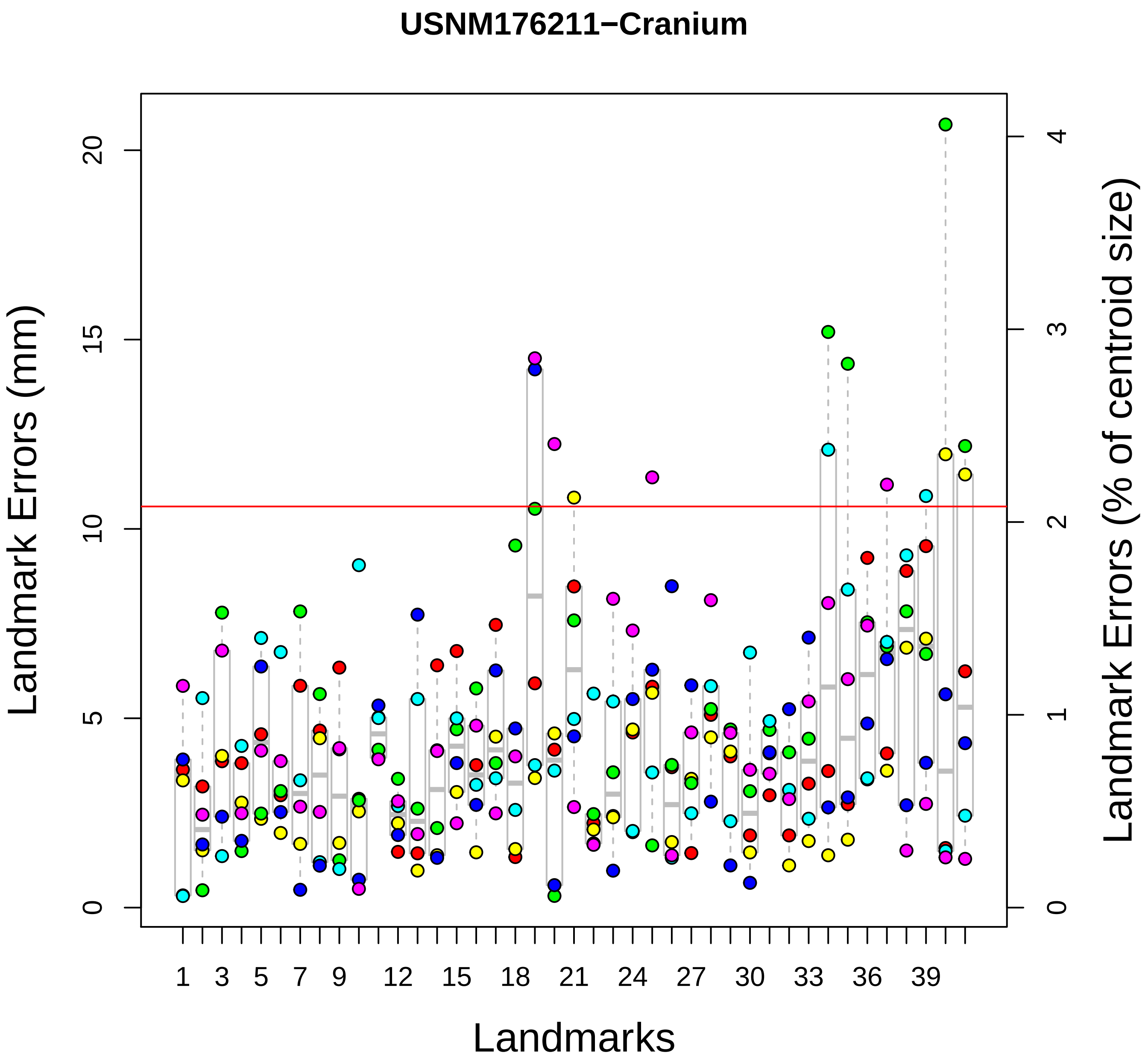

USNM176211−Cranium
Landmark Errors (mm)
Landmark Errors (% of centroid size)
Landmarks
0
5
10
15
20
0
1
2
3
4
1
3
5
7
9
12
15
18
21
24
27
30
33
36
39

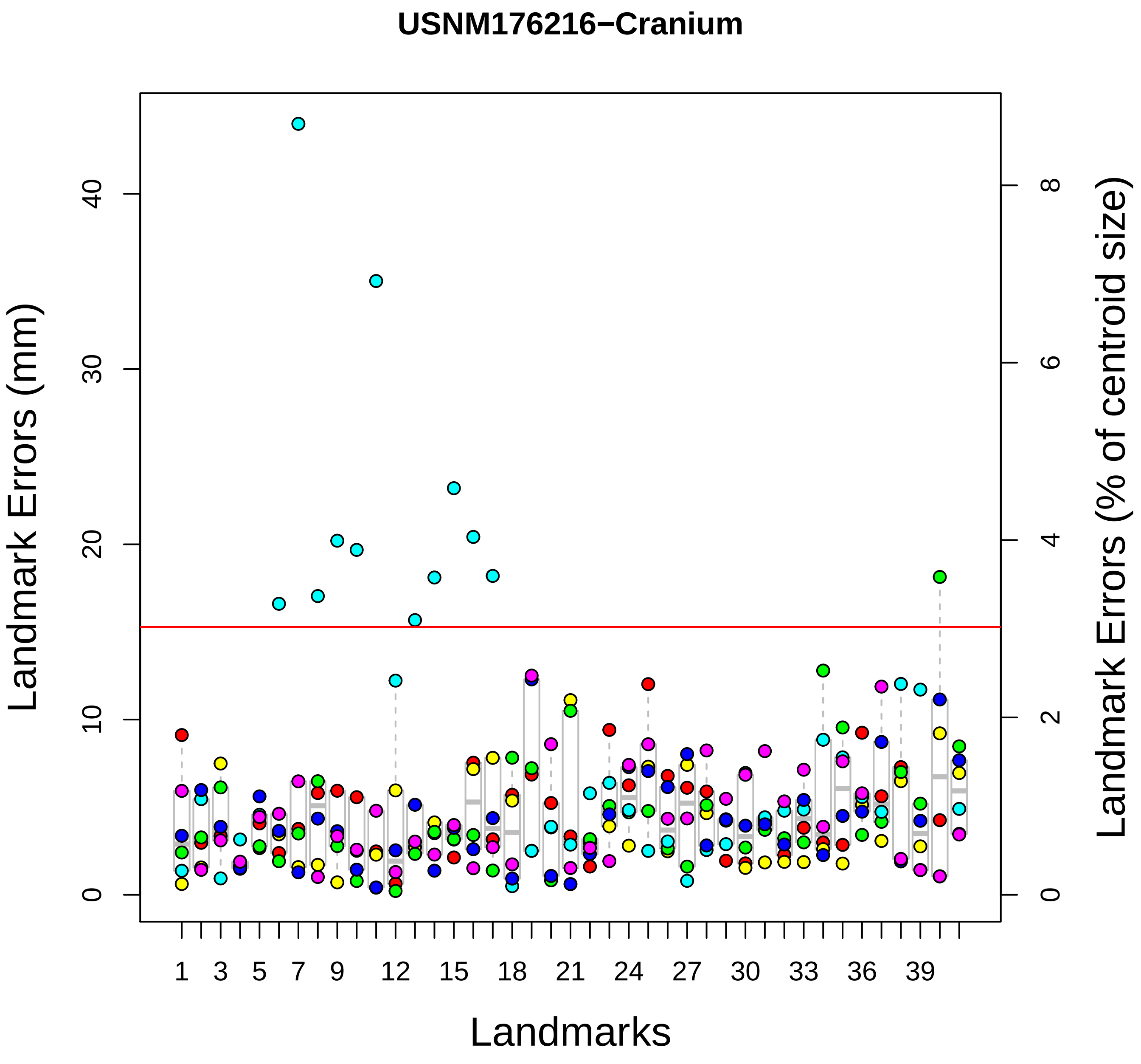

USNM176216−Cranium
Landmark Errors (mm)
Landmark Errors (% of centroid size)
Landmarks
0
10
20
30
40
0
2
4
6
8
1
3
5
7
9
12
15
18
21
24
27
30
33
36
39

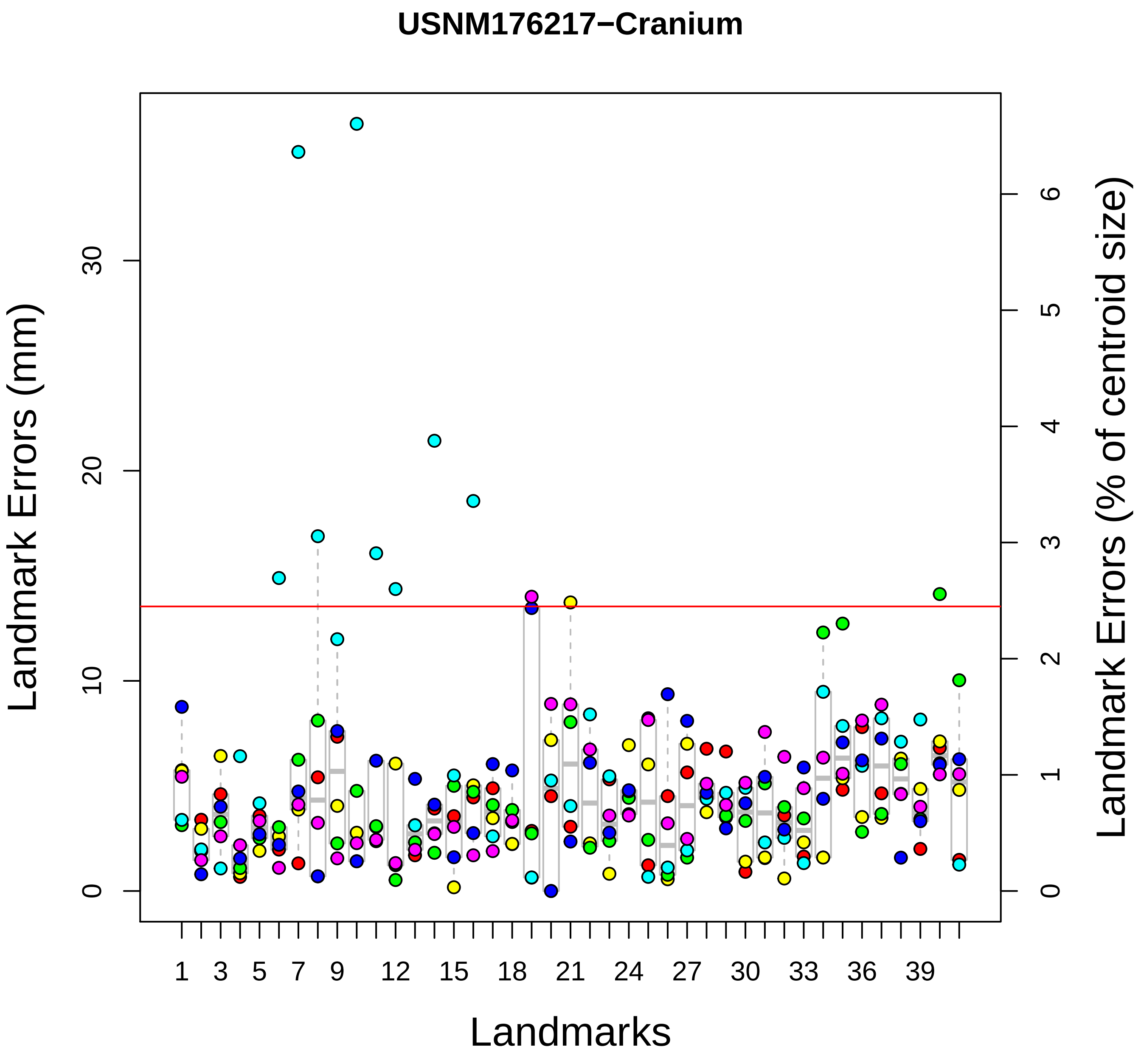
USNM176217−Cranium
Landmark Errors (mm)
Landmark Errors (% of centroid size)
Landmarks
0
10
20
30
0
1
2
3
4
5
6
1
3
5
7
9
12
15
18
21
24
27
30
33
36
39

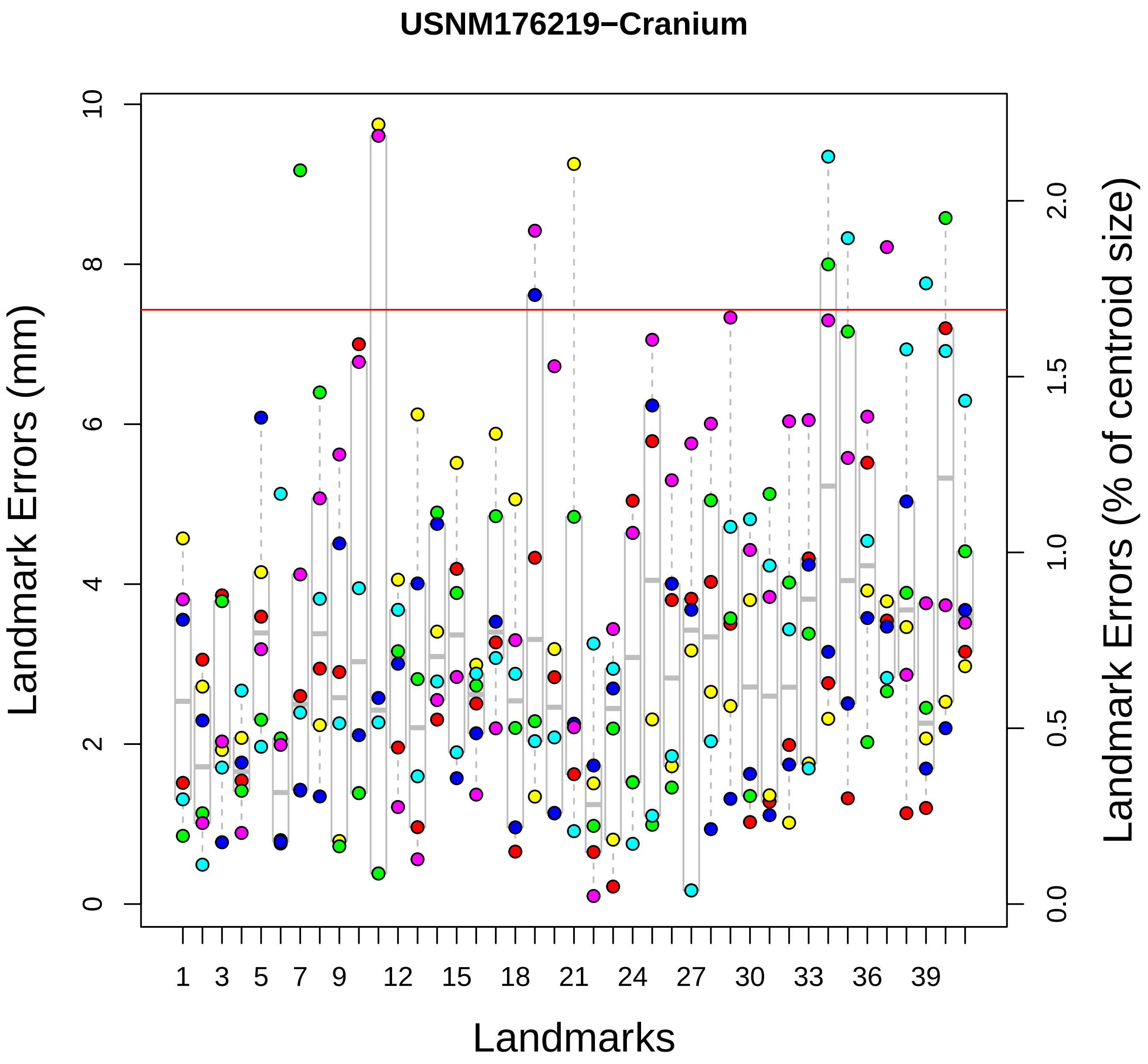

USNM176219−Cranium
Landmark Errors (mm)
Landmark Errors (% of centroid size)
Landmarks
0
2
4
6
8
10
0.0
0.5
1.0
1.5
2.0
1
3
5
7
9
12
15
18
21
24
27
30
33
36
39

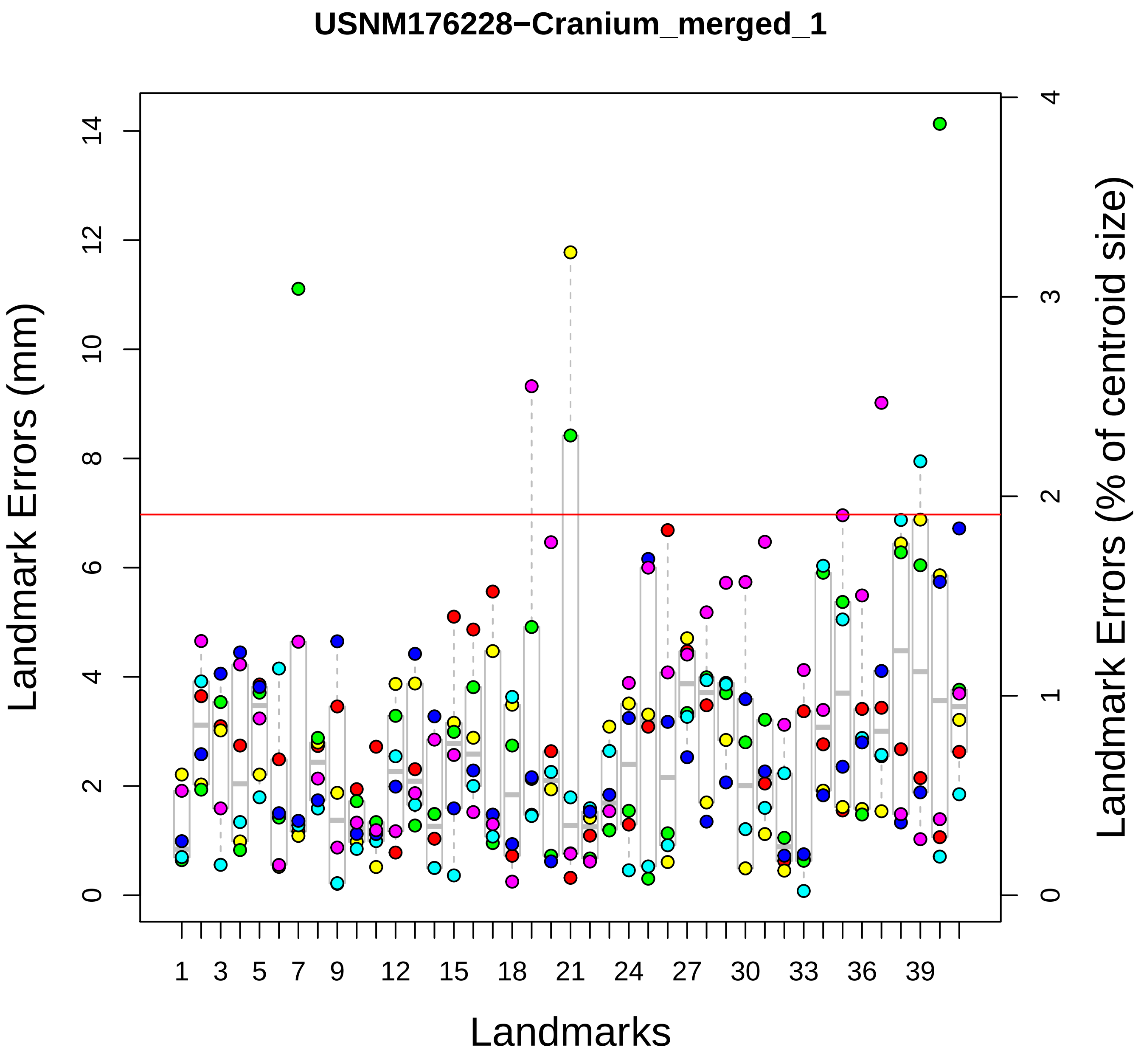

USNM176228−Cranium_merged_1
Landmark Errors (mm)
Landmark Errors (% of centroid size)
Landmarks
0
2
4
6
8
10
12
14
0
1
2
3
4
1
3
5
7
9
12
15
18
21
24
27
30
33
36
39

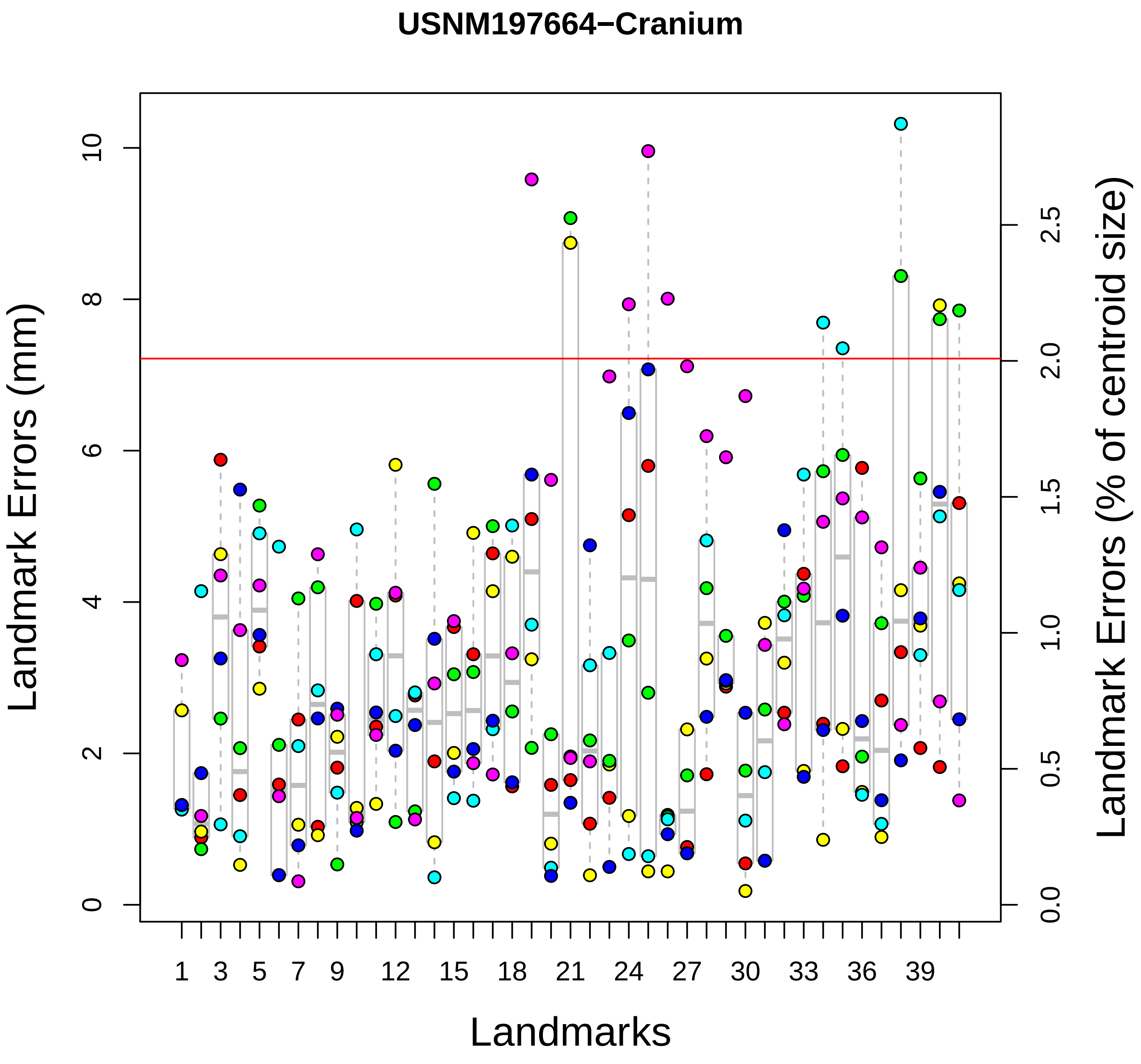

USNM197664−Cranium
Landmark Errors (mm)
Landmark Errors (% of centroid size)
Landmarks
0
2
4
6
8
10
0.0
0.5
1.0
1.5
2.0
2.5
1
3
5
7
9
12
15
18
21
24
27
30
33
36
39

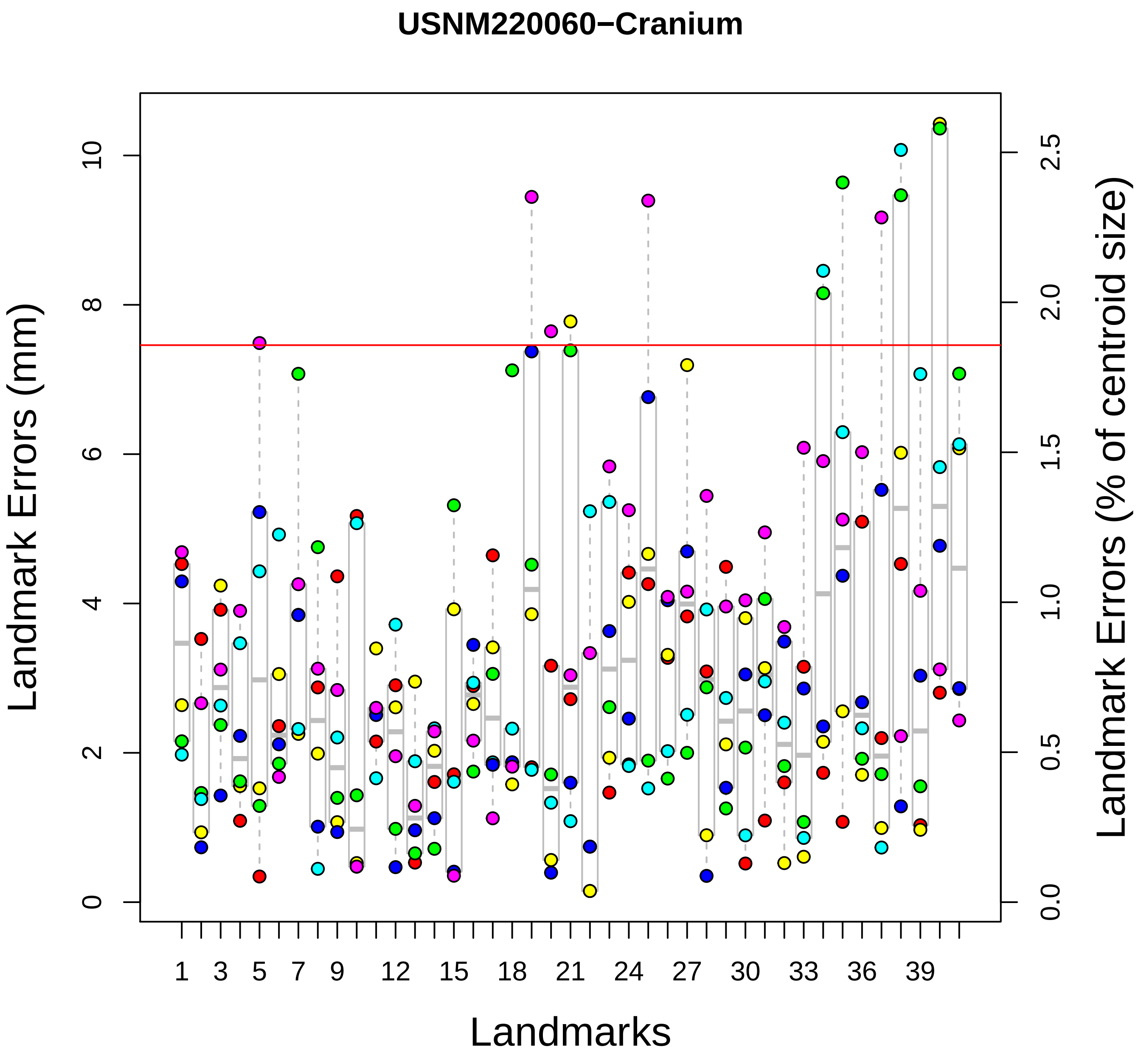

USNM220060−Cranium
Landmark Errors (mm)
Landmark Errors (% of centroid size)
Landmarks
0
2
4
6
8
10
0.0
0.5
1.0
1.5
2.0
2.5
1
3
5
7
9
12
15
18
21
24
27
30
33
36
39

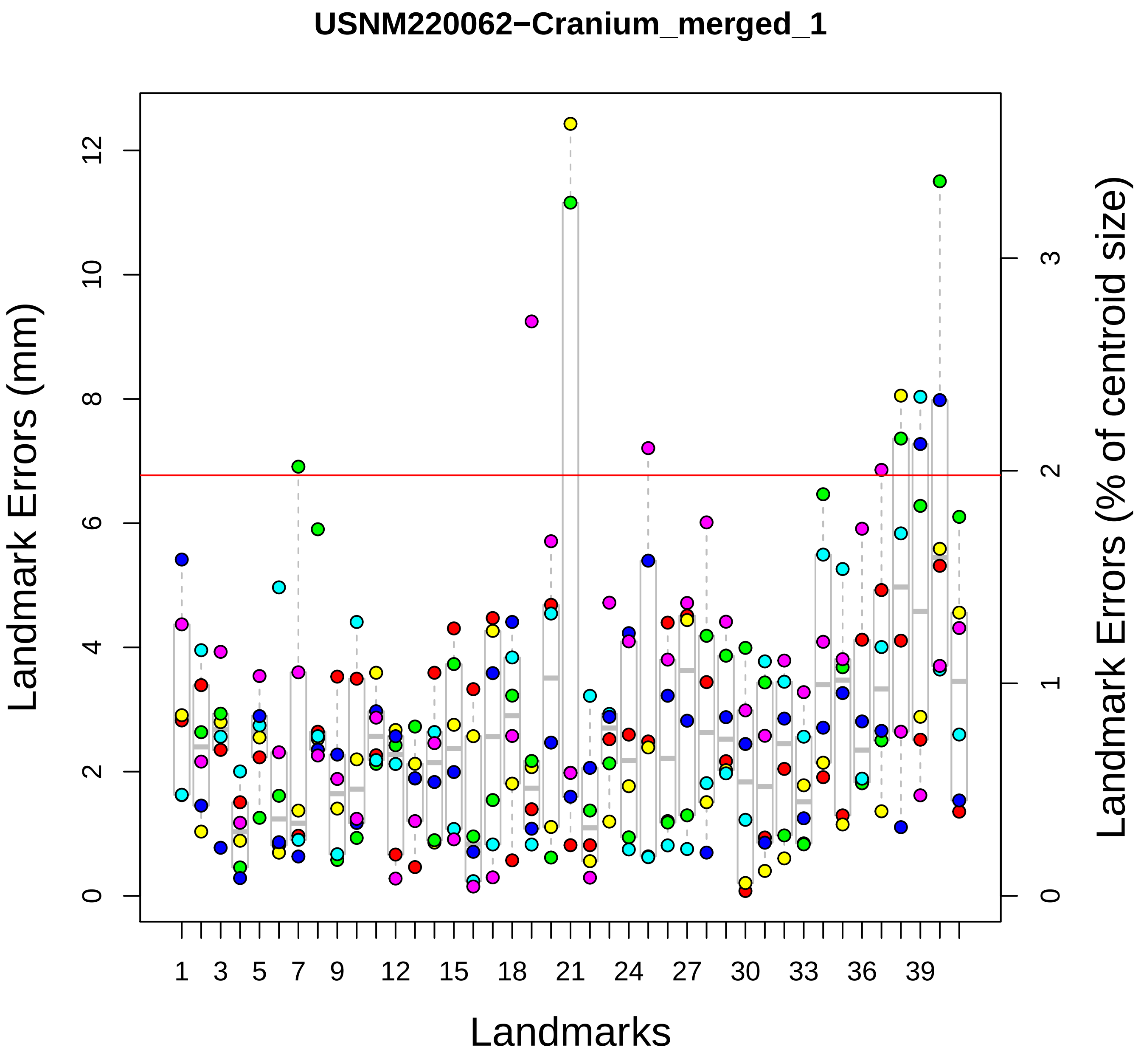

USNM220062−Cranium_merged_1
Landmark Errors (mm)
Landmark Errors (% of centroid size)
Landmarks
0
2
4
6
8
10
12
0
1
2
3
1
3
5
7
9
12
15
18
21
24
27
30
33
36
39

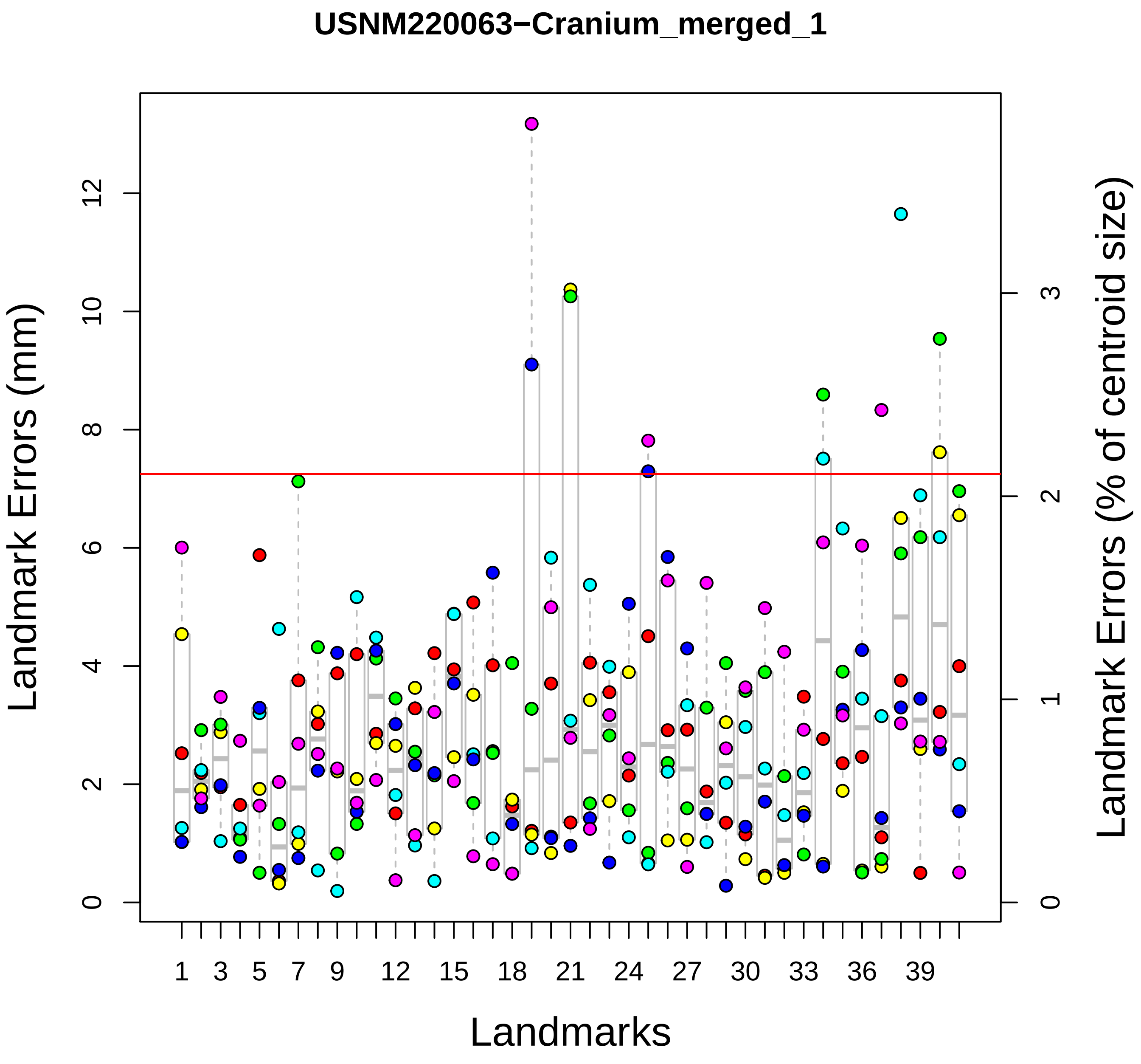

USNM220063−Cranium_merged_1
Landmark Errors (mm)
Landmark Errors (% of centroid size)
Landmarks
0
2
4
6
8
10
12
0
1
2
3
1
3
5
7
9
12
15
18
21
24
27
30
33
36
39

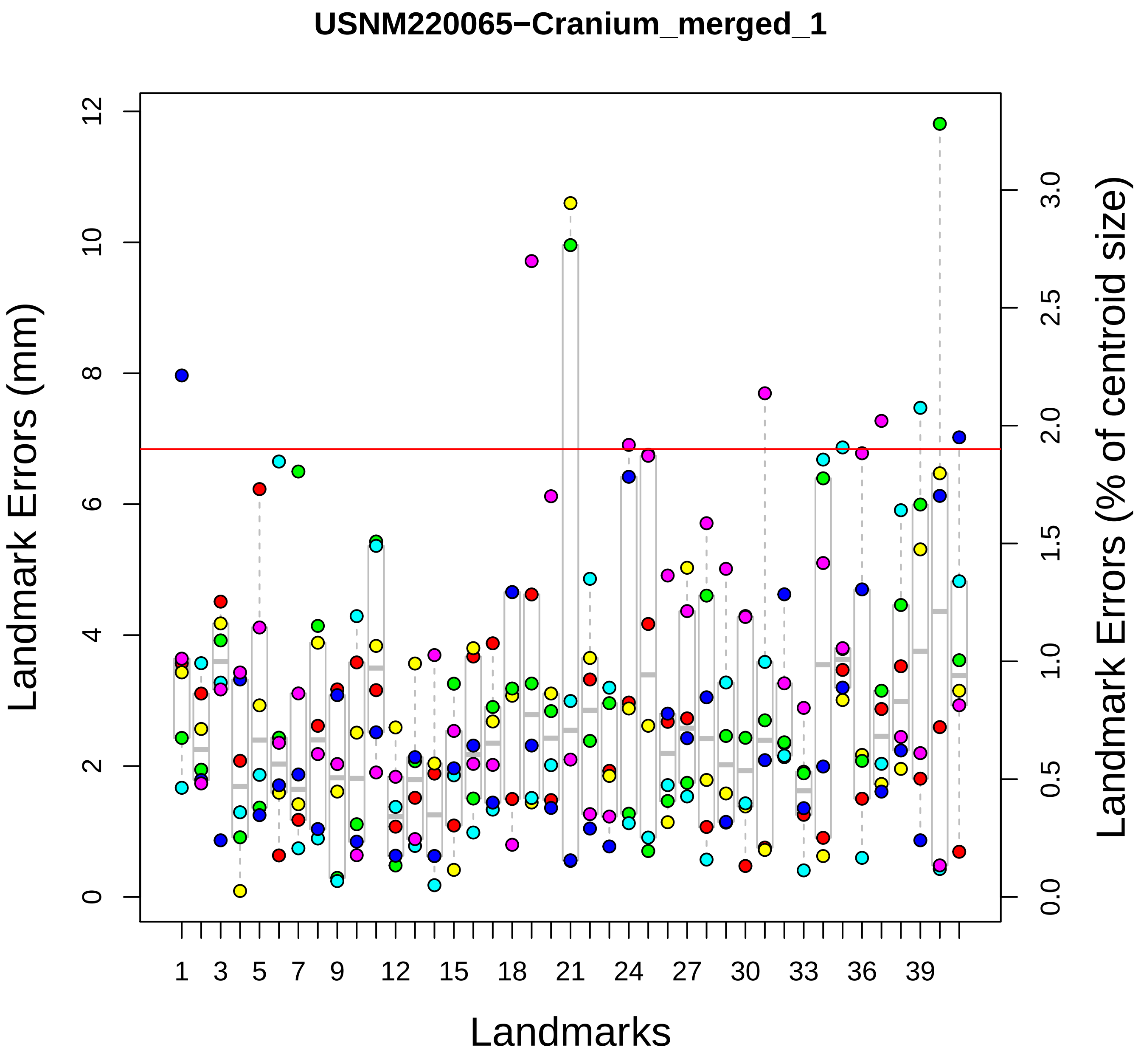

USNM220065−Cranium_merged_1
Landmark Errors (mm)
Landmark Errors (% of centroid size)
Landmarks
0
2
4
6
8
10
12
0.0
0.5
1.0
1.5
2.0
2.5
3.0
1
3
5
7
9
12
15
18
21
24
27
30
33
36
39

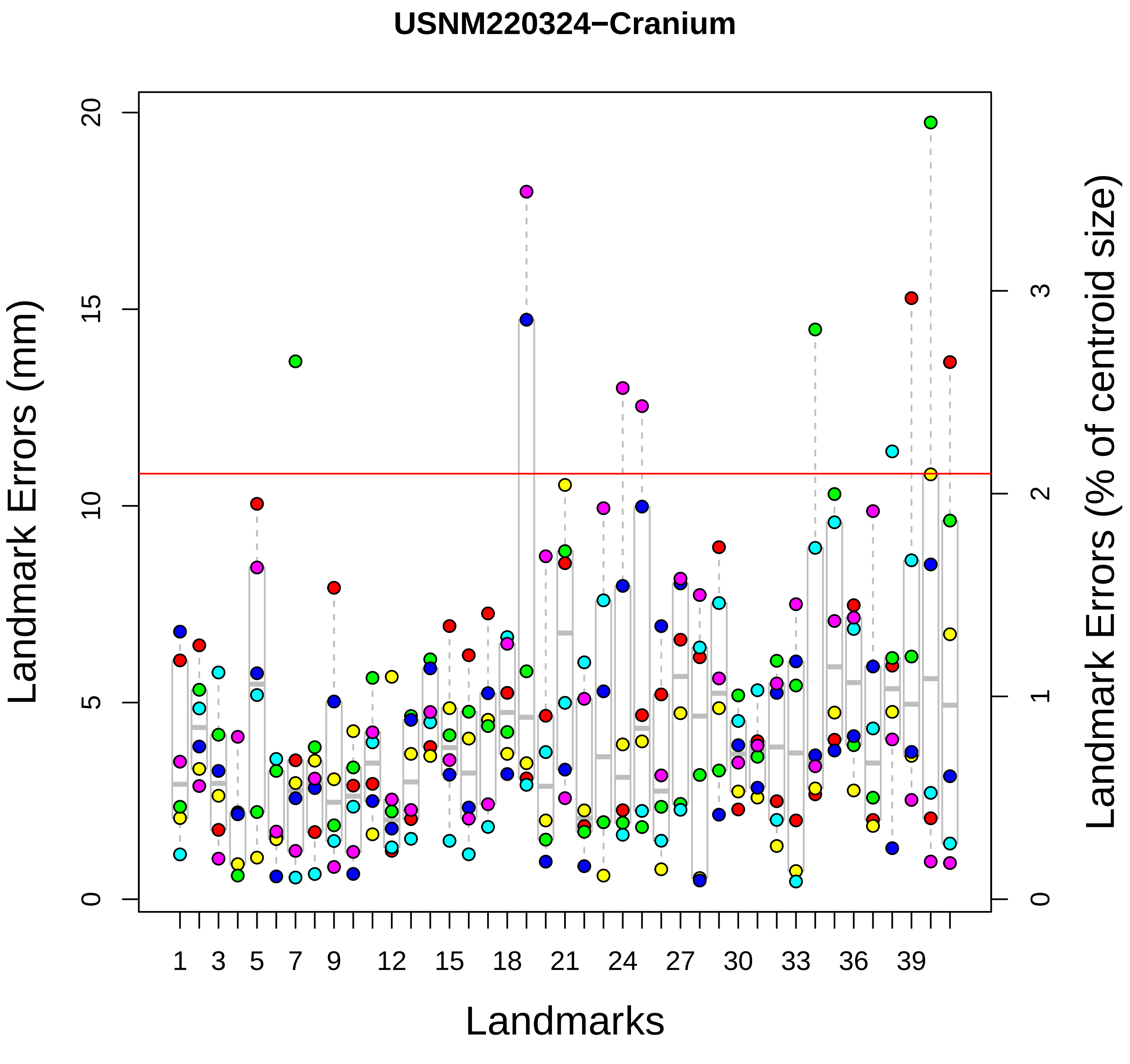

USNM220324−Cranium
Landmark Errors (mm)
Landmark Errors (% of centroid size)
Landmarks
0
5
10
15
20
0
1
2
3
1
3
5
7
9
12
15
18
21
24
27
30
33
36
39

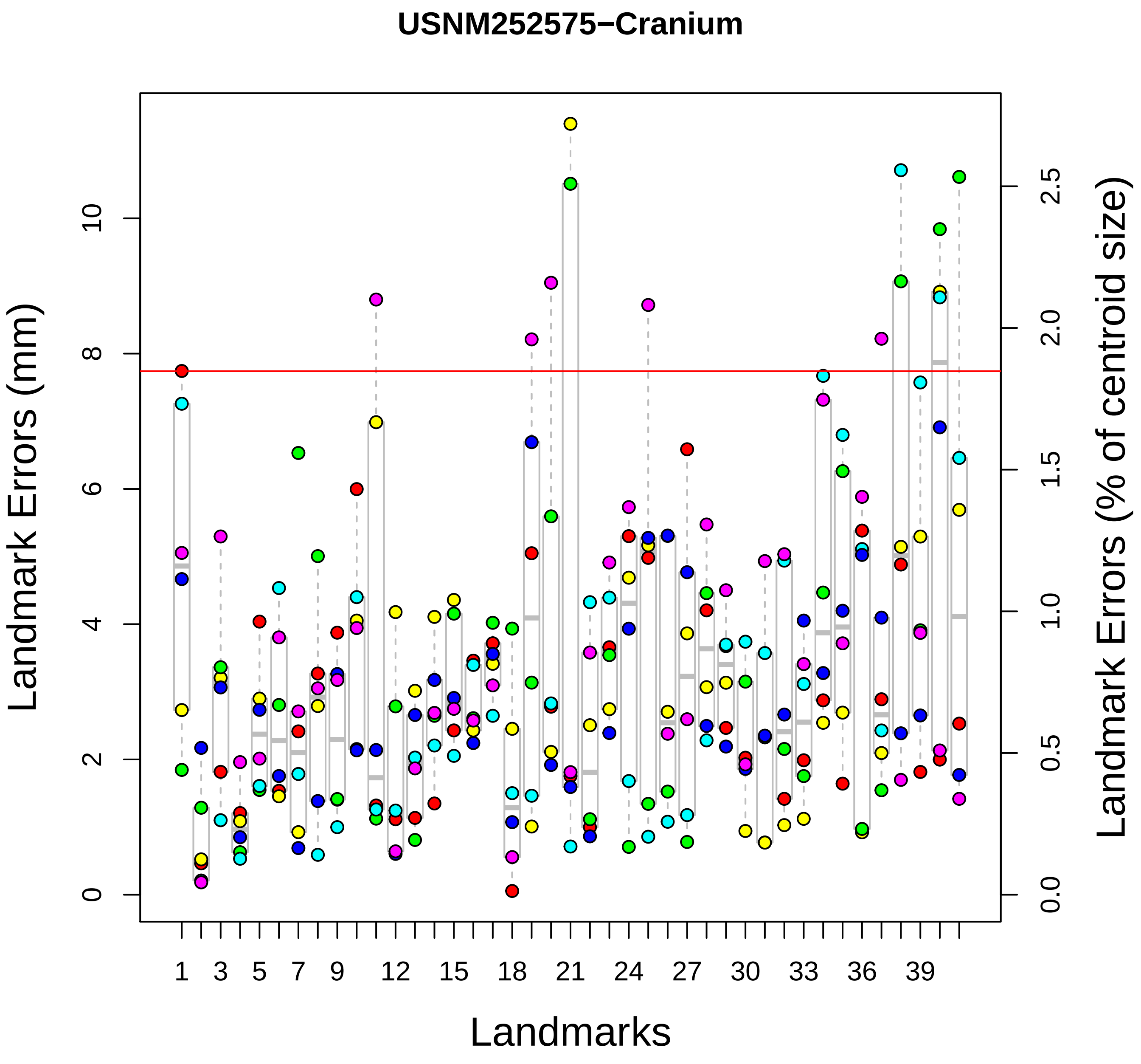
USNM252575−Cranium
Landmark Errors (mm)
Landmark Errors (% of centroid size)
Landmarks
0
2
4
6
8
10
0.0
0.5
1.0
1.5
2.0
2.5
1
3
5
7
9
12
15
18
21
24
27
30
33
36
39

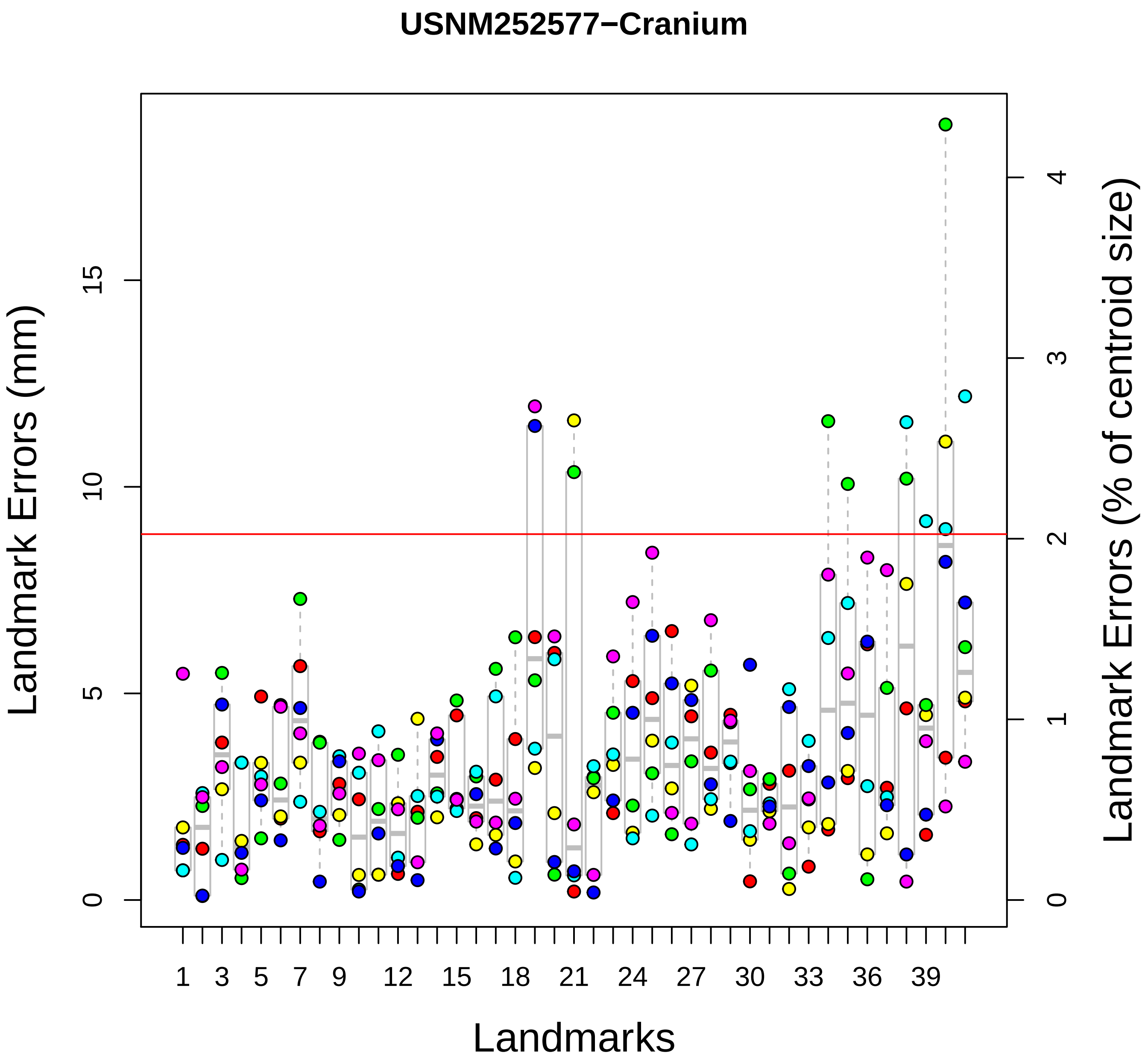

USNM252577−Cranium
Landmark Errors (mm)
Landmark Errors (% of centroid size)
Landmarks
0
5
10
15
0
1
2
3
4
1
3
5
7
9
12
15
18
21
24
27
30
33
36
39

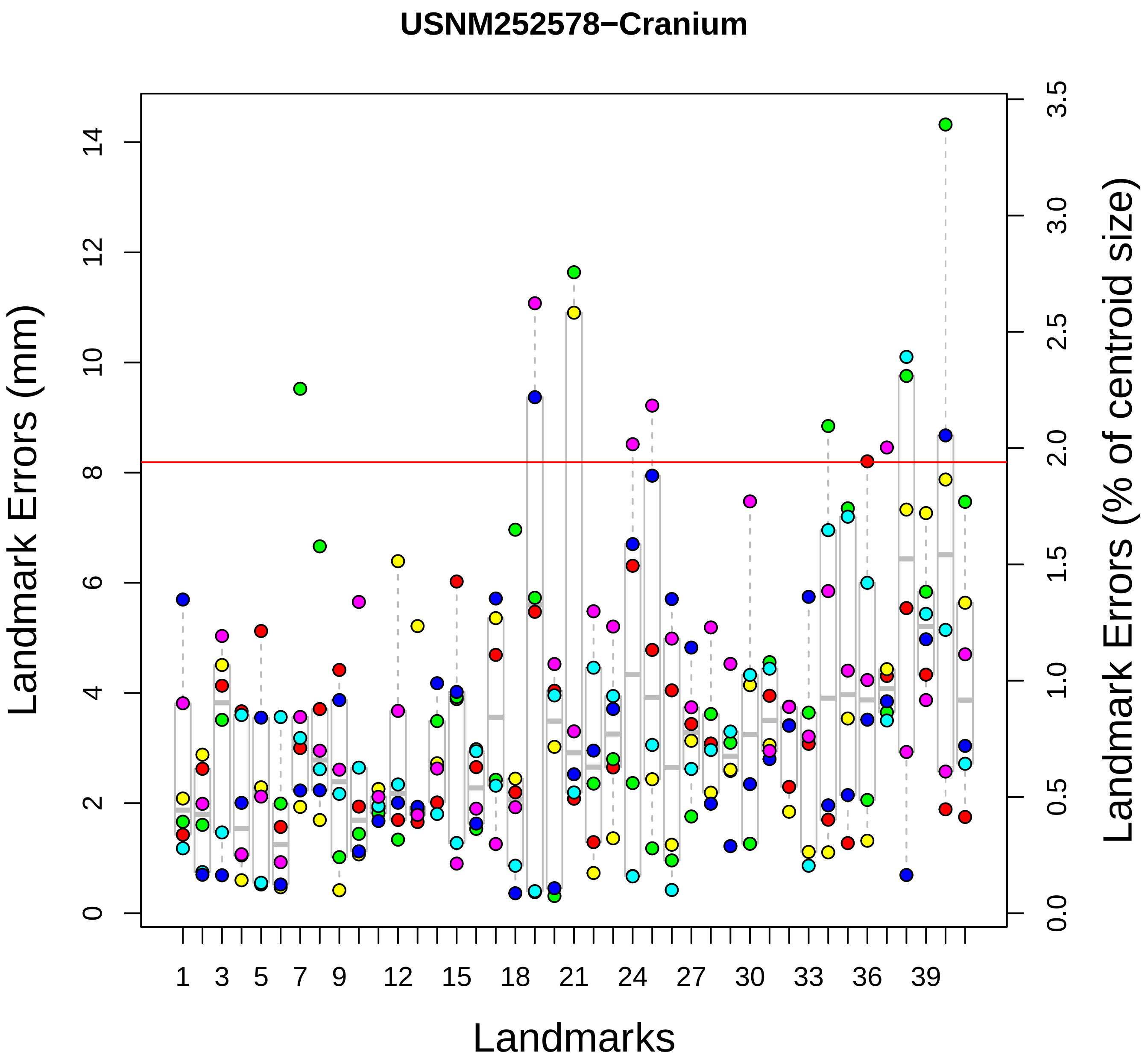
USNM252578−Cranium
Landmark Errors (mm)
Landmark Errors (% of centroid size)
Landmarks
0
2
4
6
8
10
12
14
0.0
0.5
1.0
1.5
2.0
2.5
3.0
3.5
1
3
5
7
9
12
15
18
21
24
27
30
33
36
39

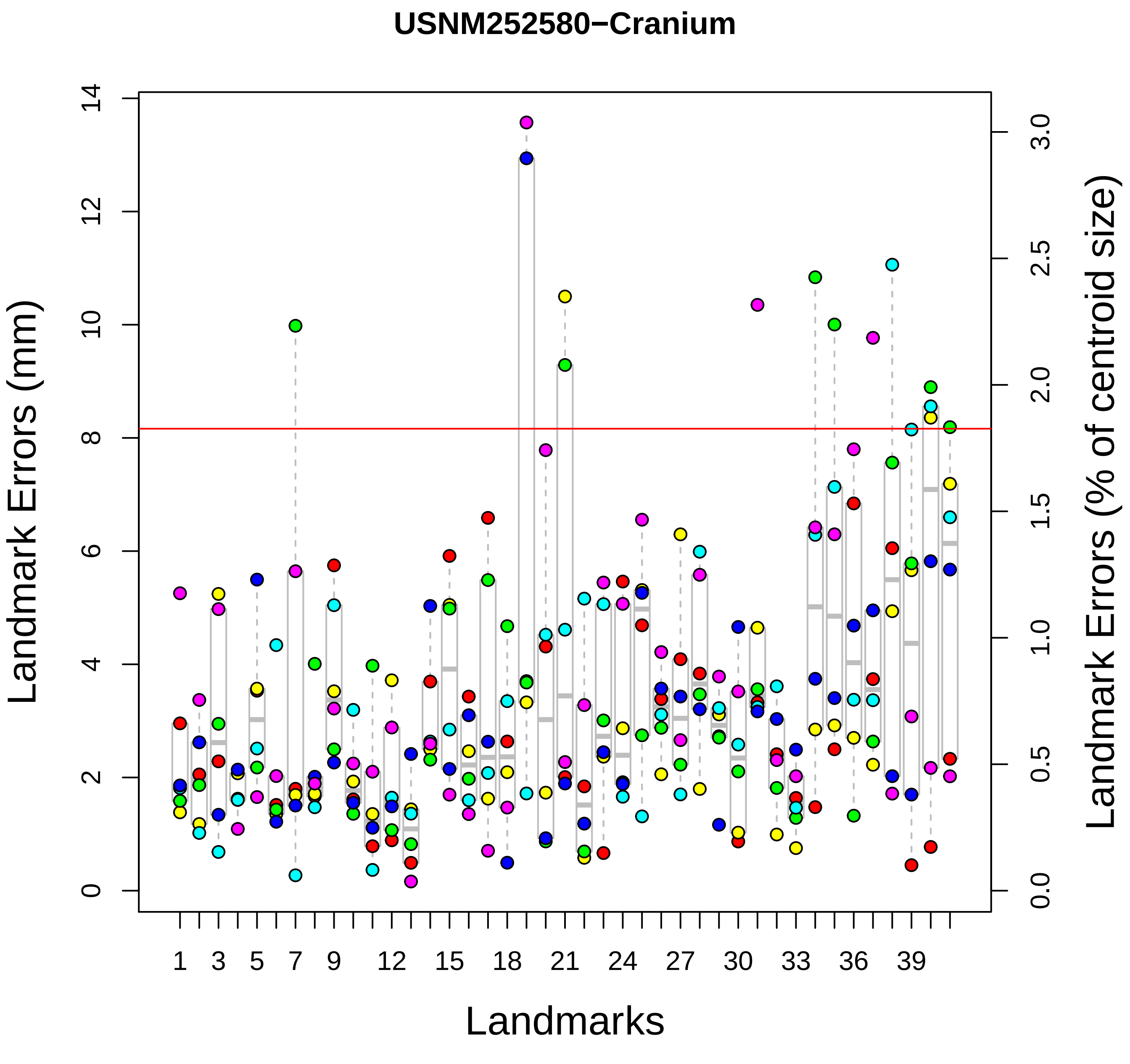
USNM252580−Cranium
Landmark Errors (mm)
0
2
4
6
8
10
12
14
Landmark Errors (% of centroid size)
0.0
0.5
1.0
1.5
2.0
2.5
3.0
Landmarks
1
3
5
7
9
12
15
18
21
24
27
30
33
36
39

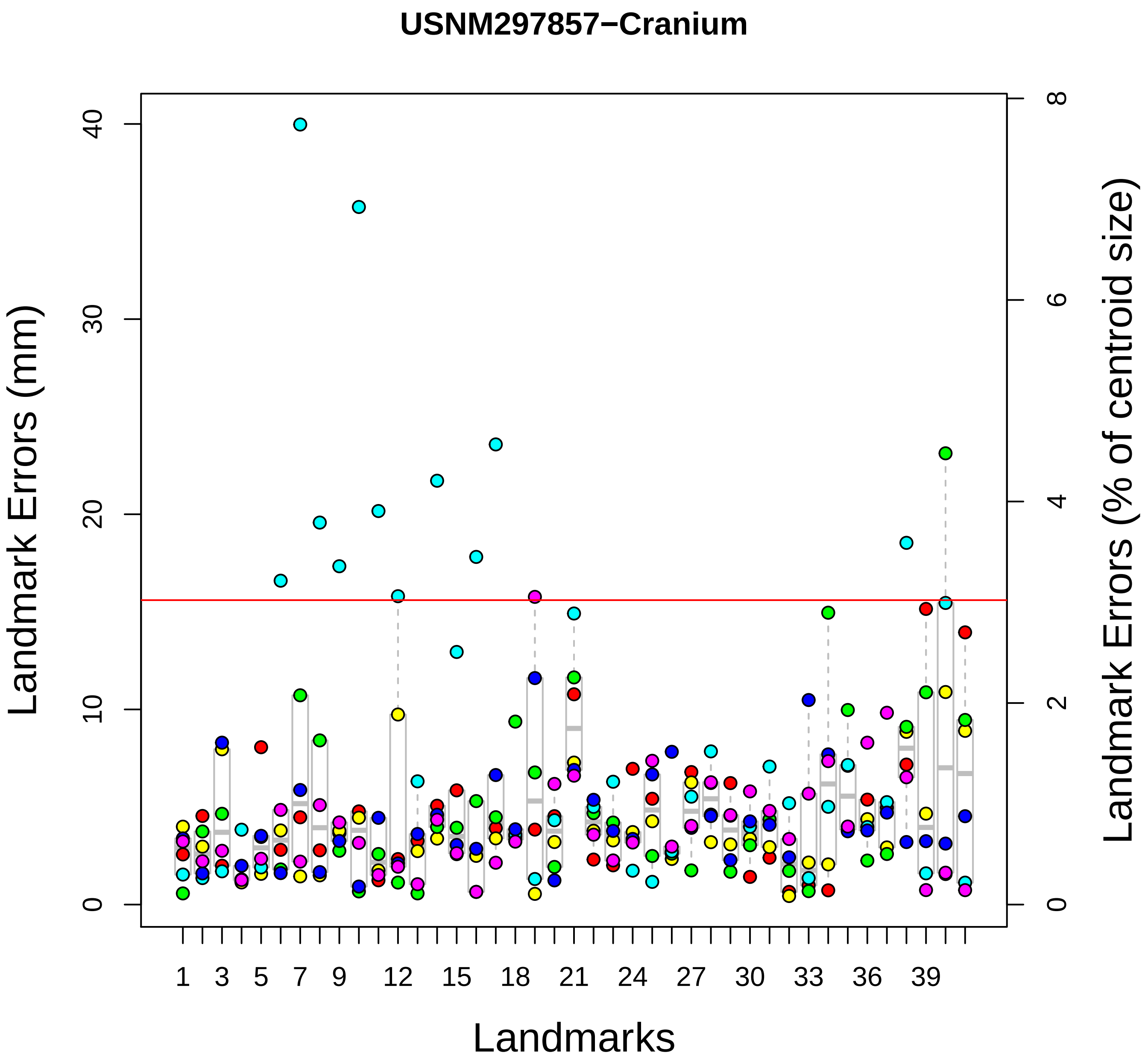

USNM297857−Cranium
Landmark Errors (mm)
Landmark Errors (% of centroid size)
Landmarks
0
10
20
30
40
0
2
4
6
8
1
3
5
7
9
12
15
18
21
24
27
30
33
36
39

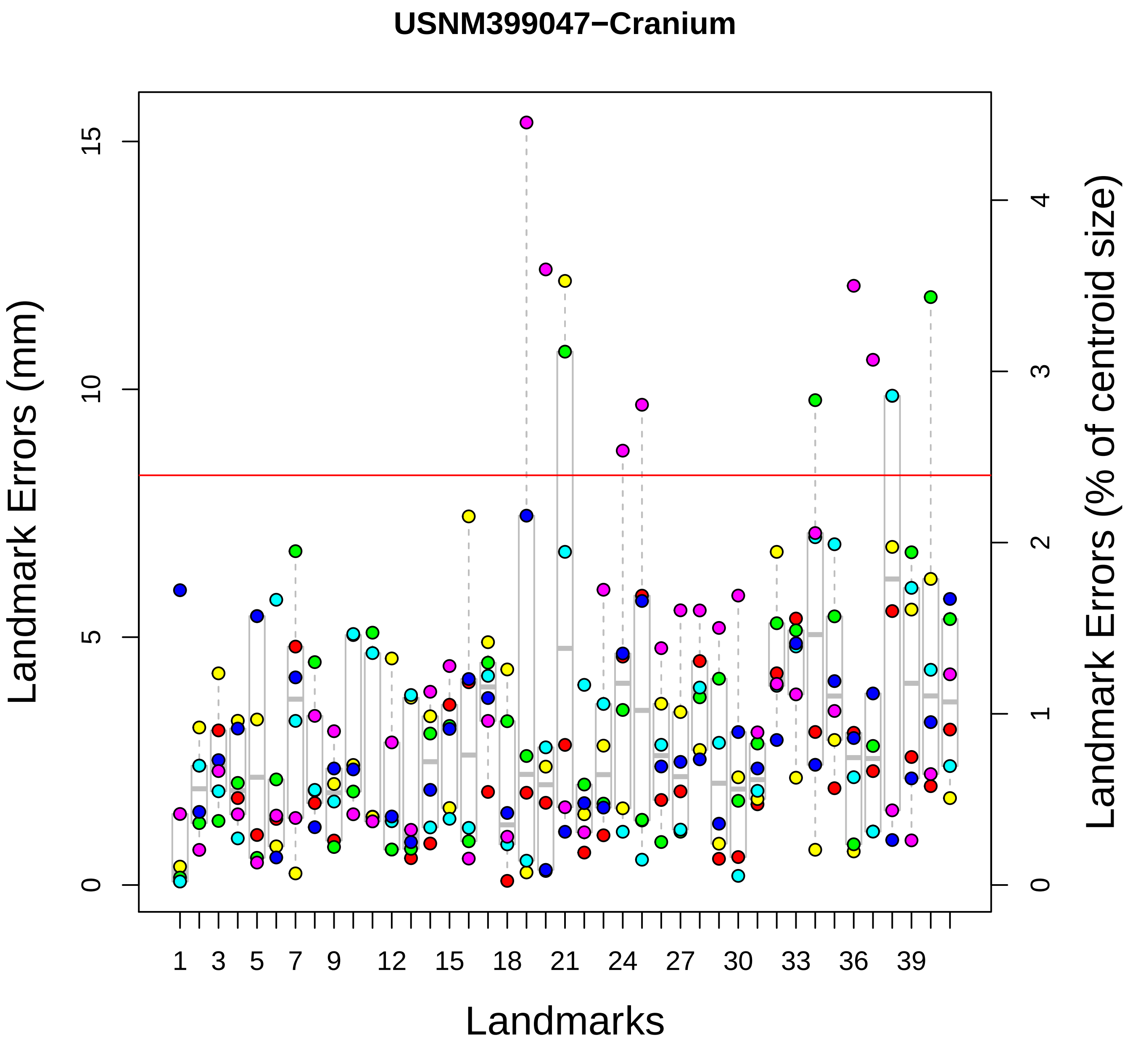
USNM399047−Cranium
Landmark Errors (mm)
Landmark Errors (% of centroid size)
Landmarks
0
5
10
15
0
1
2
3
4
1
3
5
7
9
12
15
18
21
24
27
30
33
36
39

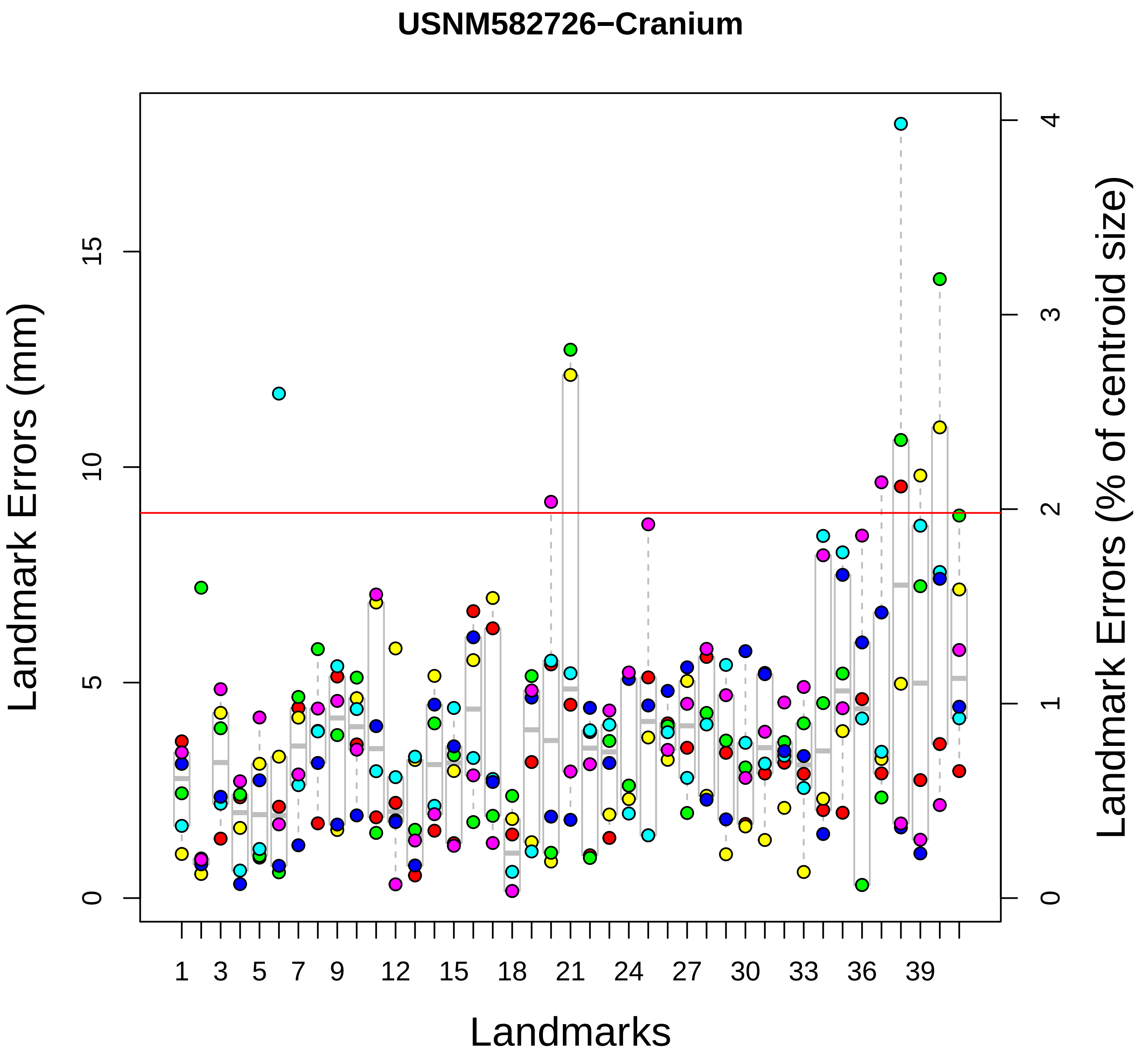
USNM582726−Cranium
Landmark Errors (mm)
Landmark Errors (% of centroid size)
Landmarks
0
5
10
15
0
1
2
3
4
1
3
5
7
9
12
15
18
21
24
27
30
33
36
39

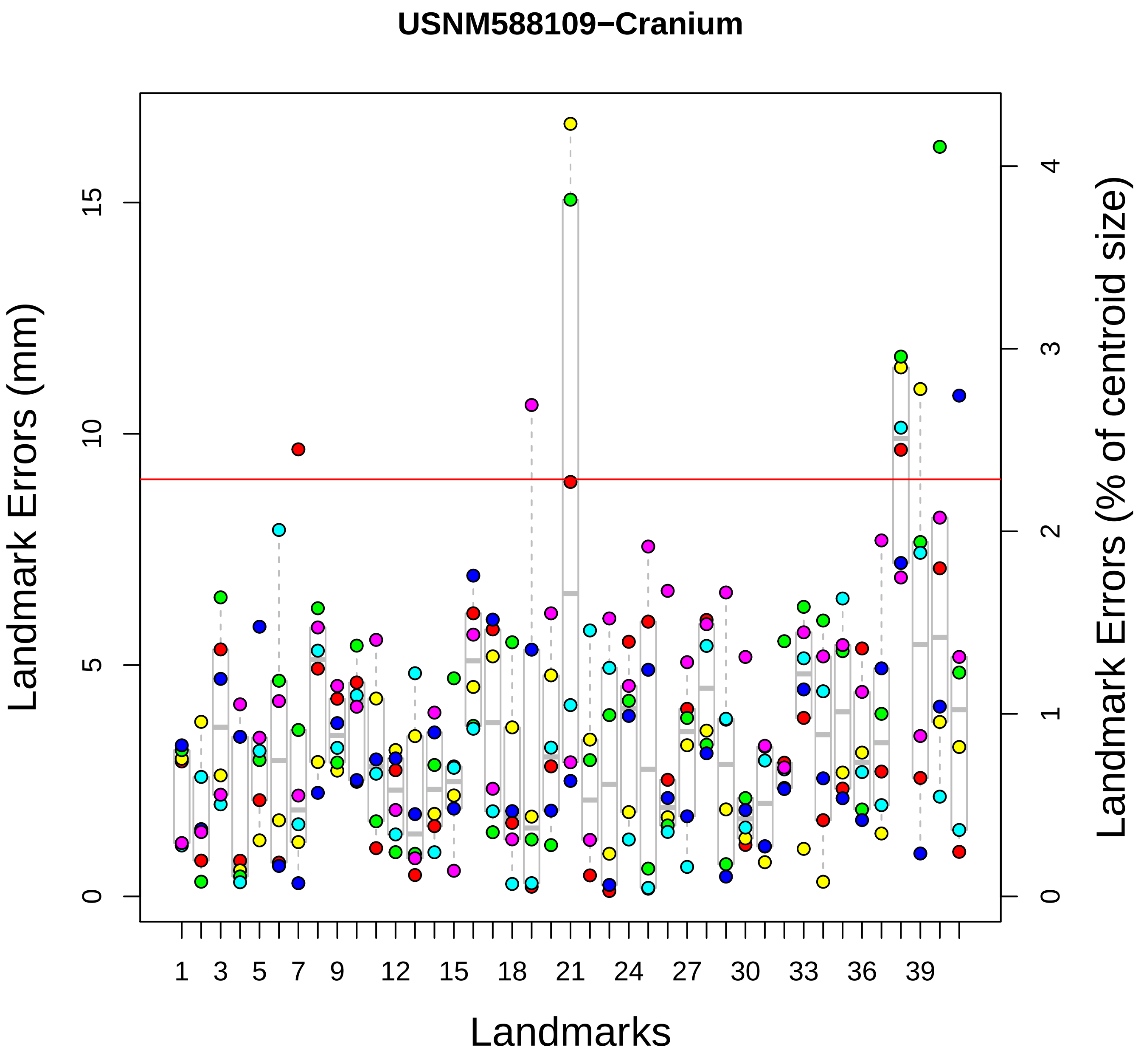
USNM588109−Cranium
Landmark Errors (mm)
Landmark Errors (% of centroid size)
Landmarks
0
5
10
15
0
1
2
3
4
1
3
5
7
9
12
15
18
21
24
27
30
33
36
39

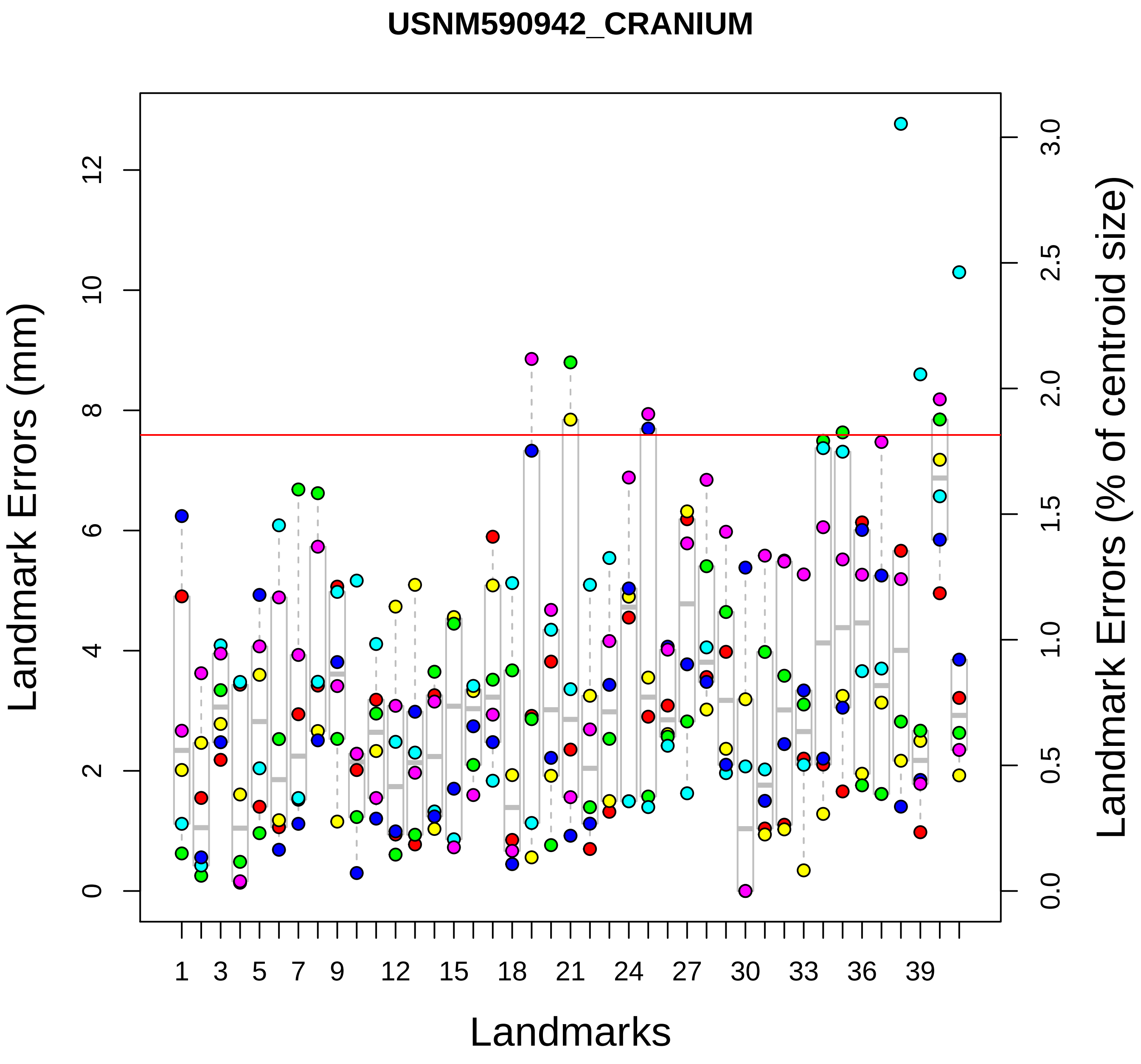

USNM590942_CRANIUM
Landmark Errors (mm)
Landmark Errors (% of centroid size)
Landmarks
0
2
4
6
8
10
12
0.0
0.5
1.0
1.5
2.0
2.5
3.0
1
3
5
7
9
12
15
18
21
24
27
30
33
36
39

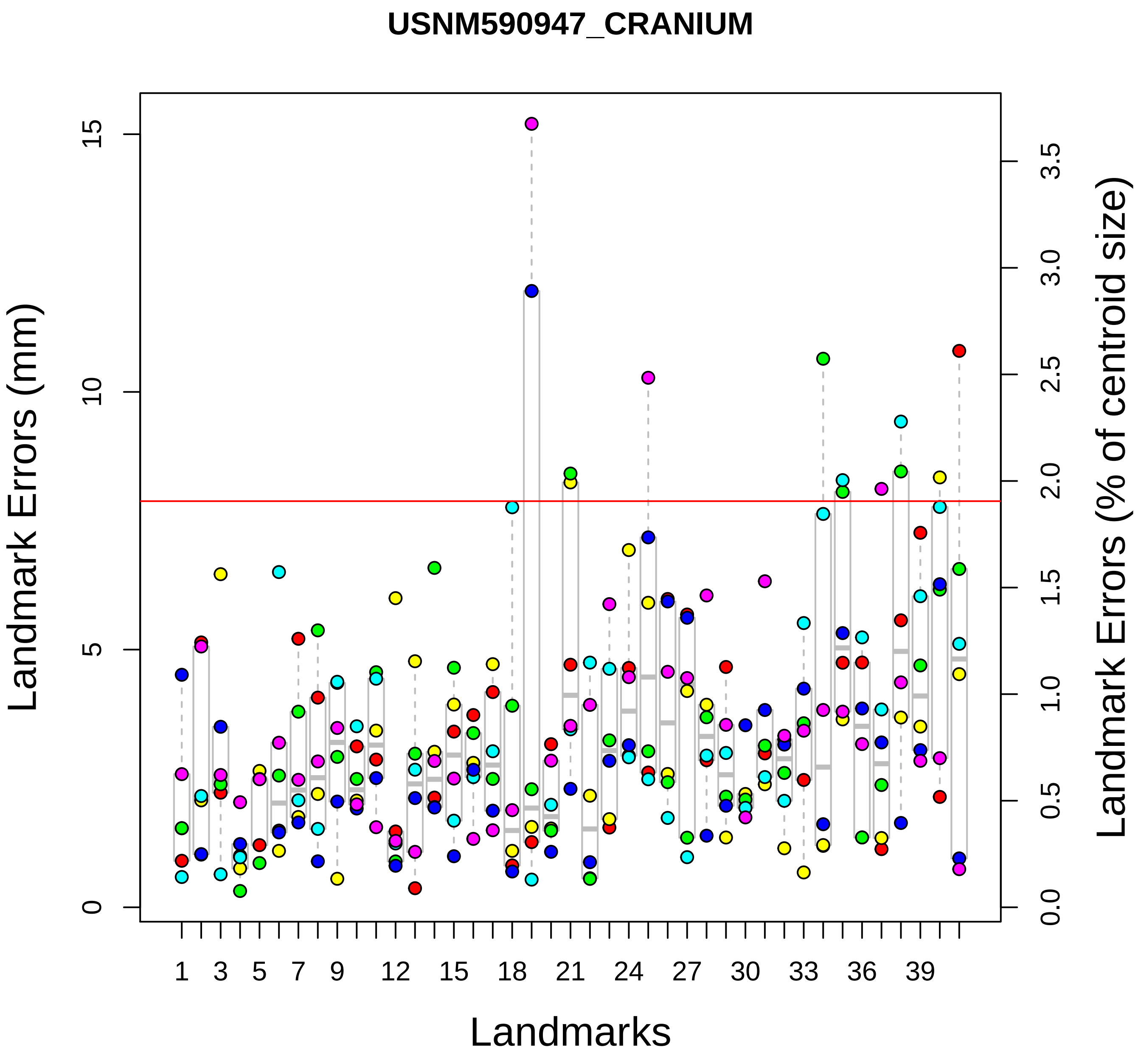
USNM590947_CRANIUM
Landmark Errors (mm)
Landmark Errors (% of centroid size)
Landmarks
0
5
10
15
0.0
0.5
1.0
1.5
2.0
2.5
3.0
3.5
1
3
5
7
9
12
15
18
21
24
27
30
33
36
39

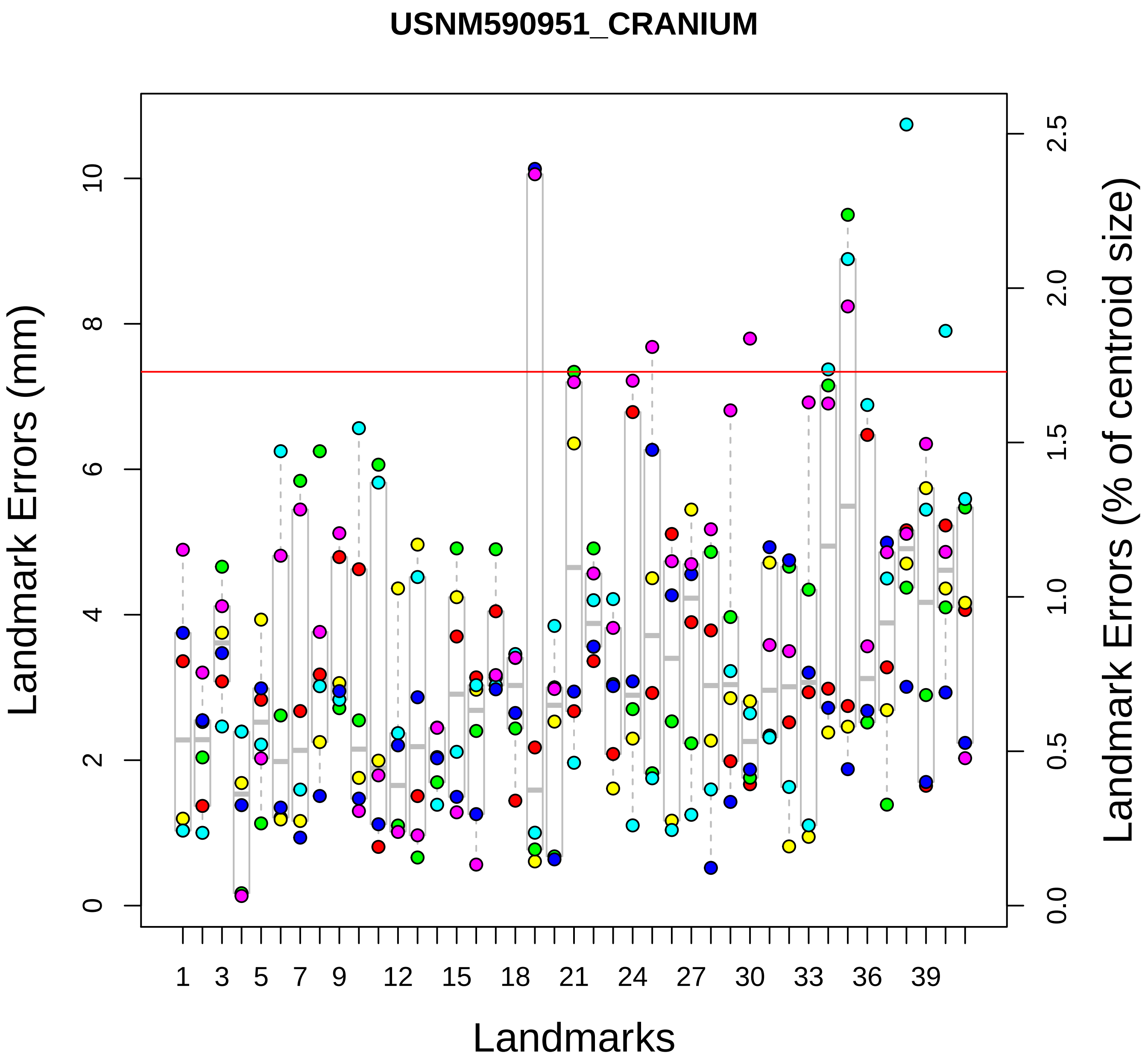
USNM590951_CRANIUM
Landmark Errors (mm)
Landmark Errors (% of centroid size)
Landmarks
0
2
4
6
8
10
0.0
0.5
1.0
1.5
2.0
2.5
1
3
5
7
9
12
15
18
21
24
27
30
33
36
39

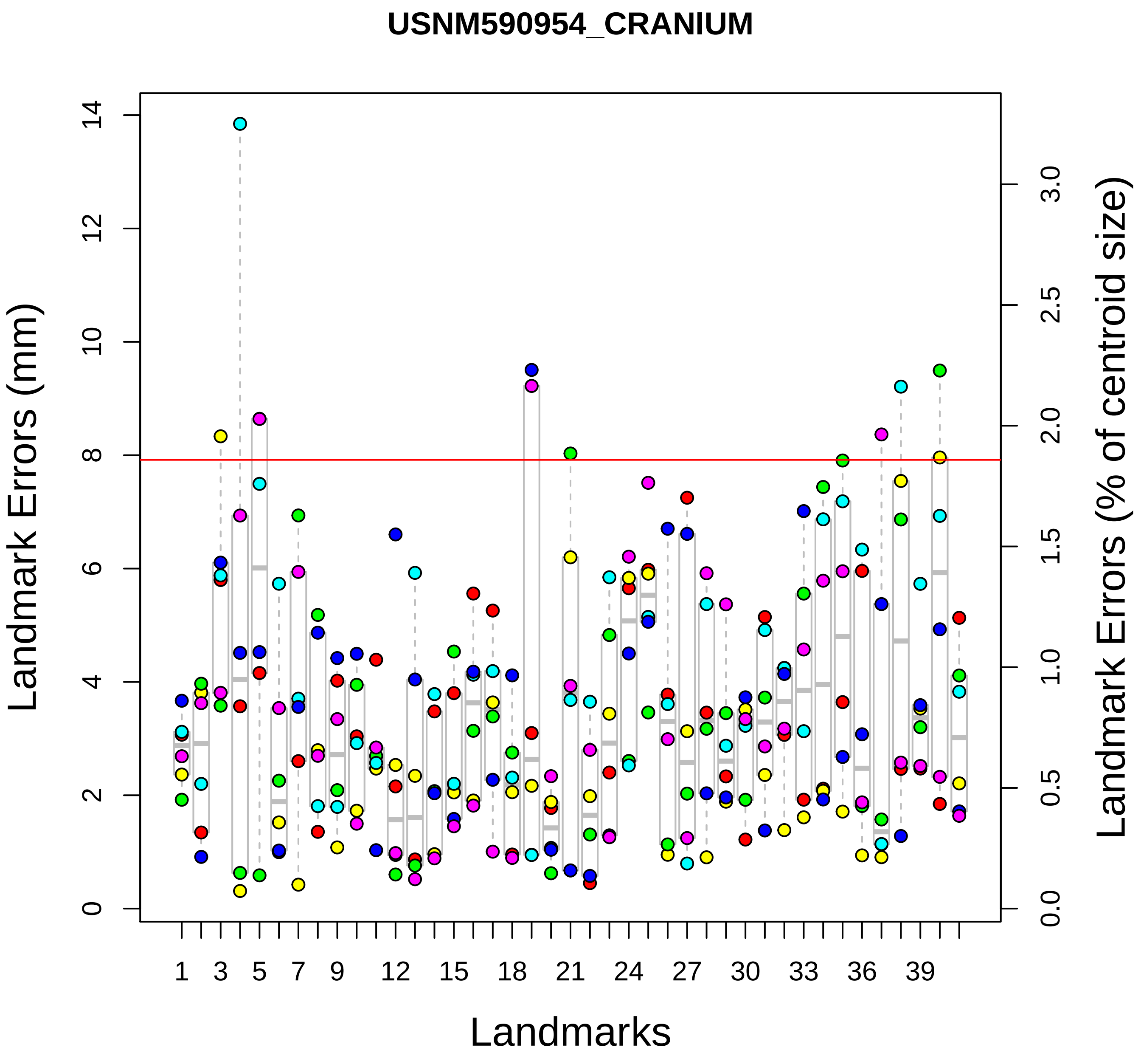
USNM590954_CRANIUM
Landmark Errors (mm)
Landmark Errors (% of centroid size)
Landmarks
0
2
4
6
8
10
12
14
0.0
0.5
1.0
1.5
2.0
2.5
3.0
1
3
5
7
9
12
15
18
21
24
27
30
33
36
39

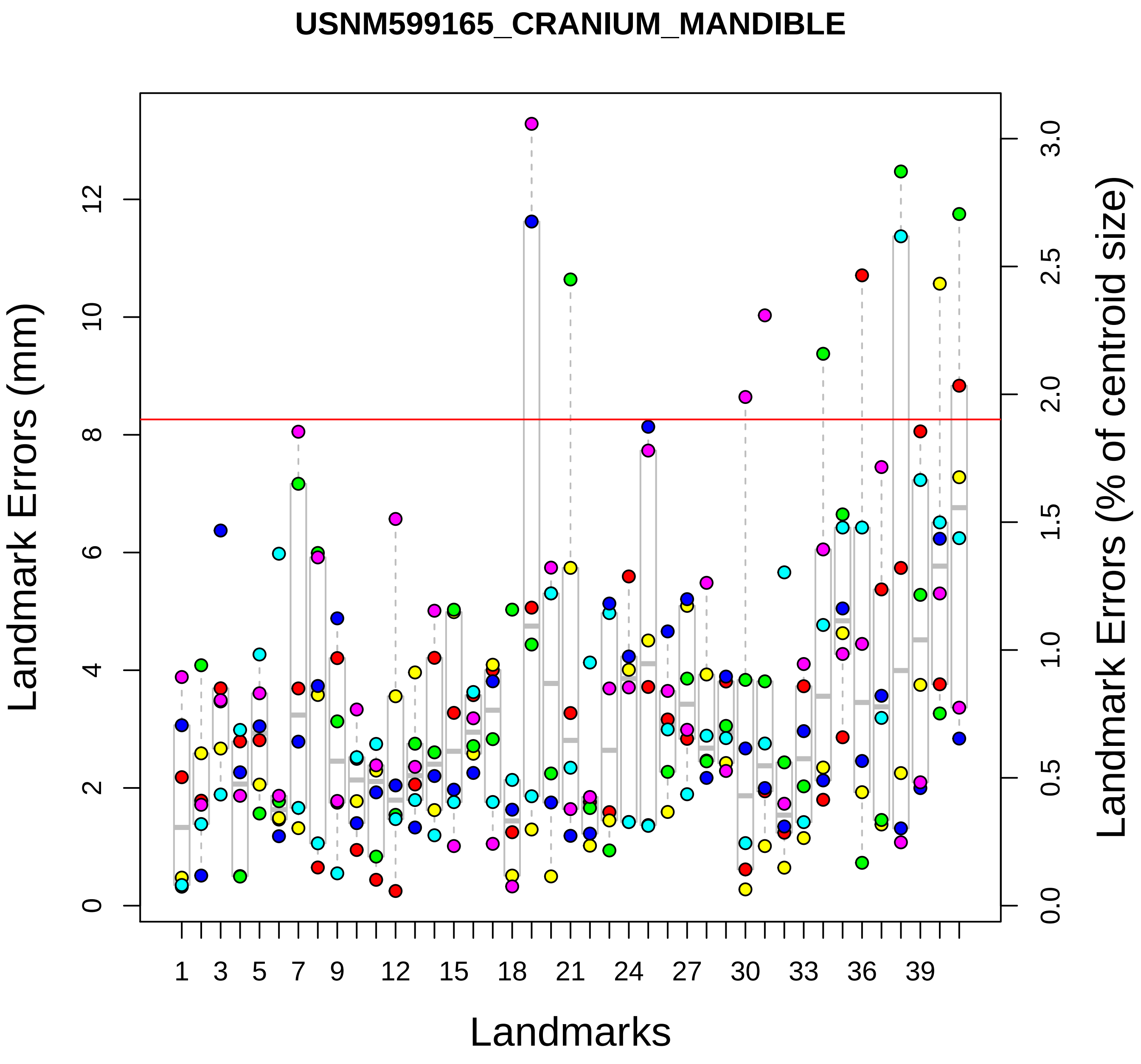

USNM599165_CRANIUM_MANDIBLE
Landmark Errors (mm)
Landmark Errors (% of centroid size)
Landmarks
0
2
4
6
8
10
12
0.0
0.5
1.0
1.5
2.0
2.5
3.0
1
3
5
7
9
12
15
18
21
24
27
30
33
36
39

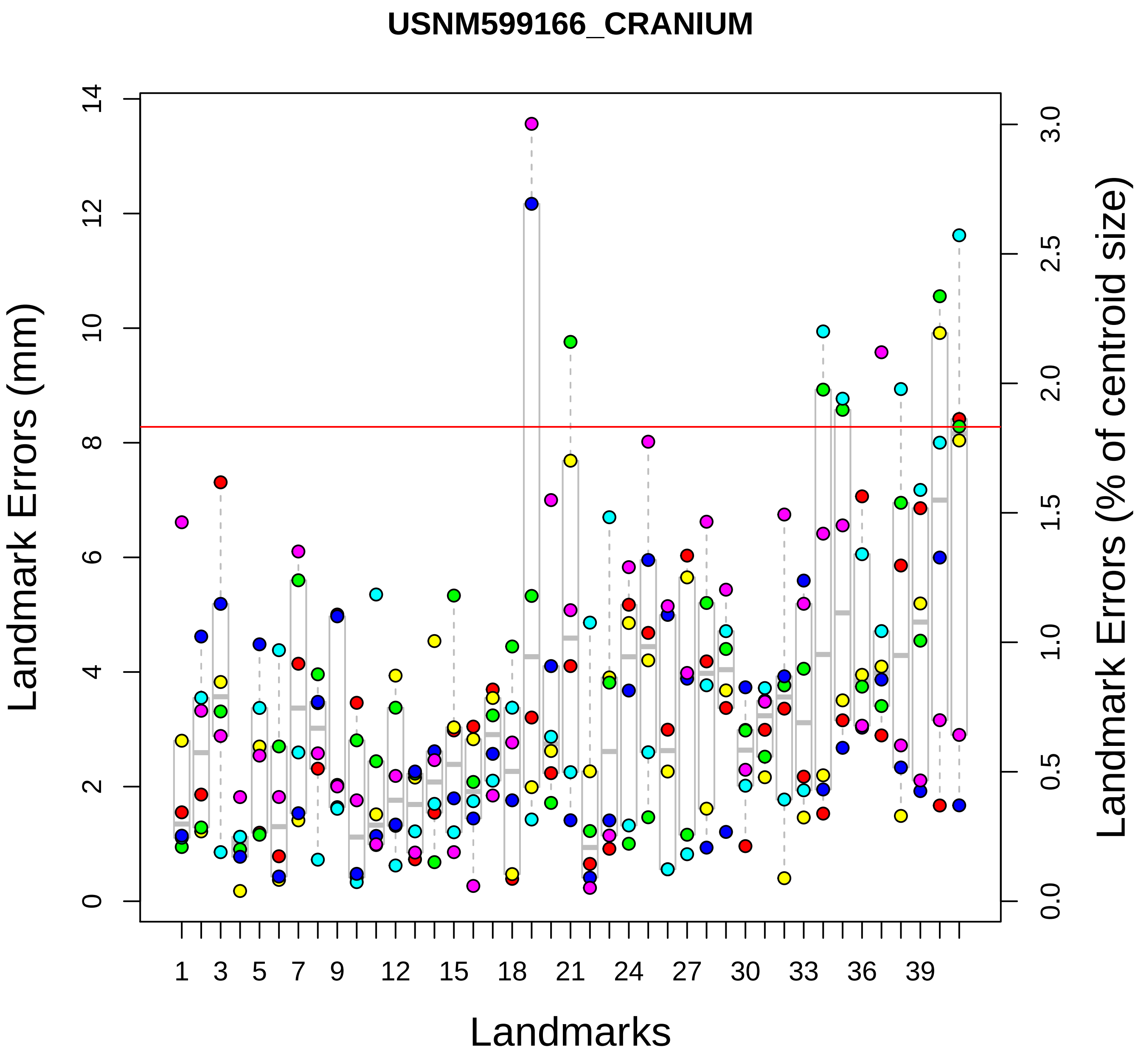
USNM599166_CRANIUM
Landmark Errors (mm)
Landmark Errors (% of centroid size)
Landmarks
0
2
4
6
8
10
12
14
0.0
0.5
1.0
1.5
2.0
2.5
3.0
1
3
5
7
9
12
15
18
21
24
27
30
33
36
39
